# Targeted delivery of mRNA to the heart via extracellular vesicles or lipid nanoparticles

**DOI:** 10.1101/2025.01.25.634881

**Authors:** Muhammad Nawaz, Benyapa Tangruksa, Sepideh Heydarkhan-Hagvall, Franziska Kohl, Hernán González-King Garibotti, Yujia Jing, Zahra Payandeh, Azadeh Reyahi, Karin Jennbacken, John Wiseman, Leif Hultin, Lennart Lindfors, Jane Synnergren, Hadi Valadi

**Affiliations:** Department of Rheumatology and Inflammation Research, Institute of Medicine, Sahlgrenska Academy, University of Gothenburg, Gothenburg, 41346, Sweden; Chief Medical Office, Global Patient Safety, BioPharmaceuticals R&D, AstraZeneca, Gaithersburg MD, USA; Systems Biology Research Center, School of Bioscience, University of Skövde, SE-541 28 Skövde, Sweden; Centre for Genomics Research, Discovery Sciences, BioPharmaceuticals R&D, AstraZeneca, Gothenburg 43183 Mölndal, Sweden; Department of Medical Biochemistry and Biophysics, Karolinska Institute, Solna, Stockholm, 171 77, Sweden; Bioscience Cardiovascular, Research and Early Development, Cardiovascular, Renal and Metabolism (CVRM), BioPharmaceuticals R&D, AstraZeneca, 431 83, Gothenburg, Mölndal, Sweden; Advanced Drug Delivery, Pharmaceutical Sciences, BioPharmaceuticals R&D, AstraZeneca, Gothenburg, 431 83 Mölndal, Sweden; Discovery Imaging, Clinical Pharmacology and Safety Sciences, BioPharmaceuticals R&D, AstraZeneca, Gothenburg, 431 83 Mölndal, Sweden; Department of Molecular and Clinical Medicine, Institute of Medicine, Sahlgrenska Academy, University of Gothenburg, Gothenburg, 41345, Sweden

**Keywords:** Targeted mRNA delivery, VEGF-A, systemic administration, lipid nanoparticles, extracellular vesicles

## Abstract

Targeted mRNA transport plays a crucial role in enhancing the therapeutic efficacy of the molecule, reducing its side effects, and minimizing off-target effects. Systemic administration of mRNA through lipid nanoparticles (LNPs) or extracellular vesicles (EVs) predominantly results in mRNA accumulation in the liver. We hypothesized that cardiac-specific EVs could more effectively target the transport of mRNA to the heart, in comparison to non-cardiac-specific EVs or LNPs. In mice, after intravenous administration, EVs from cardiac progenitor cells (CPC-EVs) were the most efficient to transport the modified mRNA, encoding vascular endothelial growth factor A (VEGF-A), to mouse heart, with minimal liver accumulation compared to non-cardiac-specific EVs or LNPs. Additionally, intracardiac injections of CPC-EVs not only demonstrate that they are the most adapted vehicle for interacting with heart tissue, delivering the mRNA to cells, and inducing maximal VEGF-A protein production, but RNA-seq analyses also revealed their minimal impact on overall gene expression, compared to LNPs or non-cardiac-specific EVs. Furthermore, immunofluorescence staining of CD31 and α-SMA, markers of microvascular density, showed increased vessel density in mouse aortic rings following the delivery of *VEGF-A* mRNA via CPC-EVs. These findings suggest that CPC-EVs are superior in mRNA targeting to heart, communication with cardiac cells, and causing minimal transcriptomic changes during *VEGF-A* mRNA delivery. Therefore, CPC-EVs could be promising vectors for heart-targeted mRNA delivery, potentially reducing liver accumulation.

## Introduction

Despite decades of dedicated research in RNA-based therapeutics, the scientific community still faces the challenge of developing vehicles that ensure the targeted delivery of RNA molecules to specific organ or cell types. Targeted delivery systems aim to maximize the therapeutic impact of drugs by ensuring their direct transport to the specific site of need, thereby enhancing efficacy, and reducing the required dosage ^1^. This focused drug delivery approach not only promises improved treatment outcomes but also diminishes the likelihood of unintended side effects ^2^.

Delivering mRNA molecules specifically to the heart poses a significant challenge; the delivery vehicle must effectively transport the mRNA to the targeted cardiac site, where it can be translated into a functional therapeutic protein. Conventional delivery vehicles, including lipid nanoparticles (LNPs) and extracellular vesicles (EVs), tend to accumulate in the liver, preventing therapeutic RNA molecules from reaching the intended organ during systemic administration ^3–6^. Therefore, targeting organs other than liver requires the development of organ- or cell-specific delivery systems.

The delivery of *VEGF-A* mRNA to cardiac tissue has emerged as a particularly promising approach for the treatment of cardiovascular diseases. This approach aims to induce neovascularization, or angiogenesis, a process with potential therapeutic benefits for patients with heart failure ^7, 8^. The therapeutic application of vascularization with VEGF-A protein has been studied for decades, both in experimental settings and in clinical trials ^9–11^. Pioneering work over the last decade has shown that chemically modified *VEGF-A* mRNA can achieve strong and temporary protein expression for several days in mouse models, larger animals, and even in initial human trials ^8, 12–17^. However, these first-in-human clinical studies relied on the direct intracardiac injections of un-encapsulated *VEGF-A* mRNA were applied without using delivery vehicles, e.g., LNPs. It has been long sought that in addition to minimally invasive localized mRNA administration, there is a critical need of investigating the mRNA administration via packing systems ^3^. However, the current RNA packaging systems have so far been designed for the efficient delivery of RNA molecules but with little success on cellular specificity. Additionally, several studies have reported that most of the systemically administered mRNAs using LNPs, or EVs are accumulated in the liver via circulatory system ^3–6^. EVs are a heterogeneous population of lipid bilayer vesicles, including exosomes and microvesicles, which are secreted by almost every cell type studied so far. They are detected in body fluids and conditioned culture media from living cells^18^. EVs play a pivotal role in intercellular communication ^19^ by transporting essential biomolecules such as RNA, lipids, and proteins between cells and organs ^20^.

In this study, we hypothesized that cardiac-specific EVs could target the transport of mRNA to the heart more effectively compared to non-cardiac-specific EVs or LNPs, as these EVs reflect properties, such as surface proteins, from their parental cells. Here, we compared EVs derived from cardiac progenitor cells (CPC-EVs), with both non-tissue-specific EVs and LNPs. The results suggest that CPC-EVs are a promising vehicle for transporting therapeutic mRNA to the heart with minimal liver accumulation. Additionally, we found that CPC-EVs were most effective in communicating with cardiac tissue regarding optimal production of the desired protein (VEGF-A) and the least alterations in gene expression, compared to other vehicles tested.

## Results

### Cellular uptake of LNPs carrying *VEGF-A* mRNA and translation in recipient cells

The cellular uptake analysis of DLin-MC3-DMA LNPs carrying *VEGF-A* mRNA was investigated in three different cell types, *in vitro*, including cardiac progenitor cells (CPCs), human umbilical vein endothelial cells (HUVECs) and a human lung epithelial cell line (HTBs). 24h post LNP uptake, the amount of *VEGF-A* mRNA was quantified in the lysates of recipient cells, and the levels of secreted VEGF-A protein were quantified in supernatants (**Figure 1A**). The results showed that significant amounts of *VEGF-A* mRNA were internalized by all three types of recipient cells and translated into VEGF-A protein (**Figure 1B-C**).

**Figure 1.**
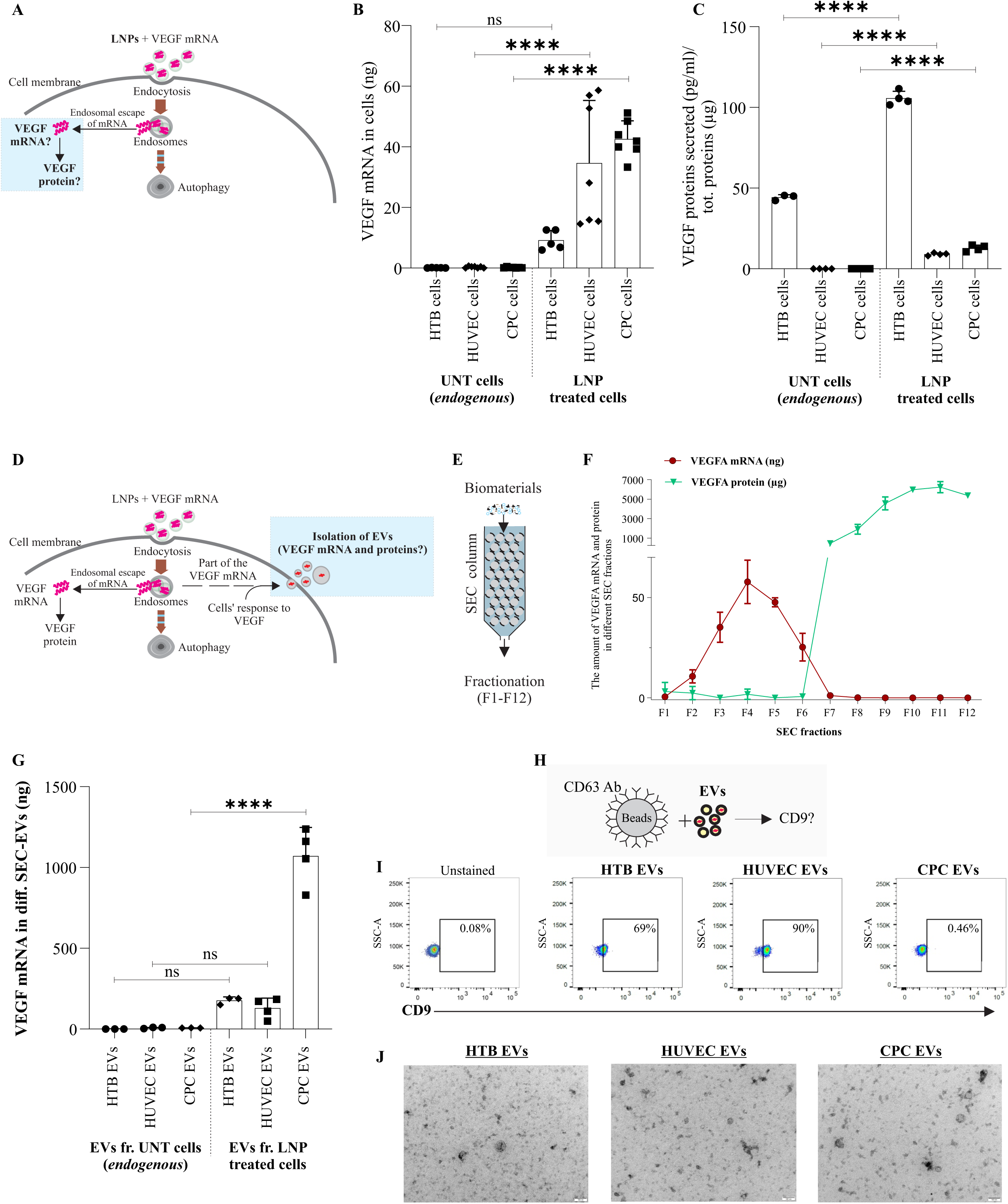
Uptake of LNP-*VEGF-A* mRNA by cells and loading into EVs, purification, and characterization. 3 µg of LNP-*VEGF-A* mRNA was administered to cardiac progenitor cells (CPCs), human umbilical vein endothelial cells (HUVECs) and a human lung epithelial cell line (HTBs). (**A**) A schematic illustration showing the uptake of LNP-mRNA by the recipient cells, escape from the endosomes for translation into protein. **(B)** Quantification of *VEGF-A* mRNA by qPCR, in the lysates of recipient cells (*n* = 6). **(C)** Quantification of VEGF-A protein by ELISA, in supernatants of recipient cells (*n* = 6). Statistically significant differences between LNP-treated and untreated samples were evaluated by applying Mann-Whitney U-test. Significant differences (\*\*\*\**p* < 0.0001) were observed between LNP-treated and untreated HUVECs and CPCs. No significant differences were observed between LNP-treated and untreated HTBs. UNT: untreated cells. 24h post LNP-mRNA uptake by cells, the conditioned media was collected, and the extracellular vesicles (EVs) secreted from these cells were isolated and purified by size exclusion chromatography (SEC), as described in the methods. **(D)** illustration showing the cellular route of mRNA loading into EVs and secretion. **(E)** Scheme showing the isolation of EVs by qEV/10 collum for size exclusion chromatography. **(F)** 12 fractions of SEC-EVs were collected and the presence of *VEGF-A* mRNA and protein in fractions of HTB-EVs, was determined by qPCR and ELISA, respectively. SEC-EVs carried substantial amount of exogenous *VEGF-A* mRNA, which was only possible to identify in fractions F1-6. **(G)** The total amounts of *VEGF-A* mRNA in SEC-EVs secreted from different cell types: HTBs, HUVECs, and CPCs. Statistically significant differences between LNP-treated and untreated samples were evaluated by applying Mann-Whitney U-test. Significant differences (\*\*\*\**p* < 0.0001). **(H)** Experimental scheme showing the immunocapture of SEC-EVs. SEC-EVs were first captured by anti-C63 antibody and then these CD63 positive EVs were further captured by CD9, conjugated with PE. **(I)** identification of EV markers; CD63 and CD9 in SEC-EVs. A 69 % of CD63^+^ HTB-EVs were also positive for CD9. A 90 % of CD63^+^ HUVEC-EVs of cells were also positive for CD9. CPC-EVs were positive for CD63, however only 0.56 % of CD63 EVs were positive for CD9. Unstained samples were used as negative control where no signals were detected. **(J)**. Transmission electron microscopy of HTB-EVs, HUVEC-EVs, and CPC-EVs. Scalebar: 100nm.

### Loading of chemically modified *VEGF-A* mRNA into extracellular vesicles

The components and chemical structure of DLin-MC3-DMA LNPs, applied for mRNA delivery in the current study, are provided in **supplementary Figure 1A**. The sequence of sense strand of chemically modified *VEGF-A* mRNA as well as sequence of the coded protein are provided in **supplementary Figure. 1B-C**. The *VEGF-A* mRNA was incorporated into EVs based on our previously established method ^21, 22^ **(Figure 1D)**. Using this method, the *VEGF-A* mRNA was loaded into three EV types secreted from CPCs (CPC-EVs), HUVECs (HUVEC-EVs) and HTBs (HTB-EVs). EVs loaded with *VEGF-A* mRNA were isolated and purified using size exclusion chromatography (SEC). The individual twelve SEC fractions of HTB-EVs were collected and analyzed for the presence of both *VEGF-A* mRNA and protein using qPCR and ELISA, respectively **(Figure 1E-F)**. The *VEGF-A* mRNA was present only in the first six fractions (F1-6), whereas the VEGF-A protein was detectable only in fractions 7 to 12 (**Figure 1F**). Additionally, after LNP-*VEGF-A mRNA* internalization, a time course experiment was performed to quantify the *VEGF-A mRNA* secretion over time (**supplementary Figure 2A-B)**. The results showed a continuous secretion of *VEGF-A mRNA into EVs,* whereby the peak was observed at 5h, with highest amounts in fraction 4. The amount of *VEGF-A* mRNA was also quantified in the individual fractions of HUVEC-EVs and CPC-EVs, which showed a significant amount of *VEGF-A* mRNA in the fractions of treated samples, compared to EVs secreted from untreated cells (unloaded EVs) (**supplementary Figure 2C-E**). Additionally, the *VEGF-A* mRNA was quantified in the pooled six fractions, showing significant amounts of the mRNA in total 6 fractions (**Figure 1G**).

### Characterization of extracellular vesicles

According to quality controls from qEV-70/10 mL columns (Izon Science Ltd, New Zealand) first 4 fractions (5 mL each) typically represent EV fractions. Since we loaded 15mL media instead of 10mL, we collected 2 extra fractions (*i.e.* 6 fractions) for EV characterization. The six SEC-EVs fractions were pooled, from all three cell types independently, and captured by anti-CD63 and anti-CD9 immunoaffinity-based method and detection by flowcytometry. The results showed that EVs which were positive for CD63, were also positive for CD9 for HTB-EV and HUVEC-EV samples. However, CPC-EVs which were positive for CD63, did not show signals for CD9 **(Figure 1H-I)**. The morphology of EVs obtained from three different cell types was examined by transmission electron microscopy and is presented in **(Figure 1J)**. Additionally, the diameters and concentrations of EVs were determined by nanoparticle tracking analysis (NTA), with mean size ranging between 149-158 nm, and mode size 146-152 nm **(supplementary Figure 3)**.

### EVs purified by size exclusion chromatography maintain their ability to deliver intact exogenous mRNA to cells

The efficacy of mRNA-loaded SEC-EVs in delivering exogenous mRNA into cells was initially determined *in vitro* (**Figure 2A**). First, we investigated the delivery of a functional mRNA, i.e., *VEGF-A* mRNA, using three SEC-EV types (HTB SEC-EVs, HUVEC SEC-EVs, and CPC SEC-EVs) to the recipient endothelial cells (HUVECs). The translation of *VEGF-A* mRNA was assessed in recipient cells, 24 hours post-delivery. Each of the three EV types successfully transported the mRNA into recipient cells, demonstrated by the production of VEGF-A protein in significant quantities when compared to untreated cells or cells treated with unloaded EVs (**Figure 2B**). The ability of SEC-EVs to deliver the intact mRNA across the cell membrane was further verified by a reporter gene. We investigated the cellular localization of EV-delivered mRNA and its translated protein in recipient cells. The Cy5-labelled *eGFP* mRNA loaded into HTB SEC-EVs was delivered to HTB-177 cells and the translation into eGFP was evaluated 24 hours after the uptake. The results showed that the EV-delivered mRNA and its protein (eGFP) were primarily localized in the cytosol of recipient cells (**Figure 2C**). These data suggest that SEC-EVs can transfer exogenous mRNAs into cells, which remains intact and results in the synthesis of a novel protein in the host cells.

**Figure 2.**
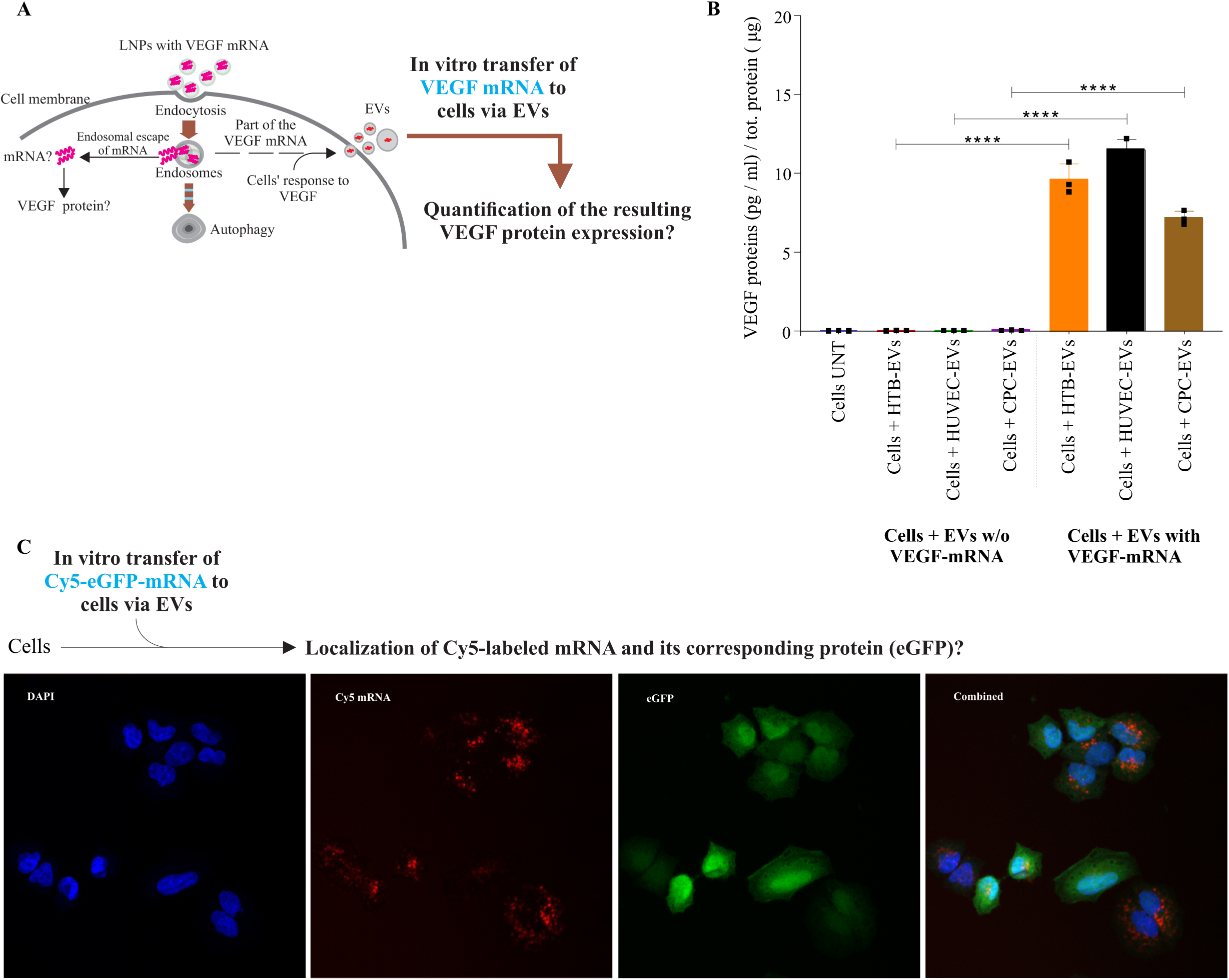
Delivery of *VEGF-A* mRNA to cells via SEC-EVs. **(A)** Schematic illustration of EV-mediated delivery to cells after loading from parent cells, which were initially exposed to LNPs. After isolation of EV-mRNA from LNP-mRNA treated cells, an equal amount of *VEGF-A* mRNA (550 ng) was delivered to HUVECs via three different EV types i.e., HTB SEC-EVs, HUVEC SEC-EVs and CPC SEC-EVs. The untreated cells and EVs which were not loaded with *VEGF-A* mRNA were used as controls. **(B)** Examination of translated *VEGF-A* mRNA after mRNA delivery via EVs. 24h post *VEGF-A* mRNA delivery *in vitro*, the supernatants from treated and untreated samples were collected and analyzed for *VEGF-A* protein using ELISA. Significantly higher levels of VEGF-A protein were detected in cells treated with *VEGF-A* mRNA via EVs. Whereases no VEGF-A protein was detected in untreated cells and the cells which were treated with EVs not carrying *VEGF-A* mRNA. One-Way ANOVA was applied to compare the statistical difference between treated (n=3) and control samples (n=3). ****p < 0.0001. **(C)** EVs containing Cy5-labelled translatable *eGFP* mRNA secreted from LNP-treated cells were delivered to recipient HTB-177 cells, *in vitro*. The cellular uptake of EV-Cy5-*eGFP* mRNA (red) and its translation into eGFP (green) were detected by confocal microscopy (n = 3). One representative image is shown. UNT: untreated, w/o: without

### Intravenous and intramuscular delivery of luciferase mRNA via SEC-EVs or LNPs

We investigated the efficiency of EVs in delivering a translatable exogenous mRNA via systemic route *in vivo* to different organs. A firefly luciferase mRNA (Fluc-mRNA) - a reporter gene, was administered to mice via HTB SEC-EVs or LNPs, and the biodistribution was analyzed (**Figure 3A**). Initially, the biodistribution of Fluc-mRNA was examined by intravenous delivery via EVs or LNPs. Although, LNPs showed a higher efficiency than HTB-EVs in transporting Fluc-mRNA to different tissues and in producing elevated levels of the luciferase enzyme, however, neither LNPs nor HTB-EVs showed specificity in targeted mRNA delivery (**Figure 3B-G**). Additionally, the uptake of mRNA by difficult-to-transfect cells was examined such as in muscle cells. The Fluc-mRNA was injected intramuscularly to examine the uptake of the mRNA by muscles via EVs or LNPs. *In vivo* imaging system (IVIS) measurements indicated that both EVs and LNPs were internalized by muscle cells, resulting in Fluc-mRNA translation into luciferase (**Figure 3H-J**).

**Figure 3.**
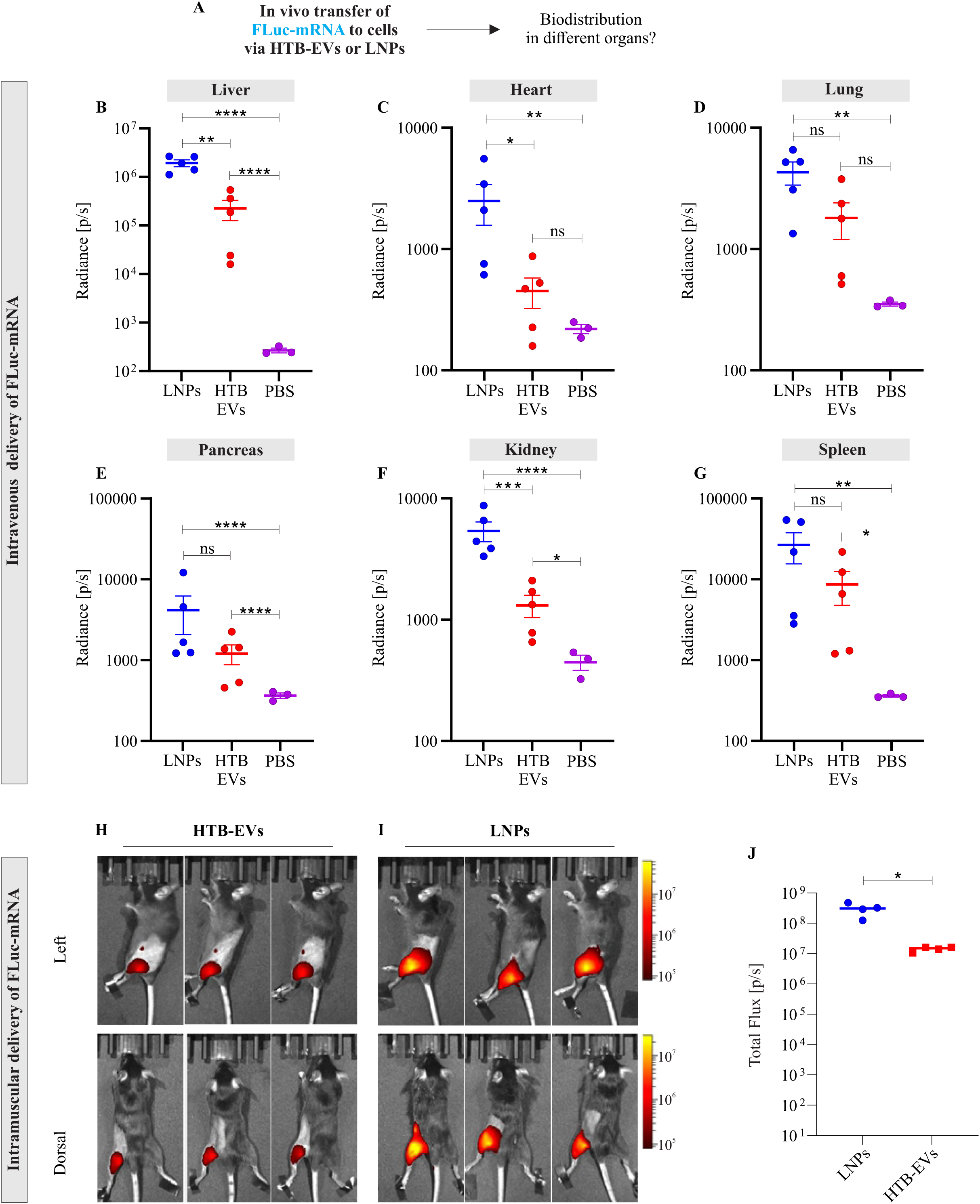
Intramuscular and intravenous delivery of luciferase mRNA via EVs or LNPs. **(A)** A schematic representing *in vivo* administration of mRNA via non-cardiac specific EVs (HTB-EVs) and examination of biodistribution. The mice were intravenously administered with EVs or LNPs containing firefly luciferase mRNA (FLuc-mRNA; 1 µg, in 100 µL). To confirm the translation of FLuc-mRNA into its protein i.e. luciferase, a luciferin (5 ml/kg) was administrated intravenously 6 h after the FLuc-mRNA delivery. The mice were terminated 20 minutes after the luciferin administration and the organs were dissected and scanned with an IVIS Spectrum within 5 minutes of termination. The total radiance was quantified as a factor of luciferase produced from FLuc-mRNA. Quantification of luciferase radiance in **(B)** liver, **(C)** heart, **(D)** lung, **(E)** pancreas, **(F)** kidney, and **(G)** spleen. One-Way ANOVA was applied to compare the statistical differences in radiance between untreated samples (n=3) with HTB-EVs or LNPs (n=3 each). *p < 0.05, **p < 0.01, ***p < 0.001, ****p < 0.0001. In a separate experiment, EVs or LNPs containing FLuc-mRNA (215 ng, in 30 µL) were administered intramuscularly to female C57bl/Ncr mice. The mice were sacrificed 6 h after Fluc-mRNA administration. After 20 minutes of luciferin administration the organs were dissected and scanned with an IVIS Spectrum within 5 minutes of termination. The radiance (luciferase-luciferin activity) was quantified in dorsal and left muscle after injections via **(H)** HTB-EVs, and **(I)** LNPs. **(J)** Total flux (radiance) of luciferase quantified after FLuc-mRNA delivery via HTB-EVs or LNPs. Nonparametric One-Way ANOVA followed by the Mann-Whitney U-test was applied to compare statistical differences between luciferase total flux after FLuc-mRNA delivery via HTB-EVs and LNPs. *p < 0.05. ns: non-significant.

### Cardiac progenitor cell derived EVs are the most efficient in tissue targeted delivery of mRNA

Previously, it has been shown that systemic administration of molecular therapies often leads to their accumulation in the liver through the circulatory system ^3–6, 23^. Therefore, the development of a delivery vehicle capable of bypassing the liver has garnered significant interest. In this context, our data demonstrate the unusual ability of cardiac-specific EVs (CPC-EVs) for liver bypass and the enhanced transport of exogenous *VEGF-A* mRNA specifically to the heart, in comparison to LNPs or non-cardiac-specific EVs.

The *VEGF-A* mRNA was intravenously administered to mice using LNPs and three different types of EVs such as cardiac specific CPC-EVs or non-cardiac-specific EVs such as HTB-EVs and HUVEC-EVs. The biodistribution of mRNA was examined across seven major organs, including the liver, heart, lung, pancreas, kidney, spleen, and thymus (**Figure 4A**). When a targeting efficacy for *VEGF-A* mRNA delivery was compared, the LNPs, HTB-EVs and HUVEC-EVs predominantly delivered the mRNA to the liver, and randomly to other studied organs such as lung, pancreas, spleen, thymus, and kidney, without exhibiting any specific targeting to a particular organ (**Figure 4B, D, F, and supplementary figures 4B, 5B**). However, CPCs showed an efficient targeted delivery of the same mRNA to the heart tissue (**Figure 4H**). Remarkably, among the total detected *VEGF-A* mRNA in all tissues, CPC-EVs delivered 56% specifically to the heart tissue, while LNPs, HTB-EVs, and HUVEC-EVs achieved only 19%, 8%, and 30%, respectively. Taking in view the fact that bypassing the liver-oriented delivery is an important aspect of targeted mRNA deliver, our data demonstrate that CPC-EVs transported a mere 11% of the detected *VEGF-A* mRNA to the liver, representing the lowest percentage compared to the other three vehicles. In contrast, the mRNA transported to liver by LNPs, HTB-EVs, and HUVEC-EVs was 37%, 41%, and 16%, respectively. This could, potentially, be of great importance in the development of targeted mRNA delivery therapies, as the ability to bypass the liver could allow for more efficient and specific delivery of therapeutic mRNA to the target tissues or organs.

**Figure 4.**
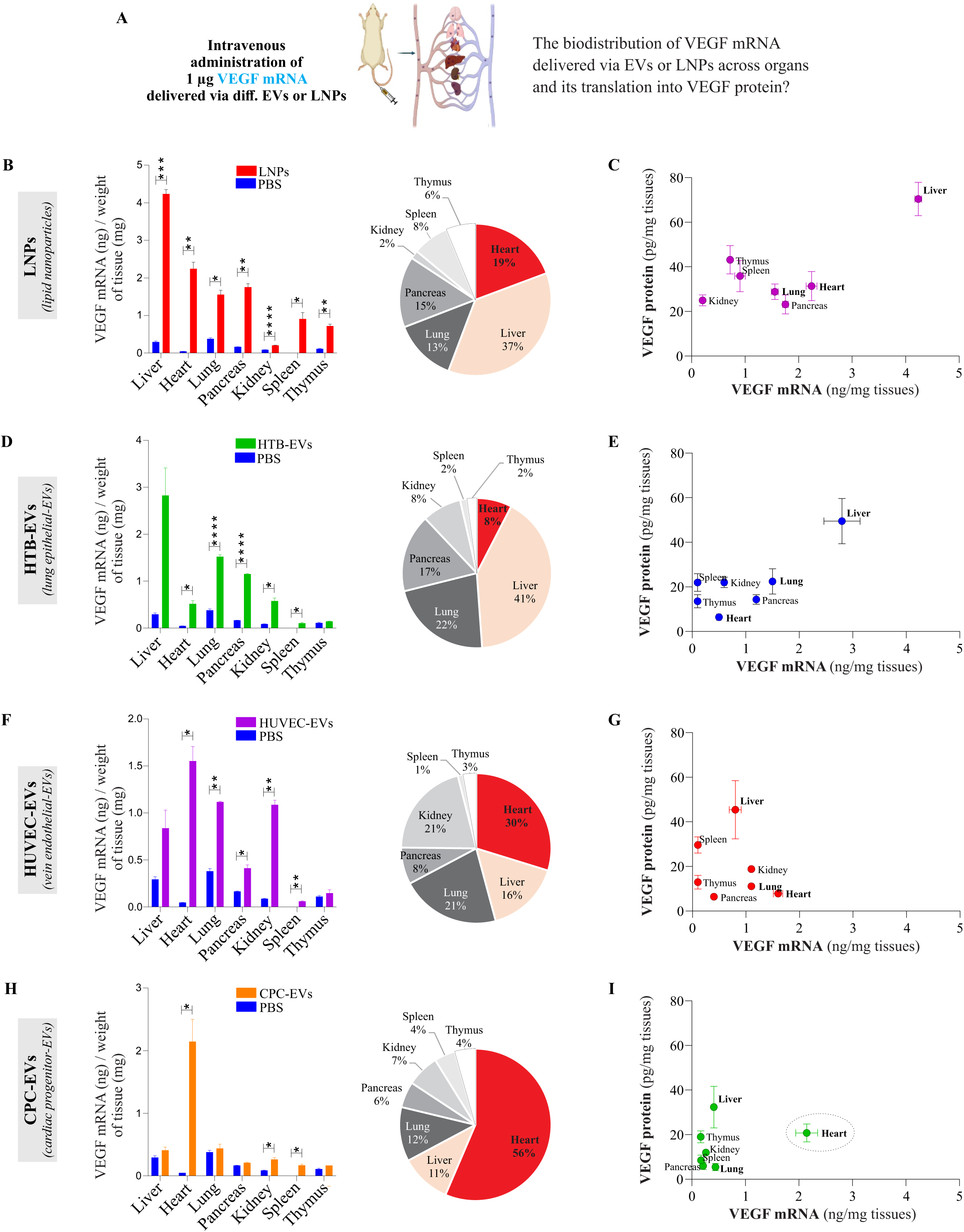
Organ targeted delivery of *VEGF-A* mRNA after intravenous delivery via SEC-EVs or LNPs. It is important to clarify that this is the amount of detected *VEGF-A* mRNA (in nanograms) per milligram of tissue from each organ. The detected amount of *VEGF-A* mRNA is not normalized against the total weight of each examined organ, due to the significant weight disparity among the organs; for example, the mouse liver weighs about 19 times more than the mouse heart**. (A)** Intravenous delivery of *VEGF-A* mRNA to C57BL/6Ncrl mice via cardiac-specific EVs or non-cardiac specific EVs or LNPs. 6h post administration, the mice were sacrificed, and the specificity of each vehicle for the delivery of *VEGF-A* mRNA was investigated in different organs (n=3). Relative percentage (%) distribution of *VEGF-A* mRNA and the *VEGF-A* mRNA amount (ng) in all organs delivered by **(B)** LNPs, **(D)** HTB-EVs, (**F**) HUVEC-EVs, and **(H)** CPC-EVs. The targeting specificity of each vehicle was compared by quantifying the *VEGF-A* mRNA and protein within the same organ and compared between other organs. (**C**) Targeting efficiency analysis showing LNPs have no targeting specificity, except for the known fact that LNP delivery is accumulated in the liver. (**E** and **G**) Targeting specificity of HTB-EVs and HUVEC-EVs shown by tissue-localization of *VEGF-A* mRNA and the levels of its protein. (**I)** Targeting efficiency of CPC-EVs to heart tissue for the localization of *VEGF-A* mRNA and the levels of its protein. The mRNA levels were normalized with tissue weight. One-Way ANOVA was applied to compare the statistical differences of *VEGF-A* mRNA between treated (n=3) and untreated (n=3) samples.

Additionally, we investigated the primary localization of the translated VEGF-A protein in mice tissues after intravenous administration. When delivery via LNPs or non-cardiac EVs or naked without vehicle, the *VEGF-A* mRNA, and its protein (VEGF-A) were primarily accumulated in the liver (**Figure 4C, E, G** and **supplementary figure 4**). In contrast, the results showed that *VEGF-A* mRNA and its protein were primarily detected in heart tissue when CPC-EVs were used as the delivery vehicle for intravenous administration (**Figure 4I**). These findings strongly suggest that CPC-EVs possess the potential to deliver a larger quantity of administered *VEGF-A* mRNA directly to the heart tissue, with minimizing accumulation in the liver, in comparison to LNPs or HTB-EVs, or HUVEC-EVs.

To further evaluate the specific delivery to the heart compared to other organs, we analyzed the ratio of *VEGF-A* mRNA or VEGF-A protein levels between the heart and other organs when LNPs and the three different EV types were used as delivery vehicles for intravenous mRNA transport (**supplementary Figures 5C and 6C**). The findings demonstrated that the ratio of *VEGF-A* mRNA in the heart compared to other organs was highest (except for heart/spleen) when CPC-EVs were used as the delivery vehicle (**supplementary Figures 5C**). The quantitative analysis of VEGF-A protein showed that although LNPs delivered efficiently to almost every organ, there was no specific targeting to a particular organ (**supplementary Figure 6B**). However, VEGF-A protein was found to be most abundantly produced in the heart relative to other organs (**supplementary Figures 6C**).

### Vehicles’ specificity, delivery, and their effects on gene expression in the heart tissue

Since the first successful heart catheterization in 1929, cardiac catheter–based delivery methods have been extensively explored in the context of gene- and cell-based therapies enabling efficient treatment of coronary artery disease^3, 24, 25^. However, the application of mRNA therapies for treating cardiovascular disease is only just beginning to emerge. In this context, in the phase IIa trials in cardiac patients undergoing open heart surgery, direct intracardiac injections of *VEGF-A* mRNA were applied in hypoperfused regions^7, 8, 26^. In these studies, the naked *VEGF-A* mRNA was injected directly into the patients’ heart tissues without applying delivery vehicles such as LNPs. Recent study showed LNP-mediated delivery of eGFP mRNA was more efficient than mRNA in citrate buffer upon intramyocardial administration in mice ^27^. Acknowledging the limitations in efficacy associated with the direct injection of naked mRNA into tissues, there is a substantial need to explore mRNA administration strategies using biological vehicles. As a result, the present study aimed to examine the specificity, delivery efficiency, and potential side effects of LNPs as well as three distinct biological vehicles when used for the direct injection of *VEGF-A* mRNA into the heart. Since mRNA delivery is categorized into two aspects of organ targeting (where the homing system facilitates the vehicle’s transport to the target organ) and vehicle-target cell communication (how effectively and safely the vehicle interacts with target cells), we also investigated *VEGF-A* mRNA delivery via direct injection of the vehicles (LNPs and the three EV types) into the mouse heart. The four vehicles (HTB-EVs, HUVEC-EVs, CPC-EVs and LNP) containing an equal amount of *VEGF-A* mRNA or naked VEGF-A mRNA (without vehicle), were directly injected into a single site of the myocardium in the left ventricle (**Figure 5A**). The results showed that, upon delivering an equal amount of *VEGF-A* mRNA, CPC-EVs induced higher VEGF protein production compared to the other vehicles or naked mRNA injection. (**Figure 5B**). However, no significant increase of the VEGF-A protein was detected in the proximate non-injected areas of the heart, liver, or the blood (**Figure 5C-E**). This indicates the efficient localized communication and uptake of CPC-EVs by heart tissue.

**Figure 5.**
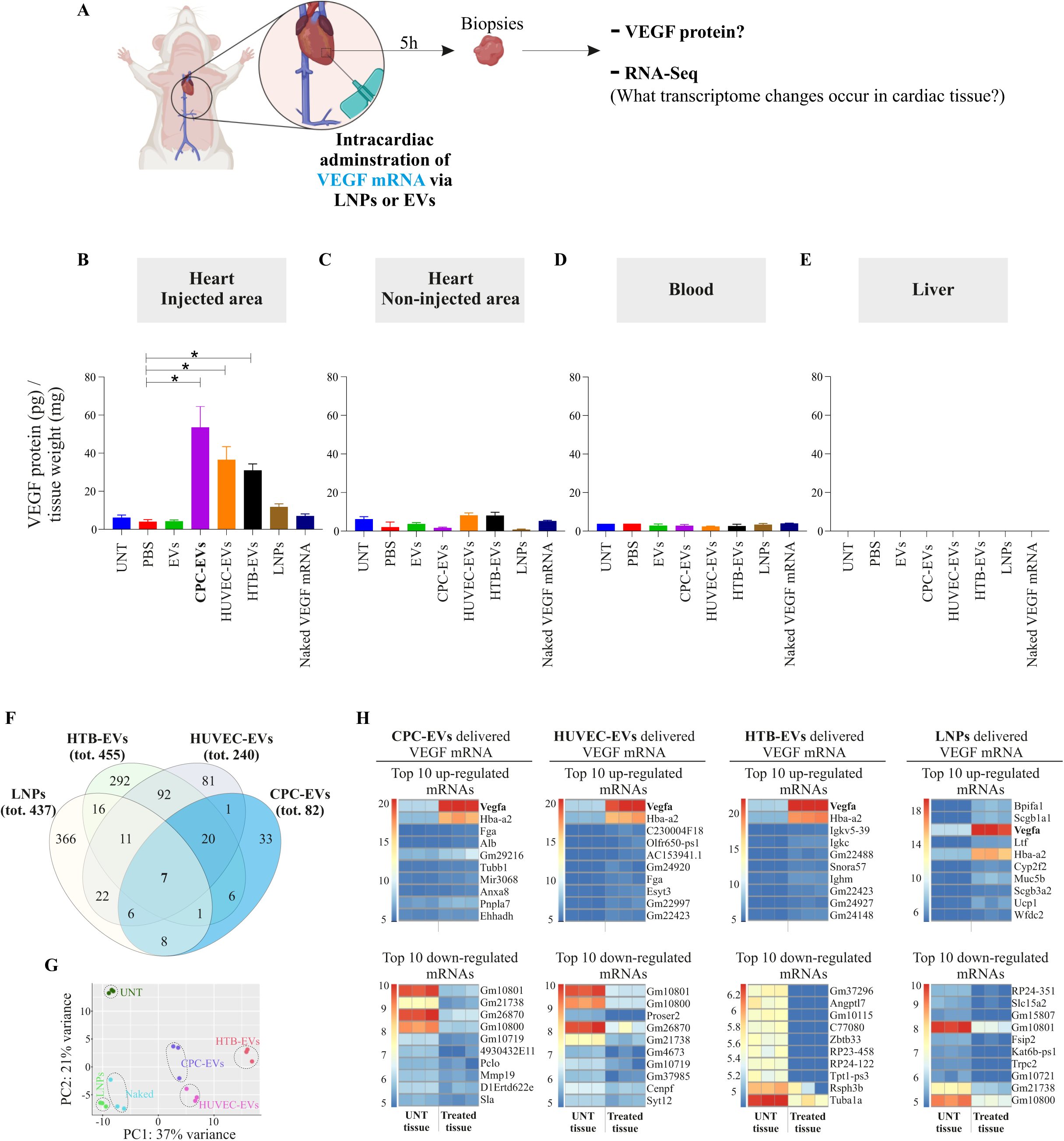
Direct intracardial injection of *VEGF-A* mRNA via cardiac EVs and non-cardiac EVs or LNPs. (**A**) *VEGF-A* mRNA was directly injected via three different EV types or LNPs into a single site of myocardium of mice, and the levels of VEGF-A protein and effect on endogenous gene expression modulations was examined in the heart tissue after EVs or LNP delivery. Untreated mice, and mice treated EVs without *VEGF-A* mRNA were used as controls. Naked *VEGF-A* mRNA without a vehicle was used to compare with delivery by vehicles. 6h post injection, mice were sacrificed, injected areas were dissected and separated from the rest of the heart and examined for the translation of injected mRNA. (**B**) Levels of VEGF-A protein in the injected areas, produced after delivery of *VEGF-A* mRNA via different vehicles. The statistically significant differences between untreated and treated groups, as well as the comparison of CPC-EVs with HTB-EVs, HUVEC-EVs or LNPs were evaluated by One-Way ANOVA (n=3). (**C-E**) Examination of spillover of VEGF-A to non-injected area, and its circulation to blood, or liver. Aorta from 12-week-old male C57BL/6Ncrl mice were collected and embedded in growth factor-reduced Matrigel and were treated with different EV types containing *VEGF-A* mRNA. Human Recombinant VEGF-A protein and Naked mRNA were used as positive controls, and PBS was used as negative control. After 12 days, aortic rings were fixed and stained for α-SMA and CD31 and analyzed by confocal microscopy. 6h post injection, mice were sacrificed, and small pieces of injected area of heart were collected, total RNA was isolated, and RNA sequencing was performed to evaluate the effects of delivery systems on gene expression in the heart. (**F**) Venn diagram showing the total changes in gene expression in the injected area, after delivery of EVs or LNPs. Overall, compared to other vehicles, the CPC-EV delivery caused less disturbance in gene expression with lowest changes in gene expression in the heart. (**G**) Principal component analysis (PCA) plot showing the similarities and differences of transcriptomic profiles in tissue after injection with three EV types, naked mRNA or LNPs. PCA plots were generated from the top 500 most variable genes. The two axes PC1 and PC2 represent the two principal components identified by the analysis. PC1 contributes to 37% of the overall variation among samples and 21 % for PC2. **(H)** top ten upregulated and top ten downregulated transcripts after mRNA delivery via EVs or LNPs. The expression intensities of the heatmaps are visualized based on variance stabilized counts.

Next, we analyzed how recipient cells or tissues responded when mRNA was delivered to the tissue using different vehicles (LNPs or the three EV types). RNA was isolated from the injected area of the heart, and the transcriptome was identified using RNA sequencing. The *VEGF-A* mRNA delivery via CPC-EVs caused the lowest changes in the gene expression in cardiac tissue (only 82 transcripts deregulated) (**Figure 5F)**. In contrast, the delivery via LNPs, HTB-EVs and HUVEC-EVs, caused much higher changes in gene expression in the heart with 437, 445 and 240 significantly differentially expressed genes (DEGs), respectively (**Figure 5F** and **supplementary Tables 1-5**). Transcripts of CPC-EV treated group showed distinct patterns in all data sets. The main variance in the transcriptomic profiles in heart tissue after injection with EVs or LNPs or naked mRNA was visualized in a principal component analysis (PCA), which shows larges differences (37% of the variance) between the EV treatments and the LNP, naked mRNA, and untreated groups (**Figure 5G**). Collectively, the data show that the delivery via CPC-EVs not only yielded the highest levels of the VEGF-A protein but also caused the lowest alterations in gene expression in the heart tissue. This suggests that CPC-EVs are more adapted to communicate with cardiac cells compared to non-cardiac specific EVs or LNPs, or delivery without vehicle. This may be due to the specific molecular signature of CPC-EVs that are recognized by target cells in the heart, leading to improved tissue specificity and efficacy. Additionally, the volcano plots showed dysregulated genes in the tissue after the injection of EVs, or LNPs or naked mRNA into the heart, where CPC-EVs exhibited least distribution in genes and differ significantly from other vehicle groups (**supplementary Figure 7A**). Interestingly, when examining the overlapping gene sets, only seven common DEGs were observed among all four delivery vehicles (**Figure 5F** and **supplementary Figure 7B**).

The bioinformatics analysis identified DEGs after treatments and the top 10 upregulated genes in the heart tissue samples revealed that when *VEGF-A* mRNA was delivered via EVs, the highest represented mRNA was *VEGF-A* (**Figure 5H**). Whereas, in case of naked *VEGF-A* mRNA or delivery via LNPs, the *VEGF-A* mRNA was not the topmost expressed gene in the heart tissue samples. CPC-EVs also show distinct profile of the top 10 downregulated genes identified after *VEGF-A* mRNA delivery. These findings suggest that EVs and LNPs exhibit distinct modes of communication with heart tissue when used for mRNA delivery.

Using further enrichment analyses, we found that CPC-EVs did not induce up-regulation of inflammatory genes in the heart, while other vehicles evoked a significant expression of inflammatory genes, specifically the LNPs (30 up-regulated and 2 down-regulated) (**Supplementary Table 6)**. Moreover, the bioinformatic analysis showed that among the genes activated following the administration of various EV types or LNPs in heart tissue, only the VEGF-A mRNA delivery via CPC-EVs was associated with “myocardial cell development”. This specific gene expression triggers the activation of genes involved in “cardiac muscle cell development” and “factors that promote cardiogenesis in vertebrates” (**Supplementary Table 7-8**). We further identified the top 10 enriched GO biological processes associated to the RNA expression changes in the heart tissue after VEGF-A mRNA delivery (**Supplementary Table 8**). The DEGs were found to be involved in “cellular development, cellular growth and proliferation, and cardiovascular disease, cancer, organismal injury and abnormalities” when *VEGF-A* mRNA was delivered to the heart tissue via different vehicles (**Supplementary Table 9**).

Given the significant challenges associated with analyzing angiogenesis during *in vivo* mRNA delivery, we examined angiogenesis using aortic ring assays *ex vivo* and CD31 and α-SMA expressions were examined as a factor of microvascular density or neo-angiogenesis (**Figure 6A**). Aorta from 12-week-old male C57BL/6Ncrl mice were collected and embedded in growth factor-reduced Matrigel and treated with different EV types containing *VEGF-A* mRNA. Human recombinant VEGF-A protein was used as a positive control. After 12 days, the aortic rings were fixed and stained for α-SMA and CD31 and analyzed by confocal microscopy (**Figure 6B-G**). Importantly, a comparison of CD31 expression after *VEGF-A* mRNA delivery between different EV types, showed that CPC-EVs induce the highest number of vessel formation and higher density of vessels (**Figure 6G**).

**Figure 6.**
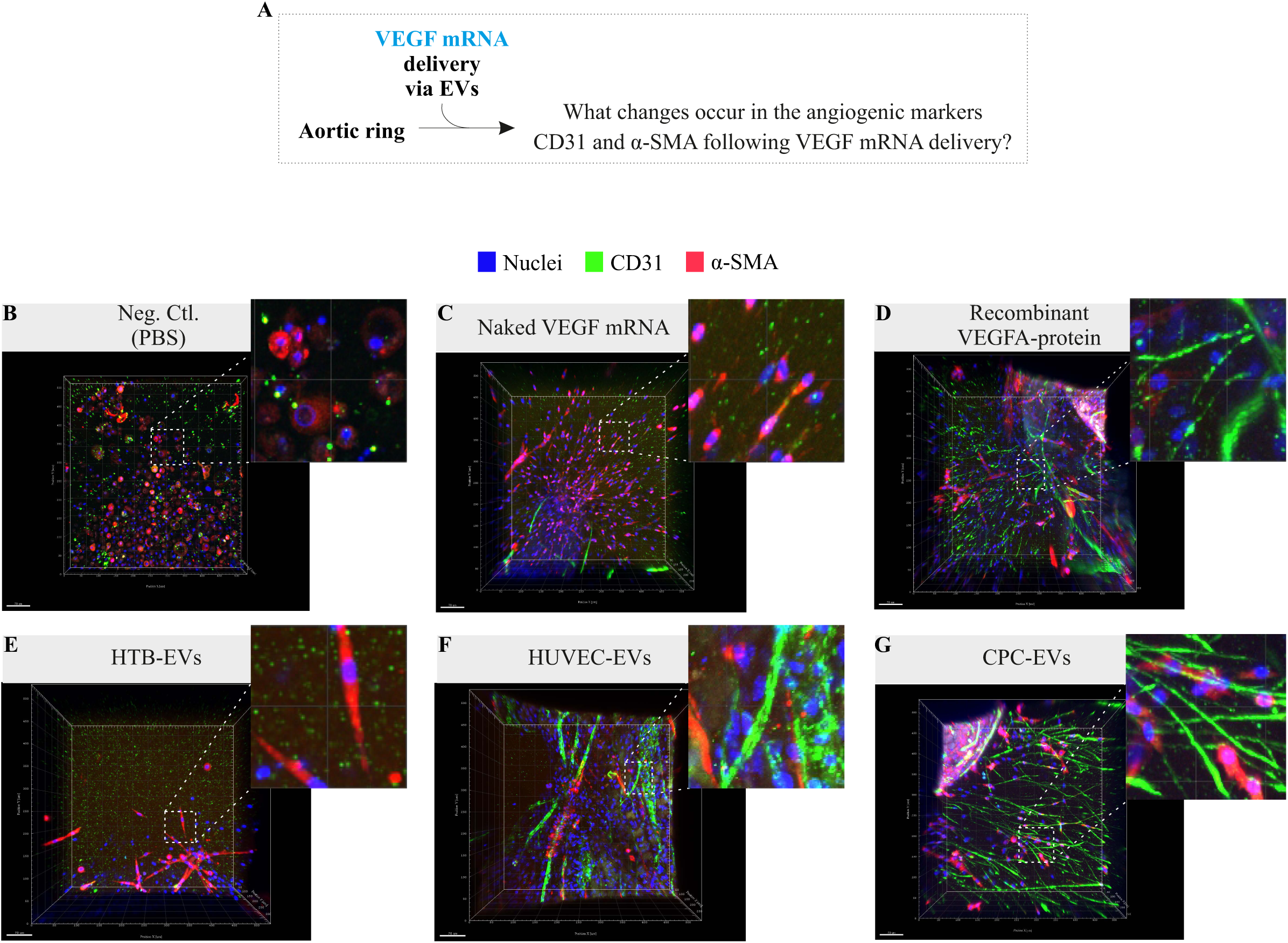
(**A**) Examination of vessel density by immunostaining of CD31 and α-SMA expressions using *ex vivo* aortic ring assay. Aorta from 12-week-old male C57BL/6Ncrl mice were collected and embedded in growth factor-reduced Matrigel and treated with *VEGF-A* mRNA delivery via different EV types. Human recombinant VEGF-A protein was used as a positive control, and PBS as negative control. After 12 days of *VEGF-A* mRNA administration, the aortic rings were fixed and immunoassayed for α-SMA and CD31 and analyzed by confocal microscopy. Examination of CD31 and α-SMA after administration of **(B)** PBS, **(C)** naked *VEGF-A* mRNA, **(D)** recombinant VEGF-A protein, or administration of *VEGF-A* mRNA via **(E)** HTB-EVs, **(F)** HUVEC-EVs and **(G)** CPC-EVs.

## Discussion

RNA-based therapies are a promising field of medical research that involves the delivery of RNA molecules to treat diseases, as has been successfully applied for the production of Covid-19 mRNA-based vaccines ^3, 23, 28–35^. However, the conventional drug delivery systems still lack the techniques for targeted transport of mRNA molecules into targeted site of action at specific organ or cell types. Targeted drug delivery for deploying the specific drug moiety should maximize the availability of the drug at specific sites with non-specific effects and the amount of drug required for greater therapeutic efficacy ^2^. The RNA molecules, like all other drugs, need to be delivered to the target cells or organs in a way that is non-immunogenic and does not cause side effects.

The current study has identified CPC-EVs as a vehicle that effectively targets cardiac tissue for the delivery of exogenous mRNA molecules, such as *VEGF-A* mRNA. Notably, CPC-EVs demonstrate both systemic and direct injection routes for mRNA delivery to the heart with minimal effects on gene expression in the cardiac tissue. It is important to clarify that Figures 3 and 4 show the amount of detected *VEGF-A* mRNA (in nanograms) per milligram of tissue from each organ. The detected amount of *VEGF-A* mRNA is not normalized against the total weight of each examined organ, due to the significant weight differences between various organs. For example, the mouse liver is roughly 19 times heavier than the mouse heart and about 500 times heavier than the adrenal glands (the adrenal glands were not included in our study). Had the adrenal glands been included, it would suggest that even though 100 moles more *VEGF-A* mRNA could potentially be detected per milligram of adrenal tissue compared to the liver, the liver’s total *VEGF-A* mRNA content would still be considerably higher, approximately five-fold more than in the adrenal glands. Therefore, we opted to report the detected *VEGF-A* mRNA levels as normalized per milligram of tissue from each organ studied. When equal amounts of *VEGF-A* mRNA were administered intravenously using LNPs or different types of EVs, the findings underscored the potential of CPC-EVs as a safe and effective approach for mRNA-based therapies in cardiac regeneration and repair. In addition, we found that CPC-EVs were also most effective in communicating with cardiac muscle cells regarding optimal production of the desired protein (VEGF-A) and with the least alterations in gene expression, compared to other vehicles tested (**Figures 5-6**). In comparison to LNPs, HTB-EVs, or HUVEC-EVs, the EVs derived from cardiac progenitor cells demonstrated a remarkable specificity or homing for the delivery of *VEGF-A* mRNA to heart and is also efficient in producing significant amounts of fresh VEGF-A protein in the heart. Several studies have reported that most of the systemically administered mRNAs using LNPs, or EVs are accumulated in the liver via circulatory system ^3–6^. In this regard, CPC-EVs are not only the lowest accumulated in the liver but also show tissue specific delivery of *VEGF-A* mRNA to heart, compared to other vehicles used in this study.

Drug delivery, including mRNA-based therapeutics, has three major attributes (i) organ targeting (where the homing system facilitates the transport of the vehicle to the target organ) (ii) mRNA delivery to target cells/tissue (vehicle-cell communication), and (iii) the effect of the delivered mRNA (how the target organ/tissue is impacted by the vehicle and its mRNA cargo). The data from the current study reasonably indicate that CPC-EVs demonstrated these three attributes with remarkable properties of tissue targeting, mRNA delivery, and lowest changes in the transcriptomics especially those associated with inflammatory responses (**Figure 7**). The data from this study also demonstrates that CPC-EVs have a different transport capacity than LNPs for the delivery of the same mRNA. Our results in this study demonstrate that upon intravenous administration CPC-EVs are the most effective at transporting administered mRNA to cardiac tissue compared to LNPs and the other EV types used in this study. We hypothesize that these EVs (CPC-EVs) may carry surface proteins recognized by the homing system, enabling their preferential transport to the heart relative to other organs examined in this study. EVs or biomimetic particles can have homing capacity as they conserve the surface characteristics of their parental cells ^36^. This is particularly interesting, given that immunofluorescence staining of CD31 and α-SMA showed that the EV-mediated delivery of *VEGF-A* mRNA to aortic rings resulted in a significantly higher vessel density (**Figure 6**). This increase in microvascular density indicates an improved angiogenic response, highlighting the potential of *VEGF-A* mRNA therapy in stimulating new blood vessel formation. These results reinforce the hypothesis that targeted mRNA delivery could potentially be an effective method for promoting vascular growth. In this regard, application of CPC-EVs with inherent tropism for the heart tissue is of paramount importance for *VEGF-A* mRNA transport to heart as well as highest levels of protein production, especially when the delivery of specific mRNA drugs to their site of action remains a critical issue.

**Figure 7.**
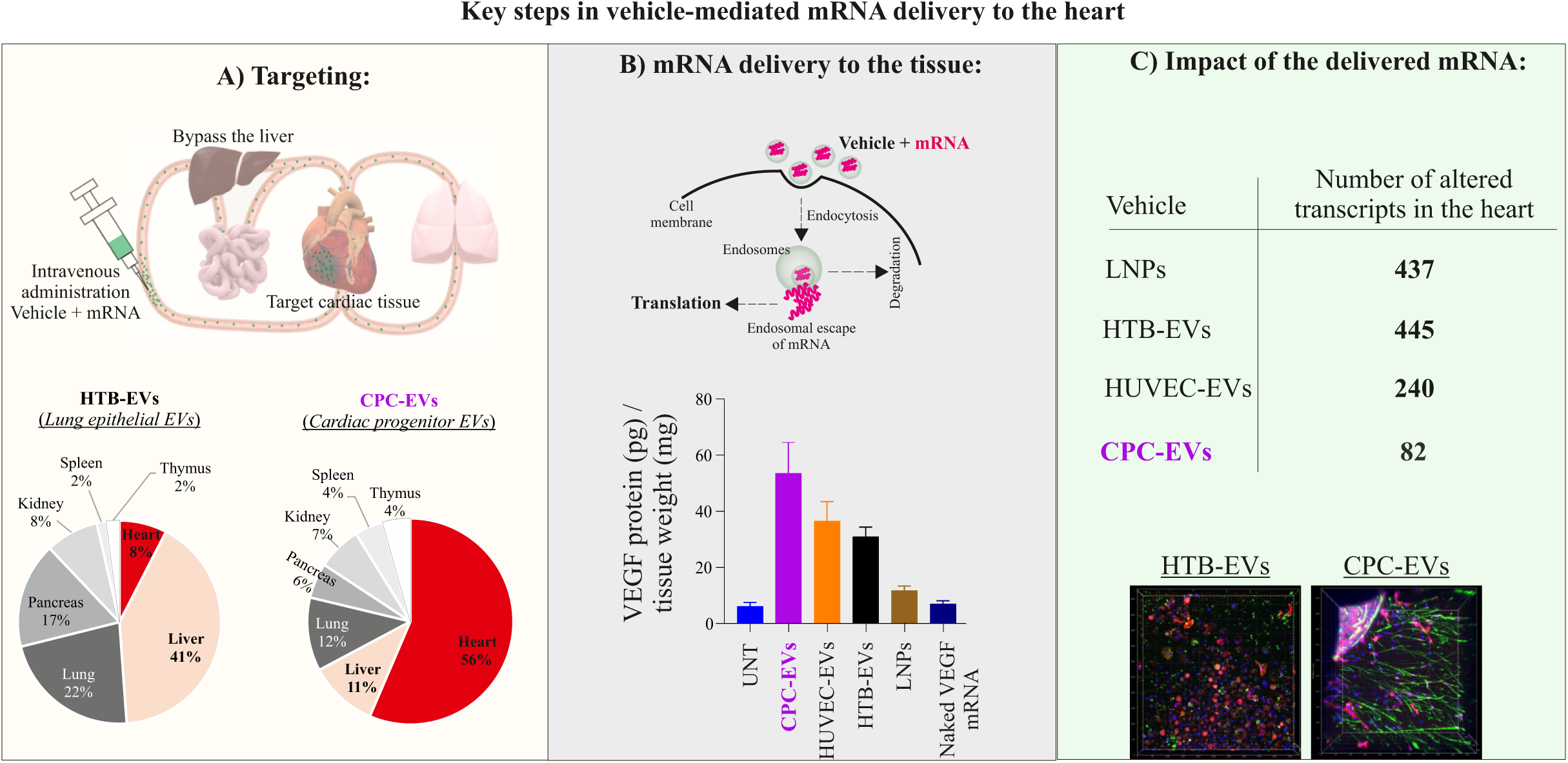
Summary of organ targeting and mRNA delivery by tissue-specific vehicles, liver bypass, heart specific delivery and side effects on mRNA changes in the injected heart tissue. Since the mRNA-therapeutics is about targeting, delivery efficiency, and the least non-specific delivery or side effects, the CPC-EVs demonstrated remarkable properties of tissue targeting, mRNA delivery, and lowest changes in the transcriptomics especially those associated with inflammatory responses.

Targeted delivery of *VEGF-A* mRNA to heart tissue is thought to be a promising approach for the treatment of cardiovascular diseases, by stimulating neovascularization (angiogenesis) ^7, 8^. Therapeutic vascularization with VEGF-A protein has been studied for decades experimentally and clinically ^9–11^. A decade-long effort by researchers demonstrated robust, transient expression of VEGF-A protein for several days using chemically modified *VEGF-A* mRNA in mouse models, large animals, and a first-in-human clinical study ^8, 12–17^. However, in first-in-human clinical study, direct intracardiac injections of naked *VEGF-A* mRNA were applied without using LNPs. It has been long sought that in addition to minimally invasive localized mRNA administration, there is a critical need of investigating the mRNA administration via packing systems ^3^. However, RNA packaging systems have so far been designed for the efficient delivery of RNA molecules but with little success on cellular specificity. Additionally, several studies have reported that most of the systemically administered mRNAs using LNPs, or EVs are accumulated in the liver via the circulatory system ^3–6^. Here, we show that tissue-specific EVs can transport the exogenous mRNA more efficiently to the target organ with least accumulation in the liver. Unlike CPC-EVs, the LNP-mRNA accumulated in liver the most, which is in line with previous studies which have reported that LNPs accumulate in the liver when administered intravenously ^4, 5^.

Some of the alternative routes of administration include inhaled delivery of mRNA nanoparticles to target lung-specific sites ^37^, oral delivery of mRNA via robotic pills to target gastrointestinal tract sites ^38^, and direct injection of mRNA to heart ^7^. It is also worth noting that although localized mRNA interventions to heart tissue have been successfully applied, new nano-formulations for organ-specific delivery of mRNA are equally meaningful. In this regard, alternative routes of administration can be a more practical solution in some contexts, especially to avoid invasive procedures for targeting organs deeply located in the body. For instance, intravenous injections of mRNA embedded in nanocarriers, such as EVs derived from cardiac cells loaded with mRNA, can be employed to target specific organs. This approach allows for non-invasive systemic delivery of mRNA to the heart, providing a promising alternative for efficient and targeted therapy. Even if it is a fraction of the delivered *VEGF-A* mRNA that reaches the heart via non-invasive routes, this would represent a lucrative advantage for treating ischemic cardiovascular diseases. Targeting the heart, therefore, could benefit patients who otherwise have no known vehicles to receive *VEGF-A* mRNA in the heart. This warrants further investigations on whether EVs from homologous organs can be used for tissue-customized mRNA delivery. Current study opens a window of opportunity, that it might be possible to identify EVs that are cell/tissue-customized for delivery to specific cells and organs. As natural vesicles, EVs possess multiple advantages in terms of drug delivery, such as biocompatibility and hypoimmunogenicity ^39^, and they have natural tendency to transport RNA between cells ^40^. To maximize the mRNA delivery at site of action, further modifications can be made to EVs such as surface modifications conjugated with receptors, peptides, or antibodies, to generate enhanced targeting. It is possible to customize EVs, especially of those secreted from parent (source) organ and given back. Moreover, homotypic EVs, derived from the same cellular source as their target organ, may offer advantages over heterotypic EVs due to their specific characteristics.

In summary, we conclude that CPC-EVs are superior in (a) mRNA targeting to the heart, (b) communication with cardiac cells, and (c) causing minimal transcriptomic changes during *VEGF-A* mRNA delivery. Thus, our data suggests that CPC-EVs may be an ideal candidate for mRNA-based heart therapies. They are not only effective in targeting the transport of mRNA to the heart but can also best interact with the target organ (heart) to optimize the production of the desired protein. However, future research is essential to understand the homing properties of CPC-EVs, which may enable the development of safe and efficient biological vehicles for targeted drug delivery to specific tissues.

## METHODS

### Construction of *VEGF-A* mRNA sequence and clean capping

The CDS sequence of *VEGF-A* 165 (isoform 11) containing an open reading frame with start and stop codon (576 nucleotides including atg start and tga stop codons, encoding for 191 amino acids of VEGF-A protein) was selected. The sequence code was provided to TriLink® Biotechnologies (CA, USA) for *VEGF-A* mRNA construct with clean cap modifications. A CleanCap® (AG, polyadenylated) method was applied which is a co-transcriptional capping method with a fully processed mature mRNA and was optimized for mammalian systems. After clean cap modifications with polyadenylation, the resulting mRNA length was 852 nucleotides. The prepared *VEGF-A* mRNA was dissolved in 1 mM sodium citrate buffer (pH, 6.4), and stored at -80 °C. The sense strand of *VEGF-A* mRNA used in this study is presented in **supplementary Fig. 1b,** and the sequence of the encoded VEGF-A protein is provided **supplementary Fig. 1c**. The VEGF-A protein encoded from the given sense strand of *VEGF-A* mRNA was obtained using the transcription and translation tool http://biomodel.uah.es/en/lab/cybertory/analysis/trans.htm.

### mRNA loading and characterization of formulated LNPs

DLin-MC3-DMA LNPs containing modified *VEGF-A* mRNA (852 nucleotides, 5meC, Ψ TriLink Biotechnologies, USA) were prepared by precipitating the mRNA with four different lipid components as described previously^21^. These components consist of an ionizable lipid, DLin-MC3-DMA (MC3), two helper lipids (DSPC and Cholesterol) and a PEGylated lipid (DMPE-PEG2000). Chemical structure of MC3-LNPs is presented in **Supplementary** Fig. 1a. A solution of *VEGF-A* mRNA in water was prepared by mixing mRNA dissolved in MilliQ-water, 100mM citrate buffer pH=3 and MilliQ-water to give a solution of 50mM citrate. Lipid solutions in ethanol (99.5%) were prepared with a composition of four lipid components [MC3:Cholesterol:DSPC:DMPE-PEG2000] = 50:38.5:10:1.5 mol% and a total lipid content of 12.5mM. The mRNA and lipid solutions were mixed in a NanoAssemblr (Precision NanoSystems, Inc., BC, Canada) microfluidic mixing system at a volume mixing ratio of 3:1 and a constant total flow rate of 12 mL/min. At the time of mixing, the ratio between the nitrogen atoms on the ionizable lipid and phosphor atoms on the mRNA chain was 3.08:1. In some preparations of LNPs, CleanCap® Cy5-*eGFP* mRNA (996 nucleotides, 5meC, Ψ) and CleanCap® *eGFP* mRNA (Trilink Biotechnology) were mixed with 1:1 ratio and encapsulated instead of *VEGF-A* mRNA. The initial 0.25 mL and the last 0.05 mL of the LNP solution prepared were discarded while the rest of the volume was collected as the sample fraction. The sample fraction was transferred immediately to a Slide-a-lyzer G2 dialysis cassette (10000 MWCO, Thermo Fisher Scientific Inc.) and dialyzed overnight at 4°C against PBS (pH7.4) to remove residual ethanol (25 v%). The volume of the PBS buffer was 600x the sample fraction volume. The dialyzed sample was collected and filtrated through a 0.22 µm sterile filter (Gillex, Merck) prior any characterization.

### Culturing and treatment of HTB-177, HUVECs, and CPC with LNP-*VEGF-A* mRNA

The HTB-177 (NCI-H460, ATCC) cell line was cultured in RPMI-1640 growth medium supplemented with 10% EV-depleted fetal bovine serum (FBS), 1% L-glutamine, and 1% penicillin-streptomycin at 37°C and 5% CO2. The medium was replaced every 48 hours, and LNP-*VEGF-A* mRNA was added for an experimental period of 24 hours. EV-depleted FBS was prepared by heat inactivating FBS at 56°C for 1 hour, followed by ultracentrifugation at 120,000 x g for 2 hours at 4°C. The EV-depleted supernatant was filtered through 0.2μm filters before incorporation into the growth medium. HUVECs (Lonza, Switzerland) were cultured in T-75cm2 culture flasks with EGM-Plus medium at 37°C and 5% CO2. The medium was replaced every 2 days, and LNP-*VEGF-A* mRNA was added when cells reached around 80% confluency. At the endpoint, the conditioned medium was collected for EV isolation, and cells were detached and processed for further analysis. iCell cardiac progenitor cells (Fujifilm Cellular Dynamics, Madison, WI, USA) were thawed, resuspended in maintenance medium, and seeded in fibronectin-coated 6-well plates. After 24 hours, the medium was replaced, and LNP-*VEGF-A* mRNA was added for an additional 24 hours. At the endpoint, conditioned medium was collected for EV isolation, and cells were detached, counted, and checked for viability. All steps were conducted under aseptic conditions.

### Characterization of the formulated LNPs

The intensity-averaged particle size (Z-average, dZ) was measured using ZetaSizer (Malvern Instruments Inc.). The measurement solution was made by diluting 20 µL of the sample fraction using 980 µL PBS (pH 7.4). The mRNA concentration and encapsulation efficiency (EE) of the final product was measured by Quant-it Ribogreen Assay Kit (Thermo Fisher Scientific).

### Culturing and Treatment of HTB-177, HUVECs, and CPC with LNP-*VEGF-A* mRNA

The human epithelial HTB-177 (NCI-H460, ATCC) cell line was cultured according to the manufacturer’s protocol. The cells were cultured in RPMI-1640 growth medium containing sodium bicarbonate, without sodium pyruvate and HEPES (Sigma Aldrich), supplemented with 10% EV-depleted Fetal bovine serum (FBS) (Sigma Aldrich), 1% of L-glutamine and 1% penicillin-streptomycin (Thermo Fisher Scientific), at 37°C and 5% CO2. The medium was replaced with a fresh medium after 48 h, followed by adding LNP-*VEGF-A* mRNA to the cells in culture for an experimental period of 24 h. The heat inactivated FBS was prepared by incubating it at 56°C for 1 hour. EV-depletion was achieved by ultracentrifugation at 120,000 x g for 2 hours at 4°C using an Optima L-100 XP ultracentrifuge with a 70Ti rotor (Beckman Coulter). The EV-depleted supernatant was then filtered through 0.2μm filters before being incorporated into the RPMI-1640 growth medium. The HUVECs cell (Lonza, Switzerland), were plated and expanded in culture medium according to the manufacturer’s instructions (CC-5035 EGM-PLUS BulletKit Medium; CC-5036 EGM-PLUS Basal Media + CC-4542 EGM-PLUS SingleQuots Kit, Lonza, Switzerland). Briefly, cells were cultured in T-75cm^2^ culture flasks with EGM-Plus medium and incubated at 37°C in 5% CO2 and 95% saturated atmospheric humidity. The culture medium was replaced with fresh media every 2 days until the cells attained around 80% confluency, and then the cells were expanded. At 80% confluency, the cells were rinsed with Ca^++^/Mg^++^ free Dulbecco’s phosphate buffered saline. TrypLE™ Express Enzyme (1X), without phenol red, was added to detach the cell layer from the flask. The enzyme activity was stopped by adding the complete culture medium to the flask. The cells were aspirated by gently pipetting and transferred to a tube and centrifuged at 1000 rpm for 5 min. The HUVEC cell pellet was resuspended with fresh culture medium and dispensed into a T-75cm^2^ culture flask at a density of 2×10^6^ cells/flask and incubated at 37°C and 5% CO2 for 24 h followed by addition of LNP-*VEGF-A* mRNA to each flask for an experimental period of additional 24 h. At the endpoint, the conditioned medium was collected for EV isolation. Also, the cells were detached from the culture flask using TrypLE™, as described before. The cells were counted and checked for their viability before centrifugation. The cell pellets were used for further analysis. All the steps were carried out under aseptic conditions.

iCell cardiac progenitor cells (R1093, Fujifilm Cellular Dynamics, Madison, WI, USA) were thawed, centrifuged (180 ×*g* for 5 min), resuspended in the maintenance medium composed of William’s E Medium, Cocktail B (Thermo Fisher Scientific) and seeded in fibronectin-coated (1 mg/mL fibronectin solution diluted in sterile D-PBS to a final concentration of 5 µg/mL immediately before use, Sigma-Aldrich, St. Louis, MO, USA) 6-well plates at a density of 5x10^5^ cells/well. Cells were then incubated at 37 °C in the ambient atmosphere supplemented with 5% CO2 and 95% relative humidity. The medium was replaced with a fresh maintenance medium after 24 h, followed by adding LNP-*VEGF-A* mRNA to each well for an experimental period of an additional 24 h. At the endpoint, the conditioned medium was collected for EV isolation. Also, the cells were detached from the culture plates using TrypLE™, as described before, counted, and checked for their viability. The pelleted cells were used for further analysis. All the steps were carried out under aseptic conditions.

### Isolation and separation of EVs from LNP treated cells

Prior to EV isolation, the cell debris was removed from the collected conditioned medium by centrifugation at 3000 x g for 15 min at 4°C on a 4K15 centrifuge (Sigma Aldrich). EVs from the conditioned media of LNP-mRNA treated cells and controls were isolated by size exclusion chromatography (SEC) using qEV-70/10 mL columns (Izon Science Ltd, New Zealand) according to manufacturer’s guidelines which suggest the collection of 4 fractions, 5 mL each. However, some modifications were applied to the original protocol, with the following amendments. (i) 15 mL media was used instead of 10 mL media. (ii) As we had 15 mL media, therefore, instead of 4 fractions, 6 fractions (5 mL each) typically represent EV fractions were collected for EV analysis. Each fraction was concentrated using 30 KDa amicon ultra/15mL centrifuge filters (cat: # UFC903024, Sigma Aldrich, now Merck), at 4000xg, for 25 min, 4 °C. The total RNA was isolated from each fraction and *VEGF-A* mRNA was detected/quantified by qPCR, in each fraction. Fractions 1-6 were analyzed for the presence of *VEGF-A* mRNA and protein. (iii) fractions 7 to 12 (non-EV fractions) were also collected to analyze any mRNA. The VEGF-A protein was analyzed in all fractions, 1-12 (15 mL from T75 flasks HTB-177 cultures, and from 6 well plates of CPC cultures: 5 wells x 3 mL = 15 mL). Briefly, after loading 15 mL of sample into the column reservoir, a 20 mL void volume was first discarded, followed by the collection of 12 fractions, each of a 5 mL collection (elution) volume.

### Quantification of *VEGF-A* mRNA in EV fractions

The total RNA from LNP-mRNA treated HTB-177, HUVECs, CPCs and their secreted EVs was isolated using miRNeasy Mini Kit (Qiagen, cat. #: 217004) according to the manufacturers’ guidelines. Total RNA was quantified by Qubit 2.0 fluorometer (Thermo Fisher Scientific) and NanoDrop 1000 (Thermo Fisher Scientific). The RNA quality was assessed by a 230/260 ratio recorded on NanoDrop. RNA samples from untreated cells and their EVs were used as controls.

LNP-*VEGF-A* mRNA from cell lysates and their secreted EVs was detected and quantified by real-time qPCR. 30-50 ng of total cellular total RNA were reverse transcribed into cDNA using high-capacity cDNA kit with RNase inhibitor (Thermo Fisher Scientific: 4374966). 45ng of cDNA was used for VEGF-A mRNA quantification using hydrolysis probes (TaqMan probe assay, Thermo Fisher Scientific, assay ID: Hs00900055_m1) on ViiA™ 7 instrument according to the manufacturer’s instructions. To generate the standard curve for absolute quantification, the VEGF-A mRNA standards were prepared using pure VEGF-A mRNA (TriLink Biotechnologies, USA). 2 µg of the pure VEGF-A mRNA was reverse transcribed into cDNA, and VEGF-A cDNA was serially diluted ten-fold (highest point: 100 ng, and lowest point: 0.0001 ng) to generate a standard curve. The assay was performed in technical triplicates. For the absolute quantification of VEGF-A mRNA, the cDNA from the cell and EV samples was interpolated against the VEGF-A standard curve with minimal R2 > 0.975. GAPDH gene (Thermo Fisher Scientific, assay ID: Hs02758991_g1) was used as internal control.

### Detection of VEGF-A protein in EV fractions

The VEGF-A protein was quantified in the individual EV fractions (F1-12) isolated from HTB-177 treated with LNP-VEGF-A mRNA. EV fractions isolated from untreated cells were used as controls. The human VEGF-A sandwich ELISA Kit (cat.#: RAB0507, Sigma Aldrich, now Merck) was used to detect and quantify the VEGF-A protein according to manufacturer’s instructions. 100 µL of EV solution or serially diluted VEGF-A protein standards were added per well. VEGF-A protein concentration (pg/mL) was recorded on ELISA reader instrument (Spectra max, 340PC, molecular devices), as the VEGF-A protein levels relative to VEGF-A standard curve. The levels of VEGF-A protein in EVs were normalized to total EV-proteins (µg).

### Characterization of EVs

#### Determination of size and concentration of SEC-EVs

The size and concentration (particle number) of EVs isolated by size exclusion chromatography from all three cell types were assessed using Nanoparticle Tracking Analysis (NTA) with an LM14c instrument from Malvern Panalytical, UK, equipped with an sCMOS camera.

Initially, The EV pellets were dissolved in 1000 µL of PBS and then further diluted 5-fold with PBS to ensure that the number of particles in the field of view remained below 100 particles per frame. Independent measurements, in scatter mode, were performed on two biological replicates from each time point. Each EV sample was analyzed through three captures, with a total of 2248 frames, using an adjusted camera level of 16 and a detection threshold of 5. The settings for Blur and Max Jump Distance were left on auto. Data acquisition and analysis were carried out using NanoSight Fluorescent NTA LM14c software version 3.2 (Malvern Panalytical, UK).

### Detection of CD63 and CD9 EV markers in SEC-EVs

EVs were initially isolated using size exclusion chromatography as described above. Subsequently, the CD63 and CD9 positive EVs were isolated using an immunoaffinity-based method. The CD63 isolation/detection reagent for cell culture medium (Thermo Fisher Scientific, cat.#: 10606D) was used to immobilize the CD63^+^ EVs to magnetic dynabeads conjugated with anti-CD63 antibody. In the binding reaction, 30 µL of CD63-antibody conjugated beads were incubated with 60 µg of EVs (beads + EVs; total volume 120 µL). As negative control, 30 µL of CD63-antibody conjugated beads alone, were incubated with an equivalent volume of PBS (no EVs). The EVs were immobilized on anti-CD63 beads and incubated at 4°C overnight (day 1). Then next day (day 2) unbound beads or EVs were washed 4 times with BSA-PBS isolation buffer (0.25% BSA dissolved in PBS). The washing steps were performed using magnetic separators (EasySep, StemCell technologies), according to the manufacturer’s protocol (cat.#: 10606D). After the final wash, the immobilized CD63^+^ EVs were suspended in 120 µL of BSA-PBS isolation buffer and then further stained with a mouse anti-human PE-CD9 antibody (BD Pharmingen™, cat.#: 555372). A 20 µL of PE-CD9 antibody was added to CD63^+^ EV solution and incubated in a sample shaker for 1 h, at room temperature (in the dark). To remove unbound CD9-antibody, the sample was washed 4 times with BSA-PBS isolation buffer using magnetic separators and finally suspended in 200 µL of isolation buffer. The immobilized CD63^+^ EVs were acquired on a BD FACSLyric system (BD Biosciences) to detect CD9^+^ EVs. The data were analyzed using FlowJo software (TreeStar Inc.). The experiment was performed in biological duplicates.

### Transmission Electron Microscopy (TEM) Analysis of SEC-EVs

The SEC-EVs isolated from HTB-177, HUVECs, and CPCs were fixed in 2% Paraformaldehyde - 0.1M phosphate buffered saline for 30 min. Subsequently, a Glow discharge technique (30 sec, 7,2V, using a Bal-Tec MED 020 Coating System) was applied over carbon-coated copper grids, and immediately, the grids were placed on top of sample drops for 15 min. Then, the grids with adherent EVs were washed in a 0.1M PBS drop. An additional fixation in 1% glutaraldehyde was performed for 5 min. After washing properly in distilled water, the grids were contrasted with 1% uranyl acetate and embedded in methylcellulose. Excess fluid was removed and allowed to dry before examination with a transmission electron microscope FEI Tecnai G2 Spirit (Thermo Fisher Scientific, Oregon, USA). All images were acquired using Radius software (Version 2.1) with a Xarosa digital camera (EMSIS GmbH, Münster, Germany).

### EV-mediated delivery of translatable *VEGF-A* mRNA to human endothelial cells *in vitro*

Further, we investigated whether EVs isolated by SEC could deliver a VEGF-A mRNA to cells. Pooled and concentrated EV fractions (F2-F6) of HTB-EVs, HUVEC-EVs and CPC-EVs containing VEGF-A mRNA (550 ng), were delivered to HUVECs. LNP-VEGF-A mRNA was used as positive control. Untreated cells and EVs without VEGF-A mRNA were used as negative controls. 24 post incubation, the production of VEGF-A protein was quantified in the conditioned medium using Human VEGF-A sandwich ELISA Kit. The presence or absence of endogenous VEGF-A mRNA was also investigated in untreated samples.

### Intramuscular and intravenous delivery of luciferase mRNA via SEC-EVs or LNPs and IVIS analysis

SEC-EVs or LNPs containing firefly luciferase mRNA (FLuc-mRNA, 215 ng, in 30 µL) were administered intramuscularly to female C57bl/Ncr mice. In a separate experiment, mice were intravenously administered with EVs or LNPs containing FLuc-mRNA (1 µg, in 100 µL). To confirm the translation of mRNA into luciferase, a luciferin (5 ml/kg) was administrated intramuscularly or intravenously 6 hours after the FLuc-mRNA delivery. The mice were terminated 20 minutes after the luciferin administration and the organs were dissected and scanned with an IVIS Spectrum within 5 minutes of termination. The total radiance was quantified as a factor of luciferase produced from FLuc-mRNA. After intramuscular delivery of FLuc-mRNA the radiance (luciferase/luciferin activity) was quantified in dorsal and left muscle of mice after Fluc-mRNA delivery via HTB-EVs, and LNPs. Nonparametric One-Way ANOVA followed by the Mann-Whitney U-test was applied to compare statistical differences between luciferase total flux after FLuc-mRNA delivery via HTB-EVs and LNPs.

### Biodistribution of *VEGF-A* mRNA delivered via EVs or LNPs

Female C57bl/Ncr mice were intravenously administered with 100 µL of EVs or LNPs containing 1 µg of VEGF-A mRNA. The mice were sacrificed 6h post administration and organs were collected and snap frozen. Total RNA and total proteins were extracted from the organs as described above. qPCR was performed to quantify the VEGF-A mRNA as described above.

VEGF-A protein was quantified using human VEGF-A sandwich ELISA Kit (cat.#: RAB0507, Sigma Aldrich, now Merck) according to manufacturer’s instructions. The total VEGF-A mRNA and protein quantified in the used tissue was normalized per total organ weight of tissue sample.

### Intramyocardial injections of *VEGF-A* mRNA via EVs or LNPs and detection of VEGF-A protein

The animal experiments followed the NIH guidelines and were approved by the Gothenburg University Animal Ethics Committee (Gothenburg Ethical Review Board number Ea 001173-2017). Male C57BL/6Ncrl mice at 10-12 weeks of age and weight of ∼25 g, were purchased from Charles River and housed on a 12 h light/12 h dark cycle, ambient temperature at 21 – 22ᵒC and 50% humidity. On the day of the injections, mice were anesthetized with 2–3% Isoflurane® mixed with oxygen, intubated, and connected to a ventilator. The mice were ventilated with air ∼800 mL/min and oxygen ∼100 mL/min (∼230 strokes/min (MiniVent Ventilator for Mice (Model 845), Harvard Apparatus, Holliston, MA). Core temperature was continuously monitored and maintained at 35 - 36.5ᵒC by a heating operating table and heating lamp controlled by rectal thermometer. Electrodes were inserted under the skin to register the heart rate and electrical activity (PharmLab, Paris, France). The mice were subjected to a left thoracotomy at the fourth intercostal space ∼2 to 3 mm to the left of the sternum. A rib spreader was used to keep the incision open. The pericardium was opened, the heart was held using an USP 8-0 suture (Braun, Kronberg im Taunus, Germany) and 40 µL of EVs or LNPs (corresponding to 50 ng VEGF-A mRNA) or PBS (no mRNA) treatments were injected in one single site in the myocardium of the left ventricle using an insulin syringe (Becton, Dickinson and Company, Franklin lakes, NJ). 50 ng of naked VEGF-A mRNA in citrate saline solution (without loading into EVs or LNPs) was also injected in separate groups. The chest was then closed, and the mice were monitored during continued maintenance of body temperature and ventilation until it regained consciousness and could be disconnected. The mice were sacrificed 6h post injection, and the heart, liver and blood were collected. For the heart, the area of injection in the left ventricle was dissected and separated from the rest of the heart and snap-frozen. The remaining parts of the heart (remote left ventricle, right ventricle and atria) were snap-frozen in a second tube and referred to as remote non-injected area for further analysis. The animals were divided into 5 groups either for vehicles loaded with VEGF-A mRNA such as HTB-EVs, HUVEC-EVs, CPC-EVs, LNPs or naked VEGF-A mRNA in citrate buffer without vehicle. Each group consisted of three individuals, and they were injected separately. Untreated naïve mice or treated with equal volume of PBS with mRNA were used as controls.

### Detection of human VEGF-A mRNA and protein in the heart and evaluation of spill over

At 6h post injection of VEGF-A mRNA via SEC-EVs, the total mRNA and total proteins from injected and non-injected heart areas as well as from liver were extracted. Total RNA from 20-70 mg of each tissue was extracted using miRNeasy Mini Kit (Qiagen, cat. #: 217004). The tissue was placed in 2mL Eppendorf tubes with 700 uL of QIAzol lysis reagent and beads and were lysed in Tissue LyserII (Qiagen) for 5 min at the maximum speed (30Hz). The RNA isolation steps were followed according to manufacturer’s guidelines provided with kit. Total extracted RNA was quantified by Qubit 2.0 fluorometer (Thermo Fisher Scientific) and NanoDrop 1000 (Thermo Fisher Scientific). The RNA quality was assessed by a 230/260 ratio recorded on NanoDrop. RNA samples from untreated cells and their EVs were used as controls. VEGF-A mRNA was quantified in each sample using qPCR, as described above.

For total protein extraction, 20-70 mg of each tissue was lysed in T-PER Tissue Protein Extraction Reagent (Thermo Fisher Scientific Cat. 78510) in the presence of 1% halt protease inhibitor cocktail, EDTA free (Thermo Fisher Scientific, cat.#: 87785), following the manufacturer’s instructions. After lysing with Tissue LyserII (Qiagen) for 5 min at speed of 30Hz, the tissue lysates were centrifuged at 10,000 x g for 15 min at 4°C to deplete tissue debris. The upper phase was transferred to a new tube and the pellet was discarded. The blood was centrifuged at 2000x g, for 5 min, 4 0C to collect the plasma. The extracted proteins from tissues, and plasma were quantified by Qubit 2.0 fluorometer (Thermo Fisher Scientific). Human VEGF-A protein was quantified by VEGF-A sandwich ELISA (Cat. #: RAB0507, Sigma Aldrich) performed separately on the samples from injected area of the left ventricle and proximate non-injected area of the heart as well as the liver, and blood. The amount of VEGF-A protein (pg/mL) in each organ was normalized to the relative organ weight (g).

### Aortic ring assay

The aortic ring assay was performed as described ^41^. Aorta from 12-week-old male C57BL/6Ncrl mice were collected. Fat tissue was removed from the aorta using forceps and blood remaining inside was washed out by flushing Opti-MEM^TM^ (ThermoFisher Scientific) supplemented with 2.5% of Fetal Bovine Serum (FBS; ThermoFisher Scientific) and 1% of Penicilin-Streptomycin (P/S; ThermoFisher Scientific). Then, the aorta was sliced into 0.5 mm rings and serum starved overnight in Opti-MEM supplemented with 1% P/S. Rings from different aortas were kept separated. The following day, aortic rings from different animals were randomized and embedded individually in 50 µL of growth factor-reduced Matrigel (Corning) on 96-well plates. After aortic rings embedding, Opti-MEM medium supplemented with 2.5% FBS and 1% P/S, with the different treatments was added to every well. A concentration of 10 ug/mL of SEC-EV total protein containing *VEGF-A* mRNA was used for treatments. Human Recombinant VEGF-A protein at 20 ng/mL was used as a positive control, and PBS was used as negative control. Maintenance media and treatments were refreshed every other day for 12 days. After 12 days, aortic rings were fixed and stained for α-SMA (Dako, M0851) and CD31 (R&D Systems, AF3628) as described ^41^.

### Confocal images acquisition

The confocal images of the aortic rings stored in Falcon 96-wellplate format were captured on an inverted Zeiss LSM880 confocal microscope equipped with Airyscan detector using Plan-Apochromat 20x/0.8 M27 air objective lens. 1 field of view image was acquired from each well and performed in triplicate with total 3 different wells per biological condition. DAPI, CD31, and α smooth muscle actin (α SMA) were imaged using 405 nm, 488 nm, and 633 nm excitation lasers, respectively. The scanning acquisition parameters are 16-bit depth, 532 µm by 532 µm x-by-y frame resolution and 8 µm z-stack thickness. The total scanning thickness is ∼250 μm.

### Images 3D volumetric projection

For presentation and qualitative display, the .czi file images obtained from Zeiss LSM880 confocal microscope were extracted and 3D-projected with Imaris version 9.0.1 (Oxford Instruments, Bitplane AG, Switzerland).

### Images skeleton quantitative analysis

The quantitative analysis of the images to determine the total branches, total junctions and branch average lengths was performed with skeleton analysis in ImageJ version1.53f51 (National Institute of Health, USA). Firstly, the multichannel .czi images were splitted into each channel. The analysis was focused only on the 488 nm/green channel (CD31) and 633 nm/far red channel (α SMA). These images were then converted into 8-bit image, morphology-processed into 2-pixel circle open, and then were stacked as z-projection sum slices. The skeletonize plugin was then chosen with the lowest intensity voxel, no prune ends elimination, largest shortest path calculated and labeled skeletons displayed. The final numerical data was eventually obtained as .csv files.

### Transcriptomic analysis of mouse-heart tissue after direct injection of *VEGF-A mRNA* via EVs or LNPs

The transcriptomic analysis of mouse heart tissue after direct injection of *VEGF-A* mRNA via EVs or LNPs involved obtaining heart tissue biopsies following injection of VEGF-A mRNA via HTB-EVs, CPC-EVs, HUVEC-EVs, or LNPs, with untreated mice as controls. Total RNA was *isolated*, and RNA-seq was performed using NovaSeq6000. The DESeq2 R-package was utilized for differential expression analysis, and explorative data analysis included PCA, Volcano plots, and Heatmap plots. Venn diagrams were employed to compare DEGs, and functional analysis was conducted on unique DEGs. Gene ontology, Ingenuity Pathway Analysis (IPA), and enrichment analysis with Enrichr were performed to identify associated pathways, diseases, and functions. Raw and processed data are available at Gen Expression Omnibus (GSE220060).

### RNA-seq

Library construction was performed using Takara SMARTer Stranded Total RNA-Seq Kit - Pico Input Mammalian kit - V3, which is specifically designed for very low input total RNA samples. Clustering was done by ’cBot’ and samples were sequenced on NovaSeq6000 (NovaSeq Control Software 1.7.5/RTA v3.4.4) with a 151nt(Read1)-19nt(Index1)-10nt(Index2)-151nt(Read2) setup using ’NovaSeqXp’ workflow in ’S4’ mode flowcell. The Bcl to FastQ conversion was performed using bcl2fastq_v2.20.0.422 from the CASAVA software suite. The quality scale used is Sanger / phred33 / Illumina 1.8+. Processing of FASTQ files was carried out by the SciLifeLab National Genomics Infrastructure at the Uppsala Multidisciplinary Center for Advanced Computational Science, Sweden. The sequenced reads were quality controlled with the FastQC software and pre-processed with Trim Galore. The processed reads were then aligned to the reference genome of Mus musculus (build GRCm38) with the STAR aligner. Read counts for genes were generated using the feature Counts library and normalized TPM values calculated with StringTie, and raw gene read counts were generated by Salmon. Technical documentation on the RNA-seq pipeline can be accessed here: https://github.com/nf-core/rnaseq. Raw and processed data are available for download at Gen Expression Omnibus (https://www.ncbi.nlm.nih.gov/geo/) accession number: GSE220060.

### Differential expression analysis

The raw gene count data generated from the Salmon tool including 60,669 transcripts from 3 tissue samples each from HTB-, HUVEC-, CPC-, LNP-*VEGF-A* mRNA treated mice were imported into R for bioinformatic analysis, and statistical testing for differential expression was carried out using the DESeq2 R-package41. Filtering and normalization of the raw counts were performed for EVs of each cell line separately within the DESeq() function in the DESeq2 package. The Wald test was used for the identification of differentially expressed genes (DEGs). P-values were adjusted for multiple testing using the Benjamin Hoch method and an adjusted-p-value (adjP) of ≤ 0.05 was considered statistically significant. A log2 fold change (log2FC) shrinkage was carried out using ashr shrinkage estimator42, to reduce the variability of the lowly expressed genes. A result table with log2FC, p-values and adjusted p-values was generated. Genes with adjP ≤ 0.05 and absolute log2FC > 1 are considered significant and are used for downstream functional and pathway analysis.

### Explorative data analysis

The gene expression dataset was further analyzed to investigate the transcriptional effect of the injection of three different vesicles with *VEGF-A* mRNA into heart tissues. The genes with no counts were filtered and the data were normalized for each cell line using the varianceStabilizingTransformation() function in the DESeq2 package.

Principal component analysis (PCA) plots were generated from the top 500 most variable genes using plotPCA() function. The two axes PC1 and PC2 represent the two principal components identified by the analysis. PC1 contributes to 37% of the overall variation among samples and 21 % for PC2. Additionally, the Volcano plots were generated using EnhancedVolcano R package.

Red and blue dots denote the significant up- and down-regulated genes passing adjusted P value and fold difference thresholds (-log10 of adjP-value ≥ 1.3, abs(logFC)>1).

MA plots were generated using plotMA() function with a shrinkage estimator from ashr package42. The output from the logFC shrinkage was used for visualization in MA plots. The top 1,000 most variable genes were selected, and samples were clustered and visualized using heatmaps to assess the reproducibility and quality of the experiment. The pheatmap R-package was used to create the heatmaps with spearman rank correlation as a distance measure.

### Venn diagram

The DEGs between the untreated control mice and the mice injected with different EVs or LNPs were compared using VennDiagram R-package to investigate the overlapped and unique DEGs in the heart tissue43. Functional analysis was carried out on the 33 unique DEGs from the CPC treatment sample using Enrichr.

### DEGs associated with inflammatory response and angiogenesis – gene ontology

A total of 364 and 266 unique Ensembl gene IDs associated with inflammatory response (GO:006954) and angiogenesis (GO:0001525) were identified using biomaRt R package, respectively. The gene IDs associated with the two GO terms were compared to the IDs of DEGs to identify how many of the DEGs are associated with inflammatory response and angiogenesis.

### Ingenuity pathway analysis

The DEGs with p ≤ 0.05 and absolute log2FC > 1 were input into Ingenuity Pathway Analysis (IPA) software 43. The analysis was carried out using the Core- and Comparison-Analysis functions. The list of top canonical pathways, top diseases and functions, and the most relevant networks associated to the expression of DEGs were obtained. IPA displays significant canonical pathways and associated diseases and functions with p-values calculated by Right-Tailed Fisher’s Exact Test, P ≤ 0.05 was deemed statistically significant. IPA uses Z-scores to predict the activity of canonical pathways. A positive Z-score suggests an activation of a pathway while a negative Z-score suggests an inhibition. an absolute Z-score of ≥ 2 is considered significant.

### Enrichment analysis with Enrichr

The gene symbols of the DEGs were input into Enrichr44-46. The Enrichr output containing the top 10 biological processes Gene Ontology (2021 version) and top 10 MGI Mammalian Phenotype Level 4 (2021 version) ranked by p-values were selected. Bar graphs with the top enriched terms and phenotypes displaying log-P-value were created and exported.

### Statistical analysis

The statistical analysis was performed by GraphPad Prism v.9 (GraphPad Software). The statistically significant differences for in vitro and in vivo data were evaluated by One-Way ANOVA and the Kruskal-Wallis test followed by the Dunn’s multiple comparison test by comparing treated and untreated samples. The individual statical parameter applied for different datasets is mentioned in the individual figure legends. Additionally, for RNA-Seq data the statistical analysis for DEGs was carried out using the DESeq2 R-package41. Filtering and normalization of the raw counts were performed for each EV type (in tissue) together within the DESeq() function in the DESeq2 package. The Wald test was used for the identification of DEGs. P-values were adjusted for multiple testing using the Benjamin Hoch method. The differentially expressed genes with false discovery rate (FDR) rate of ≤ 0.05 and absolute log2 FC > 1 were considered statistically significant. A result table with log2 fold changes, p-values, and FDR-adjusted p-values was generated and used for creating graphs.

## List of materials and reagents

### Cell lines and reagents

Human epithelial HTB-177 cells from NCI-H460, ATCC.

Human umbilical vein endothelial cells (HUVECs) from Lonza, Switzerland.

iCell cardiac progenitor cells (CPCs) from R1093, Fujifilm Cellular Dynamics, Madison, WI, USA.

RPMI-1640 growth medium from Sigma Aldrich.

1% of L-glutamine and 1% penicillin-streptomycin from Thermo Fisher Scientific. Fetal bovine serum (FBS) from Sigma Aldrich.

Human umbilical vein endothelial cells (HUVECs) from Lonza, Switzerland.

### mRNAs constructs and source

The CDS sequence of *VEGF-A* and *eGFP* mRNA provided to TriLink® Biotechnologies (CA, USA).

mRNA encoding firefly luciferase (FLuc-mRNA, Trilink Biotechnologies, USA).

### Reagents for LNP characterization

NanoAssemblr from Precision NanoSystems, Inc., BC, Canada.

Slide-a-lyzer G2 dialysis cassette (10000 MWCO, Thermo Fisher Scientific Inc.).

0.22 µm sterile filter from Gillex, Merck.

ZetaSizer from Malvern Instruments Inc.

Quant-it Ribogreen Assay Kit from Thermo Fisher Scientific.

### Chromatography and sample concentrates

qEV-70/10 mL chromatography columns from Izon Science Ltd, New Zealand.

Amicon ultra/15mL centrifuge filters from cat: # UFC903024, Sigma Aldrich, now Merck.

### Reagents for *VEGF-A* mRNA and protein analysis

miRNeasy Mini Kit (Qiagen, cat. #: 217004). Qubit 2.0 fluorometer (Thermo Fisher Scientific). NanoDrop 1000 (Thermo Fisher Scientific).

cDNA kit with RNase inhibitor (Thermo Fisher Scientific: 4374966).

mRNA quantification using hydrolysis probes (TaqMan probe assay, Thermo Fisher Scientific, assay ID: Hs00900055_m1)

*VEGF-A* mRNA standards (TriLink Biotechnologies, USA). GAPDH gene (Thermo Fisher Scientific, assay ID: Hs02758991_g1).

Human VEGF-A sandwich ELISA Kit (cat.#: RAB0507, Sigma Aldrich, now Merck). ELISA reader instrument (Spectra max, 340PC, molecular devices).

Nanoparticle Tracking Analysis (NTA) with an LM14c instrument from Malvern Panalytical, UK.

NanoSight Fluorescent NTA LM14c software version 3.2 (Malvern Panalytical, UK).

The CD63 isolation/detection reagent for cell culture medium (Thermo Fisher Scientific, cat.#: 10606D)

Proteases inhibitors (Thermo Fisher Scientific) in the Tissue LyserII (Qiagen)

### Antibodies

Mouse anti-human PE-CD9 antibody (BD Pharmingen™, cat.#: 555372)

The CD63 isolation/detection reagent (Thermo Fisher Scientific, cat.#: 10606D)

α-SMA (Dako, M0851)

CD31 (R&D Systems, AF3628)

### Other

Transmission electron microscope FEI Tecnai G2 Spirit (Thermo Fisher Scientific, Oregon, USA) Radius software (Version 2.1) with a Xarosa digital camera (EMSIS GmbH, Münster, Germany)

T-PER Tissue Protein Extraction Reagent (Thermo Fisher Scientific Cat. 78510)

## Data and materials availability

All data associated with this study are presented in the main paper or the Supplementary Materials. The source data are available on request from the corresponding author. The RNA-Seq data was deposited to NCBI repository with following accession number below.

## Accession numbers

The Raw and processed data of transcriptomics are available for download at Gene Expression Omnibus, NCBI (https://www.ncbi.nlm.nih.gov/geo/) accession number: GSE220060.

**Token for reviewers and editors:** To review GEO accession GSE220060: Go to https://www.ncbi.nlm.nih.gov/geo/query/acc.cgi?acc=GSE220060 and enter token **qvatsqokrlyrdeb** into the box

## Supplementary and supporting materials

Uploaded as additional files.

## Acknowledgements

This work has been supported by grants from the Swedish research council (VR), the Swedish Foundation of Strategic Research (SSF) in the Industrial Research Centre (FoRmulaEx – Nucleotide Functional Drug Delivery (IRC15-0065), the Swedish governmental agency for innovation systems (VINNOVA, 2017-02960), the Swedish Knowledge Foundation (20200014), the National Genomics Infrastructure in Stockholm funded by Science for Life (SciLife) Laboratory, and SNIC/Uppsala Multidisciplinary Center for Advanced Computational Science for assistance with massively parallel sequencing and access to the UPPMAX computational infrastructure, and the funding received by F.K. from the European Union’s Horizon 2020 research and innovation program under the Skłodowska Curie grant agreement No 814316. Moreover, we acknowledge Mr. Mario Soriano Navarro at the Responsable Servicio Microscopía Electrónica, Valencia Spain, for technical assistance.

## Author contributions

Conceptualization: HV

Methodology: MN, SH-H, BT, JW, FK, HGG, YJ, ZP, KJ, LH, LL, HV

Investigation: MN, SH-H, LL, JS, HV

Data analysis: MN, BT, LH, LL, JS, HV

Validation: HV & MN

Visualization: HV & MN

Funding acquisition: HV

Project administration: HV

Supervision: HV

Writing – original draft: HV and MN wrote the first draft with the input from all the co-authors.

Writing – review & editing: with the input from all the co-authors.

## Supplementary Figure Legends

**Supplementary Figure 1.**
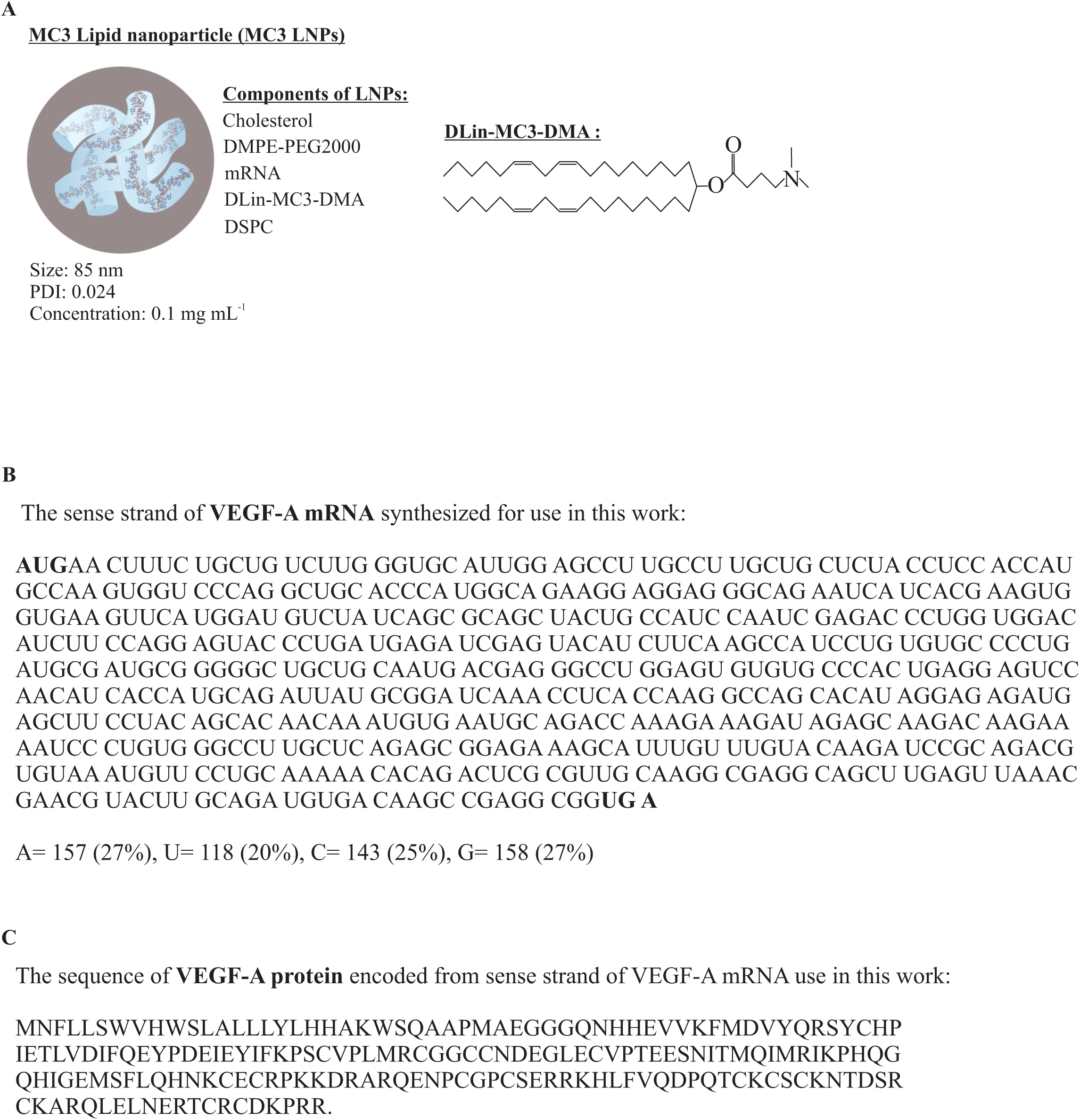
**(A)** The chemical structure and components of DLin-MC3-DMA LNPs (MC3-LNPs), and the loaded mRNA in MC3-LNPs used in the current study. **(B)** Sense strand sequence of the synthesized clean cap *VEGF-A* mRNA isoform 11, used in the current study. **(C)** Sequence of VEGF-A protein encoded from VEGFA sense strand used in the current study. PDI: polydispersity index.

**Supplementary Figure 2.**
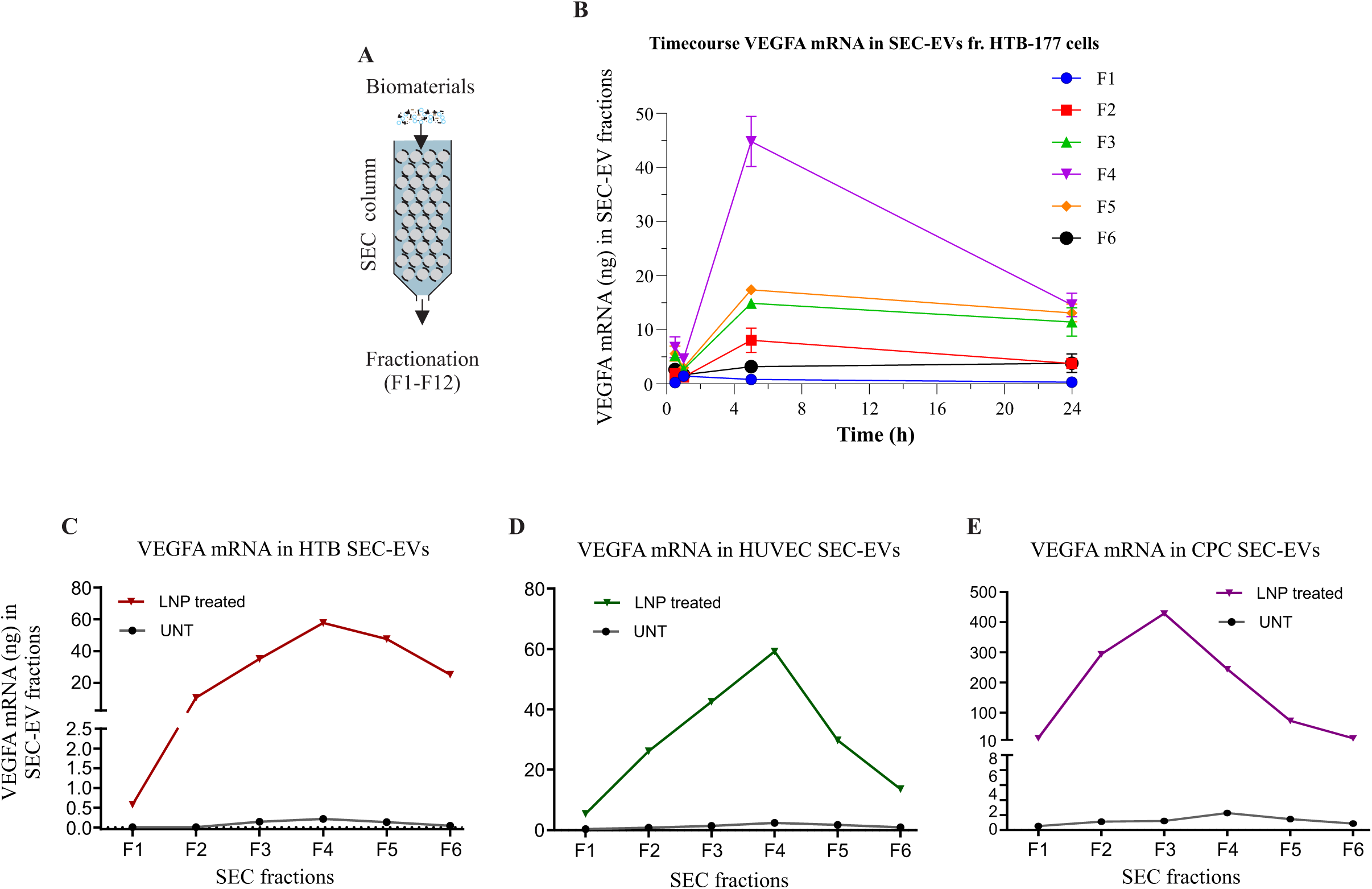
Isolation and quantification of *VEGF-A* mRNA in different EV types and time course. **(A)** 15 mL of conditioned media was loaded onto qEV/10 SEC column, and after discarding the void volume, 12 fractions of SEC-EVs were collected from sample collected at different timepoints after treatment. **(B)** Time course of *VEGF-A* mRNA secretion and quantification by qPCR in HTB-SEC EVs. Since mRNA was present only in fractions F1-6 only 6 fractions were examined. **(C-E)** *VEGF-A* mRNA in CPC-EVs, and HUVEC EVs and HTB-EVs. SEC fractions of untreated cells were used as controls, where no or negligible amounts of *VEGF-A* mRNA were detected.

**Supplementary Figure 3.**
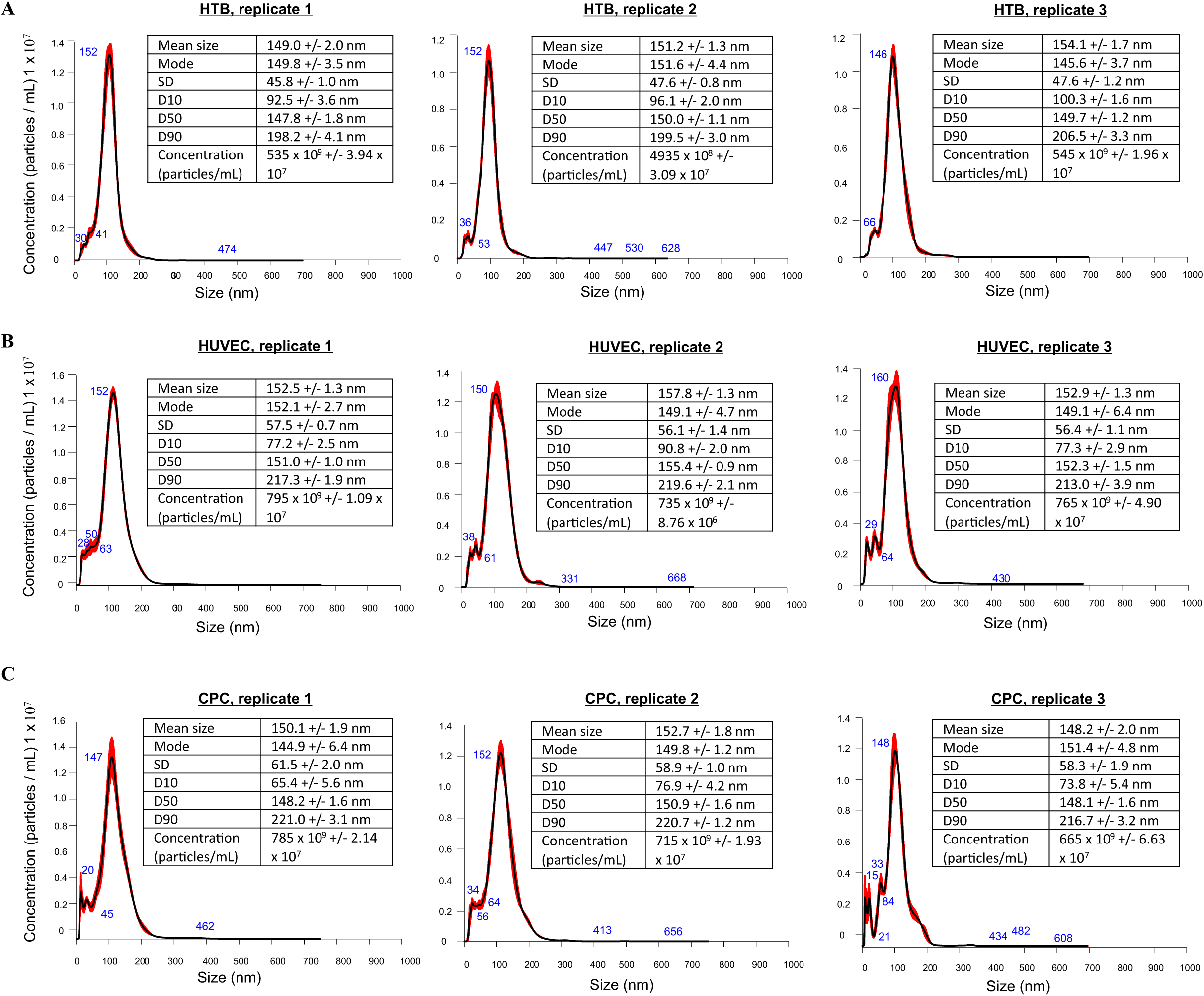
Size determination of EVs. Nanoparticle tracking analyzer (NTA)-based size distribution and numeration of (**A)** HTB SEC-EVs **(B)** HUVEC SEC-EVs, and **(C)** CPC SEC-EVs. Samples were diluted 10 times. The graphs were obtained from diluted samples, and the values were adjusted with factor 10. The adjusted values for dilution factor are presented in tables with each graph.

**Supplementary Figure 4.**
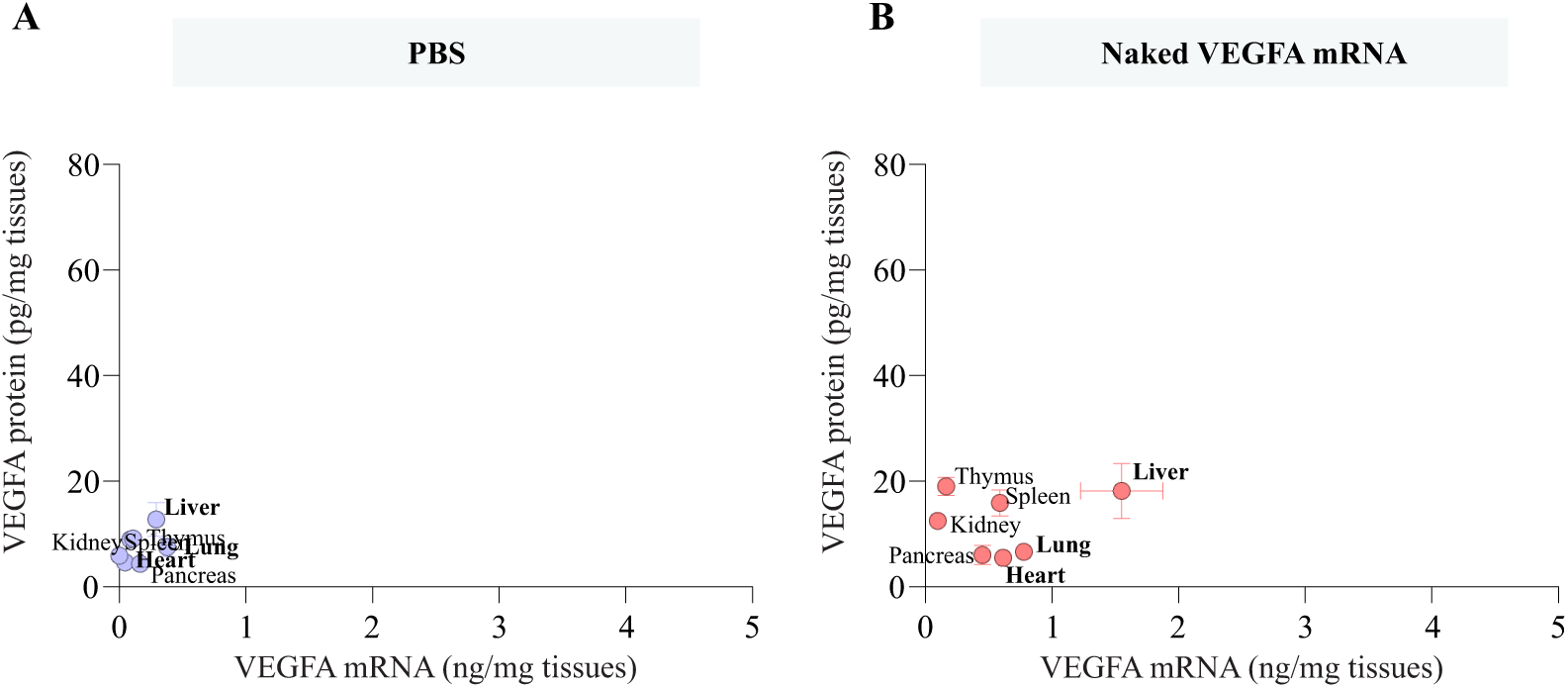
Organ specific localization of produced protein after delivery naked mRNA without a vehicle. Quantification of VEGF-A protein in organs after intravenous administration of **(A)** PBS or **(B)**naked *VEGF-A* mRNA.

**Supplementary Figure 5.**
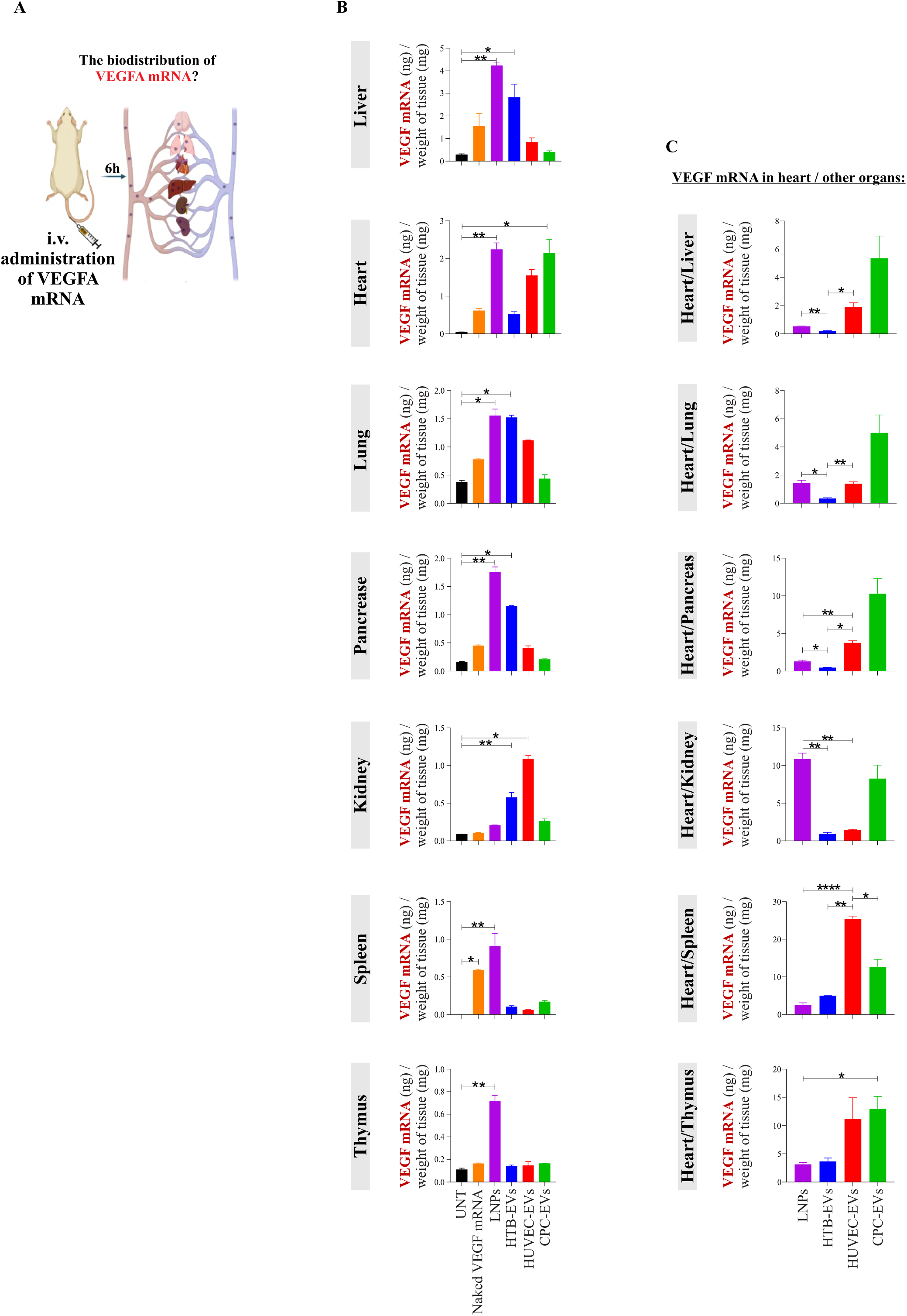
Biodistribution of *VEGF-A* mRNA after intravenous delivery via EVs or LNPs. **(A)** 1 µg of *VEGF-A* mRNA was delivered to C57BL/6Ncrl mice intravenously via CPC-EVs, or HTB-EVs or HUVEC-EVs, or LNPs. The mice were sacrificed 6 h after administration, and organs were collected and biodistribution of *VEGF-A* mRNA was examined. **(B)** A comparison for *VEGF-A* mRNA between different EV types i.e, CPC-EVs, HTB-EVs and HUVEC-EVs or LNPs within an individual organ. **(C)** Ratio between mRNA levels detected in heart/by other organs to calculate relative amounts of *VEGF-A* mRNA in heart. One-Way ANOVA test was applied to compare the statistical differences between untreated samples (n=3), and treated samples, or between each vehicle (n=3, each group). *p < 0.05, **p < 0.01, ****p < 0.0001. UNT: untreated, HTB-EVs: HTB-177 lung epithelial cell-derived EVs, HUVEC-EVs: human umbilical vein endothelial cell-derived EVs, CPC-EVs: cardiac progenitor cell-derived-EVs.

**Supplementary Figure 6.**
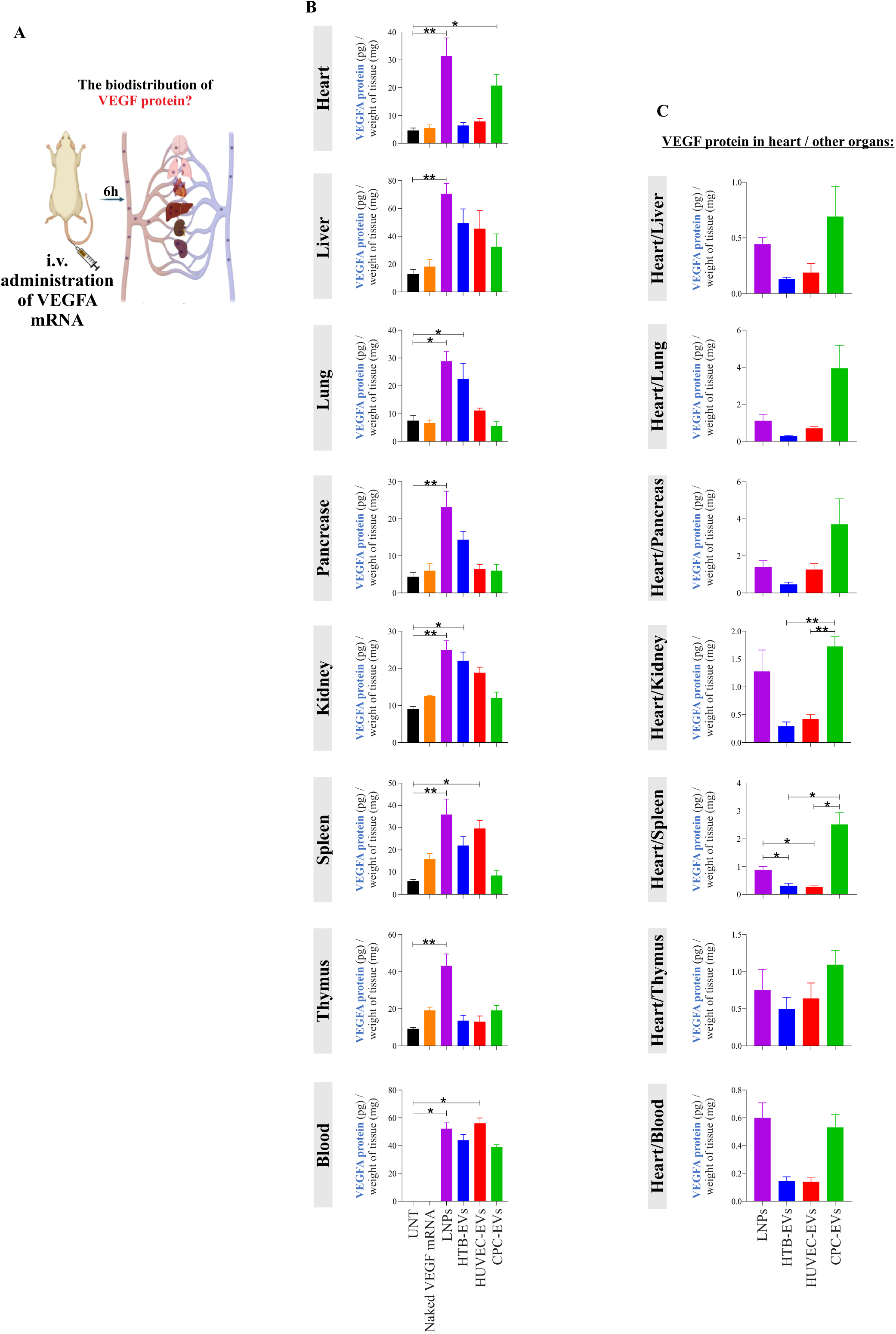
Levels of VEGF-A protein in different organs after intravenous delivery of *VEGF-A* mRNA via EVs or LNPs. **(A)** Delivery of *VEGF-A* mRNA via CPC-EVs, or HTB-EVs or HUVEC-EVs or LNPs and examination of its protein levels detected in different organs. **(B)** A comparison of VEGF-A protein between delivery by different EV types i.e. CPC-EVs, HTB-EVs, HUVEC-EVs or LNPs within an individual organ. **(C)** Ratio between VEGF-A protein levels detected in heart/by other organs to calculate relative amounts of protein in the heart. One-Way ANOVA test was applied to compare the statistical differences between untreated samples (n=3), and treated samples, or between each vehicle (n=3, each group). *p < 0.05, **p < 0.01. UNT: untreated, HTB-EVs: HTB-177 lung epithelial cell-derived EVs, HUVEC-EVs: human umbilical vein endothelial cell-derived EVs, CPC-EVs: cardiac progenitor cell-derived-EVs.

**Supplementary Figure 7.**
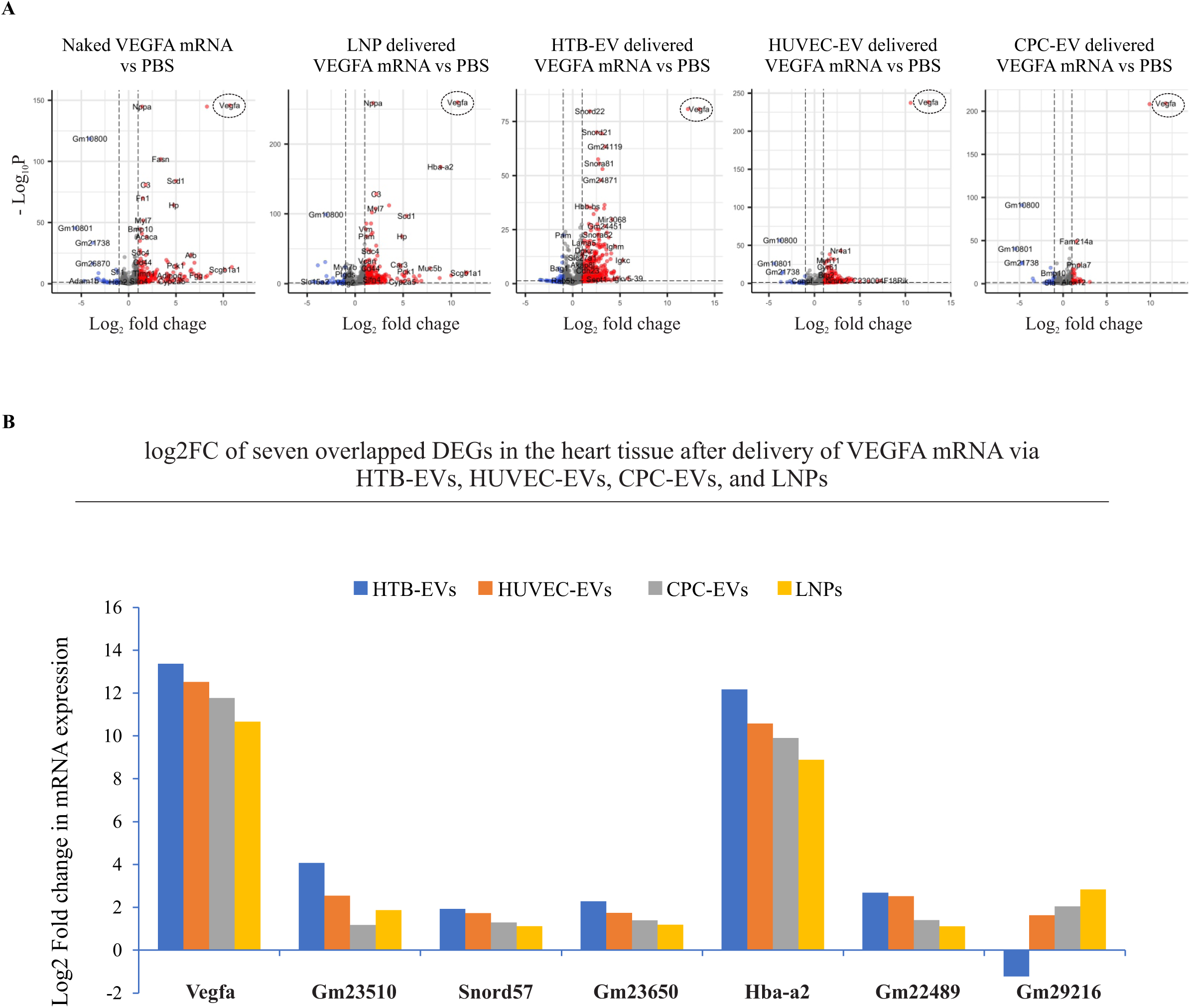
**(A)** Volcano plots show dysregulated genes in the tissue after the injection, identified using transcriptomic data. Red and blue dots denote the significant up- and down-regulated genes passing adjusted P value and fold difference thresholds (-log10 of adjP-value ≥ 1.3, abs(logFC)>1). **(B)** Shared genes between CPC-EVs, HTB-EVs, HUVEC-EVs and LNPs after *VEGF-A* mRNA delivery to heart tissue. Log2 fold changes (log2FC) in mRNA expression are shown. CPC-EVs: cardiac progenitor cell-derived extracellular vesicles (EVs), HTB-EVs: HTB-177 lung epithelial cell-derived EVs, HUVEC-EVs: human umbilical vein endothelial cell-derived EVs, LNPs: lipid nanoparticles.

