## supplementarydatatables for "Targeted delivery of mRNA to the heart via extracellular vesicles or lipid nanoparticles"

**Supplementary table 1)** List of deregulated genes after VEGF-A mRNA to cardiac tissue via CPC SEC-EVs (total 82 deregulated genes)

| | **baseMean** | **log2FC** | **lfcSE** | **stat** | **pvalue** | **padj** | **gene_symbol** |
|---|---|---|---|---|---|---|---|
| **Up regulated genes (52)** | | | | | | | |
| ENSMUSG00000023951 | 1422490,887 | 11,77072286 | 0,251045956 | 46,93978569 | 0 | 0 | Vegfa |
| ENSMUSG00000069917 | 161616,9778 | 9,908874648 | 0,319098499 | 31,1286012 | 9,8826E-213 | 5,0999E-209 | Hba-a2 |
| ENSMUSG00000028001 | 8,106012535 | 3,049787992 | 0,991612704 | 3,689561019 | 0,000224641 | 0,007967434 | Fga |
| ENSMUSG00000029368 | 12,61091204 | 2,297752619 | 0,754031598 | 3,694259726 | 0,000220528 | 0,007848903 | Alb |
| ENSMUSG00000101249 | 353,60672 | 2,044079118 | 0,476678817 | 4,74355865 | 2,09996E-06 | 0,000168013 | Gm29216 |
| ENSMUSG00000016255 | 5,729786908 | 1,991109257 | 0,528055277 | 4,291303002 | 1,77628E-05 | 0,001007305 | Tubb1 |
| ENSMUSG00000064938 | 18,99974932 | 1,934532464 | 0,378809954 | 5,49940134 | 3,81083E-08 | 5,31508E-06 | Mir3068 |
| ENSMUSG00000021950 | 5,98199759 | 1,841612569 | 0,574066462 | 3,814489228 | 0,000136465 | 0,005314921 | Anxa8 |
| ENSMUSG00000036833 | 47,32393391 | 1,800078164 | 0,176248471 | 10,41885157 | 2,03394E-25 | 1,74936E-22 | Pnpla7 |
| ENSMUSG00000022853 | 11,94833255 | 1,61292389 | 0,302535891 | 5,697875473 | 1,2131E-08 | 1,84123E-06 | Ehhadh |
| ENSMUSG00000064372 | 2887,578015 | 1,543342378 | 0,529276087 | 3,581231276 | 0,000341979 | 0,011099254 | mt-Tp |
| ENSMUSG00000034858 | 131,3496456 | 1,511231647 | 0,099330274 | 15,34805029 | 3,64992E-53 | 9,41771E-50 | Fam214a |
| ENSMUSG00000080463 | 27,49171493 | 1,408826433 | 0,475950855 | 3,618238439 | 0,000296615 | 0,009939494 | Gm22489 |
| ENSMUSG00000065289 | 27,44229996 | 1,397258918 | 0,323126687 | 4,783995831 | 1,71844E-06 | 0,000144195 | Gm23650 |
| ENSMUSG00000023912 | 11,54718517 | 1,32165868 | 0,337843337 | 4,404795101 | 1,05884E-05 | 0,00066232 | Slc25a27 |
| ENSMUSG00000040270 | 10,30985622 | 1,307731325 | 0,331244224 | 4,438842355 | 9,0444E-06 | 0,000572683 | Bach2 |
| ENSMUSG00000096349 | 5608,526165 | 1,304271113 | 0,170682629 | 7,894016329 | 2,92614E-15 | 1,31307E-12 | Gm22513 |
| ENSMUSG00000034664 | 7,511870281 | 1,297248128 | 0,391984524 | 3,89331667 | 9,88829E-05 | 0,004030008 | Itga2b |
| ENSMUSG00000065272 | 23,23785163 | 1,293982409 | 0,298521008 | 4,779753244 | 1,7551E-06 | 0,000146084 | Snord57 |
| ENSMUSG00000036528 | 31,48222706 | 1,28749118 | 0,190414895 | 7,016661916 | 2,27232E-12 | 6,17173E-10 | Ppfibp2 |
| ENSMUSG00000065881 | 27,97377145 | 1,28687561 | 0,532480385 | 3,207885108 | 0,001337149 | 0,030001555 | Gm22866 |
| ENSMUSG00000047153 | 46,8426656 | 1,284395703 | 0,144280139 | 9,115117133 | 7,85855E-20 | 4,77107E-17 | Khnyn |
| ENSMUSG00000065147 | 13,67781635 | 1,26031476 | 0,373627163 | 3,945215745 | 7,97282E-05 | 0,00341699 | Snora31 |
| ENSMUSG00000000320 | 5,595761394 | 1,207894494 | 0,443703062 | 3,424039698 | 0,000616976 | 0,01685303 | Alox12 |
| ENSMUSG00000088529 | 40,27254091 | 1,194378008 | 0,285580053 | 4,635786238 | 3,55583E-06 | 0,000262141 | Gm26083 |
| ENSMUSG00000087708 | 8,925759364 | 1,194190701 | 0,50223483 | 3,194859968 | 0,001398987 | 0,030786663 | Gm23000 |
| ENSMUSG00000032942 | 68,53435306 | 1,180112634 | 0,150271211 | 8,069987389 | 7,03054E-16 | 3,45534E-13 | Ucp3 |
| ENSMUSG00000065017 | 10,3683213 | 1,179544129 | 0,569057722 | 3,009285981 | 0,002618625 | 0,04775058 | Gm23510 |
| ENSMUSG00000053965 | 12,1641521 | 1,156674104 | 0,312951689 | 4,210113993 | 2,55242E-05 | 0,001386501 | Pde5a |
| ENSMUSG00000065905 | 12,78207471 | 1,130585998 | 0,400973393 | 3,500442584 | 0,000464486 | 0,013775756 | Gm26110 |
| ENSMUSG00000095676 | 64,96407058 | 1,130369297 | 0,370432885 | 3,671322204 | 0,000241299 | 0,008499816 | Gm25099 |
| ENSMUSG00000064925 | 34,95436303 | 1,12636159 | 0,256714038 | 4,812923808 | 1,48738E-06 | 0,000130095 | Snora62 |
| ENSMUSG00000077704 | 7,934006932 | 1,122302974 | 0,522394256 | 3,037625449 | 0,002384501 | 0,044409839 | Snord89 |
| ENSMUSG00000091089 | 5,768493625 | 1,112011527 | 0,420790231 | 3,36341728 | 0,000769839 | 0,020115212 | A930018M24Rik |
| ENSMUSG00000065904 | 7,655516212 | 1,102896574 | 0,433172595 | 3,293060421 | 0,000991032 | 0,024180706 | Gm26109 |
| ENSMUSG00000105884 | 21,26150902 | 1,098295834 | 0,392857026 | 3,47219494 | 0,000516221 | 0,014966069 | SNORA50 |
| ENSMUSG00000053964 | 140,7923853 | 1,093273376 | 0,121523929 | 9,201259301 | 3,53779E-20 | 2,35353E-17 | Lgals4 |
| ENSMUSG00000053310 | 8,117702539 | 1,086110865 | 0,481456111 | 3,102467811 | 0,001919144 | 0,038914513 | Nrgn |
| ENSMUSG00000063011 | 6,400087865 | 1,077675856 | 0,476576237 | 3,109416364 | 0,001874573 | 0,038236112 | Msln |
| ENSMUSG00000093843 | 43,51834069 | 1,072148089 | 0,217953879 | 5,303135516 | 1,1383E-07 | 1,4503E-05 | Gm25939 |
| ENSMUSG00000106585 | 15,6136049 | 1,060009201 | 0,340372021 | 3,722161393 | 0,000197525 | 0,007178354 | SNORA17 |
| ENSMUSG00000032633 | 50,1558344 | 1,055535049 | 0,166924395 | 6,599248716 | 4,13247E-11 | 9,91888E-09 | Flcn |
| ENSMUSG00000064453 | 105,7179131 | 1,051899657 | 0,141225182 | 7,697639871 | 1,38602E-14 | 4,76838E-12 | Snord21 |
| ENSMUSG00000092730 | 32,1211032 | 1,049047358 | 0,249759654 | 4,646833712 | 3,37069E-06 | 0,00025028 | Snora24 |
| ENSMUSG00000065353 | 28,89440024 | 1,047568345 | 0,284159436 | 4,200230156 | 2,66644E-05 | 0,001433349 | Snora73b |
| ENSMUSG00000003282 | 9,079643453 | 1,040109182 | 0,358542625 | 3,54823806 | 0,000387818 | 0,012315891 | Plag1 |
| ENSMUSG00000027875 | 13,01643593 | 1,032916903 | 0,282937019 | 4,175504804 | 2,97326E-05 | 0,001557716 | Hmgcs2 |
| ENSMUSG00000028150 | 25,74438622 | 1,021811406 | 0,2089804 | 5,263357999 | 1,41448E-07 | 1,75889E-05 | Rorc |
| ENSMUSG00000080478 | 11,41314764 | 1,021502886 | 0,425748755 | 3,188131322 | 0,001431955 | 0,031378357 | Snord23 |
| ENSMUSG00000065794 | 30,64566922 | 1,00884555 | 0,27371114 | 4,193760429 | 2,74368E-05 | 0,001467228 | Gm25514 |
| ENSMUSG00000032175 | 11,03292056 | 1,004668792 | 0,346699572 | 3,540928001 | 0,000398722 | 0,012508248 | Tyk2 |
| ENSMUSG00000077167 | 42,54522666 | 1,002299507 | 0,224733427 | 4,876897521 | 1,07767E-06 | 9,75673E-05 | Gm24119 |

| | | | | | | | |
|---|---|---|---|---|---|---|---|
| **Down regulated genes (30)** | | | | | | | |
| ENSMUSG00000075015 | 224,3024598 | -5,478498618 | 0,396230832 | -13,97251585 | 2,29378E-44 | 3,94569E-41 | Gm10801 |
| ENSMUSG00000095280 | 42,34764648 | -4,771371596 | 0,44137211 | -10,99348868 | 4,10737E-28 | 5,29901E-25 | Gm21738 |
| ENSMUSG00000097312 | 378,5215611 | -4,719179103 | 0,444464012 | -10,80469149 | 3,27055E-27 | 3,38E-24 | Gm26870 |
| ENSMUSG00000075014 | 192,3854893 | -4,705152025 | 0,227949796 | -20,73477846 | 1,68228E-95 | 5,7876E-92 | Gm10800 |
| ENSMUSG00000095547 | 11,90461058 | -3,479961598 | 0,633009185 | -5,839629888 | 5,23169E-09 | 8,85185E-07 | Gm10719 |
| ENSMUSG00000046958 | 5,988228091 | -3,351826241 | 0,876988226 | -4,287640409 | 1,80581E-05 | 0,001018458 | 4930432E11Rik |
| ENSMUSG00000061601 | 5,988481954 | -1,721973372 | 0,662439073 | -3,282830754 | 0,001027703 | 0,024840579 | Pclo |
| ENSMUSG00000025355 | 15,05107335 | -1,587219556 | 0,461179802 | -3,977474104 | 6,96512E-05 | 0,003098578 | Mmp19 |
| ENSMUSG00000044768 | 17,56404763 | -1,578971162 | 0,374195214 | -4,663738004 | 3,10517E-06 | 0,000235465 | D1Ertd622e |
| ENSMUSG00000022372 | 6,554582313 | -1,498280723 | 0,650748395 | -3,052613739 | 0,002268577 | 0,04343968 | Sla |
| ENSMUSG00000064357 | 6156,972475 | -1,432805679 | 0,152206234 | -9,640815578 | 5,37591E-22 | 3,9632E-19 | mt-Atp6 |
| ENSMUSG00000025591 | 6,897700688 | -1,32118906 | 0,577049435 | -3,055725579 | 0,002245166 | 0,043117185 | Tma16 |
| ENSMUSG00000092201 | 19,66687913 | -1,304560041 | 0,328740417 | -4,440569206 | 8,97212E-06 | 0,000571613 | A530058N18Rik |
| ENSMUSG00000089940 | 6,049083508 | -1,280156678 | 0,56056686 | -3,072736802 | 0,002121055 | 0,041539669 | Gm4117 |
| ENSMUSG00000029380 | 9,977435053 | -1,275665799 | 0,562665061 | -3,038290839 | 0,002379242 | 0,044409839 | Cxcl1 |
| ENSMUSG00000075324 | 14,84775228 | -1,186185282 | 0,396704773 | -3,609269874 | 0,00030706 | 0,010157585 | Fign |
| ENSMUSG00000026580 | 19,71991735 | -1,175981796 | 0,426357218 | -3,396221717 | 0,00068323 | 0,018300361 | Selp |
| ENSMUSG00000022868 | 12,8943545 | -1,172305659 | 0,468050014 | -3,216187489 | 0,001299059 | 0,029519311 | Ahsg |
| ENSMUSG00000103560 | 11,47958481 | -1,168882847 | 0,462483664 | -3,23061102 | 0,001235259 | 0,0285215 | Gm38070 |
| ENSMUSG00000105050 | 9,322572717 | -1,158521531 | 0,450876151 | -3,277478561 | 0,001047387 | 0,025198321 | RP23-458G12.2 |
| ENSMUSG00000067279 | 41,22263115 | -1,137894672 | 0,198274221 | -6,059969228 | 1,36148E-09 | 2,60218E-07 | Ppp1r3c |
| ENSMUSG00000030046 | 162,9789317 | -1,117367208 | 0,144340716 | -7,978820871 | 1,47738E-15 | 6,93092E-13 | Bmp10 |
| ENSMUSG00000057134 | 10,03073497 | -1,109539559 | 0,475476394 | -3,099175537 | 0,0019406 | 0,039272418 | Ado |
| ENSMUSG00000073902 | 9,975820698 | -1,069776145 | 0,467604644 | -3,058097382 | 0,002227472 | 0,042971469 | Gm1966 |
| ENSMUSG00000046709 | 14,19564364 | -1,036891099 | 0,384085962 | -3,363896774 | 0,000768503 | 0,020115212 | Mapk10 |
| ENSMUSG00000025511 | 17,73847242 | -1,035371121 | 0,36381815 | -3,484209462 | 0,000493593 | 0,014390886 | Tspan4 |
| ENSMUSG00000026822 | 33,98022017 | -1,027710409 | 0,237065407 | -4,769713913 | 1,84488E-06 | 0,000151119 | Lcn2 |
| ENSMUSG00000064367 | 8607,413997 | -1,025362899 | 0,166713381 | -6,461817095 | 1,03453E-10 | 2,27179E-08 | mt-Nd5 |
| ENSMUSG00000019122 | 16,51217109 | -1,018199088 | 0,432508433 | -3,095852282 | 0,001962481 | 0,039406161 | Ccl9 |
| ENSMUSG00000001018 | 10,74863618 | -1,001472063 | 0,414327372 | -3,149132706 | 0,001637558 | 0,0347048 | Snapin |

| **Supplementary table 2)** List of deregulated genes after VEGF-A mRNA to cardiac tissue via HTB SEC-EVs (total 445 genes) | | | | | | | |
|---|---|---|---|---|---|---|---|
| | **baseMean** | **log2FC** | **lfcSE** | **stat** | **pvalue** | **padj** | **gene_symbol** |
| **Up regulated genes (399)** | | | | | | | |
| ENSMUSG00000023951 | 1422490,887 | 13,3705963 | 0,25104582 | 53,3071116 | 0 | 0 | Vegfa |
| ENSMUSG00000069917 | 161616,9778 | 12,1738261 | 0,31909418 | 38,2142472 | 0 | 0 | Hba-a2 |
| ENSMUSG00000076569 | 1,374851203 | 5,82020915 | 1,60304988 | 3,81984314 | 0,000134 | 0,0036636 | Igkv5-39 |
| ENSMUSG00000076609 | 8,962699561 | 5,28967014 | 0,74154796 | 7,40725546 | 1,29E-13 | 1,703E-11 | Igkc |
| ENSMUSG00000080461 | 2,004943033 | 4,72464879 | 1,33593691 | 4,0814359 | 4,48E-05 | 0,0014547 | Gm22488 |
| ENSMUSG00000070167 | 1,539331061 | 4,57492662 | 1,40151121 | 3,82357299 | 0,000132 | 0,0036155 | Snora57 |
| ENSMUSG00000076617 | 15,54118982 | 4,48822495 | 0,49702273 | 9,2564409 | 2,11E-20 | 5,73E-18 | Ighm |
| ENSMUSG00000064795 | 3,958143162 | 4,36986795 | 0,86697475 | 5,42465198 | 5,81E-08 | 3,596E-06 | Gm22423 |
| ENSMUSG00000065331 | 1,763032613 | 4,36811056 | 1,55872528 | 3,52450359 | 0,000424 | 0,009537 | Gm24927 |
| ENSMUSG00000064696 | 3,345524363 | 4,3533239 | 0,84195379 | 5,54893814 | 2,87E-08 | 1,899E-06 | Gm24148 |
| ENSMUSG00000097597 | 1,625181522 | 4,33455295 | 1,40549611 | 3,67780233 | 0,000235 | 0,0058405 | Gm26674 |
| ENSMUSG00000064923 | 2,849277753 | 4,27569691 | 0,91542363 | 5,07780519 | 3,82E-07 | 2,026E-05 | Gm22042 |
| ENSMUSG00000064938 | 18,99974932 | 4,15625409 | 0,35069975 | 12,0202544 | 2,78E-33 | 2,361E-30 | Mir3068 |
| ENSMUSG00000077711 | 3,966238704 | 4,07729847 | 0,84834083 | 5,21672461 | 1,82E-07 | 1,031E-05 | AF357399 |
| ENSMUSG00000065017 | 10,3683213 | 4,07021387 | 0,50073421 | 8,37365749 | 5,59E-17 | 1,104E-14 | Gm23510 |
| ENSMUSG00000093300 | 2,782452769 | 4,06770477 | 0,85022396 | 5,18795901 | 2,13E-07 | 1,176E-05 | Gm23799 |
| ENSMUSG00000106489 | 1,392283915 | 4,05751946 | 1,76996903 | 3,10642731 | 0,001894 | 0,0309128 | SCARNA15 |
| ENSMUSG00000065344 | 9,475164474 | 3,99979346 | 0,54708305 | 7,58391075 | 3,35E-14 | 4,744E-12 | Gm25581 |
| ENSMUSG00000064899 | 12,97961924 | 3,81305103 | 0,37868879 | 10,2659235 | 1E-24 | 4,26E-22 | Snord118 |
| ENSMUSG00000065100 | 1,700773271 | 3,79949443 | 1,31005769 | 3,5669168 | 0,000361 | 0,0083268 | Gm26132 |
| ENSMUSG00000086411 | 1,715470514 | 3,69438906 | 1,37147084 | 3,38114187 | 0,000722 | 0,0146307 | Gm14236 |
| ENSMUSG00000065881 | 27,97377145 | 3,62835567 | 0,51920392 | 7,26919628 | 3,62E-13 | 4,473E-11 | Gm22866 |
| ENSMUSG00000064432 | 16,36035055 | 3,62361247 | 0,69208642 | 5,61490243 | 1,97E-08 | 1,332E-06 | Gm25587 |
| ENSMUSG00000088895 | 12,42763093 | 3,61752014 | 0,34172202 | 10,7798229 | 4,29E-27 | 2,21E-24 | Gm22519 |
| ENSMUSG00000065767 | 492,8097076 | 3,58042436 | 0,44638568 | 8,26279796 | 1,42E-16 | 2,599E-14 | Gm23849 |
| ENSMUSG00000064380 | 10,85707926 | 3,56426111 | 0,41317218 | 8,84979213 | 8,77E-19 | 2,041E-16 | Gm26448 |
| ENSMUSG00000088208 | 2,554959133 | 3,5476855 | 0,87009997 | 4,54968764 | 5,37E-06 | 0,0002178 | Gm23751 |
| ENSMUSG00000094812 | 198,8642739 | 3,53810153 | 0,728312 | 5,25680685 | 1,47E-07 | 8,461E-06 | Gm22614 |
| ENSMUSG00000093107 | 4,259545953 | 3,53296546 | 0,7222086 | 5,28662626 | 1,25E-07 | 7,279E-06 | Mir1839 |
| ENSMUSG00000064858 | 1,326556582 | 3,47880334 | 1,52793087 | 3,10101854 | 0,001929 | 0,0313378 | Snora43 |
| ENSMUSG00000037681 | 4,887815546 | 3,45954571 | 0,83571531 | 4,61664001 | 3,9E-06 | 0,0001622 | Esyt3 |
| ENSMUSG00000087775 | 1,618543338 | 3,42851716 | 1,06989747 | 3,78573769 | 0,000153 | 0,0041338 | Rprl2 |
| ENSMUSG00000064981 | 7,295363686 | 3,42688076 | 0,59134374 | 6,13090395 | 8,74E-10 | 7,462E-08 | Snora70 |
| ENSMUSG00000064949 | 5,215524998 | 3,42685419 | 0,6245477 | 5,84383462 | 5,1E-09 | 3,834E-07 | Snora61 |
| ENSMUSG00000077167 | 42,54522666 | 3,41467248 | 0,19793723 | 17,3637859 | 1,55E-67 | 3,732E-64 | Gm24119 |
| ENSMUSG00000065287 | 25,90726638 | 3,40135163 | 0,3025839 | 11,413416 | 3,58E-30 | 2,463E-27 | Gm24451 |
| ENSMUSG00000075760 | 4,551544179 | 3,40068152 | 0,75752402 | 4,92735525 | 8,34E-07 | 4,133E-05 | Gm25414 |
| ENSMUSG00000065282 | 7,16376203 | 3,39933671 | 0,53138306 | 6,70622442 | 2E-11 | 2,059E-09 | Gm24455 |
| ENSMUSG00000065016 | 13,90162065 | 3,39229863 | 0,36482008 | 9,50877424 | 1,93E-21 | 5,928E-19 | Snora3 |
| ENSMUSG00000065195 | 1,677637988 | 3,38908838 | 1,11242386 | 3,67439173 | 0,000238 | 0,0058914 | Gm26071 |
| ENSMUSG00000058207 | 6,107215412 | 3,38441485 | 1,28719848 | 3,3435194 | 0,000827 | 0,0163522 | Serpina3k |
| ENSMUSG00000065353 | 28,89440024 | 3,36415817 | 0,25621112 | 13,2787624 | 3,07E-40 | 4,033E-37 | Snora73b |
| ENSMUSG00000105884 | 21,26150902 | 3,34931908 | 0,37210089 | 9,21526206 | 3,11E-20 | 8,002E-18 | SNORA50 |
| ENSMUSG00000092819 | 5,468590285 | 3,32007766 | 0,64821943 | 5,49420982 | 3,92E-08 | 2,536E-06 | Gm23639 |
| ENSMUSG00000064604 | 25,49092606 | 3,31092478 | 0,25810533 | 12,9728329 | 1,74E-38 | 1,798E-35 | Snora44 |
| ENSMUSG00000065725 | 5,069375325 | 3,25573615 | 0,52564966 | 6,50976985 | 7,53E-11 | 7,214E-09 | Gm26165 |
| ENSMUSG00000065147 | 13,67781635 | 3,2361516 | 0,35023268 | 9,44931173 | 3,41E-21 | 9,844E-19 | Snora31 |
| ENSMUSG00000036492 | 2,759072759 | 3,22486652 | 0,94421282 | 3,97900783 | 6,92E-05 | 0,0021248 | Rnf39 |
| ENSMUSG00000065036 | 4,964269336 | 3,17390701 | 0,59635461 | 5,68019738 | 1,35E-08 | 9,564E-07 | Gm22362 |
| ENSMUSG00000105863 | 14,10540587 | 3,14396262 | 0,39741491 | 8,15706397 | 3,43E-16 | 6,192E-14 | SNORA17 |
| ENSMUSG00000093843 | 43,51834069 | 3,14373971 | 0,19926872 | 15,8996871 | 6,37E-57 | 1,021E-53 | Gm25939 |
| ENSMUSG00000080478 | 11,41314764 | 3,13428231 | 0,3975387 | 8,1298557 | 4,3E-16 | 7,564E-14 | Snord23 |
| ENSMUSG00000064841 | 3,408933412 | 3,08939433 | 1,48224225 | 3,02797185 | 0,002462 | 0,0372272 | Gm26205 |

| | | | | | | | |
|---|---|---|---|---|---|---|---|
| ENSMUSG00000094306 | 320,6151439 | 3,0869468 | 0,67944427 | 4,96618323 | 6,83E-07 | 3,47E-05 | Gm24924 |
| ENSMUSG00000089093 | 3,847474124 | 3,07916638 | 0,74941803 | 4,58350214 | 4,57E-06 | 0,0001864 | Snord11 |
| ENSMUSG00000096349 | 5608,526165 | 3,07880104 | 0,17059189 | 18,1588223 | 1,09E-73 | 3,156E-70 | Gm22513 |
| ENSMUSG00000065327 | 4,231812401 | 3,07626811 | 0,61751448 | 5,37396926 | 7,7E-08 | 4,73E-06 | Gm22058 |
| ENSMUSG00000093660 | 3,927213196 | 3,06189146 | 0,71184624 | 4,75630909 | 1,97E-06 | 9,004E-05 | Gm26505 |
| ENSMUSG00000088351 | 3,705327555 | 3,05825891 | 0,88768352 | 4,00725678 | 6,14E-05 | 0,0019146 | Gm24016 |
| ENSMUSG00000091021 | 12,84391564 | 3,03926741 | 0,33756496 | 9,2188603 | 3E-20 | 7,879E-18 | Gm17300 |
| ENSMUSG00000087708 | 8,925759364 | 3,0364227 | 0,48978917 | 6,51115689 | 7,46E-11 | 7,214E-09 | Gm23000 |
| ENSMUSG00000064967 | 3,07438834 | 2,99140816 | 0,89659522 | 3,91248919 | 9,13E-05 | 0,0026794 | Gm24405 |
| ENSMUSG00000094655 | 9,824739753 | 2,96603819 | 0,40028184 | 7,67401891 | 1,67E-14 | 2,506E-12 | Gm25360 |
| ENSMUSG00000077704 | 7,934006932 | 2,96131306 | 0,51224404 | 6,11865962 | 9,44E-10 | 8,011E-08 | Snord89 |
| ENSMUSG00000094668 | 48,22127104 | 2,91972959 | 0,19446937 | 15,1417324 | 8,59E-52 | 1,24E-48 | Gm24871 |
| ENSMUSG00000095915 | 20,3894278 | 2,9165942 | 0,31546944 | 9,45140939 | 3,34E-21 | 9,844E-19 | n-R5s115 |
| ENSMUSG00000106585 | 15,6136049 | 2,90185425 | 0,32429968 | 9,16589052 | 4,91E-20 | 1,223E-17 | SNORA17 |
| ENSMUSG00000065006 | 4,207117856 | 2,87825753 | 0,88899763 | 3,84149398 | 0,000122 | 0,0034068 | Gm26325 |
| ENSMUSG00000065160 | 5,396925525 | 2,87796615 | 0,65553505 | 4,83748594 | 1,31E-06 | 6,221E-05 | Mir5117 |
| ENSMUSG00000089387 | 3,933426246 | 2,87788051 | 0,69177777 | 4,62733906 | 3,7E-06 | 0,0001563 | Gm24679 |
| ENSMUSG00000064626 | 5,603283881 | 2,86646746 | 1,16839793 | 3,24750733 | 0,001164 | 0,0207416 | Gm22306 |
| ENSMUSG00000064493 | 5,083777215 | 2,8550695 | 0,61051115 | 5,0911338 | 3,56E-07 | 1,909E-05 | Snora28 |
| ENSMUSG00000093852 | 223,6999874 | 2,82100593 | 0,31677085 | 9,11547368 | 7,83E-20 | 1,884E-17 | Gm24312 |
| ENSMUSG00000099937 | 3,030968904 | 2,81822488 | 0,77471102 | 4,16817677 | 3,07E-05 | 0,0010329 | Gm28872 |
| ENSMUSG00000087935 | 163,0081786 | 2,80218728 | 0,1735581 | 16,2739146 | 1,51E-59 | 2,727E-56 | Snora81 |
| ENSMUSG00000096391 | 13,38102652 | 2,800895 | 0,37950101 | 7,63908619 | 2,19E-14 | 3,157E-12 | Gm23686 |
| ENSMUSG00000064387 | 17,68345081 | 2,79941555 | 0,2868389 | 9,95554513 | 2,39E-23 | 9,058E-21 | Snora73a |
| ENSMUSG00000077575 | 13,59695674 | 2,79359182 | 0,36705168 | 7,86462233 | 3,7E-15 | 6,003E-13 | Gm24888 |
| ENSMUSG00000061780 | 10,27620268 | 2,79011878 | 0,85553191 | 3,86604861 | 0,000111 | 0,0031177 | Cfd |
| ENSMUSG00000094830 | 7,372834482 | 2,77061406 | 0,60795962 | 4,97496279 | 6,53E-07 | 3,351E-05 | n-R5s194 |
| ENSMUSG00000065018 | 41,53330332 | 2,76298373 | 0,25357956 | 11,0722929 | 1,71E-28 | 9,869E-26 | Gm23511 |
| ENSMUSG00000065794 | 30,64566922 | 2,75734121 | 0,2644866 | 10,6099966 | 2,68E-26 | 1,288E-23 | Gm25514 |
| ENSMUSG00000065899 | 23,36055205 | 2,75688562 | 0,27681909 | 10,1516509 | 3,26E-24 | 1,306E-21 | Gm24523 |
| ENSMUSG00000064853 | 10,33667829 | 2,74140005 | 0,39517947 | 7,21969402 | 5,21E-13 | 6,266E-11 | Gm23442 |
| ENSMUSG00000065232 | 6,686279019 | 2,71234476 | 0,59668516 | 4,9779568 | 6,43E-07 | 3,312E-05 | Gm22973 |
| ENSMUSG00000065089 | 4,222177772 | 2,71076844 | 0,68414391 | 4,44798858 | 8,67E-06 | 0,0003257 | Gm26493 |
| ENSMUSG00000088595 | 21,93640221 | 2,69016095 | 0,28845353 | 9,53464697 | 1,5E-21 | 4,823E-19 | Gm24601 |
| ENSMUSG00000092805 | 459,0491398 | 2,68374017 | 0,16315789 | 16,5556252 | 1,46E-61 | 3,006E-58 | Gm26461 |
| ENSMUSG00000080463 | 27,49171493 | 2,68174533 | 0,48312645 | 5,88633328 | 3,95E-09 | 3,064E-07 | Gm22489 |
| ENSMUSG00000064558 | 4,425181996 | 2,67230767 | 0,65802091 | 4,5368495 | 5,71E-06 | 0,0002302 | Gm24620 |
| ENSMUSG00000075833 | 7,367051468 | 2,66440927 | 0,52733281 | 5,43207907 | 5,57E-08 | 3,486E-06 | Gm22003 |
| ENSMUSG00000064925 | 34,95436303 | 2,64321919 | 0,25504368 | 10,5491427 | 5,13E-26 | 2,386E-23 | Snora62 |
| ENSMUSG00000065734 | 9,376377712 | 2,6351462 | 0,45154116 | 6,16774116 | 6,93E-10 | 6,022E-08 | Snord49a |
| ENSMUSG00000093413 | 4,939002413 | 2,60675831 | 0,59291287 | 4,84000056 | 1,3E-06 | 6,184E-05 | Snora15 |
| ENSMUSG00000065041 | 31,59933999 | 2,6046151 | 0,22802094 | 11,5919979 | 4,52E-31 | 3,437E-28 | Gm24044 |
| ENSMUSG00000064987 | 2,149371264 | 2,60444932 | 1,15840805 | 3,08595487 | 0,002029 | 0,0323899 | Gm25260 |
| ENSMUSG00000065124 | 6,368891383 | 2,58524457 | 1,03854308 | 3,26599605 | 0,001091 | 0,0198754 | Snora65 |
| ENSMUSG00000070178 | 2,471954991 | 2,57411175 | 0,99120996 | 3,328434 | 0,000873 | 0,0170087 | n-R5s71 |
| ENSMUSG00000096037 | 6,128992231 | 2,56622477 | 0,50183681 | 5,49369002 | 3,94E-08 | 2,536E-06 | n-R5s136 |
| ENSMUSG00000077563 | 21,82030259 | 2,56306511 | 0,27990098 | 9,36930644 | 7,3E-21 | 2,066E-18 | Snora68 |
| ENSMUSG00000088168 | 3,030621924 | 2,5533221 | 0,9691611 | 3,37885055 | 0,000728 | 0,0147119 | Gm23540 |
| ENSMUSG00000080465 | 62,49188432 | 2,551166 | 0,2325566 | 11,1432933 | 7,72E-29 | 4,643E-26 | Gm22486 |
| ENSMUSG00000000486 | 2,401866565 | 2,53923903 | 0,89722052 | 3,5174049 | 0,000436 | 0,0097059 | Sept1 |
| ENSMUSG00000064453 | 105,7179131 | 2,53797156 | 0,13990108 | 18,2441935 | 2,3E-74 | 8,302E-71 | Snord21 |
| ENSMUSG00000064994 | 19,96007836 | 2,51869224 | 0,30099208 | 8,59718121 | 8,17E-18 | 1,709E-15 | Gm22422 |
| ENSMUSG00000064798 | 84,53151868 | 2,50961924 | 0,34200311 | 7,60014849 | 2,96E-14 | 4,226E-12 | n-R5s193 |
| ENSMUSG00000096147 | 22,31712811 | 2,50434563 | 0,6195883 | 4,51379233 | 6,37E-06 | 0,0002546 | Gm25818 |
| ENSMUSG00000030276 | 4,314077674 | 2,50324716 | 0,59616675 | 4,65774257 | 3,2E-06 | 0,0001373 | Ttll3 |
| ENSMUSG00000077323 | 52,73866949 | 2,50078426 | 0,19672291 | 12,8622071 | 7,34E-38 | 7,066E-35 | Rnu11 |
| ENSMUSG00000065862 | 5,691323301 | 2,49890272 | 0,66637839 | 4,2553559 | 2,09E-05 | 0,0007311 | Gm24029 |

| | | | | | | | |
|---|---|---|---|---|---|---|---|
| ENSMUSG00000065591 | 6,785837062 | 2,49582282 | 0,56875979 | 4,82845159 | 1,38E-06 | 6,468E-05 | Gm22753 |
| ENSMUSG00000084480 | 4,457895003 | 2,49216282 | 0,59120994 | 4,66935789 | 3,02E-06 | 0,0001305 | n-R5s155 |
| ENSMUSG00000064451 | 21,65225676 | 2,49007688 | 0,30763056 | 8,32996093 | 8,09E-17 | 1,536E-14 | Snora23 |
| ENSMUSG00000065905 | 12,78207471 | 2,48070634 | 0,41099107 | 6,35312094 | 2,11E-10 | 1,927E-08 | Gm26110 |
| ENSMUSG00000064721 | 9,364890437 | 2,48057368 | 0,41542375 | 6,29033284 | 3,17E-10 | 2,857E-08 | Gm25855 |
| ENSMUSG00000094665 | 2,988224316 | 2,47205209 | 0,85555064 | 3,56162179 | 0,000369 | 0,008483 | Gm24873 |
| ENSMUSG00000064857 | 2,926727204 | 2,4717465 | 0,81080783 | 3,6943577 | 0,00022 | 0,0055326 | n-R5s25 |
| ENSMUSG00000064500 | 5,151920242 | 2,46484246 | 0,68535234 | 4,13068295 | 3,62E-05 | 0,0011917 | Gm25296 |
| ENSMUSG00000064455 | 11,58953306 | 2,46364051 | 0,34738922 | 7,38011418 | 1,58E-13 | 2,056E-11 | n-R5s56 |
| ENSMUSG00000106943 | 18,80795598 | 2,45942621 | 0,30071784 | 8,39868312 | 4,52E-17 | 9,05E-15 | Snora26 |
| ENSMUSG00000065686 | 2,818140135 | 2,44458391 | 0,98154558 | 3,2662944 | 0,00109 | 0,0198754 | Snora5c |
| ENSMUSG00000077734 | 17,66551817 | 2,41440319 | 0,53484408 | 4,93455452 | 8,03E-07 | 3,998E-05 | Snord83b |
| ENSMUSG00000044757 | 4,271766854 | 2,37961964 | 0,59437573 | 4,47335353 | 7,7E-06 | 0,0002995 | Gm6430 |
| ENSMUSG00000064382 | 16,73454241 | 2,37279221 | 0,33031889 | 7,45740888 | 8,82E-14 | 1,201E-11 | Gm26447 |
| ENSMUSG00000029368 | 12,61091204 | 2,3718935 | 0,89189101 | 3,41346811 | 0,000641 | 0,0133955 | Alb |
| ENSMUSG00000094377 | 454,9040834 | 2,37004616 | 0,49376819 | 5,19628839 | 2,03E-07 | 1,137E-05 | Gm24407 |
| ENSMUSG00000088529 | 40,27254091 | 2,3661284 | 0,29142481 | 8,35228587 | 6,7E-17 | 1,306E-14 | Gm26083 |
| ENSMUSG00000064280 | 5,613819757 | 2,36177778 | 0,55237715 | 4,7292177 | 2,25E-06 | 0,0001016 | Ccdc146 |
| ENSMUSG00000075770 | 2,36543625 | 2,33445207 | 0,9707702 | 3,18928794 | 0,001426 | 0,0244152 | Gm22589 |
| ENSMUSG00000096613 | 3,338047779 | 2,33423401 | 0,79019315 | 3,60632186 | 0,000311 | 0,0073352 | n-R5s102 |
| ENSMUSG00000064702 | 436,9746472 | 2,32936159 | 0,2828533 | 8,46310763 | 2,6E-17 | 5,367E-15 | Gm24950 |
| ENSMUSG00000092687 | 3,363375564 | 2,32128523 | 1,01592151 | 3,13354448 | 0,001727 | 0,0285821 | Gm25744 |
| ENSMUSG00000064634 | 2,430513647 | 2,31692336 | 0,85759768 | 3,42696325 | 0,00061 | 0,0129343 | Gm22620 |
| ENSMUSG00000024301 | 2,785998826 | 2,31634715 | 0,80380336 | 3,55326167 | 0,00038 | 0,0087294 | Kifc5b |
| ENSMUSG00000064655 | 19,62564621 | 2,31140601 | 0,29585361 | 8,05713596 | 7,81E-16 | 1,358E-13 | Gm25788 |
| ENSMUSG00000041642 | 10,01857433 | 2,30111846 | 0,4132258 | 5,9099008 | 3,42E-09 | 2,685E-07 | Kif21b |
| ENSMUSG00000071068 | 3,677536195 | 2,29594297 | 0,76042147 | 3,66348232 | 0,000249 | 0,006096 | Treml2 |
| ENSMUSG00000065259 | 10,10354532 | 2,29537712 | 0,38995108 | 6,2120835 | 5,23E-10 | 4,629E-08 | Snora30 |
| ENSMUSG00000065893 | 2,217589294 | 2,28518024 | 0,84261347 | 3,4174546 | 0,000632 | 0,0132392 | n-R5s151 |
| ENSMUSG00000065289 | 27,44229996 | 2,28261091 | 0,34172539 | 6,96507295 | 3,28E-12 | 3,644E-10 | Gm23650 |
| ENSMUSG00000098343 | 27,15683121 | 2,27625433 | 0,48502174 | 5,10035109 | 3,39E-07 | 1,832E-05 | Mir6240 |
| ENSMUSG00000064901 | 23,13601044 | 2,27562819 | 0,27012385 | 8,65012032 | 5,14E-18 | 1,125E-15 | Snora21 |
| ENSMUSG00000095449 | 8,833449917 | 2,27023145 | 0,43111571 | 5,6280079 | 1,82E-08 | 1,241E-06 | n-R5s112 |
| ENSMUSG00000075918 | 23,24090728 | 2,26149305 | 0,69934316 | 3,82386667 | 0,000131 | 0,0036155 | n-R5s2 |
| ENSMUSG00000065097 | 24,73127733 | 2,25892396 | 1,19846254 | 2,95672867 | 0,003109 | 0,0446015 | Snora16a |
| ENSMUSG00000105131 | 8,144433502 | 2,25841161 | 0,94104083 | 3,20216885 | 0,001364 | 0,0235167 | AC069308.1 |
| ENSMUSG00000065878 | 8,31408116 | 2,25598393 | 0,45165873 | 5,37596274 | 7,62E-08 | 4,698E-06 | Snord34 |
| ENSMUSG00000086358 | 3,069782771 | 2,24526179 | 0,80135384 | 3,49619806 | 0,000472 | 0,0104617 | Gm13270 |
| ENSMUSG00000065752 | 4,09893881 | 2,23697545 | 0,72950741 | 3,68658709 | 0,000227 | 0,005676 | Gm23344 |
| ENSMUSG00000080518 | 2,370798277 | 2,23575058 | 0,97663203 | 3,1359289 | 0,001713 | 0,0283833 | Gm22193 |
| ENSMUSG00000102189 | 19,20248899 | 2,21964225 | 0,31703911 | 7,27497824 | 3,46E-13 | 4,348E-11 | Gm37194 |
| ENSMUSG00000064512 | 22,00693642 | 2,21576402 | 0,31777026 | 7,24225098 | 4,41E-13 | 5,352E-11 | Gm22456 |
| ENSMUSG00000065208 | 16,81496102 | 2,21549561 | 0,28535027 | 8,01289277 | 1,12E-15 | 1,925E-13 | Gm24616 |
| ENSMUSG00000023034 | 38,31313267 | 2,21480522 | 0,23674447 | 9,57412221 | 1,03E-21 | 3,369E-19 | Nr4a1 |
| ENSMUSG00000038578 | 2,027156097 | 2,21394973 | 1,03944336 | 3,04364276 | 0,002337 | 0,0359533 | Susd1 |
| ENSMUSG00000065637 | 27,23273663 | 2,21030763 | 0,2375403 | 9,50870302 | 1,93E-21 | 5,928E-19 | Gm26397 |
| ENSMUSG00000064745 | 2,463180288 | 2,21010979 | 0,92222035 | 3,20439999 | 0,001353 | 0,0233911 | Gm24708 |
| ENSMUSG00000096887 | 301,2283616 | 2,20840548 | 0,1789037 | 12,496697 | 7,78E-36 | 7,019E-33 | Gm20594 |
| ENSMUSG00000064350 | 129,7963446 | 2,20343888 | 0,64700688 | 3,96562961 | 7,32E-05 | 0,0022168 | mt-Ty |
| ENSMUSG00000065118 | 10,55808504 | 2,18854315 | 0,40666594 | 5,7344869 | 9,78E-09 | 7,129E-07 | Gm23297 |
| ENSMUSG00000080542 | 10,26796215 | 2,1826843 | 0,48122664 | 4,9490493 | 7,46E-07 | 3,737E-05 | Gm22710 |
| ENSMUSG00000106474 | 5,729241121 | 2,17507043 | 0,74353515 | 3,59588542 | 0,000323 | 0,0075614 | AC124613.1 |
| ENSMUSG00000095299 | 3,349516812 | 2,16262632 | 0,84252882 | 3,30094804 | 0,000964 | 0,0182248 | Gm25380 |
| ENSMUSG00000077192 | 5,095159473 | 2,15360079 | 0,62347279 | 4,01088785 | 6,05E-05 | 0,0018993 | Snora17 |
| ENSMUSG00000090207 | 2,289382733 | 2,15210941 | 0,78014121 | 3,45142122 | 0,000558 | 0,0120289 | 4930524O07Rik |
| ENSMUSG00000070063 | 7,317639795 | 2,15107547 | 0,49353196 | 4,7927216 | 1,65E-06 | 7,562E-05 | Snora33 |
| ENSMUSG00000095676 | 64,96407058 | 2,14982205 | 0,37621171 | 6,04318876 | 1,51E-09 | 1,225E-07 | Gm25099 |

| | | | | | | | |
|---|---|---|---|---|---|---|---|
| ENSMUSG00000080610 | 15,82873069 | 2,14974514 | 0,40939773 | 5,61411046 | 1,98E-08 | 1,332E-06 | Snord110 |
| ENSMUSG00000064513 | 4,489561192 | 2,1472867 | 0,80228938 | 3,3994103 | 0,000675 | 0,0139071 | Gm22457 |
| ENSMUSG00000064403 | 14,66454684 | 2,14175505 | 0,34819853 | 6,45745899 | 1,06E-10 | 1,004E-08 | Gm23928 |
| ENSMUSG00000065295 | 8,832152187 | 2,13853989 | 0,4998611 | 4,72402274 | 2,31E-06 | 0,0001039 | Gm24134 |
| ENSMUSG00000092774 | 3,147633239 | 2,12747595 | 0,71738043 | 3,62121554 | 0,000293 | 0,0069711 | Gm26197 |
| ENSMUSG00000064725 | 3,517154575 | 2,11378657 | 0,68511667 | 3,70302183 | 0,000213 | 0,0054128 | Snord96a |
| ENSMUSG00000065939 | 3,150183019 | 2,10800277 | 0,78114737 | 3,40696923 | 0,000657 | 0,0136396 | Snora2b |
| ENSMUSG00000064748 | 14,09198339 | 2,10432814 | 0,35791583 | 6,2045576 | 5,49E-10 | 4,827E-08 | n-R5s28 |
| ENSMUSG00000065226 | 2,585256949 | 2,10206409 | 0,97174831 | 3,04484683 | 0,002328 | 0,035854 | Gm25791 |
| ENSMUSG00000080486 | 15,3424984 | 2,10074956 | 0,42688382 | 5,30411066 | 1,13E-07 | 6,642E-06 | Snord100 |
| ENSMUSG00000095590 | 19,09137139 | 2,08944002 | 0,2773389 | 7,78327204 | 7,07E-15 | 1,109E-12 | Gm24305 |
| ENSMUSG00000065642 | 5,96161128 | 2,08112882 | 0,50745731 | 4,56551594 | 4,98E-06 | 0,0002025 | Snora69 |
| ENSMUSG00000016255 | 5,729786908 | 2,07974001 | 0,63054076 | 3,88932604 | 0,000101 | 0,0028957 | Tubb1 |
| ENSMUSG00000065258 | 2,64621214 | 2,07972998 | 1,01078625 | 3,01238886 | 0,002592 | 0,0387217 | Gm23969 |
| ENSMUSG00000065304 | 7,173473588 | 2,0685981 | 0,49151376 | 4,6603679 | 3,16E-06 | 0,000136 | Gm23245 |
| ENSMUSG00000095118 | 34,50880434 | 2,04681043 | 0,28646483 | 7,40635579 | 1,3E-13 | 1,703E-11 | Snord14d |
| ENSMUSG00000077254 | 12,32525102 | 2,03156909 | 0,37823697 | 5,70989953 | 1,13E-08 | 8,157E-07 | Gm26079 |
| ENSMUSG00000029715 | 3,642698801 | 2,01847974 | 0,63875479 | 3,76406846 | 0,000167 | 0,004451 | Pop7 |
| ENSMUSG00000020072 | 3,489528335 | 2,01438087 | 0,68485071 | 3,59210907 | 0,000328 | 0,0076219 | Pbld2 |
| ENSMUSG00000043673 | 3,328506059 | 2,01270263 | 0,77816259 | 3,32896252 | 0,000872 | 0,0169994 | Kcns3 |
| ENSMUSG00000088185 | 94,86790498 | 2,01206466 | 0,17954233 | 11,3689265 | 5,97E-30 | 3,917E-27 | Scarna2 |
| ENSMUSG00000070883 | 3,936481227 | 2,00692626 | 0,69295573 | 3,57229699 | 0,000354 | 0,0081837 | Ccdc173 |
| ENSMUSG00000092674 | 9,659737136 | 2,00436104 | 0,43723063 | 4,99162791 | 5,99E-07 | 3,108E-05 | Gm24105 |
| ENSMUSG00000077222 | 19,3413445 | 2,00244585 | 0,2898765 | 7,16966065 | 7,52E-13 | 8,821E-11 | Gm22270 |
| ENSMUSG00000064714 | 2,489874184 | 1,98415711 | 0,82711176 | 3,20391838 | 0,001356 | 0,0234022 | n-R5s62 |
| ENSMUSG00000092730 | 32,1211032 | 1,98129067 | 0,26663203 | 7,68032816 | 1,59E-14 | 2,41E-12 | Snora24 |
| ENSMUSG00000065649 | 20,46479172 | 1,97871072 | 0,88086243 | 3,09142095 | 0,001992 | 0,0320476 | Snora74a |
| ENSMUSG00000037318 | 2,932546948 | 1,97622223 | 0,79646652 | 3,27049143 | 0,001074 | 0,0196615 | Traf3ip3 |
| ENSMUSG00000037922 | 3,924341801 | 1,96904572 | 0,66212878 | 3,61726994 | 0,000298 | 0,0070666 | Bank1 |
| ENSMUSG00000099021 | 119334,8796 | 1,95844721 | 0,38183149 | 5,48797978 | 4,07E-08 | 2,596E-06 | Rn7s1 |
| ENSMUSG00000093112 | 22,07708965 | 1,9569292 | 0,31301858 | 6,54993677 | 5,76E-11 | 5,729E-09 | Gm22701 |
| ENSMUSG00000077239 | 11,76065557 | 1,95146822 | 0,4533239 | 4,74232065 | 2,11E-06 | 9,558E-05 | Snord66 |
| ENSMUSG00000033730 | 2,374722874 | 1,94365806 | 0,90598745 | 3,02845791 | 0,002458 | 0,0372216 | Egr3 |
| ENSMUSG00000066554 | 2,168409105 | 1,93857162 | 0,95799603 | 2,98563581 | 0,00283 | 0,0414601 | Gm10167 |
| ENSMUSG00000091825 | 8,375245978 | 1,93694478 | 0,44891856 | 4,75502434 | 1,98E-06 | 9,033E-05 | Gm5778 |
| ENSMUSG00000092680 | 40,73762269 | 1,93272389 | 0,2793502 | 7,18772747 | 6,59E-13 | 7,857E-11 | Snord55 |
| ENSMUSG00000065061 | 48,96277463 | 1,92726888 | 0,1780415 | 10,9924899 | 4,15E-28 | 2,305E-25 | Gm22884 |
| ENSMUSG00000065272 | 23,23785163 | 1,92032787 | 0,33458244 | 6,06667111 | 1,31E-09 | 1,065E-07 | Snord57 |
| ENSMUSG00000064647 | 9,536556805 | 1,89518396 | 0,42552439 | 4,87383942 | 1,09E-06 | 5,247E-05 | Gm23301 |
| ENSMUSG00000064563 | 23,43147612 | 1,88758294 | 0,27546679 | 7,11532537 | 1,12E-12 | 1,289E-10 | Snora7a |
| ENSMUSG00000089161 | 5,814263714 | 1,87162725 | 0,54264382 | 3,99533791 | 6,46E-05 | 0,0020049 | Gm24130 |
| ENSMUSG00000065911 | 557,5757125 | 1,86862168 | 0,24595214 | 7,85401772 | 4,03E-15 | 6,461E-13 | Gm24447 |
| ENSMUSG00000077709 | 31,79778462 | 1,86719075 | 0,26002527 | 7,43718532 | 1,03E-13 | 1,374E-11 | Snora64 |
| ENSMUSG00000064966 | 253,214058 | 1,86312084 | 0,14439627 | 13,0526029 | 6,14E-39 | 6,817E-36 | Snord15b |
| ENSMUSG00000049892 | 6,385627477 | 1,86306611 | 0,50822193 | 4,17918768 | 2,93E-05 | 0,000991 | Rasd1 |
| ENSMUSG00000065876 | 12,6076353 | 1,86068829 | 0,43648829 | 4,70245245 | 2,57E-06 | 0,0001134 | Gm25682 |
| ENSMUSG00000088158 | 2,338203204 | 1,85206879 | 0,9182526 | 2,95054269 | 0,003172 | 0,0453242 | Scarna3b |
| ENSMUSG00000053656 | 43,30292181 | 1,84981218 | 0,19008332 | 9,89372293 | 4,43E-23 | 1,599E-20 | Dancr |
| ENSMUSG00000096211 | 10,36954759 | 1,83973987 | 0,38288431 | 5,18962796 | 2,11E-07 | 1,17E-05 | Gm23330 |
| ENSMUSG00000065729 | 8,208656833 | 1,8350461 | 0,59950295 | 3,6815517 | 0,000232 | 0,0057679 | Gm26176 |
| ENSMUSG00000040345 | 3,843867714 | 1,82878915 | 0,66561511 | 3,42548069 | 0,000614 | 0,012986 | Arhgap9 |
| ENSMUSG00000064768 | 7,273968164 | 1,81791095 | 0,54702706 | 3,88870401 | 0,000101 | 0,0028972 | Snord60 |
| ENSMUSG00000065102 | 2,981666418 | 1,81743935 | 0,7405695 | 3,22887255 | 0,001243 | 0,0218415 | n-R5s50 |
| ENSMUSG00000065087 | 316,2631235 | 1,80950764 | 0,09354374 | 19,4376294 | 3,71E-84 | 1,784E-80 | Snord22 |
| ENSMUSG00000028864 | 5,972236427 | 1,80724306 | 0,54855182 | 3,87195047 | 0,000108 | 0,0030551 | Hgf |
| ENSMUSG00000092746 | 662,8368482 | 1,80683109 | 0,35678323 | 5,43005833 | 5,63E-08 | 3,504E-06 | Rn7s6 |
| ENSMUSG00000083579 | 2,856397205 | 1,80541191 | 0,78510405 | 3,11373513 | 0,001847 | 0,0302945 | Gm15538 |

| | | | | | | | |
|---|---|---|---|---|---|---|---|
| ENSMUSG00000077565 | 31,54936904 | 1,80313974 | 0,26886358 | 6,98198122 | 2,91E-12 | 3,256E-10 | Gm23201 |
| ENSMUSG00000038894 | 46,24735837 | 1,79458985 | 0,21803233 | 8,45562068 | 2,78E-17 | 5,642E-15 | Irs2 |
| ENSMUSG00000087968 | 27,9323877 | 1,78537087 | 0,25476403 | 7,26881316 | 3,63E-13 | 4,473E-11 | Gm25395 |
| ENSMUSG00000096263 | 35,29474267 | 1,78464165 | 0,44587426 | 4,46170289 | 8,13E-06 | 0,0003104 | Snord14c |
| ENSMUSG00000087819 | 11,86584853 | 1,77564733 | 0,4210854 | 4,65175646 | 3,29E-06 | 0,0001405 | Gm25117 |
| ENSMUSG00000065824 | 15,59831079 | 1,76720156 | 0,30326037 | 6,14373329 | 8,06E-10 | 6,924E-08 | Gm26315 |
| ENSMUSG00000065922 | 29,08561728 | 1,76274592 | 0,2353203 | 7,72980148 | 1,08E-14 | 1,671E-12 | n-R5-8s1 |
| ENSMUSG00000064999 | 8,415697505 | 1,76052615 | 0,44226659 | 4,45384949 | 8,43E-06 | 0,0003195 | Gm26035 |
| ENSMUSG00000020928 | 5,037134763 | 1,75065377 | 0,60521083 | 3,54246024 | 0,000396 | 0,0090516 | Higd1b |
| ENSMUSG00000056753 | 3,052100954 | 1,7493253 | 0,68404958 | 3,28589249 | 0,001017 | 0,018881 | C330011M18Rik |
| ENSMUSG00000088088 | 382,2648965 | 1,74434324 | 0,24059998 | 7,49576349 | 6,59E-14 | 9,146E-12 | Rmrp |
| ENSMUSG00000023341 | 5,473287161 | 1,74376295 | 0,56533618 | 3,70135307 | 0,000214 | 0,0054371 | Mx2 |
| ENSMUSG00000038491 | 3,440190414 | 1,7409372 | 0,62508339 | 3,44386253 | 0,000573 | 0,0122785 | Cdk19os |
| ENSMUSG00000064344 | 1065,613708 | 1,74092642 | 0,51318364 | 3,9336874 | 8,37E-05 | 0,0024891 | mt-Tm |
| ENSMUSG00000065822 | 64,89446235 | 1,72753306 | 0,18657759 | 9,45442086 | 3,25E-21 | 9,765E-19 | Snord15a |
| ENSMUSG00000096243 | 557,7681099 | 1,72140283 | 0,48094892 | 4,09227665 | 4,27E-05 | 0,0013915 | Gm24265 |
| ENSMUSG00000094245 | 3,297370413 | 1,71882463 | 0,85494049 | 2,93697996 | 0,003314 | 0,0466615 | Gm23966 |
| ENSMUSG00000030263 | 4,347787751 | 1,71801985 | 0,64836712 | 3,36961743 | 0,000753 | 0,0150657 | Lrmp |
| ENSMUSG00000077714 | 87,32491953 | 1,7153161 | 0,14961347 | 11,6270026 | 3E-31 | 2,409E-28 | Snord17 |
| ENSMUSG00000097971 | 30295,17545 | 1,68816914 | 0,20086845 | 8,664937 | 4,52E-18 | 1,003E-15 | Gm26917 |
| ENSMUSG00000006574 | 3,517131249 | 1,68514217 | 0,81781477 | 3,00905245 | 0,002621 | 0,0390283 | Slc4a1 |
| ENSMUSG00000092702 | 3,230004619 | 1,67843273 | 0,80023934 | 2,98332401 | 0,002851 | 0,0416477 | Gm24514 |
| ENSMUSG00000064941 | 15,02204267 | 1,67591871 | 0,38265569 | 4,79424057 | 1,63E-06 | 7,529E-05 | Gm23238 |
| ENSMUSG00000097769 | 101,591868 | 1,64652069 | 0,18410216 | 9,14215245 | 6,12E-20 | 1,497E-17 | Snhg4 |
| ENSMUSG00000065778 | 6,857692203 | 1,64611387 | 0,68422491 | 3,18086392 | 0,001468 | 0,0250177 | Gm22154 |
| ENSMUSG00000064349 | 52,00563968 | 1,6415737 | 0,66713187 | 3,20537987 | 0,001349 | 0,0233675 | mt-Tc |
| ENSMUSG00000077611 | 29,42107908 | 1,63591024 | 0,25518711 | 6,69200728 | 2,2E-11 | 2,253E-09 | Gm23946 |
| ENSMUSG00000034173 | 18,95137885 | 1,63302284 | 0,2823576 | 6,1122593 | 9,82E-10 | 8,29E-08 | Zbed5 |
| ENSMUSG00000065845 | 22,25192062 | 1,63075124 | 0,63072019 | 3,28629624 | 0,001015 | 0,018881 | Gm25189 |
| ENSMUSG00000030838 | 18,59936215 | 1,62150632 | 0,38963382 | 4,603622 | 4,15E-06 | 0,0001712 | Ush1c |
| ENSMUSG00000020169 | 12,04743858 | 1,62112664 | 0,35279864 | 4,99395119 | 5,92E-07 | 3,082E-05 | Best3 |
| ENSMUSG00000084700 | 4,41659536 | 1,62045399 | 0,72260674 | 3,0721178 | 0,002125 | 0,0334487 | Gm22984 |
| ENSMUSG00000064372 | 2887,578015 | 1,6202219 | 0,52959147 | 3,65793049 | 0,000254 | 0,0062085 | mt-Tp |
| ENSMUSG00000064751 | 13,24867882 | 1,62017967 | 0,36065465 | 4,90017617 | 9,58E-07 | 4,668E-05 | Gm26330 |
| ENSMUSG00000049230 | 4,604213877 | 1,61829076 | 0,59716821 | 3,3990007 | 0,000676 | 0,0139071 | Gm9833 |
| ENSMUSG00000065176 | 142,5141792 | 1,61050222 | 0,1673994 | 9,80675564 | 1,05E-22 | 3,706E-20 | Rnu12 |
| ENSMUSG00000037221 | 7,209914035 | 1,60538079 | 0,45133843 | 4,07962199 | 4,51E-05 | 0,0014628 | Mospd3 |
| ENSMUSG00000029279 | 3,145152635 | 1,60201747 | 0,77103106 | 2,97348695 | 0,002944 | 0,0426608 | Brdt |
| ENSMUSG00000088789 | 40,59452935 | 1,59855654 | 0,38185543 | 4,61816447 | 3,87E-06 | 0,0001619 | Scarna13 |
| ENSMUSG00000073940 | 169,5864615 | 1,59824421 | 0,26052912 | 6,41978789 | 1,36E-10 | 1,271E-08 | Hbb-bt |
| ENSMUSG00000028341 | 20,37763617 | 1,59439962 | 0,34698928 | 4,99396496 | 5,92E-07 | 3,082E-05 | Nr4a3 |
| ENSMUSG00000065392 | 7,66748375 | 1,59133025 | 0,49197076 | 3,80887297 | 0,00014 | 0,0038083 | Gm25894 |
| ENSMUSG00000089901 | 16,87021311 | 1,59046903 | 0,30930304 | 5,49641142 | 3,88E-08 | 2,52E-06 | Gm8113 |
| ENSMUSG00000064823 | 9,928041742 | 1,58863895 | 0,51287003 | 3,68898937 | 0,000225 | 0,0056408 | Snord82 |
| ENSMUSG00000012819 | 19,12378364 | 1,58817594 | 0,29856584 | 5,66185175 | 1,5E-08 | 1,054E-06 | Cdh23 |
| ENSMUSG00000064844 | 12,00129496 | 1,58347267 | 0,49712134 | 3,76111487 | 0,000169 | 0,0044873 | Gm26202 |
| ENSMUSG00000064342 | 460,3851252 | 1,57800692 | 0,50811832 | 3,68606807 | 0,000228 | 0,0056763 | mt-Ti |
| ENSMUSG00000080365 | 2,79040573 | 1,57621421 | 0,74481808 | 3,00566509 | 0,00265 | 0,0392631 | Gm25776 |
| ENSMUSG00000083076 | 3,904471275 | 1,57521284 | 0,6353502 | 3,23110864 | 0,001233 | 0,0217012 | Gm11408 |
| ENSMUSG00000030889 | 35,26293915 | 1,57232233 | 0,22170086 | 7,35201485 | 1,95E-13 | 2,493E-11 | Vwa3a |
| ENSMUSG00000094976 | 6,38774661 | 1,56843062 | 0,66204774 | 3,14899638 | 0,001638 | 0,0272695 | Gm25257 |
| ENSMUSG00000065738 | 50,38805975 | 1,56017501 | 0,22253608 | 7,26403031 | 3,76E-13 | 4,595E-11 | Gm24494 |
| ENSMUSG00000065050 | 13,09861974 | 1,54712629 | 0,75401644 | 2,9378027 | 0,003305 | 0,0466274 | Gm25704 |
| ENSMUSG00000065196 | 20,21421003 | 1,54706876 | 0,38501491 | 4,47080931 | 7,79E-06 | 0,0003017 | Snord85 |
| ENSMUSG00000093044 | 5,6147063 | 1,54396661 | 0,57529089 | 3,37676944 | 0,000733 | 0,0147822 | Snord8 |
| ENSMUSG00000052305 | 284,260898 | 1,542816 | 0,11846438 | 13,1627884 | 1,44E-39 | 1,728E-36 | Hbb-bs |
| ENSMUSG00000035697 | 23,24080599 | 1,53768131 | 0,28211431 | 5,78781249 | 7,13E-09 | 5,25E-07 | Hmha1 |

| | | | | | | | |
|---|---|---|---|---|---|---|---|
| ENSMUSG00000065628 | 55,73680266 | 1,53664287 | 0,2566062 | 6,28597862 | 3,26E-10 | 2,92E-08 | Snord33 |
| ENSMUSG00000021250 | 32,82810577 | 1,51651911 | 0,24167753 | 6,56038852 | 5,37E-11 | 5,378E-09 | Fos |
| ENSMUSG00000035692 | 3,391294518 | 1,50350267 | 0,75897455 | 2,92805082 | 0,003411 | 0,0475129 | Isg15 |
| ENSMUSG00000084708 | 215,0483683 | 1,49674882 | 0,16469058 | 9,28297905 | 1,65E-20 | 4,574E-18 | Gm22988 |
| ENSMUSG00000065336 | 13,51270618 | 1,49525106 | 0,35672575 | 4,62408251 | 3,76E-06 | 0,0001583 | Snora34 |
| ENSMUSG00000080237 | 6,635496861 | 1,49441135 | 0,4944777 | 3,62901222 | 0,000285 | 0,0068088 | Gm14239 |
| ENSMUSG00000064829 | 5,222729443 | 1,49388801 | 0,66058771 | 3,08893468 | 0,002009 | 0,0322093 | Gm22879 |
| ENSMUSG00000047767 | 18,97584198 | 1,49277162 | 0,30626911 | 5,22971512 | 1,7E-07 | 9,684E-06 | Atg16l2 |
| ENSMUSG00000052415 | 10,29605027 | 1,48726247 | 0,41936739 | 4,06129083 | 4,88E-05 | 0,001572 | Tchh |
| ENSMUSG00000093178 | 13,62797525 | 1,48405141 | 0,40321383 | 4,17570719 | 2,97E-05 | 0,001004 | Snord87 |
| ENSMUSG00000004934 | 14,83438305 | 1,47363561 | 0,37280987 | 4,39887555 | 1,09E-05 | 0,0004047 | Pias4 |
| ENSMUSG00000042489 | 3,371662371 | 1,47164277 | 0,64056155 | 3,10071801 | 0,001931 | 0,0313378 | Clspn |
| ENSMUSG00000062031 | 6,376921549 | 1,46008651 | 0,53385266 | 3,4208838 | 0,000624 | 0,0131305 | Athl1 |
| ENSMUSG00000059195 | 34,25682387 | 1,45868905 | 0,22812034 | 6,65658723 | 2,8E-11 | 2,828E-09 | Gm12715 |
| ENSMUSG00000088948 | 91,06811477 | 1,45232402 | 0,20439558 | 7,35430231 | 1,92E-13 | 2,473E-11 | Gm23262 |
| ENSMUSG00000064816 | 48,25209774 | 1,43711455 | 0,45018914 | 3,74729265 | 0,000179 | 0,0046732 | Gm22357 |
| ENSMUSG00000030830 | 6,082743 | 1,43396218 | 0,55967707 | 3,28267931 | 0,001028 | 0,0189997 | Itgal |
| ENSMUSG00000064694 | 63,33626646 | 1,43197246 | 0,31057881 | 4,98924287 | 6,06E-07 | 3,135E-05 | Gm24146 |
| ENSMUSG00000092051 | 7,06722228 | 1,43184917 | 0,50601076 | 3,47946638 | 0,000502 | 0,0110355 | Gm17229 |
| ENSMUSG00000080683 | 6,088509937 | 1,4298609 | 0,52393231 | 3,39892432 | 0,000677 | 0,0139071 | Gm23991 |
| ENSMUSG00000092898 | 8,867486138 | 1,42754567 | 0,57730684 | 3,1964424 | 0,001391 | 0,0239572 | Gm23806 |
| ENSMUSG00000009292 | 4,324013114 | 1,42097256 | 0,67653931 | 2,98393183 | 0,002846 | 0,0416072 | Trpm2 |
| ENSMUSG00000037103 | 3,8294806 | 1,40816694 | 0,69323249 | 2,93250038 | 0,003362 | 0,0470644 | Dcaf15 |
| ENSMUSG00000022503 | 9,501847022 | 1,40762589 | 0,43522553 | 3,78599134 | 0,000153 | 0,0041338 | Nubp1 |
| ENSMUSG00000036528 | 31,48222706 | 1,39156799 | 0,23947177 | 6,10583791 | 1,02E-09 | 8,53E-08 | Ppfibp2 |
| ENSMUSG00000080615 | 49,14678177 | 1,37689237 | 0,44201156 | 3,67174041 | 0,000241 | 0,0059326 | Snord99 |
| ENSMUSG00000004127 | 4,931170335 | 1,37558897 | 0,56437632 | 3,17421456 | 0,001502 | 0,0254179 | Trmt10a |
| ENSMUSG00000084799 | 6,072642721 | 1,37381544 | 0,5294649 | 3,29659088 | 0,000979 | 0,0184615 | Ino80dos |
| ENSMUSG00000025815 | 10,56688377 | 1,37251586 | 0,4491976 | 3,64203712 | 0,00027 | 0,0065494 | Dhtkd1 |
| ENSMUSG00000064336 | 193,2162664 | 1,36960803 | 0,58466948 | 3,09346266 | 0,001978 | 0,0318634 | mt-Tf |
| ENSMUSG00000089542 | 80,86930545 | 1,36928756 | 0,16255248 | 8,63220841 | 6,02E-18 | 1,296E-15 | Gm25835 |
| ENSMUSG00000098374 | 22,4987015 | 1,36914371 | 0,60768402 | 3,03500618 | 0,002405 | 0,0367701 | Gm28043 |
| ENSMUSG00000065273 | 7,161471608 | 1,36559382 | 0,57095167 | 3,15460536 | 0,001607 | 0,0269371 | Gm25128 |
| ENSMUSG00000055413 | 5,658315996 | 1,362205 | 0,64857619 | 2,93173938 | 0,003371 | 0,0471342 | H2-Q5 |
| ENSMUSG00000103405 | 133,4635255 | 1,36015307 | 0,12936961 | 10,6797612 | 1,27E-26 | 6,3E-24 | Dst |
| ENSMUSG00000064371 | 271,2855655 | 1,35904125 | 0,48187959 | 3,44045668 | 0,000581 | 0,0124156 | mt-Tt |
| ENSMUSG00000026730 | 6,129852001 | 1,35869131 | 0,49896483 | 3,3904873 | 0,000698 | 0,0142813 | Pter |
| ENSMUSG00000040675 | 9,907699002 | 1,357133 | 0,42384611 | 3,75839916 | 0,000171 | 0,0045197 | Mthfd1l |
| ENSMUSG00000002825 | 4,334116437 | 1,34334607 | 0,67365801 | 2,90681677 | 0,003651 | 0,0497559 | Qtrt1 |
| ENSMUSG00000080538 | 63,83146242 | 1,34274683 | 0,39045317 | 3,94393081 | 8,02E-05 | 0,0023999 | Gm25541 |
| ENSMUSG00000051984 | 23,50933496 | 1,33833518 | 0,29153934 | 4,97316496 | 6,59E-07 | 3,359E-05 | Sec31b |
| ENSMUSG00000028461 | 19,26621428 | 1,33557872 | 0,31070369 | 4,70535658 | 2,53E-06 | 0,0001122 | Ccdc107 |
| ENSMUSG00000067629 | 10,08963135 | 1,32563173 | 0,41926816 | 3,72534847 | 0,000195 | 0,0050533 | Syngap1 |
| ENSMUSG00000065228 | 13,52813435 | 1,32559904 | 0,49894549 | 3,33096786 | 0,000865 | 0,0169002 | Gm25789 |
| ENSMUSG00000084744 | 43,83030721 | 1,32150758 | 0,21269021 | 6,49052314 | 8,55E-11 | 8,121E-09 | Gm25291 |
| ENSMUSG00000064968 | 11,83775737 | 1,31120562 | 0,44975043 | 3,52908462 | 0,000417 | 0,0094027 | Snord47 |
| ENSMUSG00000040852 | 15,55179232 | 1,30984958 | 0,34639132 | 4,24408604 | 2,19E-05 | 0,0007632 | Plekhh2 |
| ENSMUSG00000001750 | 17,4745821 | 1,30383986 | 0,31086458 | 4,63695354 | 3,54E-06 | 0,0001501 | Tcirg1 |
| ENSMUSG00000064871 | 10,46335385 | 1,29354242 | 0,46227671 | 3,42204771 | 0,000622 | 0,0131123 | Snord58b |
| ENSMUSG00000064339 | 1179305,311 | 1,2844881 | 0,18128985 | 7,32469768 | 2,39E-13 | 3,031E-11 | mt-Rnr2 |
| ENSMUSG00000105780 | 12,0516847 | 1,28326897 | 0,46420813 | 3,39801377 | 0,000679 | 0,0139336 | SNORD77 |
| ENSMUSG00000026270 | 7,630660006 | 1,28106477 | 0,49043478 | 3,29102756 | 0,000998 | 0,0187082 | Capn10 |
| ENSMUSG00000004626 | 6,549682959 | 1,26936604 | 0,49680425 | 3,25532577 | 0,001133 | 0,0203548 | Stxbp2 |
| ENSMUSG00000034858 | 131,3496456 | 1,26070642 | 0,1288136 | 9,93223082 | 3,01E-23 | 1,115E-20 | Fam214a |
| ENSMUSG00000065145 | 192,9465894 | 1,25391602 | 0,25621599 | 5,25862732 | 1,45E-07 | 8,411E-06 | Vaultrc5 |
| ENSMUSG00000106106 | 139027,0953 | 1,25221394 | 0,57731149 | 2,95325014 | 0,003144 | 0,0450177 | Rn18s-rs5 |
| ENSMUSG00000089672 | 14,0767768 | 1,25011875 | 0,35585595 | 3,98986271 | 6,61E-05 | 0,0020473 | Gp49a |

| | | | | | | | |
|---|---|---|---|---|---|---|---|
| ENSMUSG00000093183 | 29,30489432 | 1,24840218 | 0,50691126 | 3,15552891 | 0,001602 | 0,0268832 | Gm25687 |
| ENSMUSG00000033222 | 5,593706507 | 1,24790922 | 0,5556129 | 3,03877296 | 0,002375 | 0,0364293 | Ttf2 |
| ENSMUSG00000066491 | 8,185777943 | 1,24013773 | 0,49904178 | 3,17792123 | 0,001483 | 0,0251838 | Gm6265 |
| ENSMUSG00000079484 | 55,939642 | 1,23871045 | 0,18789806 | 6,8491636 | 7,43E-12 | 7,941E-10 | Phyhd1 |
| ENSMUSG00000024042 | 53,56221682 | 1,23168124 | 0,18915471 | 6,77304458 | 1,26E-11 | 1,319E-09 | Sik1 |
| ENSMUSG00000031842 | 12,8723286 | 1,22998022 | 0,38809987 | 3,70930123 | 0,000208 | 0,0053084 | Pde4c |
| ENSMUSG00000026632 | 13,41362362 | 1,22527741 | 0,34398859 | 4,04458156 | 5,24E-05 | 0,0016772 | Tatdn3 |
| ENSMUSG00000026628 | 11,41625934 | 1,22184223 | 0,38085852 | 3,75479083 | 0,000173 | 0,0045769 | Atf3 |
| ENSMUSG00000091498 | 9,432024084 | 1,21159181 | 0,42594588 | 3,45427023 | 0,000552 | 0,0119562 | Mpc1-ps |
| ENSMUSG00000003549 | 15,82166754 | 1,21146558 | 0,33073885 | 4,1354034 | 3,54E-05 | 0,0011728 | Ercc1 |
| ENSMUSG00000025478 | 9,101961264 | 1,21094625 | 0,40454718 | 3,57006144 | 0,000357 | 0,0082406 | Dpysl4 |
| ENSMUSG00000014856 | 6,690243341 | 1,19521808 | 0,4961108 | 3,12578447 | 0,001773 | 0,029287 | Tmem208 |
| ENSMUSG00000015647 | 138,6425012 | 1,19223335 | 0,12541333 | 9,68397867 | 3,53E-22 | 1,212E-19 | Lama5 |
| ENSMUSG00000033590 | 34,91876205 | 1,19105699 | 0,21329869 | 5,90515532 | 3,52E-09 | 2,748E-07 | Myo5c |
| ENSMUSG00000090222 | 10,70782269 | 1,18844098 | 0,39315147 | 3,59129047 | 0,000329 | 0,0076219 | Gm16340 |
| ENSMUSG00000072809 | 4,830836592 | 1,18722288 | 0,57842608 | 2,9097368 | 0,003617 | 0,0493867 | 9330160F10Rik |
| ENSMUSG00000040479 | 100,6171537 | 1,18654039 | 0,1387686 | 8,73686313 | 2,4E-18 | 5,404E-16 | Dgkz |
| ENSMUSG00000030428 | 10,11743727 | 1,18600214 | 0,38884916 | 3,60810526 | 0,000308 | 0,0072969 | Ttyh1 |
| ENSMUSG00000002064 | 11,73560353 | 1,18237374 | 0,35117964 | 3,86526219 | 0,000111 | 0,0031217 | Sdf2 |
| ENSMUSG00000031722 | 67,59733656 | 1,1739533 | 0,43485682 | 3,33802192 | 0,000844 | 0,0165666 | Hp |
| ENSMUSG00000074886 | 28,68800563 | 1,17296335 | 0,2655282 | 4,79763246 | 1,61E-06 | 7,426E-05 | Grk6 |
| ENSMUSG00000020899 | 28,0991221 | 1,16561607 | 0,2405834 | 5,19565393 | 2,04E-07 | 1,137E-05 | Pfas |
| ENSMUSG00000038260 | 21,52240379 | 1,16043613 | 0,28261343 | 4,52716699 | 5,98E-06 | 0,0002403 | Trpm4 |
| ENSMUSG00000064366 | 76,58517786 | 1,15622653 | 0,32963772 | 3,9814668 | 6,85E-05 | 0,0021122 | mt-Tl2 |
| ENSMUSG00000004565 | 15,71953678 | 1,15397694 | 0,35463321 | 3,77873138 | 0,000158 | 0,0042282 | Pnpla6 |
| ENSMUSG00000043889 | 8,68015513 | 1,14604843 | 0,54104297 | 2,91531015 | 0,003553 | 0,0488832 | Gm8399 |
| ENSMUSG00000030231 | 53,74395136 | 1,1392668 | 0,19375764 | 6,16147993 | 7,21E-10 | 6,228E-08 | Plekha5 |
| ENSMUSG00000029022 | 15,68629489 | 1,13325114 | 0,33488261 | 3,88565098 | 0,000102 | 0,0029222 | Miip |
| ENSMUSG00000064355 | 1577,236323 | 1,13074105 | 0,43746726 | 3,22857042 | 0,001244 | 0,0218415 | mt-Tk |
| ENSMUSG00000064359 | 15114,82149 | 1,12665273 | 0,29327175 | 4,27312084 | 1,93E-05 | 0,0006785 | mt-Tg |
| ENSMUSG00000085385 | 14,52408016 | 1,12509136 | 0,33105364 | 3,88411278 | 0,000103 | 0,0029233 | Snhg17 |
| ENSMUSG00000070699 | 7,052957962 | 1,10831528 | 0,46544873 | 3,0898837 | 0,002002 | 0,0321781 | Sars2 |
| ENSMUSG00000031488 | 7,692889135 | 1,10032256 | 0,51562163 | 2,94144448 | 0,003267 | 0,0463104 | Rab11fip1 |
| ENSMUSG00000097204 | 6,609246285 | 1,09040682 | 0,52618557 | 2,91440885 | 0,003564 | 0,0489059 | Gm17690 |
| ENSMUSG00000032942 | 68,53435306 | 1,08695554 | 0,18884989 | 5,99071868 | 2,09E-09 | 1,684E-07 | Ucp3 |
| ENSMUSG00000070392 | 9,971586606 | 1,07964329 | 0,40155061 | 3,30454128 | 0,000951 | 0,0180164 | Gm20634 |
| ENSMUSG00000026950 | 41,49472096 | 1,07358145 | 0,2181635 | 5,20066306 | 1,99E-07 | 1,119E-05 | Neb |
| ENSMUSG00000088856 | 6,256976774 | 1,07342242 | 0,50867609 | 2,91460967 | 0,003561 | 0,0489059 | Gm24727 |
| ENSMUSG00000040860 | 17,60318901 | 1,07147543 | 0,30054456 | 4,0238011 | 5,73E-05 | 0,0018123 | Crocc |
| ENSMUSG00000095159 | 12,08165 | 1,06981265 | 0,3865696 | 3,37072567 | 0,00075 | 0,0150473 | Tubb4b-ps1 |
| ENSMUSG00000036833 | 47,32393391 | 1,06270521 | 0,23404142 | 4,90954739 | 9,13E-07 | 4,466E-05 | Pnpla7 |
| ENSMUSG00000044471 | 80,77230255 | 1,05477304 | 0,15830276 | 6,89650911 | 5,33E-12 | 5,783E-10 | Lincpint |
| ENSMUSG00000002625 | 83,57258152 | 1,05279347 | 0,16364826 | 6,68420656 | 2,32E-11 | 2,36E-09 | Akap8l |
| ENSMUSG00000039164 | 8,494090426 | 1,05207871 | 0,4354464 | 3,09424695 | 0,001973 | 0,0318148 | Naif1 |
| ENSMUSG00000071647 | 12,69112282 | 1,048141 | 0,40161241 | 3,23407014 | 0,00122 | 0,0215827 | Eml3 |
| ENSMUSG00000064338 | 1552,500567 | 1,04720034 | 0,4177989 | 3,15022359 | 0,001631 | 0,027218 | mt-Tv |
| ENSMUSG00000044991 | 17,77481736 | 1,04172502 | 0,36686353 | 3,41968613 | 0,000627 | 0,0131693 | 1110034G24Rik |
| ENSMUSG00000025270 | 29,10269343 | 1,03930432 | 0,30655796 | 3,87554198 | 0,000106 | 0,0030222 | Alas2 |
| ENSMUSG00000002031 | 11,6812932 | 1,03896186 | 0,38859995 | 3,29718432 | 0,000977 | 0,0184466 | Ift46 |
| ENSMUSG00000092837 | 1604,560964 | 1,03180795 | 0,28242633 | 4,10521004 | 4,04E-05 | 0,0013249 | Rpph1 |
| ENSMUSG00000030279 | 17,61130384 | 1,03099949 | 0,32047982 | 3,73825161 | 0,000185 | 0,0048182 | C2cd5 |
| ENSMUSG00000031808 | 129,669034 | 1,03043394 | 0,13601067 | 7,78405382 | 7,02E-15 | 1,109E-12 | Slc27a1 |
| ENSMUSG00000061589 | 126,9475153 | 1,02807834 | 0,12680926 | 8,30126849 | 1,03E-16 | 1,93E-14 | Dot1l |
| ENSMUSG00000043059 | 12,99024184 | 1,02597762 | 0,34375573 | 3,51738009 | 0,000436 | 0,0097059 | Zfp513 |
| ENSMUSG00000063511 | 180,869185 | 1,02516428 | 0,12885169 | 8,15466198 | 3,5E-16 | 6,238E-14 | Snrnp70 |
| ENSMUSG00000026213 | 16,73651247 | 1,02473974 | 0,3153915 | 3,75040087 | 0,000177 | 0,0046324 | Stk11ip |
| ENSMUSG00000051864 | 25,72104098 | 1,02303403 | 0,2537995 | 4,42736558 | 9,54E-06 | 0,0003566 | Tbc1d22a |

| | | | | | | | |
|---|---|---|---|---|---|---|---|
| ENSMUSG00000107018 | 10,75498864 | 1,01483457 | 0,42863361 | 3,0551735 | 0,002249 | 0,0349521 | RP23-149A1.2 |
| ENSMUSG00000025145 | 26,03021083 | 1,00854664 | 0,27369017 | 4,10094586 | 4,11E-05 | 0,0013465 | Lrrc45 |
| ENSMUSG00000032295 | 34,49001593 | 1,0034686 | 0,23646718 | 4,61711664 | 3,89E-06 | 0,0001622 | Man2c1 |
| ENSMUSG00000021431 | 24,45890935 | 1,00298516 | 0,27852313 | 4,04319757 | 5,27E-05 | 0,0016834 | Snrnp48 |
| **Down regulated genes (46)** | | | | | | | |
| ENSMUSG00000104116 | 6,529266347 | -3,45157676 | 1,23410107 | -3,30466369 | 0,0009509 | 0,0180164 | Gm37296 |
| ENSMUSG00000028989 | 6,451067114 | -3,3759533 | 1,228765186 | -3,28225151 | 0,0010298 | 0,01900421 | Angptl7 |
| ENSMUSG00000062319 | 5,068327002 | -3,22616364 | 1,231355681 | -3,21100037 | 0,0013227 | 0,0229981 | Gm10115 |
| ENSMUSG00000050390 | 5,713060689 | -2,84628877 | 1,22187405 | -3,0593149 | 0,0022184 | 0,03461004 | C77080 |
| ENSMUSG00000048047 | 5,476446868 | -2,75036997 | 1,23209349 | -3,0094028 | 0,0026176 | 0,03902831 | Zbtb33 |
| ENSMUSG00000105940 | 5,491056093 | -2,69808191 | 1,229130195 | -2,99278595 | 0,0027644 | 0,04074931 | RP23-458G12.1 |
| ENSMUSG00000107286 | 4,740337579 | -2,69518412 | 1,240291594 | -2,98114284 | 0,0028717 | 0,04177641 | RP24-122N24.2 |
| ENSMUSG00000084319 | 5,793614545 | -2,57968501 | 1,25054427 | -2,92937177 | 0,0033965 | 0,04740289 | Tpt1-ps3 |
| ENSMUSG00000023806 | 15,6458291 | -1,79783861 | 0,751071302 | -3,06426856 | 0,002182 | 0,03415278 | Rsph3b |
| ENSMUSG00000072235 | 34,65542525 | -1,61611364 | 0,474786757 | -3,90381909 | 9,469E-05 | 0,00275488 | Tuba1a |
| ENSMUSG00000094614 | 11,83647857 | -1,60873417 | 0,70722613 | -2,98827485 | 0,0028056 | 0,04118738 | Gm21769 |
| ENSMUSG00000022965 | 21,64751993 | -1,47644084 | 0,502337294 | -3,48412168 | 0,0004938 | 0,01089507 | Ifngr2 |
| ENSMUSG00000028416 | 89,34290557 | -1,42787998 | 0,245939325 | -6,10588143 | 1,022E-09 | 8,5304E-08 | Bag1 |
| ENSMUSG00000038991 | 24,7240612 | -1,41360872 | 0,511825591 | -3,35299724 | 0,0007994 | 0,01586844 | Txndc5 |
| ENSMUSG00000026511 | 20,53676869 | -1,33993096 | 0,505932399 | -3,25815664 | 0,0011214 | 0,02022838 | Srp9 |
| ENSMUSG00000022206 | 53,044906 | -1,33449799 | 0,327184802 | -4,49007715 | 7,12E-06 | 0,00027996 | Npr3 |
| ENSMUSG00000078652 | 36,94685657 | -1,28502661 | 0,398588982 | -3,7400055 | 0,000184 | 0,00479339 | Psme3 |
| ENSMUSG00000047675 | 103,3754278 | -1,28023878 | 0,244799546 | -5,54850396 | 2,881E-08 | 1,8985E-06 | Rps8 |
| ENSMUSG00000028165 | 24,73089098 | -1,27767511 | 0,478575672 | -3,2628136 | 0,0011031 | 0,01994879 | Cisd2 |
| ENSMUSG00000068479 | 33,52078188 | -1,26444603 | 0,483328667 | -3,22130277 | 0,0012761 | 0,02234866 | Mfap1a |
| ENSMUSG00000027782 | 49,28226126 | -1,23812257 | 0,344589353 | -4,0527194 | 5,063E-05 | 0,00162351 | Kpna4 |
| ENSMUSG00000101249 | 353,60672 | -1,22275733 | 0,541359712 | -3,00647591 | 0,0026429 | 0,03919878 | Gm29216 |
| ENSMUSG00000026436 | 57,97841956 | -1,21528444 | 0,301231309 | -4,47781191 | 7,541E-06 | 0,00029492 | Elk4 |
| ENSMUSG00000030224 | 36,31553506 | -1,20645759 | 0,370985473 | -3,74983016 | 0,000177 | 0,00463453 | Strap |
| ENSMUSG00000092201 | 19,66687913 | -1,20411778 | 0,505638829 | -3,02907732 | 0,002453 | 0,03722158 | A530058N18Rik |
| ENSMUSG00000037152 | 31,14331553 | -1,20173689 | 0,42842694 | -3,3842129 | 0,0007138 | 0,01450877 | Ndufc1 |
| ENSMUSG00000101939 | 213,5155819 | -1,19351391 | 0,461277024 | -3,25111955 | 0,0011495 | 0,02053051 | Gm28438 |
| ENSMUSG00000044951 | 51,05133579 | -1,12190082 | 0,294024959 | -4,23840332 | 2,251E-05 | 0,00078092 | Mylk4 |
| ENSMUSG00000032412 | 27,81053964 | -1,12095433 | 0,479330822 | -2,98784116 | 0,0028096 | 0,04120396 | Atp1b3 |
| ENSMUSG00000028234 | 46,42076275 | -1,11461871 | 0,328294545 | -3,87268015 | 0,0001076 | 0,00305192 | Rps20 |
| ENSMUSG00000101609 | 180,154871 | -1,10435402 | 0,161588906 | -7,06539229 | 1,602E-12 | 1,8344E-10 | Kcnq1ot1 |
| ENSMUSG00000061086 | 537,0582248 | -1,09228084 | 0,140271308 | -7,93983458 | 2,025E-15 | 3,32E-13 | Myl4 |
| ENSMUSG00000078866 | 45,9124198 | -1,09210715 | 0,306073485 | -4,00895329 | 6,099E-05 | 0,00190916 | Gm14420 |
| ENSMUSG00000102869 | 26,7266431 | -1,07425264 | 0,386371709 | -3,333954 | 0,0008562 | 0,01678031 | 2900097C17Rik |
| ENSMUSG00000032328 | 48,90765603 | -1,06274601 | 0,307955022 | -3,9071479 | 9,339E-05 | 0,00273375 | Tmem30a |
| ENSMUSG00000056091 | 19,65222393 | -1,05669913 | 0,46882056 | -2,91620322 | 0,0035432 | 0,04883655 | St3gal5 |
| ENSMUSG00000054408 | 29,04073287 | -1,05499903 | 0,396270682 | -3,24019408 | 0,0011945 | 0,02117641 | Spcs3 |
| ENSMUSG00000031751 | 41,40090517 | -1,04021228 | 0,33449713 | -3,60304511 | 0,0003145 | 0,00739203 | Amfr |
| ENSMUSG00000026623 | 62,3147291 | -1,02679545 | 0,261444925 | -4,30471078 | 1,672E-05 | 0,00059578 | Lpgat1 |
| ENSMUSG00000030753 | 35,68370012 | -1,02512626 | 0,356135832 | -3,41862738 | 0,0006294 | 0,0132014 | Prkrir |
| ENSMUSG00000034850 | 28,58486585 | -1,01917437 | 0,40662679 | -3,09999629 | 0,0019352 | 0,031379 | Tmem127 |
| ENSMUSG00000000711 | 26,95055521 | -1,01588146 | 0,426060809 | -3,00762144 | 0,002633 | 0,03909152 | Rab5b |
| ENSMUSG00000072704 | 41,30308636 | -1,01585497 | 0,336674422 | -3,51947943 | 0,0004324 | 0,00966105 | Smim10l1 |
| ENSMUSG00000105434 | 58,36396477 | -1,0084266 | 0,249579033 | -4,42524481 | 9,633E-06 | 0,00035922 | RP23-407I6.6 |
| ENSMUSG00000029390 | 78,78133049 | -1,0030085 | 0,246358047 | -4,46261779 | 8,096E-06 | 0,00030992 | Tmed2 |
| ENSMUSG00000026335 | 890,7236321 | -1,00277111 | 0,09702975 | -10,5089508 | 7,856E-26 | 3,5429E-23 | Pam |

**Supplementary table 3)** List of deregulated genes after VEGF-A mRNA to cardiac tissue via HUVEC SEC-EVs (total 240 deregulated genes)

| | **baseMean** | **log2FC** | **lfcSE** | **stat** | **pvalue** | **padj** | **gene_symbol** |
|---|---|---|---|---|---|---|---|
| **Up regulated genes (227)** | | | | | | | |
| ENSMUSG00000023951 | 1422490,89 | 12,524042 | 0,2510462 | 49,937662 | 0 | 0 | Vegfa |
| ENSMUSG00000069917 | 161616,978 | 10,57573 | 0,319101 | 33,213826 | 6,8E-242 | 4,706E-238 | Hba-a2 |
| ENSMUSG00000071753 | 1,94739131 | 7,2429607 | 1,5769112 | 4,5882829 | 4,469E-06 | 0,0003239 | C230004F18Rik |
| ENSMUSG00000081859 | 1,65834851 | 6,9081996 | 1,7434073 | 4,1125599 | 3,913E-05 | 0,00210767 | Olfr650-ps1 |
| ENSMUSG00000107372 | 2,22895128 | 5,9474118 | 1,6292673 | 3,8881779 | 0,000101 | 0,00438288 | AC153941.1 |
| ENSMUSG00000064952 | 2,04363543 | 4,9867255 | 1,5722366 | 3,8165699 | 0,0001353 | 0,00570613 | Gm24920 |
| ENSMUSG00000028001 | 8,10601253 | 4,3010698 | 0,9841554 | 4,8165538 | 1,461E-06 | 0,00012694 | Fga |
| ENSMUSG00000037681 | 4,88781555 | 4,0816146 | 0,7661635 | 5,7037293 | 1,172E-08 | 1,7831E-06 | Esyt3 |
| ENSMUSG00000064779 | 3,66073205 | 3,9928741 | 0,8308095 | 5,2145729 | 1,842E-07 | 2,1432E-05 | Gm22997 |
| ENSMUSG00000064795 | 3,95814316 | 3,9015849 | 0,8673104 | 4,9275688 | 8,326E-07 | 7,7875E-05 | Gm22423 |
| ENSMUSG00000027513 | 16,0863807 | 3,8050973 | 0,65324 | 6,1766025 | 6,55E-10 | 1,2592E-07 | Pck1 |
| ENSMUSG00000064632 | 4,28461658 | 3,7165057 | 0,8564029 | 4,8038213 | 1,557E-06 | 0,00013384 | Gm25142 |
| ENSMUSG00000096434 | 1,60784205 | 3,65967 | 1,4620526 | 3,241721 | 0,0011881 | 0,03012254 | Gm22129 |
| ENSMUSG00000064350 | 129,796345 | 3,5887162 | 0,6406231 | 5,9564649 | 2,578E-09 | 4,5165E-07 | mt-Ty |
| ENSMUSG00000061780 | 10,2762027 | 3,5781446 | 0,7468067 | 5,2053811 | 1,936E-07 | 2,1967E-05 | Cfd |
| ENSMUSG00000105131 | 8,1444335 | 3,4006026 | 0,8869078 | 4,3591555 | 1,306E-05 | 0,0008333 | AC069308.1 |
| ENSMUSG00000064432 | 16,3603506 | 3,2457081 | 0,6902944 | 5,1280422 | 2,928E-07 | 3,1912E-05 | Gm25587 |
| ENSMUSG00000077588 | 4,32269404 | 3,1861285 | 0,6634794 | 5,2179251 | 1,809E-07 | 2,1227E-05 | Gm24372 |
| ENSMUSG00000031710 | 11,7501427 | 3,1514764 | 1,0873139 | 3,5541509 | 0,0003792 | 0,01246861 | Ucp1 |
| ENSMUSG00000094131 | 2,00766134 | 3,1310028 | 1,0802438 | 3,5609814 | 0,0003695 | 0,0121776 | Gm22265 |
| ENSMUSG00000064437 | 2,46594876 | 3,1259186 | 1,0830427 | 3,5472074 | 0,0003893 | 0,01271133 | Snord49b |
| ENSMUSG00000029368 | 12,610912 | 3,0650293 | 0,7820897 | 4,4252585 | 9,633E-06 | 0,00064731 | Alb |
| ENSMUSG00000075760 | 4,55154418 | 3,0372076 | 0,7504311 | 4,5312454 | 5,864E-06 | 0,00040996 | Gm25414 |
| ENSMUSG00000065006 | 4,20711786 | 3,0003618 | 0,8495585 | 4,1000449 | 4,131E-05 | 0,00219085 | Gm26325 |
| ENSMUSG00000064987 | 2,14937126 | 2,9549003 | 1,1049022 | 3,3862456 | 0,0007086 | 0,02034976 | Gm25260 |
| ENSMUSG00000077711 | 3,9662387 | 2,9492776 | 0,8882324 | 3,9175292 | 8,946E-05 | 0,00402082 | AF357399 |
| ENSMUSG00000087943 | 183,401334 | 2,9359023 | 1,1368323 | 3,3687668 | 0,0007551 | 0,02115829 | Gm24245 |
| ENSMUSG00000073867 | 224,269978 | 2,9247909 | 0,7845048 | 4,2541094 | 2,099E-05 | 0,00124163 | AA474408 |
| ENSMUSG00000023034 | 38,3131327 | 2,9036896 | 0,2029558 | 14,4454 | 2,68E-47 | 9,2747E-44 | Nr4a1 |
| ENSMUSG00000077527 | 2,11726666 | 2,8895771 | 1,0652143 | 3,4205076 | 0,000625 | 0,01868787 | Gm25506 |
| ENSMUSG00000087708 | 8,92575936 | 2,882936 | 0,4736474 | 6,4061481 | 1,492E-10 | 3,1784E-08 | Gm23000 |
| ENSMUSG00000088168 | 3,03062192 | 2,8662744 | 0,9071401 | 3,7874977 | 0,0001522 | 0,0062508 | Gm23540 |
| ENSMUSG00000064349 | 52,0056397 | 2,8576633 | 0,653057 | 4,8285266 | 1,375E-06 | 0,00012128 | mt-Tc |
| ENSMUSG00000084667 | 12,5727597 | 2,8180496 | 0,8074025 | 4,0589606 | 4,929E-05 | 0,00247226 | Gm23212 |
| ENSMUSG00000088609 | 10,5069397 | 2,7882622 | 0,7300162 | 4,3376296 | 1,44E-05 | 0,00090583 | Gm24187 |
| ENSMUSG00000035769 | 31,5087615 | 2,7261091 | 0,7765461 | 4,0780956 | 4,541E-05 | 0,00233664 | Xylb |
| ENSMUSG00000064938 | 18,9997493 | 2,7046596 | 0,3911602 | 7,1973633 | 6,139E-13 | 1,6342E-10 | Mir3068 |
| ENSMUSG00000064336 | 193,216266 | 2,6644797 | 0,579954 | 5,0211976 | 5,135E-07 | 5,2268E-05 | mt-Tf |
| ENSMUSG00000064342 | 460,385125 | 2,6449393 | 0,5063229 | 5,5995897 | 2,149E-08 | 3,0663E-06 | mt-Ti |
| ENSMUSG00000098973 | 1728,61971 | 2,6339801 | 0,7564972 | 4,0475464 | 5,176E-05 | 0,00257101 | Mir6236 |
| ENSMUSG00000086324 | 89,4795022 | 2,6068298 | 0,5427419 | 5,2121761 | 1,866E-07 | 2,153E-05 | Gm15564 |
| ENSMUSG00000077756 | 4,95944624 | 2,5890404 | 0,7029922 | 4,2243743 | 2,396E-05 | 0,00137629 | Snord90 |
| ENSMUSG00000064346 | 1199,39623 | 2,5743381 | 0,6029144 | 4,727589 | 2,272E-06 | 0,00018721 | mt-Tw |
| ENSMUSG00000064352 | 1000,07658 | 2,5673541 | 0,6280981 | 4,5592059 | 5,135E-06 | 0,00036451 | mt-Ts1 |
| ENSMUSG00000077391 | 2,79634973 | 2,5659029 | 0,7334914 | 4,0681247 | 4,739E-05 | 0,00239439 | Gm24336 |
| ENSMUSG00000064372 | 2887,57802 | 2,5640041 | 0,5292925 | 5,2467084 | 1,548E-07 | 1,832E-05 | mt-Tp |
| ENSMUSG00000033831 | 7,23983722 | 2,5501772 | 0,8448125 | 3,6721113 | 0,0002406 | 0,00887183 | Fgb |
| ENSMUSG00000065017 | 10,3683213 | 2,5480878 | 0,5602743 | 4,988205 | 6,094E-07 | 6,0693E-05 | Gm23510 |

| | | | | | | | |
|---|---|---|---|---|---|---|---|
| ENSMUSG00000065160 | 5,39692553 | 2,5296418 | 0,645647 | 4,4235657 | 9,709E-06 | 0,00064925 | Mir5117 |
| ENSMUSG00000098343 | 27,1568312 | 2,5294461 | 0,4684867 | 5,7681614 | 8,014E-09 | 1,3177E-06 | Mir6240 |
| ENSMUSG00000065778 | 6,8576922 | 2,5267972 | 0,6055271 | 4,6505772 | 3,31E-06 | 0,00025494 | Gm22154 |
| ENSMUSG00000080463 | 27,4917149 | 2,516637 | 0,4762901 | 5,6606583 | 1,508E-08 | 2,2207E-06 | Gm22489 |
| ENSMUSG00000064500 | 5,15192024 | 2,5144291 | 0,651025 | 4,3700554 | 1,242E-05 | 0,00081493 | Gm25296 |
| ENSMUSG00000065881 | 27,9737715 | 2,5108602 | 0,5315974 | 5,1415507 | 2,725E-07 | 2,9936E-05 | Gm22866 |
| ENSMUSG00000022878 | 4,58847187 | 2,5075914 | 0,9318031 | 3,4154899 | 0,0006367 | 0,01895371 | Adipoq |
| ENSMUSG00000040998 | 5,52955822 | 2,5046898 | 0,4986093 | 5,4121477 | 6,227E-08 | 8,1325E-06 | Npnt |
| ENSMUSG00000004939 | 4,64071867 | 2,4734606 | 0,5907819 | 4,6502688 | 3,315E-06 | 0,00025494 | Nmrk2 |
| ENSMUSG00000065232 | 6,68627902 | 2,4492606 | 0,5845063 | 4,6676071 | 3,047E-06 | 0,00023832 | Gm22973 |
| ENSMUSG00000051851 | 1,65805045 | 2,4472554 | 1,0791559 | 3,1308598 | 0,001743 | 0,0393579 | Cxx1c |
| ENSMUSG00000033196 | 3,76961288 | 2,4376384 | 0,7744488 | 3,7774392 | 0,0001584 | 0,00647024 | Myh2 |
| ENSMUSG00000026768 | 15,7502569 | 2,4344894 | 0,3529786 | 7,1825262 | 6,843E-13 | 1,7874E-10 | Itga8 |
| ENSMUSG00000065460 | 3,82380977 | 2,4169718 | 0,7992265 | 3,6716661 | 0,000241 | 0,00887183 | Mir133a-2 |
| ENSMUSG00000075833 | 7,36705147 | 2,3679472 | 0,5167164 | 5,0181367 | 5,218E-07 | 5,272E-05 | Gm22003 |
| ENSMUSG00000077704 | 7,93400693 | 2,3480588 | 0,5214173 | 4,9426693 | 7,706E-07 | 7,3065E-05 | Snord89 |
| ENSMUSG00000065344 | 9,47516447 | 2,3435396 | 0,6142139 | 4,3367365 | 1,446E-05 | 0,00090583 | Gm25581 |
| ENSMUSG00000028341 | 20,3776362 | 2,3418198 | 0,2940837 | 8,2115112 | 2,184E-16 | 9,1625E-14 | Nr4a3 |
| ENSMUSG00000102189 | 19,202489 | 2,3304533 | 0,2917341 | 8,2360844 | 1,779E-16 | 7,6975E-14 | Gm37194 |
| ENSMUSG00000099624 | 2,62472614 | 2,3243756 | 0,7524526 | 3,7290127 | 0,0001922 | 0,00762068 | Gm7889 |
| ENSMUSG00000089855 | 73,1387976 | 2,30903 | 0,6473376 | 4,1157503 | 3,859E-05 | 0,00208684 | Gm15662 |
| ENSMUSG00000055775 | 5,2039782 | 2,3052374 | 0,5683114 | 4,5422473 | 5,566E-06 | 0,0003911 | Myh8 |
| ENSMUSG00000064344 | 1065,61371 | 2,3008423 | 0,5125941 | 4,922612 | 8,54E-07 | 7,9339E-05 | mt-Tm |
| ENSMUSG00000065729 | 8,20865683 | 2,2790684 | 0,5471749 | 4,6380137 | 3,518E-06 | 0,0002661 | Gm26176 |
| ENSMUSG00000093107 | 4,25954595 | 2,2759656 | 0,7740307 | 3,6108005 | 0,0003053 | 0,01045946 | Mir1839 |
| ENSMUSG00000064925 | 34,954363 | 2,2489524 | 0,2538517 | 9,0786915 | 1,099E-19 | 6,6138E-17 | Snora62 |
| ENSMUSG00000064343 | 433,168375 | 2,2228299 | 0,5085641 | 4,815653 | 1,467E-06 | 0,00012694 | mt-Tq |
| ENSMUSG00000021624 | 1,75503963 | 2,2152094 | 1,0621781 | 3,0361688 | 0,0023961 | 0,04976843 | Cd180 |
| ENSMUSG00000064967 | 3,07438834 | 2,2032946 | 0,9105026 | 3,2258457 | 0,001256 | 0,03121866 | Gm24405 |
| ENSMUSG00000096887 | 301,228362 | 2,1660027 | 0,1766896 | 12,422507 | 1,973E-35 | 3,4137E-32 | Gm20594 |
| ENSMUSG00000064371 | 271,285565 | 2,1624568 | 0,4786348 | 4,9480375 | 7,497E-07 | 7,2066E-05 | mt-Tt |
| ENSMUSG00000037071 | 410,526941 | 2,1490058 | 0,2708163 | 8,1888505 | 2,637E-16 | 1,0738E-13 | Scd1 |
| ENSMUSG00000065228 | 13,5281344 | 2,1314642 | 0,4347174 | 5,2978426 | 1,172E-07 | 1,4229E-05 | Gm25789 |
| ENSMUSG00000093183 | 29,3048943 | 2,1074565 | 0,4786509 | 4,8416975 | 1,287E-06 | 0,00011497 | Gm25687 |
| ENSMUSG00000064340 | 705,35181 | 2,0981126 | 0,5439123 | 4,3614297 | 1,292E-05 | 0,0008333 | mt-Tl1 |
| ENSMUSG00000075864 | 2,76461768 | 2,0227975 | 0,85009 | 3,2170377 | 0,0012952 | 0,03152042 | Gm26002 |
| ENSMUSG00000096774 | 2,41544228 | 2,0036752 | 0,8426813 | 3,2169506 | 0,0012956 | 0,03152042 | Gm22113 |
| ENSMUSG00000037362 | 15,0202097 | 1,9952601 | 0,2947277 | 7,055953 | 1,714E-12 | 4,3145E-10 | Nov |
| ENSMUSG00000059430 | 4,91243738 | 1,9866546 | 0,5856791 | 3,9713951 | 7,145E-05 | 0,00332583 | Actg2 |
| ENSMUSG00000065794 | 30,6456692 | 1,9566432 | 0,2773881 | 7,3287432 | 2,323E-13 | 6,8426E-11 | Gm25514 |
| ENSMUSG00000095118 | 34,5088043 | 1,9496174 | 0,2745891 | 7,3740825 | 1,655E-13 | 5,0906E-11 | Snord14d |
| ENSMUSG00000064338 | 1552,50057 | 1,9452921 | 0,4171226 | 5,0753198 | 3,868E-07 | 4,0879E-05 | mt-Tv |
| ENSMUSG00000064512 | 22,0069364 | 1,943302 | 0,3105188 | 6,5699477 | 5,033E-11 | 1,106E-08 | Gm22456 |
| ENSMUSG00000037313 | 4,00158996 | 1,9318396 | 0,5848462 | 3,8934996 | 9,881E-05 | 0,00430895 | Tacc3 |
| ENSMUSG00000065096 | 3,00219829 | 1,9252764 | 0,836448 | 3,146391 | 0,001653 | 0,03789549 | Gm23680 |
| ENSMUSG00000094976 | 6,38774661 | 1,9172464 | 0,6086164 | 3,7662303 | 0,0001657 | 0,00668867 | Gm25257 |
| ENSMUSG00000106474 | 5,72924112 | 1,9162887 | 0,7264879 | 3,3880873 | 0,0007038 | 0,02025564 | AC124613.1 |
| ENSMUSG00000093843 | 43,5183407 | 1,8974318 | 0,2247611 | 8,6770013 | 4,063E-18 | 2,1635E-15 | Gm25939 |
| ENSMUSG00000027559 | 13,0336545 | 1,8891564 | 0,5857663 | 3,8395488 | 0,0001233 | 0,00526635 | Car3 |
| ENSMUSG00000021250 | 32,8281058 | 1,8888271 | 0,2114288 | 9,1564021 | 5,366E-20 | 3,3762E-17 | Fos |
| ENSMUSG00000077323 | 52,7386695 | 1,8784713 | 0,2022842 | 9,4937093 | 2,23E-21 | 1,6244E-18 | Rnu11 |
| ENSMUSG00000064364 | 416,823833 | 1,8592608 | 0,529206 | 4,0607707 | 4,891E-05 | 0,00246209 | mt-Th |
| ENSMUSG00000080615 | 49,1467818 | 1,8423482 | 0,4265072 | 4,7713657 | 1,83E-06 | 0,0001554 | Snord99 |

| | | | | | | | |
|---|---|---|---|---|---|---|---|
| ENSMUSG00000075918 | 23,2409073 | 1,8408309 | 0,6975628 | 3,3751321 | 0,0007378 | 0,02092911 | n-R5s2 |
| ENSMUSG00000031722 | 67,5973366 | 1,8341338 | 0,364907 | 5,4130829 | 6,195E-08 | 8,1325E-06 | Hp |
| ENSMUSG00000064369 | 135,018515 | 1,8321229 | 0,4538074 | 4,5137728 | 6,368E-06 | 0,00044079 | mt-Te |
| ENSMUSG00000064453 | 105,717913 | 1,8234332 | 0,1461342 | 12,632354 | 1,4E-36 | 2,7691E-33 | Snord21 |
| ENSMUSG00000080542 | 10,2679621 | 1,8179859 | 0,4749866 | 4,3332133 | 1,469E-05 | 0,0009122 | Gm22710 |
| ENSMUSG00000064558 | 4,425182 | 1,8154934 | 0,6816449 | 3,4048705 | 0,000662 | 0,01937301 | Gm24620 |
| ENSMUSG00000065089 | 4,22217777 | 1,813677 | 0,7093264 | 3,3178085 | 0,0009073 | 0,02448205 | Gm26493 |
| ENSMUSG00000080478 | 11,4131476 | 1,7927343 | 0,4490988 | 4,4864596 | 7,242E-06 | 0,00049382 | Snord23 |
| ENSMUSG00000048096 | 25,5515498 | 1,7913409 | 0,2371382 | 7,795445 | 6,418E-15 | 2,3381E-12 | Lmod1 |
| ENSMUSG00000035378 | 3,55342214 | 1,7776035 | 0,7670309 | 3,1716359 | 0,0015158 | 0,03538554 | Shq1 |
| ENSMUSG00000065541 | 6,54782728 | 1,755107 | 0,7260247 | 3,216203 | 0,001299 | 0,03154721 | Mir24-2 |
| ENSMUSG00000065295 | 8,83215219 | 1,7534951 | 0,4934134 | 4,1007756 | 4,118E-05 | 0,00219085 | Gm24134 |
| ENSMUSG00000065591 | 6,78583706 | 1,743899 | 0,5858616 | 3,6374898 | 0,0002753 | 0,00975739 | Gm22753 |
| ENSMUSG00000077698 | 17,2875545 | 1,7434353 | 0,6812045 | 3,3157555 | 0,000914 | 0,02461462 | Snord12 |
| ENSMUSG00000094668 | 48,221271 | 1,7368754 | 0,216297 | 8,2721474 | 1,316E-16 | 5,8751E-14 | Gm24871 |
| ENSMUSG00000065289 | 27,4423 | 1,7351724 | 0,3452326 | 5,4099393 | 6,305E-08 | 8,1565E-06 | Gm23650 |
| ENSMUSG00000065272 | 23,2378516 | 1,7334967 | 0,3218304 | 5,7489679 | 8,979E-09 | 1,4453E-06 | Snord57 |
| ENSMUSG00000084700 | 4,41659536 | 1,7289052 | 0,682422 | 3,2926721 | 0,0009924 | 0,02627331 | Gm22984 |
| ENSMUSG00000092805 | 459,04914 | 1,715615 | 0,1652639 | 10,569905 | 4,109E-26 | 3,7922E-23 | Gm26461 |
| ENSMUSG00000106471 | 4,50560388 | 1,7134415 | 0,8037447 | 3,0472868 | 0,0023092 | 0,04880286 | mmu-mir-486b |
| ENSMUSG00000042436 | 10,7201552 | 1,7131549 | 0,3859573 | 4,8772706 | 1,076E-06 | 9,7961E-05 | Mfap4 |
| ENSMUSG00000064899 | 12,9796192 | 1,7117275 | 0,4672555 | 4,1973871 | 2,7E-05 | 0,00153818 | Snord118 |
| ENSMUSG00000080538 | 63,8314624 | 1,7105151 | 0,3790367 | 4,9369484 | 7,935E-07 | 7,4728E-05 | Gm25541 |
| ENSMUSG00000035202 | 51864,9952 | 1,7098246 | 0,7413869 | 3,1411685 | 0,0016828 | 0,03843949 | Lars2 |
| ENSMUSG00000065041 | 31,59934 | 1,7083652 | 0,2448941 | 7,2480952 | 4,227E-13 | 1,1702E-10 | Gm24044 |
| ENSMUSG00000043673 | 3,32850606 | 1,7073088 | 0,755764 | 3,1317222 | 0,0017378 | 0,03935121 | Kcns3 |
| ENSMUSG00000038418 | 86,2974387 | 1,6807549 | 0,1244978 | 13,642498 | 2,238E-42 | 5,1626E-39 | Egr1 |
| ENSMUSG00000065767 | 492,809708 | 1,6771489 | 0,4482938 | 4,2520241 | 2,118E-05 | 0,00124791 | Gm23849 |
| ENSMUSG00000064604 | 25,4909261 | 1,676408 | 0,3094395 | 5,7665499 | 8,091E-09 | 1,3177E-06 | Snora44 |
| ENSMUSG00000065378 | 7,23998795 | 1,6700662 | 0,584543 | 3,5296373 | 0,0004161 | 0,01336539 | Gm22744 |
| ENSMUSG00000077734 | 17,6655182 | 1,6699971 | 0,541766 | 3,7058714 | 0,0002107 | 0,00810066 | Snord83b |
| ENSMUSG00000105780 | 12,0516847 | 1,6602893 | 0,4128053 | 4,5038741 | 6,673E-06 | 0,00045954 | SNORD77 |
| ENSMUSG00000096263 | 35,2947427 | 1,6398349 | 0,4391775 | 4,2446098 | 2,19E-05 | 0,00128443 | Snord14c |
| ENSMUSG00000101249 | 353,60672 | 1,6312198 | 0,4794927 | 3,9612372 | 7,456E-05 | 0,00344757 | Gm29216 |
| ENSMUSG00000076281 | 36,8869971 | 1,6302722 | 0,7242635 | 3,1092651 | 0,0018755 | 0,04149297 | Gm24270 |
| ENSMUSG00000095676 | 64,9640706 | 1,6273241 | 0,3770513 | 4,757302 | 1,962E-06 | 0,00016561 | Gm25099 |
| ENSMUSG00000053963 | 3,0072257 | 1,6221144 | 0,7555736 | 3,057708 | 0,0022304 | 0,04749997 | 6330403A02Rik |
| ENSMUSG00000065287 | 25,9072664 | 1,6169302 | 0,350234 | 5,0364585 | 4,742E-07 | 4,9358E-05 | Gm24451 |
| ENSMUSG00000025889 | 11,6433288 | 1,5914875 | 0,4231749 | 4,2761298 | 1,902E-05 | 0,00113962 | Snca |
| ENSMUSG00000021741 | 2,85197994 | 1,5702133 | 0,6673579 | 3,1673408 | 0,0015384 | 0,0357917 | Gm5457 |
| ENSMUSG00000096349 | 5608,52617 | 1,5657574 | 0,1709758 | 9,3565274 | 8,24E-21 | 5,7033E-18 | Gm22513 |
| ENSMUSG00000065016 | 13,9016206 | 1,5612416 | 0,4335395 | 4,1395997 | 3,479E-05 | 0,00188869 | Snora3 |
| ENSMUSG00000064493 | 5,08377722 | 1,56057 | 0,6637371 | 3,187785 | 0,0014337 | 0,03392531 | Snora28 |
| ENSMUSG00000064853 | 10,3366783 | 1,550958 | 0,4398665 | 4,0740131 | 4,621E-05 | 0,00235178 | Gm23442 |
| ENSMUSG00000064748 | 14,0919834 | 1,5497556 | 0,3628932 | 4,7215461 | 2,341E-06 | 0,00019091 | n-R5s28 |
| ENSMUSG00000098178 | 941714,759 | 1,5459216 | 0,6279274 | 3,2276184 | 0,0012483 | 0,03119057 | Yam1 |
| ENSMUSG00000064542 | 6,52838823 | 1,5388763 | 0,5336167 | 3,5504499 | 0,0003846 | 0,01258546 | Gm22980 |
| ENSMUSG00000064355 | 1577,23632 | 1,5347983 | 0,4370113 | 4,054824 | 5,017E-05 | 0,00250734 | mt-Tk |
| ENSMUSG00000020689 | 14,6011632 | 1,5108185 | 0,362501 | 4,6294179 | 3,667E-06 | 0,00027309 | Itgb3 |
| ENSMUSG00000032085 | 38,4358139 | 1,5044764 | 0,2391176 | 6,5904435 | 4,385E-11 | 9,7909E-09 | Tagln |
| ENSMUSG00000018830 | 220,175708 | 1,5030728 | 0,1251381 | 12,155672 | 5,352E-34 | 7,9228E-31 | Myh11 |
| ENSMUSG00000065734 | 9,37637771 | 1,5002164 | 0,4925184 | 3,6840573 | 0,0002296 | 0,0086585 | Snord49a |
| ENSMUSG00000064823 | 9,92804174 | 1,4953685 | 0,4903068 | 3,6753454 | 0,0002375 | 0,00887183 | Snord82 |

| | | | | | | | |
|---|---|---|---|---|---|---|---|
| ENSMUSG00000065145 | 192,946589 | 1,4750404 | 0,2516428 | 6,1842426 | 6,24E-10 | 1,2167E-07 | Vaultrc5 |
| ENSMUSG00000064366 | 76,5851779 | 1,4722672 | 0,3187089 | 5,0277964 | 4,961E-07 | 5,0875E-05 | mt-Tl2 |
| ENSMUSG00000064844 | 12,001295 | 1,4683855 | 0,4783617 | 3,7024995 | 0,0002135 | 0,00816377 | Gm26202 |
| ENSMUSG00000059195 | 34,2568239 | 1,4659316 | 0,208932 | 7,2880571 | 3,145E-13 | 8,8837E-11 | Gm12715 |
| ENSMUSG00000077563 | 21,8203026 | 1,4652358 | 0,3102591 | 5,1233585 | 3,001E-07 | 3,246E-05 | Snora68 |
| ENSMUSG00000065147 | 13,6778163 | 1,4470194 | 0,4292617 | 3,9458629 | 7,951E-05 | 0,0036447 | Snora31 |
| ENSMUSG00000028195 | 91,0100192 | 1,4448027 | 0,1418624 | 10,371157 | 3,354E-25 | 2,9022E-22 | Cyr61 |
| ENSMUSG00000076258 | 630,0724 | 1,4307234 | 0,5378895 | 3,3690373 | 0,0007543 | 0,02115829 | Gm23935 |
| ENSMUSG00000079225 | 6,44614378 | 1,4300716 | 0,6243413 | 3,1166219 | 0,0018294 | 0,0408449 | Gm9531 |
| ENSMUSG00000064816 | 48,2520977 | 1,4283911 | 0,4421235 | 3,8160321 | 0,0001356 | 0,00570613 | Gm22357 |
| ENSMUSG00000028464 | 80,8241015 | 1,4277003 | 0,1417416 | 10,235474 | 1,375E-24 | 1,0576E-21 | Tpm2 |
| ENSMUSG00000064339 | 1179305,31 | 1,4229729 | 0,1812889 | 8,0853427 | 6,199E-16 | 2,4518E-13 | mt-Rnr2 |
| ENSMUSG00000080610 | 15,8287307 | 1,4012989 | 0,4218185 | 3,8964243 | 9,762E-05 | 0,00430104 | Snord110 |
| ENSMUSG00000025470 | 4,5574763 | 1,3952511 | 0,5404598 | 3,3262909 | 0,0008801 | 0,02398273 | Zfp511 |
| ENSMUSG00000105884 | 21,261509 | 1,3938586 | 0,4259167 | 3,8583184 | 0,0001142 | 0,00489304 | SNORA50 |
| ENSMUSG00000080486 | 15,3424984 | 1,3849928 | 0,4371541 | 3,7702051 | 0,0001631 | 0,00662164 | Snord100 |
| ENSMUSG00000064829 | 5,22272944 | 1,3849503 | 0,6244618 | 3,1010017 | 0,0019287 | 0,04237874 | Gm22879 |
| ENSMUSG00000064655 | 19,6256462 | 1,3849255 | 0,318415 | 4,7856762 | 1,704E-06 | 0,00014562 | Gm25788 |
| ENSMUSG00000077565 | 31,549369 | 1,3779786 | 0,2677419 | 5,5153287 | 3,481E-08 | 4,8191E-06 | Gm23201 |
| ENSMUSG00000105863 | 14,1054059 | 1,37619 | 0,4613383 | 3,6182409 | 0,0002966 | 0,01026501 | SNORA17 |
| ENSMUSG00000057378 | 36,8434561 | 1,3749775 | 0,2285211 | 6,3192645 | 2,628E-10 | 5,3501E-08 | Ryr3 |
| ENSMUSG00000064721 | 9,36489044 | 1,3711364 | 0,4559695 | 3,6467584 | 0,0002656 | 0,00949942 | Gm25855 |
| ENSMUSG00000064563 | 23,4314761 | 1,3679383 | 0,2785349 | 5,2943444 | 1,194E-07 | 1,4378E-05 | Snora7a |
| ENSMUSG00000052305 | 284,260898 | 1,3608367 | 0,1148474 | 12,005485 | 3,325E-33 | 3,8357E-30 | Hbb-bs |
| ENSMUSG00000064359 | 15114,8215 | 1,3418577 | 0,2932158 | 4,9669906 | 6,8E-07 | 6,624E-05 | mt-Tg |
| ENSMUSG00000005483 | 16,8398541 | 1,3412041 | 0,286806 | 5,0816028 | 3,743E-07 | 3,9853E-05 | Dnajb1 |
| ENSMUSG00000106585 | 15,6136049 | 1,3392693 | 0,3871599 | 4,0199853 | 5,82E-05 | 0,00282697 | SNORA17 |
| ENSMUSG00000088895 | 12,4276309 | 1,3390117 | 0,4423889 | 3,65707 | 0,0002551 | 0,0092449 | Gm22519 |
| ENSMUSG00000044937 | 4,8992507 | 1,333503 | 0,5731769 | 3,1614007 | 0,0015701 | 0,03640741 | Ttc41 |
| ENSMUSG00000088948 | 91,0681148 | 1,3317748 | 0,1983314 | 6,9918504 | 2,713E-12 | 6,706E-10 | Gm23262 |
| ENSMUSG00000065196 | 20,21421 | 1,328605 | 0,3745906 | 4,0771557 | 4,559E-05 | 0,00233741 | Snord85 |
| ENSMUSG00000020423 | 59,1933829 | 1,3045768 | 0,1772483 | 7,6148528 | 2,64E-14 | 8,7011E-12 | Btg2 |
| ENSMUSG00000077575 | 13,5969567 | 1,296534 | 0,4210728 | 3,6938007 | 0,0002209 | 0,0084019 | Gm24888 |
| ENSMUSG00000068745 | 28,9253189 | 1,2948375 | 0,240454 | 5,7324122 | 9,901E-09 | 1,54E-06 | Mybphl |
| ENSMUSG00000093178 | 13,6279753 | 1,2947902 | 0,3836858 | 3,9396402 | 8,16E-05 | 0,00371593 | Snord87 |
| ENSMUSG00000064968 | 11,8377574 | 1,2764513 | 0,4173941 | 3,6878681 | 0,0002261 | 0,00855319 | Snord47 |
| ENSMUSG00000087385 | 4,42598282 | 1,26493 | 0,5552621 | 3,1113777 | 0,0018622 | 0,04137714 | AA415398 |
| ENSMUSG00000077167 | 42,5452267 | 1,2646404 | 0,2560164 | 5,3161547 | 1,06E-07 | 1,2983E-05 | Gm24119 |
| ENSMUSG00000065905 | 12,7820747 | 1,2645259 | 0,4505798 | 3,484808 | 0,0004925 | 0,01532032 | Gm26110 |
| ENSMUSG00000064768 | 7,27396816 | 1,2480941 | 0,5468967 | 3,1145908 | 0,001842 | 0,04106094 | Snord60 |
| ENSMUSG00000035783 | 83,857258 | 1,2448887 | 0,1680265 | 7,6542815 | 1,944E-14 | 6,7276E-12 | Acta2 |
| ENSMUSG00000039828 | 16,798452 | 1,2441515 | 0,3235866 | 4,330492 | 1,488E-05 | 0,00091943 | Wdr70 |
| ENSMUSG00000091825 | 8,37524598 | 1,2420496 | 0,4606679 | 3,4052229 | 0,0006611 | 0,01937301 | Gm5778 |
| ENSMUSG00000092680 | 40,7376227 | 1,2319378 | 0,2864808 | 4,7305795 | 2,239E-06 | 0,00018558 | Snord55 |
| ENSMUSG00000092730 | 32,1211032 | 1,2297723 | 0,2779573 | 4,8367245 | 1,32E-06 | 0,00011713 | Snora24 |
| ENSMUSG00000040435 | 45,2662035 | 1,2277098 | 0,210693 | 6,1398718 | 8,259E-10 | 1,5661E-07 | Ppp1r15a |
| ENSMUSG00000038894 | 46,2473584 | 1,2251157 | 0,2220334 | 5,8426089 | 5,139E-09 | 8,6754E-07 | Irs2 |
| ENSMUSG00000053560 | 29,0980229 | 1,2207482 | 0,2346175 | 5,5635655 | 2,643E-08 | 3,7336E-06 | Ier2 |
| ENSMUSG00000090698 | 20,1144695 | 1,2167559 | 0,2657962 | 4,9805871 | 6,339E-07 | 6,2681E-05 | Apold1 |
| ENSMUSG00000065118 | 10,558085 | 1,2137735 | 0,4347719 | 3,4704732 | 0,0005195 | 0,01598227 | Gm23297 |
| ENSMUSG00000065353 | 28,8944002 | 1,2110594 | 0,3264722 | 4,2138651 | 2,51E-05 | 0,001436 | Snora73b |
| ENSMUSG00000065701 | 98,1930442 | 1,204675 | 0,4345064 | 3,4348819 | 0,0005928 | 0,01787864 | Rny1 |
| ENSMUSG00000090877 | 12,6242533 | 1,1875496 | 0,3870788 | 3,6802261 | 0,000233 | 0,0087591 | Hspa1b |

| | | | | | | | |
|---|---|---|---|---|---|---|---|
| ENSMUSG00000088595 | 21,9364022 | 1,1687616 | 0,3367503 | 4,005314 | 6,194E-05 | 0,00297697 | Gm24601 |
| ENSMUSG00000064901 | 23,1360104 | 1,1662153 | 0,3004961 | 4,3590527 | 1,306E-05 | 0,0008333 | Snora21 |
| ENSMUSG00000019997 | 301,96488 | 1,1341136 | 0,1013233 | 11,356112 | 6,915E-30 | 6,8379E-27 | Ctgf |
| ENSMUSG00000039377 | 9,07990666 | 1,1235182 | 0,4020109 | 3,4509909 | 0,0005585 | 0,01699289 | Hlx |
| ENSMUSG00000065878 | 8,31408116 | 1,0898016 | 0,4916199 | 3,0638972 | 0,0021847 | 0,04667184 | Snord34 |
| ENSMUSG00000091997 | 14,0909246 | 1,0831044 | 0,3027856 | 4,0969196 | 4,187E-05 | 0,0021954 | Gm6611 |
| ENSMUSG00000045996 | 7,63315069 | 1,0810471 | 0,4907261 | 3,0438168 | 0,002336 | 0,04899528 | Polr2k |
| ENSMUSG00000026532 | 40,7113804 | 1,0801321 | 0,2029946 | 5,6830263 | 1,323E-08 | 1,9912E-06 | Spta1 |
| ENSMUSG00000080465 | 62,4918843 | 1,0782261 | 0,2547916 | 4,6528829 | 3,273E-06 | 0,00025456 | Gm22486 |
| ENSMUSG00000064353 | 293,328967 | 1,0717311 | 0,3812412 | 3,4522628 | 0,0005559 | 0,01698765 | mt-Td |
| ENSMUSG00000020527 | 13,8628319 | 1,0706276 | 0,3181716 | 3,9048369 | 9,429E-05 | 0,00422408 | Myo19 |
| ENSMUSG00000050440 | 10,0251812 | 1,0697504 | 0,4714437 | 3,0986266 | 0,0019442 | 0,04258473 | Hamp |
| ENSMUSG00000073940 | 169,586462 | 1,0535084 | 0,2614913 | 4,4631538 | 8,076E-06 | 0,00054803 | Hbb-bt |
| ENSMUSG00000037411 | 85,8369724 | 1,0447123 | 0,1375176 | 7,848986 | 4,194E-15 | 1,5692E-12 | Serpine1 |
| ENSMUSG00000037243 | 15,3325039 | 1,0421297 | 0,3576565 | 3,5303983 | 0,0004149 | 0,013358 | Zfp692 |
| ENSMUSG00000065899 | 23,360552 | 1,0269258 | 0,3334025 | 3,6738075 | 0,000239 | 0,00887183 | Gm24523 |
| ENSMUSG00000010936 | 14,523247 | 1,0208161 | 0,3146805 | 3,7902568 | 0,0001505 | 0,00621867 | Vac14 |
| ENSMUSG00000024190 | 57,5638455 | 1,0083781 | 0,1864843 | 5,7357441 | 9,709E-09 | 1,5272E-06 | Dusp1 |
| ENSMUSG00000030838 | 18,5993621 | 1,0056246 | 0,3939621 | 3,2538123 | 0,0011387 | 0,0292443 | Ush1c |
| ENSMUSG00000064337 | 546894,166 | 1,0024266 | 0,1419467 | 7,3105416 | 2,661E-13 | 7,6733E-11 | mt-Rnr1 |
| **Down regulated genes (13)** | | | | | | | |
| | **baseMean** | **log2FC** | **lfcSE** | **stat** | **pvalue** | **padj** | **gene_symbol** |
| ENSMUSG00000075015 | 224,30246 | -4,389288 | 0,3929886 | -11,36085 | 6,55E-30 | 6,8379E-27 | Gm10801 |
| ENSMUSG00000075014 | 192,385489 | -3,761894 | 0,2298853 | -16,4621 | 6,867E-61 | 3,1687E-57 | Gm10800 |
| ENSMUSG00000045319 | 5,57076151 | -3,688588 | 1,2198345 | -3,45904 | 0,0005421 | 0,01663936 | Proser2 |
| ENSMUSG00000097312 | 378,521561 | -3,559438 | 0,4418655 | -8,30547 | 9,943E-17 | 4,6927E-14 | Gm26870 |
| ENSMUSG00000095280 | 42,3476465 | -3,493979 | 0,4062446 | -8,814035 | 1,207E-18 | 6,9631E-16 | Gm21738 |
| ENSMUSG00000097750 | 9,19754524 | -2,990667 | 1,2036547 | -3,219057 | 0,0012861 | 0,03145566 | Gm4673 |
| ENSMUSG00000095547 | 11,9046106 | -2,679067 | 0,687543 | -4,359243 | 1,305E-05 | 0,0008333 | Gm10719 |
| ENSMUSG00000104211 | 4,78200832 | -2,563598 | 1,0672274 | -3,070612 | 0,0021362 | 0,04584735 | Gm37985 |
| ENSMUSG00000026605 | 28,5935471 | -1,35121 | 0,3257345 | -4,573079 | 4,806E-06 | 0,00034516 | Cenpf |
| ENSMUSG00000049303 | 17,3284393 | -1,21921 | 0,46568 | -3,282572 | 0,0010286 | 0,02707139 | Syt12 |
| ENSMUSG00000064357 | 6156,97248 | -1,045298 | 0,1527862 | -7,092993 | 1,312E-12 | 3,3644E-10 | mt-Atp6 |
| ENSMUSG00000039349 | 17,8142139 | -1,039668 | 0,4453991 | -3,044851 | 0,002328 | 0,04890126 | C130074G19Rik |
| ENSMUSG00000101939 | 213,515582 | -1,027405 | 0,4521994 | -3,065685 | 0,0021717 | 0,04653728 | Gm28438 |

| **Supplementary table 4)** List of deregulated genes after VEGF-A mRNA to cardiac tissue via LNPs (total 437) | | | | | | | |
|---|---|---|---|---|---|---|---|
| | **baseMean** | **log2FC** | **lfcSE** | **stat** | **pvalue** | **padj** | **gene_symbol** |
| **Up regulated genes (365)** | | | | | | | |
| ENSMUSG00000027483 | 22,4968764 | 11,6478929 | 1,196345712 | 8,878064678 | 6,80337E-19 | 1,1829E-16 | Bpifa1 |
| ENSMUSG00000024653 | 18,41835271 | 11,5363767 | 1,204792879 | 8,727327322 | 2,60762E-18 | 4,0032E-16 | Scgb1a1 |
| ENSMUSG00000023951 | 1422490,887 | 10,6643785 | 0,251046413 | 42,53787336 | 0 | 0 | Vegfa |
| ENSMUSG00000032496 | 8,019986346 | 9,99870826 | 1,202532764 | 7,689597925 | 1,47598E-14 | 1,5214E-12 | Ltf |
| ENSMUSG00000069917 | 161616,9778 | 8,88693418 | 0,319101618 | 27,93338569 | 1,0493E-171 | 5,048E-168 | Hba-a2 |
| ENSMUSG00000052974 | 3,646646261 | 8,78954985 | 1,266652866 | 6,411478939 | 1,44115E-10 | 8,2857E-09 | Cyp2f2 |
| ENSMUSG00000066108 | 24,19261272 | 7,79859762 | 0,764796628 | 10,39105657 | 2,7232E-25 | 7,8597E-23 | Muc5b |
| ENSMUSG00000038791 | 1,533006653 | 6,90029486 | 1,438249837 | 4,652036906 | 3,28672E-06 | 7,9984E-05 | Scgb3a2 |
| ENSMUSG00000031710 | 11,75014271 | 6,69250326 | 0,920933521 | 7,531070572 | 5,0326E-14 | 4,8096E-12 | Ucp1 |
| ENSMUSG00000017723 | 1,313658774 | 6,49759246 | 1,631725049 | 4,02035542 | 5,81104E-05 | 0,00099478 | Wfdc2 |
| ENSMUSG00000030017 | 4,733111454 | 6,31551315 | 1,124940424 | 5,946785257 | 2,7346E-09 | 1,2981E-07 | Reg3g |
| ENSMUSG00000026390 | 1,405585351 | 6,13698364 | 1,680091737 | 3,828707538 | 0,000128818 | 0,00199033 | Marco |
| ENSMUSG00000030278 | 7,091233119 | 6,11765859 | 0,884586636 | 7,20690852 | 5,72364E-13 | 4,5135E-11 | Cidec |
| ENSMUSG00000030546 | 4,66970807 | 5,56971014 | 1,105557357 | 5,403853725 | 6,52241E-08 | 2,3531E-06 | Plin1 |
| ENSMUSG00000061780 | 10,27620268 | 5,35059958 | 0,635048174 | 8,665445589 | 4,4975E-18 | 6,4903E-16 | Cfd |
| ENSMUSG00000037071 | 410,5269406 | 5,34662703 | 0,251795399 | 21,32435267 | 6,7481E-101 | 1,082E-97 | Scd1 |
| ENSMUSG00000027513 | 16,08638075 | 5,31491951 | 0,587301544 | 9,277083038 | 1,74182E-20 | 3,5403E-18 | Pck1 |
| ENSMUSG00000027485 | 5,045329794 | 5,13403422 | 0,745716164 | 7,149991054 | 8,67836E-13 | 6,7332E-11 | Bpifb1 |
| ENSMUSG00000054986 | 1,668836031 | 5,10785512 | 1,416014826 | 4,105001225 | 4,04313E-05 | 0,00072842 | Sec14l3 |
| ENSMUSG00000076609 | 8,962699561 | 5,08819172 | 0,696166372 | 7,575479505 | 3,57804E-14 | 3,5126E-12 | Igkc |
| ENSMUSG00000005547 | 1,781961448 | 4,93061625 | 1,53890247 | 3,791502952 | 0,000149738 | 0,0022746 | Cyp2a5 |
| ENSMUSG00000064057 | 5,995743084 | 4,90609125 | 0,62496019 | 8,094465014 | 5,75166E-16 | 6,8597E-14 | Scgb3a1 |
| ENSMUSG00000031722 | 67,59733656 | 4,87561573 | 0,274631561 | 17,86662092 | 2,14607E-71 | 1,9356E-68 | Hp |
| ENSMUSG00000076617 | 15,54118982 | 4,74748457 | 0,441977747 | 10,95511096 | 6,28014E-28 | 2,2657E-25 | Ighm |
| ENSMUSG00000067149 | 2,129741248 | 4,73549197 | 1,200585307 | 4,406804507 | 1,04907E-05 | 0,00022462 | Jchain |
| ENSMUSG00000027559 | 13,03365451 | 4,58029464 | 0,405390636 | 11,48424918 | 1,58305E-30 | 7,3693E-28 | Car3 |
| ENSMUSG00000022878 | 4,588471869 | 4,38519215 | 0,753609625 | 6,137778128 | 8,36836E-10 | 4,2373E-08 | Adipoq |
| ENSMUSG00000026417 | 4,073495189 | 3,72719917 | 0,704516098 | 5,655796584 | 1,55125E-08 | 6,47E-07 | Pigr |
| ENSMUSG00000025153 | 170,5779424 | 3,53507145 | 0,154915606 | 22,92013824 | 2,9265E-116 | 8,446E-113 | Fasn |
| ENSMUSG00000021950 | 5,98199759 | 3,52027946 | 0,49624795 | 7,367063633 | 1,74427E-13 | 1,5349E-11 | Anxa8 |
| ENSMUSG00000023043 | 8,74186796 | 3,21161576 | 0,408040641 | 8,109861623 | 5,06774E-16 | 6,1456E-14 | Krt18 |
| ENSMUSG00000079620 | 2,964693647 | 3,20786056 | 0,762993448 | 4,66682405 | 3,05891E-06 | 7,4946E-05 | Muc4 |
| ENSMUSG00000029372 | 4,62087332 | 3,15485425 | 0,53322437 | 6,247592417 | 4,16828E-10 | 2,2034E-08 | Ppbp |
| ENSMUSG00000020911 | 7,577937675 | 3,10155576 | 0,434788306 | 7,414100883 | 1,22453E-13 | 1,1401E-11 | Krt19 |
| ENSMUSG00000033576 | 2,932997999 | 3,09481612 | 0,873594503 | 4,080499656 | 4,4939E-05 | 0,00080262 | Apol6 |
| ENSMUSG00000042985 | 4,656223055 | 3,09373711 | 0,616732503 | 5,415808939 | 6,10122E-08 | 2,2122E-06 | Upk3b |
| ENSMUSG00000002997 | 7,856569755 | 3,07692182 | 0,401742242 | 7,919611133 | 2,38255E-15 | 2,7288E-13 | Prkar2b |
| ENSMUSG00000027070 | 4,896084986 | 3,0757821 | 0,594796919 | 5,548607287 | 2,87954E-08 | 1,1201E-06 | Lrp2 |
| ENSMUSG00000024371 | 1,50344304 | 3,07153839 | 1,387208319 | 3,064282977 | 0,002181924 | 0,02096072 | C2 |
| ENSMUSG00000040809 | 4,996507879 | 2,93761302 | 0,544207883 | 5,75352305 | 8,74025E-09 | 3,8291E-07 | Chil3 |
| ENSMUSG00000041220 | 7,788570463 | 2,89831837 | 0,437039858 | 6,927155831 | 4,29386E-12 | 3,0828E-10 | Elovl6 |
| ENSMUSG00000091694 | 2,310615337 | 2,85796947 | 0,798339152 | 4,113021243 | 3,90514E-05 | 0,00070709 | Apol11b |
| ENSMUSG00000101249 | 353,60672 | 2,83867313 | 0,475577908 | 6,298304604 | 3,00919E-10 | 1,6264E-08 | Gm29216 |
| ENSMUSG00000056071 | 6,850853806 | 2,80224487 | 0,402311937 | 7,233160891 | 4,7188E-13 | 3,8257E-11 | S100a9 |
| ENSMUSG00000033022 | 5,437504488 | 2,78389216 | 0,47955252 | 6,132064106 | 8,67461E-10 | 4,3618E-08 | Cdo1 |
| ENSMUSG00000019987 | 6,200283373 | 2,7711961 | 0,431432603 | 6,722096426 | 1,79128E-11 | 1,1968E-09 | Arg1 |
| ENSMUSG00000090231 | 5,885814193 | 2,76569748 | 0,464993086 | 6,269800863 | 3,6151E-10 | 1,9322E-08 | Cfb |
| ENSMUSG00000031489 | 1,608203768 | 2,71339983 | 1,272509199 | 2,9895515 | 0,002793873 | 0,02545353 | Adrb3 |
| ENSMUSG00000060962 | 1,395767894 | 2,67775581 | 1,0115638 | 3,340566639 | 0,000836076 | 0,00974808 | Dmkn |
| ENSMUSG00000053310 | 8,117702539 | 2,6203213 | 0,399365317 | 6,863673061 | 6,71121E-12 | 4,6787E-10 | Nrgn |
| ENSMUSG00000043110 | 3,691337808 | 2,6053365 | 0,671801431 | 4,380811332 | 1,18238E-05 | 0,00024765 | Lrrn4 |
| ENSMUSG00000006574 | 3,517131249 | 2,60317382 | 0,622287557 | 4,639592302 | 3,49097E-06 | 8,4386E-05 | Slc4a1 |

| | | | | | | | |
|---|---|---|---|---|---|---|---|
| ENSMUSG00000018927 | 25,14974754 | 2,56715814 | 0,23982968 | 10,90183526 | 1,12956E-27 | 3,9758E-25 | Ccl6 |
| ENSMUSG00000061048 | 4,081052242 | 2,53876433 | 0,671497679 | 4,290780295 | 1,78046E-05 | 0,00035101 | Cdh3 |
| ENSMUSG00000031373 | 3,424417428 | 2,51999312 | 0,732137142 | 3,98945305 | 6,62258E-05 | 0,0011204 | Car5b |
| ENSMUSG00000005360 | 8,559935419 | 2,49579528 | 0,365360482 | 7,112436885 | 1,14012E-12 | 8,7053E-11 | Slc1a3 |
| ENSMUSG00000047139 | 7,864555315 | 2,48374707 | 0,335754644 | 7,655898748 | 1,91965E-14 | 1,9509E-12 | Cd24a |
| ENSMUSG00000003528 | 6,637198629 | 2,46042856 | 0,417207505 | 6,225428407 | 4,80242E-10 | 2,5293E-08 | Slc25a1 |
| ENSMUSG00000021922 | 20,49153041 | 2,42217241 | 0,477466992 | 5,456830048 | 4,8471E-08 | 1,789E-06 | Itih4 |
| ENSMUSG00000028001 | 8,106012535 | 2,40116397 | 0,983760722 | 3,174352879 | 0,00150171 | 0,0156245 | Fga |
| ENSMUSG00000046844 | 1,495077783 | 2,39748116 | 1,378283498 | 2,718278212 | 0,006562263 | 0,04941971 | Vat1l |
| ENSMUSG00000049382 | 14,94020527 | 2,37019468 | 0,322383449 | 7,612913019 | 2,67986E-14 | 2,6671E-12 | Krt8 |
| ENSMUSG00000026180 | 3,867090046 | 2,32286558 | 0,523896549 | 4,868360778 | 1,12528E-06 | 3,0639E-05 | Cxcr2 |
| ENSMUSG00000042045 | 35,39985993 | 2,30996857 | 0,16841897 | 13,86073154 | 1,09552E-43 | 7,1861E-41 | Sln |
| ENSMUSG00000050440 | 10,0251812 | 2,29728257 | 0,327778886 | 7,284018483 | 3,24021E-13 | 2,7045E-11 | Hamp |
| ENSMUSG00000025479 | 5,638596614 | 2,28700182 | 0,494978368 | 5,039835834 | 4,65931E-07 | 1,3979E-05 | Cyp2e1 |
| ENSMUSG00000025150 | 10,15844685 | 2,27550228 | 0,305310284 | 7,710830911 | 1,25001E-14 | 1,3167E-12 | Cbr2 |
| ENSMUSG00000065344 | 9,475164474 | 2,25523451 | 0,531160723 | 4,706573565 | 2,51915E-06 | 6,3114E-05 | Gm25581 |
| ENSMUSG00000053897 | 21,18068601 | 2,24966926 | 0,236944306 | 9,695086237 | 3,1637E-22 | 7,2469E-20 | Slc39a8 |
| ENSMUSG00000055653 | 9,698513982 | 2,22975089 | 0,366328725 | 6,402835702 | 1,52517E-10 | 8,6653E-09 | Gpc3 |
| ENSMUSG00000026582 | 7,278462018 | 2,22818044 | 0,342791233 | 6,790731728 | 1,11566E-11 | 7,5587E-10 | Sele |
| ENSMUSG00000050335 | 8,530868177 | 2,19915363 | 0,345871943 | 6,658539022 | 2,76563E-11 | 1,7978E-09 | Lgals3 |
| ENSMUSG00000024164 | 263,9459887 | 2,19853082 | 0,089883301 | 24,51962026 | 9,1249E-133 | 3,292E-129 | C3 |
| ENSMUSG00000020532 | 70,92226137 | 2,19038229 | 0,137247085 | 16,06833088 | 4,25387E-58 | 3,4104E-55 | Acaca |
| ENSMUSG00000029482 | 14,47989806 | 2,18883998 | 0,253872683 | 8,842505972 | 9,35953E-19 | 1,5706E-16 | Aacs |
| ENSMUSG00000059430 | 4,912437384 | 2,18122646 | 0,482495624 | 4,941945427 | 7,73469E-07 | 2,1886E-05 | Actg2 |
| ENSMUSG00000041607 | 5,901831153 | 2,17935378 | 0,497217115 | 4,827703803 | 1,38116E-06 | 3,6979E-05 | Mbp |
| ENSMUSG00000064632 | 4,284616582 | 2,16663095 | 0,84275818 | 3,320075922 | 0,00089993 | 0,01034812 | Gm25142 |
| ENSMUSG00000027907 | 13,85280805 | 2,15953873 | 0,251328122 | 8,784564894 | 1,56971E-18 | 2,5076E-16 | S100a11 |
| ENSMUSG00000020469 | 184,8003693 | 2,12440801 | 0,095002198 | 22,44681875 | 1,3746E-111 | 3,306E-108 | Myl7 |
| ENSMUSG00000045763 | 3,345474645 | 2,0958526 | 0,660452275 | 3,762925197 | 0,000167937 | 0,0025088 | Basp1 |
| ENSMUSG00000038725 | 33,33321092 | 2,0930659 | 0,164648941 | 12,86245503 | 7,32138E-38 | 4,5937E-35 | Pkhd1l1 |
| ENSMUSG00000025889 | 11,64332885 | 2,08708503 | 0,323567841 | 6,745051552 | 1,52973E-11 | 1,0316E-09 | Snca |
| ENSMUSG00000031517 | 6,763198528 | 2,07372497 | 0,390840436 | 5,667262814 | 1,45097E-08 | 6,0692E-07 | Gpm6a |
| ENSMUSG00000023078 | 3,07450001 | 2,05639863 | 0,612716462 | 3,917653761 | 8,9415E-05 | 0,00145473 | Cxcl13 |
| ENSMUSG00000037681 | 4,887815546 | 2,02839165 | 0,775588354 | 3,333965575 | 0,000856173 | 0,00992404 | Esyt3 |
| ENSMUSG00000011263 | 1,614038545 | 2,0282716 | 0,957675623 | 3,003034091 | 0,002673025 | 0,0244917 | Exoc3l2 |
| ENSMUSG00000056054 | 2,73699154 | 2,02448466 | 0,679946107 | 3,60902468 | 0,00030735 | 0,00417643 | S100a8 |
| ENSMUSG00000035385 | 11,72803848 | 2,00595349 | 0,341342331 | 6,201955306 | 5,57659E-10 | 2,9158E-08 | Ccl2 |
| ENSMUSG00000068745 | 28,92531893 | 2,00353107 | 0,172460312 | 11,78690724 | 4,5598E-32 | 2,2691E-29 | Mybphl |
| ENSMUSG00000031906 | 15,34442213 | 2,00163669 | 0,288505461 | 7,213142198 | 5,46753E-13 | 4,3353E-11 | Smpd3 |
| ENSMUSG00000025270 | 29,10269343 | 1,99052316 | 0,18947733 | 10,68745048 | 1,1653E-26 | 3,6558E-24 | Alas2 |
| ENSMUSG00000047798 | 7,929406089 | 1,98239562 | 0,358697198 | 5,868617029 | 4,39445E-09 | 2,0005E-07 | Cd300lf |
| ENSMUSG00000074677 | 1,733066901 | 1,98000882 | 0,851450003 | 3,129052555 | 0,001753709 | 0,01757485 | Sirpb1c |
| ENSMUSG00000021069 | 9,143016836 | 1,97164527 | 0,335300176 | 6,208190909 | 5,3598E-10 | 2,8126E-08 | Pygl |
| ENSMUSG00000033220 | 2,98314865 | 1,94000927 | 0,613505036 | 3,758662545 | 0,000170824 | 0,00254929 | Rac2 |
| ENSMUSG00000035769 | 31,50876154 | 1,93434063 | 0,768953514 | 3,285650689 | 0,001017471 | 0,01145107 | Xylb |
| ENSMUSG00000021728 | 3,947550909 | 1,9236222 | 0,475304062 | 4,516561151 | 6,2852E-06 | 0,00014172 | Emb |
| ENSMUSG00000063011 | 6,400087865 | 1,9222776 | 0,411895635 | 5,078344837 | 3,80737E-07 | 1,1816E-05 | Msln |
| ENSMUSG00000026579 | 7,398850774 | 1,92164702 | 0,437835572 | 4,822777142 | 1,41573E-06 | 3,7625E-05 | F5 |
| ENSMUSG00000088609 | 10,50693966 | 1,91803383 | 0,706865246 | 3,420425958 | 0,000625232 | 0,00760769 | Gm24187 |
| ENSMUSG00000026580 | 19,71991735 | 1,9152799 | 0,205712643 | 9,50348123 | 2,02988E-21 | 4,3721E-19 | Selp |
| ENSMUSG00000001349 | 8,486717922 | 1,91161164 | 0,36151779 | 5,648152495 | 1,62181E-08 | 6,7061E-07 | Cnn1 |
| ENSMUSG00000025105 | 1,261183055 | 1,90528217 | 1,298508247 | 2,723247991 | 0,006464351 | 0,0488935 | Bnc1 |
| ENSMUSG00000022425 | 7,828630414 | 1,87540035 | 0,327690861 | 6,069302217 | 1,28467E-09 | 6,3273E-08 | Enpp2 |
| ENSMUSG00000065017 | 10,3683213 | 1,8746316 | 0,501408715 | 4,252866288 | 2,11052E-05 | 0,00040882 | Gm23510 |
| ENSMUSG00000041616 | 1241,81611 | 1,87204061 | 0,054143653 | 34,67611489 | 1,8053E-263 | 1,303E-259 | Nppa |
| ENSMUSG00000068876 | 4,075512016 | 1,84868044 | 0,555030426 | 3,903960712 | 9,46311E-05 | 0,00152243 | Cgn |

| | | | | | | | |
|---|---|---|---|---|---|---|---|
| ENSMUSG00000031613 | 9,563374938 | 1,84707024 | 0,367708823 | 5,396719003 | 6,78706E-08 | 2,4304E-06 | Hpgd |
| ENSMUSG00000016255 | 5,729786908 | 1,83855834 | 0,503677303 | 4,16339041 | 3,13557E-05 | 0,00058613 | Tubb1 |
| ENSMUSG00000058715 | 4,324292696 | 1,83465891 | 0,557655583 | 3,863511923 | 0,000111768 | 0,00175892 | Fcer1g |
| ENSMUSG00000032068 | 1,750515353 | 1,82546179 | 0,861911147 | 2,990443904 | 0,002785723 | 0,0253953 | Plet1 |
| ENSMUSG00000052305 | 284,260898 | 1,81339378 | 0,098384915 | 18,54537884 | 8,88723E-77 | 9,8655E-74 | Hbb-bs |
| ENSMUSG00000048387 | 5,632114072 | 1,81271137 | 0,481653714 | 4,265398439 | 1,99546E-05 | 0,00038914 | Osr1 |
| ENSMUSG00000079293 | 2,147434453 | 1,80558212 | 0,76238039 | 3,141593332 | 0,001680312 | 0,01707234 | Clec7a |
| ENSMUSG00000044258 | 27,63903637 | 1,80031441 | 0,203077677 | 9,070702099 | 1,18253E-19 | 2,2162E-17 | Ctla2a |
| ENSMUSG00000028583 | 11,40415412 | 1,79343479 | 0,274313205 | 6,820261863 | 9,08747E-12 | 6,2448E-10 | Pdpn |
| ENSMUSG00000020467 | 8,715339278 | 1,78163368 | 0,324385826 | 5,833962045 | 5,41264E-09 | 2,4258E-07 | Efemp1 |
| ENSMUSG00000061816 | 41,49802862 | 1,77913311 | 0,164161197 | 11,01120756 | 3,3745E-28 | 1,2815E-25 | Myl1 |
| ENSMUSG00000044468 | 21,12661031 | 1,77865188 | 0,237425247 | 7,747984186 | 9,33628E-15 | 9,9801E-13 | Fam46c |
| ENSMUSG00000047228 | 1,430682699 | 1,77165662 | 1,04899571 | 2,807599779 | 0,004991221 | 0,04014956 | BC048546 |
| ENSMUSG00000025362 | 32,04993157 | 1,76398556 | 0,169725345 | 10,57421912 | 3,92433E-26 | 1,2049E-23 | Rps26 |
| ENSMUSG00000026193 | 177,125616 | 1,75448993 | 0,080566193 | 21,87231812 | 4,7673E-106 | 9,828E-103 | Fn1 |
| ENSMUSG00000028871 | 1,759036526 | 1,74403721 | 0,897380604 | 2,885181331 | 0,003911883 | 0,03316826 | Rspo1 |
| ENSMUSG00000000204 | 11,05395878 | 1,73908287 | 0,334312735 | 5,56400876 | 2,63647E-08 | 1,0452E-06 | Slfn4 |
| ENSMUSG00000045545 | 3,398153585 | 1,72344613 | 0,576851589 | 3,60345808 | 0,000314011 | 0,00424695 | Krt14 |
| ENSMUSG00000024810 | 21,96506239 | 1,72097781 | 0,203271478 | 8,691697045 | 3,57065E-18 | 5,258E-16 | Il33 |
| ENSMUSG00000006014 | 77,43773137 | 1,7165805 | 0,173564366 | 10,08462726 | 6,46099E-24 | 1,6358E-21 | Prg4 |
| ENSMUSG00000039232 | 3,752230321 | 1,70987521 | 0,544692575 | 3,736498442 | 0,000186601 | 0,00275341 | Stx11 |
| ENSMUSG00000006651 | 3,574376104 | 1,70731169 | 0,609793134 | 3,465317306 | 0,000529606 | 0,00662283 | Aplp1 |
| ENSMUSG00000026581 | 6,962678554 | 1,70567371 | 0,400756479 | 4,696440984 | 2,64734E-06 | 6,5736E-05 | Sell |
| ENSMUSG00000073940 | 169,5864615 | 1,69628987 | 0,248083285 | 7,109739427 | 1,16262E-12 | 8,8304E-11 | Hbb-bt |
| ENSMUSG00000040747 | 5,308527066 | 1,69569968 | 0,380513318 | 4,871497376 | 1,10756E-06 | 3,0271E-05 | Cd53 |
| ENSMUSG00000069792 | 2,836542271 | 1,69385605 | 0,67377334 | 3,260495592 | 0,001112177 | 0,01230815 | Wfdc17 |
| ENSMUSG00000032554 | 28,34339752 | 1,68862795 | 0,184946233 | 9,339600453 | 9,66983E-21 | 1,9935E-18 | Trf |
| ENSMUSG00000026347 | 6,761091701 | 1,6854472 | 0,373190159 | 4,926913831 | 8,35386E-07 | 2,3546E-05 | Tmem163 |
| ENSMUSG00000019122 | 16,51217109 | 1,68309137 | 0,223237539 | 7,787474302 | 6,83619E-15 | 7,4737E-13 | Ccl9 |
| ENSMUSG00000029304 | 2,972295013 | 1,67722431 | 0,639635241 | 3,335608053 | 0,000851131 | 0,00987353 | Spp1 |
| ENSMUSG00000078763 | 4,436151526 | 1,67002963 | 0,441072837 | 4,280684825 | 1,86319E-05 | 0,00036682 | Slfn1 |
| ENSMUSG00000022523 | 9,461648421 | 1,66535284 | 0,344278916 | 5,226350946 | 1,72888E-07 | 5,7093E-06 | Fgf12 |
| ENSMUSG00000030046 | 162,9789317 | 1,65776904 | 0,082657089 | 20,12746081 | 4,24181E-90 | 6,1214E-87 | Bmp10 |
| ENSMUSG00000061086 | 537,0582248 | 1,65251069 | 0,091137821 | 18,20375512 | 4,81883E-74 | 4,9672E-71 | Myl4 |
| ENSMUSG00000015947 | 4,053638716 | 1,65104923 | 0,604659253 | 3,412362073 | 0,000644025 | 0,00781659 | Fcgr1 |
| ENSMUSG00000033777 | 3,2606068 | 1,65044446 | 0,533042163 | 3,692571433 | 0,000221998 | 0,00317781 | Tlr13 |
| ENSMUSG00000043263 | 5,555757069 | 1,6463152 | 0,431389921 | 4,294499836 | 1,75088E-05 | 0,00034612 | Pyhin1 |
| ENSMUSG00000054675 | 4,752452399 | 1,63131213 | 0,491488941 | 3,881140005 | 0,000103968 | 0,0016542 | Tmem119 |
| ENSMUSG00000020077 | 19,10700704 | 1,6191407 | 0,209265423 | 7,977016388 | 1,49913E-15 | 1,7401E-13 | Srgn |
| ENSMUSG00000025804 | 6,739222374 | 1,61481997 | 0,345389398 | 5,070967922 | 3,95797E-07 | 1,2205E-05 | Ccr1 |
| ENSMUSG00000049103 | 12,74637067 | 1,61159355 | 0,275321372 | 6,171180373 | 6,7782E-10 | 3,481E-08 | Ccr2 |
| ENSMUSG00000005087 | 37,11019066 | 1,60501614 | 0,15773108 | 10,35565341 | 3,94484E-25 | 1,1162E-22 | Cd44 |
| ENSMUSG00000015340 | 7,00018577 | 1,60091809 | 0,343306055 | 5,06013114 | 4,18968E-07 | 1,2779E-05 | Cybb |
| ENSMUSG00000020120 | 21,63777704 | 1,59631196 | 0,211280371 | 7,788245132 | 6,79463E-15 | 7,4737E-13 | Plek |
| ENSMUSG00000021957 | 41,80574575 | 1,596271 | 0,18564329 | 8,817741747 | 1,16791E-18 | 1,9152E-16 | Tkt |
| ENSMUSG00000017009 | 79,90837048 | 1,59261352 | 0,108067776 | 14,88360364 | 4,21188E-50 | 3,0391E-47 | Sdc4 |
| ENSMUSG00000071068 | 3,677536195 | 1,58006609 | 0,662127767 | 3,151957092 | 0,001621801 | 0,01650891 | Treml2 |
| ENSMUSG00000036098 | 8,866556258 | 1,57024432 | 0,356957572 | 4,829240457 | 1,37055E-06 | 3,6763E-05 | Myrf |
| ENSMUSG00000041959 | 34,52283913 | 1,54684815 | 0,181817023 | 8,724177179 | 2,68123E-18 | 4,0305E-16 | S100a10 |
| ENSMUSG00000017737 | 2,117051183 | 1,54123489 | 0,655471254 | 3,132561534 | 0,001732881 | 0,01743878 | Mmp9 |
| ENSMUSG00000005800 | 3,598369225 | 1,54115659 | 0,524390079 | 3,555095377 | 0,000377842 | 0,00499326 | Mmp8 |
| ENSMUSG00000020644 | 8,080847635 | 1,5345152 | 0,311701642 | 5,296284506 | 1,18183E-07 | 4,0192E-06 | Id2 |
| ENSMUSG00000021215 | 22,92951528 | 1,53355255 | 0,206086976 | 7,693258465 | 1,43434E-14 | 1,4891E-12 | Net1 |
| ENSMUSG00000015134 | 7,206443101 | 1,53252532 | 0,374534259 | 4,535939406 | 5,73476E-06 | 0,00013095 | Aldh1a3 |
| ENSMUSG00000026655 | 6,598438264 | 1,52573655 | 0,36528808 | 4,620989634 | 3,81914E-06 | 9,1248E-05 | Fam107b |
| ENSMUSG00000079012 | 3,091837458 | 1,52149968 | 0,7009236 | 3,012323534 | 0,002592561 | 0,02390179 | Serpina3m |

| | | | | | | | |
|---|---|---|---|---|---|---|---|
| ENSMUSG00000000303 | 1,761990896 | 1,51596157 | 0,812429145 | 2,828924501 | 0,004670472 | 0,03801442 | Cdh1 |
| ENSMUSG00000041653 | 4,515370534 | 1,50724284 | 0,570218467 | 3,336145365 | 0,000849487 | 0,00986822 | Pnpla3 |
| ENSMUSG00000034845 | 20,73428785 | 1,49867875 | 0,192114613 | 8,038105962 | 9,12377E-16 | 1,0704E-13 | Plvap |
| ENSMUSG00000028859 | 8,938665417 | 1,49785988 | 0,409298969 | 4,16661163 | 3,09161E-05 | 0,00057942 | Csf3r |
| ENSMUSG00000028270 | 20,80296876 | 1,49569419 | 0,209977522 | 7,381283514 | 1,56771E-13 | 1,414E-11 | Gbp2 |
| ENSMUSG00000031400 | 8,076223868 | 1,49568209 | 0,3382747 | 4,835193575 | 1,33016E-06 | 3,5813E-05 | G6pdx |
| ENSMUSG00000067818 | 41,76298962 | 1,49244096 | 0,158633853 | 9,604035105 | 7,68746E-22 | 1,7334E-19 | Myl9 |
| ENSMUSG00000004266 | 9,033485928 | 1,49110502 | 0,329861748 | 4,9233092 | 8,50928E-07 | 2,3891E-05 | Ptpn6 |
| ENSMUSG00000038642 | 23,24173667 | 1,48365363 | 0,189427096 | 8,077320342 | 6,62055E-16 | 7,8312E-14 | Ctss |
| ENSMUSG00000069919 | 44,77837254 | 1,48343465 | 0,173226315 | 8,783749363 | 1,58114E-18 | 2,5076E-16 | Hba-a1 |
| ENSMUSG00000069516 | 51,21819076 | 1,47557639 | 0,132731549 | 11,2962459 | 1,36948E-29 | 5,8126E-27 | Lyz2 |
| ENSMUSG00000040152 | 142,671601 | 1,47198595 | 0,115934112 | 12,84403808 | 9,28995E-38 | 5,586E-35 | Thbs1 |
| ENSMUSG00000032271 | 4,53297038 | 1,46652433 | 0,509249173 | 3,512522864 | 0,000443874 | 0,00570904 | Nnmt |
| ENSMUSG00000020654 | 5,514593381 | 1,45459216 | 0,556381906 | 3,327292414 | 0,000876943 | 0,01011604 | Adcy3 |
| ENSMUSG00000046805 | 12,53910034 | 1,45122393 | 0,316429004 | 4,978609669 | 6,40427E-07 | 1,8596E-05 | Mpeg1 |
| ENSMUSG00000030162 | 3,359256839 | 1,45072151 | 0,56422796 | 3,285971337 | 0,001016314 | 0,0114492 | Olr1 |
| ENSMUSG00000054555 | 3,741939934 | 1,45023276 | 0,591185852 | 3,200055177 | 0,001374013 | 0,01453768 | Adam12 |
| ENSMUSG00000023019 | 63,14137518 | 1,44911669 | 0,19006373 | 7,875458491 | 3,39496E-15 | 3,7979E-13 | Gpd1 |
| ENSMUSG00000037922 | 3,924341801 | 1,44420514 | 0,552035702 | 3,308997876 | 0,000936305 | 0,01068973 | Bank1 |
| ENSMUSG00000030849 | 3,901820487 | 1,43562665 | 0,547388012 | 3,318710089 | 0,000904343 | 0,01038231 | Fgfr2 |
| ENSMUSG00000026163 | 3,824295991 | 1,43062967 | 0,476625433 | 3,613536999 | 0,000302048 | 0,00411213 | Sphkap |
| ENSMUSG00000000958 | 3,874005122 | 1,42141075 | 0,63534608 | 3,041205775 | 0,002356327 | 0,0222686 | Slc7a7 |
| ENSMUSG00000065721 | 2,949406564 | 1,42107196 | 0,725603311 | 2,879515902 | 0,003982862 | 0,03367117 | Gm26168 |
| ENSMUSG00000093938 | 3,31879918 | 1,41499524 | 0,569275575 | 3,213036266 | 0,001313397 | 0,01401896 | Evi2b |
| ENSMUSG00000024679 | 3,885369408 | 1,41323496 | 0,57613786 | 3,179811195 | 0,00147371 | 0,01537752 | Ms4a6d |
| ENSMUSG00000065160 | 5,396925525 | 1,41274918 | 0,598929772 | 3,130326629 | 0,00174612 | 0,01751683 | Mir5117 |
| ENSMUSG00000003355 | 1,694213842 | 1,40884191 | 0,857017775 | 2,730502655 | 0,006323782 | 0,04800553 | Fkbp11 |
| ENSMUSG00000031584 | 18,73315924 | 1,40697127 | 0,213065798 | 6,879090247 | 6,0236E-12 | 4,2197E-10 | Gsr |
| ENSMUSG00000031994 | 7,970485698 | 1,4016271 | 0,339506813 | 4,572695046 | 4,81491E-06 | 0,00011225 | Adamts8 |
| ENSMUSG00000001131 | 13,53345326 | 1,40035519 | 0,258382639 | 5,752439566 | 8,79647E-09 | 3,8351E-07 | Timp1 |
| ENSMUSG00000029019 | 128,653974 | 1,39563518 | 0,126956653 | 11,13406852 | 8,56356E-29 | 3,34E-26 | Nppb |
| ENSMUSG00000039236 | 3,637517395 | 1,39363036 | 0,566905067 | 3,200438369 | 0,001372187 | 0,01453768 | Isg20 |
| ENSMUSG00000030830 | 6,082743 | 1,39188363 | 0,432644376 | 3,786801097 | 0,000152599 | 0,00231288 | Itgal |
| ENSMUSG00000024526 | 12,92856069 | 1,3809266 | 0,302704425 | 4,967261427 | 6,7905E-07 | 1,9599E-05 | Cidea |
| ENSMUSG00000000290 | 15,81041461 | 1,37775407 | 0,229540384 | 6,289680838 | 3,18119E-10 | 1,7066E-08 | Itgb2 |
| ENSMUSG00000020186 | 22,08538517 | 1,37612493 | 0,226341216 | 6,380573296 | 1,76426E-10 | 9,8682E-09 | Csrp2 |
| ENSMUSG00000024659 | 22,93126746 | 1,37470629 | 0,189477584 | 7,501798838 | 6,29479E-14 | 5,9373E-12 | Anxa1 |
| ENSMUSG00000029373 | 10,66431056 | 1,37139221 | 0,302732331 | 4,946190232 | 7,568E-07 | 2,1541E-05 | Pf4 |
| ENSMUSG00000029810 | 17,36579551 | 1,37074727 | 0,299013409 | 4,980585173 | 6,33923E-07 | 1,8481E-05 | Tmem176b |
| ENSMUSG00000086324 | 89,47950218 | 1,36942478 | 0,540583036 | 3,257582285 | 0,001123657 | 0,01240665 | Gm15564 |
| ENSMUSG00000032085 | 38,4358139 | 1,35759353 | 0,203736577 | 6,932353144 | 4,13897E-12 | 2,9865E-10 | Tagln |
| ENSMUSG00000028874 | 4,3402443 | 1,35599655 | 0,464535124 | 3,531119235 | 0,000413805 | 0,00537984 | Fgr |
| ENSMUSG00000028019 | 6,533733795 | 1,34653691 | 0,395380296 | 3,936324045 | 8,27392E-05 | 0,00135837 | Pdgfc |
| ENSMUSG00000030772 | 79,69488335 | 1,34209364 | 0,121891686 | 11,18045769 | 5,08272E-29 | 2,0375E-26 | Dkk3 |
| ENSMUSG00000025044 | 13,43616422 | 1,33143801 | 0,285112884 | 5,050185386 | 4,41382E-07 | 1,3353E-05 | Msr1 |
| ENSMUSG00000018774 | 3,043981748 | 1,32958096 | 0,538132994 | 3,199734854 | 0,001375541 | 0,01453768 | Cd68 |
| ENSMUSG00000027078 | 10,69444482 | 1,32931673 | 0,311529737 | 4,692883408 | 2,69381E-06 | 6,6566E-05 | Ube2l6 |
| ENSMUSG00000046721 | 6,953572823 | 1,32802926 | 0,378152772 | 4,027940313 | 5,62676E-05 | 0,00096782 | Rpl14-ps1 |
| ENSMUSG00000084667 | 12,57275972 | 1,32598431 | 0,796701695 | 2,77326872 | 0,005549627 | 0,04340741 | Gm23212 |
| ENSMUSG00000032487 | 6,958798729 | 1,31801782 | 0,3503452 | 4,245203122 | 2,18395E-05 | 0,00042135 | Ptgs2 |
| ENSMUSG00000096768 | 63,77359812 | 1,30732358 | 0,214442911 | 6,398639485 | 1,56767E-10 | 8,8718E-09 | Erdr1 |
| ENSMUSG00000021998 | 27,17852608 | 1,30660787 | 0,196605999 | 6,91748195 | 4,59742E-12 | 3,2844E-10 | Lcp1 |
| ENSMUSG00000031488 | 7,692889135 | 1,30498066 | 0,380436145 | 3,957665046 | 7,5686E-05 | 0,00125832 | Rab11fip1 |
| ENSMUSG00000023046 | 19,00458387 | 1,30451721 | 0,235797302 | 5,862324453 | 4,56432E-09 | 2,0648E-07 | Igfbp6 |
| ENSMUSG00000027195 | 15,4545639 | 1,30253869 | 0,247582863 | 5,607957264 | 2,04729E-08 | 8,299E-07 | Hsd17b12 |
| ENSMUSG00000024349 | 7,395598148 | 1,30163855 | 0,355151845 | 4,152966631 | 3,28193E-05 | 0,00060954 | Tmem173 |

| | | | | | | | |
|---|---|---|---|---|---|---|---|
| ENSMUSG00000021792 | 12,53584842 | 1,29914388 | 0,273231795 | 5,154326337 | 2,54544E-07 | 8,1089E-06 | Fam213a |
| ENSMUSG00000020689 | 14,60116317 | 1,28921535 | 0,298732534 | 4,729346237 | 2,25244E-06 | 5,7026E-05 | Itgb3 |
| ENSMUSG00000038871 | 18,21049461 | 1,28726532 | 0,242224976 | 5,652315583 | 1,58301E-08 | 6,5829E-07 | Bpgm |
| ENSMUSG00000030342 | 23,10992621 | 1,28572943 | 0,196637905 | 6,810952041 | 9,6955E-12 | 6,6311E-10 | Cd9 |
| ENSMUSG00000058258 | 6,150646298 | 1,28512891 | 0,386252215 | 3,875379429 | 0,000106459 | 0,00169197 | Idi1 |
| ENSMUSG00000035373 | 8,32153972 | 1,27397993 | 0,322269751 | 4,412715186 | 1,02082E-05 | 0,00021922 | Ccl7 |
| ENSMUSG00000049001 | 6,408005506 | 1,27343412 | 0,38502801 | 3,845033659 | 0,000120536 | 0,00187643 | Ndnf |
| ENSMUSG00000079227 | 10,577535 | 1,27189817 | 0,309688156 | 4,546294237 | 5,45987E-06 | 0,00012546 | Ccr5 |
| ENSMUSG00000035232 | 3,53079252 | 1,27188204 | 0,549908179 | 3,08057974 | 0,00206598 | 0,02006337 | Pdk3 |
| ENSMUSG00000106471 | 4,505603885 | 1,27009145 | 0,739368609 | 2,76526647 | 0,005687631 | 0,04427088 | mmu-mir-486b |
| ENSMUSG00000018339 | 283,3483263 | 1,27003012 | 0,084009429 | 15,24243844 | 1,84841E-52 | 1,4039E-49 | Gpx3 |
| ENSMUSG00000026479 | 11,23857691 | 1,26868257 | 0,325002364 | 4,361033008 | 1,2945E-05 | 0,00026802 | Lamc2 |
| ENSMUSG00000027962 | 34,94169789 | 1,26330342 | 0,197149763 | 6,686034635 | 2,29299E-11 | 1,511E-09 | Vcam1 |
| ENSMUSG00000059173 | 12,15588582 | 1,26236489 | 0,293119268 | 4,720074381 | 2,35758E-06 | 5,948E-05 | Pde1a |
| ENSMUSG00000053279 | 26,75523047 | 1,26235865 | 0,198928012 | 6,615818841 | 3,69501E-11 | 2,3805E-09 | Aldh1a1 |
| ENSMUSG00000020917 | 49,30751794 | 1,26139078 | 0,137163918 | 9,343292546 | 9,3384E-21 | 1,9531E-18 | Acly |
| ENSMUSG00000021614 | 95,44729287 | 1,25555534 | 0,101184008 | 12,55077432 | 3,93642E-36 | 2,2723E-33 | Vcan |
| ENSMUSG00000046275 | 4,244272882 | 1,25166015 | 0,557069054 | 3,03203672 | 0,002429097 | 0,02279213 | Tusc5 |
| ENSMUSG00000098112 | 7,734807957 | 1,25095066 | 0,397761391 | 3,710916052 | 0,000206511 | 0,00299513 | Bin2 |
| ENSMUSG00000079435 | 47,83995992 | 1,24787501 | 0,148310939 | 8,635551672 | 5,84443E-18 | 8,2687E-16 | Rpl36a |
| ENSMUSG00000026827 | 13,03903634 | 1,24712669 | 0,307177434 | 4,503638676 | 6,67997E-06 | 0,00014946 | Gpd2 |
| ENSMUSG00000041559 | 6,859665254 | 1,24610189 | 0,342694583 | 4,120539129 | 3,77987E-05 | 0,00068699 | Fmod |
| ENSMUSG00000059854 | 1,88367974 | 1,24560884 | 0,68909651 | 2,785151564 | 0,005350271 | 0,04223729 | Hydin |
| ENSMUSG00000013584 | 19,8354973 | 1,2402932 | 0,229026588 | 5,753214534 | 8,75622E-09 | 3,8291E-07 | Aldh1a2 |
| ENSMUSG00000040998 | 5,529558216 | 1,24012415 | 0,472004145 | 3,302188847 | 0,000959335 | 0,0109095 | Npnt |
| ENSMUSG00000023832 | 2,841061082 | 1,2359595 | 0,621547333 | 2,877319329 | 0,004010695 | 0,0338272 | Acat2 |
| ENSMUSG00000023913 | 7,660428185 | 1,23262078 | 0,337255147 | 4,148208659 | 3,35087E-05 | 0,00062155 | Pla2g7 |
| ENSMUSG00000030688 | 46,79984652 | 1,22996508 | 0,141724653 | 8,887366852 | 6,25741E-19 | 1,1012E-16 | Stard10 |
| ENSMUSG00000025491 | 3,229737933 | 1,2275058 | 0,531354609 | 3,071566874 | 0,002129385 | 0,02059594 | Ifitm1 |
| ENSMUSG00000064432 | 16,36035055 | 1,22115121 | 0,690245392 | 2,789777916 | 0,005274421 | 0,04172981 | Gm25587 |
| ENSMUSG00000022636 | 8,104780878 | 1,21946783 | 0,321602986 | 4,257521408 | 2,06706E-05 | 0,00040094 | Alcam |
| ENSMUSG00000025321 | 7,309616496 | 1,21644718 | 0,334747153 | 4,123354348 | 3,73395E-05 | 0,00068208 | Itgb8 |
| ENSMUSG00000065591 | 6,785837062 | 1,21609736 | 0,508507982 | 3,139342461 | 0,001693274 | 0,01716134 | Gm22753 |
| ENSMUSG00000079017 | 4,476627136 | 1,21454777 | 0,432578845 | 3,431714673 | 0,000599778 | 0,00734131 | Ifi27l2a |
| ENSMUSG00000058427 | 3,769741405 | 1,21311643 | 0,452526479 | 3,33972472 | 0,000838615 | 0,00975972 | Cxcl2 |
| ENSMUSG00000026070 | 2,922813609 | 1,21279582 | 0,552590857 | 3,00574711 | 0,002649292 | 0,02433605 | Il18r1 |
| ENSMUSG00000026395 | 22,98766227 | 1,21229848 | 0,202162678 | 6,296824973 | 3,03804E-10 | 1,6359E-08 | Ptprc |
| ENSMUSG00000004730 | 5,763150612 | 1,21192969 | 0,392819172 | 3,656311574 | 0,00025587 | 0,0035676 | Adgre1 |
| ENSMUSG00000035783 | 83,85725805 | 1,20876967 | 0,143189353 | 8,673236555 | 4,20008E-18 | 6,1224E-16 | Acta2 |
| ENSMUSG00000035493 | 42,71630751 | 1,20691288 | 0,152105581 | 8,148919995 | 3,67189E-16 | 4,568E-14 | Tgfbi |
| ENSMUSG00000056220 | 8,587538725 | 1,20548828 | 0,355658122 | 3,912301291 | 9,14208E-05 | 0,00147903 | Pla2g4a |
| ENSMUSG00000030088 | 11,96012943 | 1,2051065 | 0,322815173 | 4,207663987 | 2,58024E-05 | 0,0004893 | Aldh1l1 |
| ENSMUSG00000002111 | 6,636005111 | 1,20190724 | 0,465151567 | 3,273352698 | 0,001062798 | 0,01187092 | Spi1 |
| ENSMUSG00000001025 | 18,26613808 | 1,19791255 | 0,238542598 | 5,373470325 | 7,72355E-08 | 2,7251E-06 | S100a6 |
| ENSMUSG00000032231 | 55,87239209 | 1,19786946 | 0,13694434 | 8,945207592 | 3,71251E-19 | 6,6142E-17 | Anxa2 |
| ENSMUSG00000017969 | 23,65537387 | 1,1974974 | 0,238037755 | 5,378581053 | 7,50752E-08 | 2,6554E-06 | Ptgis |
| ENSMUSG00000031246 | 30,33384325 | 1,19487579 | 0,190705152 | 6,531702728 | 6,50261E-11 | 4,0102E-09 | Sh3bgrl |
| ENSMUSG00000000594 | 9,832475864 | 1,19274099 | 0,326480173 | 4,138922687 | 3,48941E-05 | 0,00064393 | Gm2a |
| ENSMUSG00000046733 | 2,302038693 | 1,19192113 | 0,708508855 | 2,721557639 | 0,006497505 | 0,0490149 | Gprc5a |
| ENSMUSG00000019814 | 14,99800352 | 1,18938533 | 0,255451723 | 5,036127223 | 4,75045E-07 | 1,4223E-05 | Ltv1 |
| ENSMUSG00000065289 | 27,44229996 | 1,18650032 | 0,312984853 | 4,260606652 | 2,03873E-05 | 0,00039704 | Gm23650 |
| ENSMUSG00000054966 | 6,23203552 | 1,18579285 | 0,37239312 | 3,747464028 | 0,000178631 | 0,00265209 | Lmntd1 |
| ENSMUSG00000020473 | 50,44285533 | 1,18438756 | 0,14703809 | 8,300809242 | 1,03405E-16 | 1,369E-14 | Aebp1 |
| ENSMUSG00000026335 | 890,7236321 | 1,18356177 | 0,066567037 | 17,89435533 | 1,30498E-71 | 1,2555E-68 | Pam |
| ENSMUSG00000009376 | 6,782498983 | 1,1812357 | 0,400019748 | 3,543742272 | 0,000394491 | 0,00518007 | Met |
| ENSMUSG00000039405 | 10,870178 | 1,17695972 | 0,332162716 | 4,033157671 | 5,50323E-05 | 0,00095339 | Prss23 |

| | | | | | | | |
|---|---|---|---|---|---|---|---|
| ENSMUSG00000036446 | 12,87394273 | 1,17099679 | 0,331502674 | 4,029145509 | 5,598E-05 | 0,00096517 | Lum |
| ENSMUSG00000024892 | 30,35245391 | 1,16778362 | 0,223850705 | 5,553484515 | 2,8003E-08 | 1,0952E-06 | Pcx |
| ENSMUSG00000026068 | 3,050873269 | 1,16760699 | 0,515809746 | 3,027693941 | 0,002464275 | 0,02297284 | Il18rap |
| ENSMUSG00000058624 | 60,96375993 | 1,1671348 | 0,138475975 | 8,642695097 | 5,49023E-18 | 7,8445E-16 | Gda |
| ENSMUSG00000009687 | 18,66157948 | 1,16711052 | 0,24633534 | 5,112404 | 3,18085E-07 | 9,9789E-06 | Fxyd5 |
| ENSMUSG00000008540 | 20,50227226 | 1,16659804 | 0,223579497 | 5,55815256 | 2,72645E-08 | 1,0721E-06 | Mgst1 |
| ENSMUSG00000041362 | 7,186390989 | 1,16522875 | 0,466934622 | 3,199620413 | 0,001376087 | 0,01453768 | Shtn1 |
| ENSMUSG00000030671 | 11,31129276 | 1,16430768 | 0,315545641 | 4,167417391 | 3,0807E-05 | 0,00057812 | Pde3b |
| ENSMUSG00000040254 | 6,218302275 | 1,16252756 | 0,464568881 | 3,198709175 | 0,001380443 | 0,01456448 | Sema3d |
| ENSMUSG00000037742 | 513,0553403 | 1,16095965 | 0,05797776 | 20,10163007 | 7,14091E-90 | 9,3682E-87 | Eef1a1 |
| ENSMUSG00000086806 | 21,87357035 | 1,16016375 | 0,299928558 | 4,311706065 | 1,62E-05 | 0,0003256 | Gm13054 |
| ENSMUSG00000030220 | 8,425596498 | 1,15792519 | 0,367216719 | 3,700834415 | 0,000214892 | 0,00309183 | Arhgdib |
| ENSMUSG00000026768 | 15,75025688 | 1,15769157 | 0,340880884 | 3,913187447 | 9,10858E-05 | 0,00147526 | Itga8 |
| ENSMUSG00000052749 | 6,156644279 | 1,15583721 | 0,362555229 | 3,739581862 | 0,000184327 | 0,00272943 | Trim30b |
| ENSMUSG00000060802 | 61,19088353 | 1,15168519 | 0,133726083 | 8,783739744 | 1,58127E-18 | 2,5076E-16 | B2m |
| ENSMUSG00000018830 | 220,1757078 | 1,14745831 | 0,11500821 | 10,13600301 | 3,82428E-24 | 1,0034E-21 | Myh11 |
| ENSMUSG00000026365 | 363,6825892 | 1,14056566 | 0,129833823 | 9,01995556 | 1,88166E-19 | 3,4372E-17 | Cfh |
| ENSMUSG00000040612 | 6,247595637 | 1,14037863 | 0,374524081 | 3,615921527 | 0,000299281 | 0,0040783 | Ildr2 |
| ENSMUSG00000020922 | 17,90214262 | 1,13899734 | 0,21957707 | 5,527644045 | 3,2456E-08 | 1,2358E-06 | Lsm12 |
| ENSMUSG00000058546 | 110,8371003 | 1,13145159 | 0,111650414 | 10,30694478 | 6,55522E-25 | 1,8192E-22 | Rpl23a |
| ENSMUSG00000078784 | 7,865451833 | 1,12939725 | 0,362216366 | 3,674899167 | 0,000237943 | 0,00336768 | 1810022K09Rik |
| ENSMUSG00000023367 | 5,772838526 | 1,12893375 | 0,406015918 | 3,40312613 | 0,000666195 | 0,00803163 | Tmem176a |
| ENSMUSG00000060703 | 5,436103697 | 1,12048724 | 0,389190958 | 3,485773213 | 0,000490717 | 0,00622279 | Cd302 |
| ENSMUSG00000020018 | 10,77997178 | 1,11924711 | 0,293346954 | 4,268510546 | 1,96783E-05 | 0,00038479 | Snrpf |
| ENSMUSG00000039497 | 12,15244593 | 1,11852062 | 0,252263049 | 4,831133755 | 1,35758E-06 | 3,6483E-05 | Dse |
| ENSMUSG00000027533 | 39,27992537 | 1,11832876 | 0,157237842 | 7,355454219 | 1,90279E-13 | 1,6642E-11 | Fabp5 |
| ENSMUSG00000065272 | 23,23785163 | 1,11617954 | 0,284212655 | 4,370478835 | 1,23974E-05 | 0,00025705 | Snord57 |
| ENSMUSG00000080463 | 27,49171493 | 1,11315197 | 0,468555356 | 3,11530727 | 0,001837534 | 0,01826271 | Gm22489 |
| ENSMUSG00000064945 | 242,7667032 | 1,11093942 | 0,507459248 | 2,985772101 | 0,002828634 | 0,0257053 | Rny3 |
| ENSMUSG00000028656 | 48,7389992 | 1,10277807 | 0,138629125 | 8,198447502 | 2,43511E-16 | 3,0558E-14 | Cap1 |
| ENSMUSG00000026728 | 371,1687967 | 1,10269323 | 0,057721441 | 19,18459667 | 4,97788E-82 | 5,9863E-79 | Vim |
| ENSMUSG00000073489 | 24,83426103 | 1,10061036 | 0,215756613 | 5,43876048 | 5,36525E-08 | 1,9548E-06 | Ifi204 |
| ENSMUSG00000021136 | 11,59357892 | 1,09715759 | 0,284771232 | 4,299721759 | 1,71013E-05 | 0,00034047 | Smoc1 |
| ENSMUSG00000023039 | 3,782661654 | 1,0964395 | 0,501412719 | 2,970271148 | 0,00297537 | 0,02670247 | Krt7 |
| ENSMUSG00000028369 | 33,26406672 | 1,08650524 | 0,216954122 | 5,356539376 | 8,4831E-08 | 2,9714E-06 | Svep1 |
| ENSMUSG00000028015 | 8,740742289 | 1,0853389 | 0,325738265 | 3,850623819 | 0,000117817 | 0,00183808 | Ctso |
| ENSMUSG00000030587 | 3,981728029 | 1,08441379 | 0,53835417 | 2,863633685 | 0,00418812 | 0,03501667 | 2200002D01Rik |
| ENSMUSG00000030786 | 38,76329038 | 1,08362549 | 0,155264232 | 7,258953563 | 3,90096E-13 | 3,1805E-11 | Itgam |
| ENSMUSG00000025492 | 30,12730163 | 1,08088136 | 0,172554956 | 6,539057357 | 6,19078E-11 | 3,8508E-09 | Ifitm3 |
| ENSMUSG00000079419 | 5,917412332 | 1,07967721 | 0,489931238 | 2,989039178 | 0,002798562 | 0,02548016 | Ms4a6c |
| ENSMUSG00000031673 | 9,557114836 | 1,07708327 | 0,272786721 | 4,38342404 | 1,16828E-05 | 0,00024541 | Cdh11 |
| ENSMUSG00000049313 | 25,79768835 | 1,07554328 | 0,18859694 | 6,000654315 | 1,96524E-09 | 9,5489E-08 | Sorl1 |
| ENSMUSG00000032116 | 31,13271691 | 1,07544121 | 0,185033497 | 6,107094069 | 1,01461E-09 | 5,0489E-08 | Stt3a |
| ENSMUSG00000028108 | 29,33561641 | 1,0737964 | 0,222211304 | 5,189568328 | 2,10782E-07 | 6,7746E-06 | Ecm1 |
| ENSMUSG00000052397 | 83,53547722 | 1,07354386 | 0,111787417 | 9,782236045 | 1,34212E-22 | 3,228E-20 | Ezr |
| ENSMUSG00000022371 | 19,65797419 | 1,07274173 | 0,216989865 | 5,289687316 | 1,22526E-07 | 4,1409E-06 | Col14a1 |
| ENSMUSG00000032186 | 9,473692363 | 1,07179757 | 0,328993309 | 3,768466963 | 0,000164253 | 0,00247285 | Tmod2 |
| ENSMUSG00000031762 | 92,56978177 | 1,07144533 | 0,111041968 | 9,853525132 | 6,61812E-23 | 1,6467E-20 | Mt2 |
| ENSMUSG00000072620 | 14,14787127 | 1,06470172 | 0,253558651 | 4,611481956 | 3,99808E-06 | 9,4895E-05 | Slfn2 |
| ENSMUSG00000085272 | 30,07617693 | 1,06239408 | 0,201614577 | 5,593479046 | 2,22564E-08 | 8,9466E-07 | Sbk3 |
| ENSMUSG00000057113 | 104,1610709 | 1,06195997 | 0,116918594 | 9,25453934 | 2,15156E-20 | 4,225E-18 | Npm1 |
| ENSMUSG00000034463 | 4,909071622 | 1,06112112 | 0,438507863 | 3,122158539 | 0,001795302 | 0,01794105 | Scara3 |
| ENSMUSG00000024912 | 2,995247771 | 1,06101783 | 0,617816062 | 2,723241811 | 0,006464472 | 0,0488935 | Fosl1 |
| ENSMUSG00000038633 | 16,87632992 | 1,0585197 | 0,256208151 | 4,552585453 | 5,29906E-06 | 0,0001221 | Degs1 |
| ENSMUSG00000039001 | 45,37877921 | 1,05823224 | 0,140389271 | 7,761600918 | 8,38639E-15 | 9,0316E-13 | Rps21 |
| ENSMUSG00000031827 | 8,60461989 | 1,05770073 | 0,337446405 | 3,678893385 | 0,000234248 | 0,00332542 | Cotl1 |

| | | | | | | | |
|---|---|---|---|---|---|---|---|
| ENSMUSG00000055172 | 19,11521343 | 1,05636158 | 0,204677337 | 5,493150519 | 3,94826E-08 | 1,4799E-06 | C1ra |
| ENSMUSG00000006360 | 15,08136703 | 1,05115357 | 0,270982909 | 4,318604278 | 1,57019E-05 | 0,00031709 | Crip1 |
| ENSMUSG00000028936 | 59,42800461 | 1,05099688 | 0,137313543 | 7,88400614 | 3,17049E-15 | 3,5745E-13 | Rpl22 |
| ENSMUSG00000022351 | 3,126630189 | 1,05090568 | 0,55386641 | 2,796097186 | 0,005172384 | 0,04112544 | Sqle |
| ENSMUSG00000036438 | 68,59607496 | 1,04854108 | 0,113211536 | 9,436529923 | 3,85329E-21 | 8,1775E-19 | Calm2 |
| ENSMUSG00000032548 | 19,0091958 | 1,0473944 | 0,215969644 | 5,196768148 | 2,02783E-07 | 6,5761E-06 | Slco2a1 |
| ENSMUSG00000071866 | 111,6231052 | 1,04731047 | 0,101763271 | 10,44769534 | 1,50129E-25 | 4,4215E-23 | Ppia |
| ENSMUSG00000065228 | 13,52813435 | 1,04712379 | 0,411380067 | 3,222639555 | 0,001270153 | 0,01369404 | Gm25789 |
| ENSMUSG00000052681 | 35,4134466 | 1,04026968 | 0,161412484 | 6,699769542 | 2,08748E-11 | 1,3882E-09 | Rap1b |
| ENSMUSG00000030208 | 35,62702457 | 1,04014195 | 0,175758792 | 6,193917875 | 5,86868E-10 | 3,0407E-08 | Emp1 |
| ENSMUSG00000027009 | 8,266629693 | 1,03713311 | 0,331053361 | 3,668454245 | 0,000244021 | 0,00344567 | Itga4 |
| ENSMUSG00000059498 | 8,06323928 | 1,03320788 | 0,363275576 | 3,439520795 | 0,000582745 | 0,00717542 | Fcgr3 |
| ENSMUSG00000028343 | 17,11399278 | 1,03241213 | 0,218934696 | 5,073020602 | 3,9155E-07 | 1,2099E-05 | Erp44 |
| ENSMUSG00000029561 | 18,22800495 | 1,03077233 | 0,258879132 | 4,399439943 | 1,08531E-05 | 0,00023032 | Oasl2 |
| ENSMUSG00000027508 | 8,178801988 | 1,03011082 | 0,337363822 | 3,609689822 | 0,000306563 | 0,00416967 | Pag1 |
| ENSMUSG00000022283 | 160,2735082 | 1,02432309 | 0,084350379 | 12,26553789 | 1,38688E-34 | 7,6977E-32 | Pabpc1 |
| ENSMUSG00000012428 | 44,68959905 | 1,02304358 | 0,145485688 | 7,273294048 | 3,50826E-13 | 2,9096E-11 | Steap4 |
| ENSMUSG00000033099 | 14,22002504 | 1,02076229 | 0,256630604 | 4,399492243 | 1,08504E-05 | 0,00023032 | Nol12 |
| ENSMUSG00000013160 | 16,81993061 | 1,01888184 | 0,258381038 | 4,373081091 | 1,22505E-05 | 0,00025527 | Atp6v0d1 |
| ENSMUSG00000005054 | 5,394145122 | 1,01871702 | 0,398192357 | 3,222320196 | 0,001271569 | 0,01369404 | Cstb |
| ENSMUSG00000030844 | 4,495877553 | 1,01777741 | 0,489496592 | 2,894871055 | 0,003793143 | 0,0323707 | Rgs10 |
| ENSMUSG00000079641 | 22,02341195 | 1,01753574 | 0,20095765 | 5,398716336 | 6,71194E-08 | 2,4095E-06 | Rpl39 |
| ENSMUSG00000016256 | 17,54503514 | 1,01597505 | 0,238603851 | 4,65727983 | 3,20415E-06 | 7,8107E-05 | Ctsz |
| ENSMUSG00000064356 | 404,6097395 | 1,01518496 | 0,350033441 | 3,486262286 | 0,00048982 | 0,00621729 | mt-Atp8 |
| ENSMUSG00000049744 | 3,94503207 | 1,01350815 | 0,527491493 | 2,801579951 | 0,005085304 | 0,04067961 | Arhgap15 |
| ENSMUSG00000037072 | 28,03023405 | 1,01340254 | 0,193418775 | 5,55838721 | 2,72279E-08 | 1,0721E-06 | Sep15 |
| ENSMUSG00000054641 | 8,677427976 | 1,01331157 | 0,311491634 | 3,771500958 | 0,000162269 | 0,00244691 | Mmrn1 |
| ENSMUSG00000019948 | 3,977702292 | 1,01127159 | 0,515758249 | 2,822093389 | 0,004771128 | 0,03865926 | Actr6 |
| ENSMUSG00000067235 | 4,474638729 | 1,01040241 | 0,465607759 | 2,945458674 | 0,003224762 | 0,02849758 | H2-Q10 |
| ENSMUSG00000060063 | 6,06035577 | 1,00994677 | 0,406058123 | 3,168229194 | 0,001533706 | 0,01590008 | Alox5ap |
| ENSMUSG00000029174 | 57,45011382 | 1,00467701 | 0,12348295 | 8,364684698 | 6,02759E-17 | 8,1294E-15 | Tbc1d1 |
| ENSMUSG00000005973 | 15,52691604 | 1,00278749 | 0,226952682 | 4,790099244 | 1,66699E-06 | 4,3659E-05 | Rcn1 |
| ENSMUSG00000098343 | 27,15683121 | 1,00068962 | 0,463204828 | 2,951339879 | 0,003163985 | 0,0280636 | Mir6240 |
| **Down regulated genes (72)** | | | | | | | |
| ENSMUSG00000106986 | 3,095684012 | -4,496178299 | 1,272552861 | -4,06639882 | 4,77452E-05 | 0,000842863 | RP24-351E22.1 |
| ENSMUSG00000022899 | 7,246742896 | -4,187572834 | 1,225608418 | -3,966039321 | 7,30768E-05 | 0,001220569 | Slc15a2 |
| ENSMUSG00000082277 | 1,784903271 | -4,089346408 | 1,368583529 | -3,394398882 | 0,000687794 | 0,008252194 | Gm15807 |
| ENSMUSG00000075015 | 224,3024598 | -3,889794014 | 0,350195139 | -11,29947558 | 1,32004E-29 | 5,77257E-27 | Gm10801 |
| ENSMUSG00000075249 | 1,573186903 | -3,628284491 | 1,44376094 | -3,259233816 | 0,001117136 | 0,012353551 | Fsip2 |
| ENSMUSG00000099400 | 1,479452894 | -3,454757001 | 1,454772361 | -3,054282479 | 0,002255995 | 0,021517694 | Kat6b-ps1 |
| ENSMUSG00000100254 | 3,310836415 | -3,434032161 | 1,339330903 | -3,159799247 | 0,001578779 | 0,016229737 | Trpc2 |
| ENSMUSG00000096519 | 1,418111752 | -3,369618976 | 1,537190283 | -2,960873324 | 0,003067681 | 0,027383796 | Gm10721 |
| ENSMUSG00000095280 | 42,34764648 | -3,119768686 | 0,258801317 | -12,22156604 | 2,38447E-34 | 1,27445E-31 | Gm21738 |
| ENSMUSG00000075014 | 192,3854893 | -3,028157969 | 0,141132601 | -21,55289923 | 4,9733E-103 | 8,9713E-100 | Gm10800 |
| ENSMUSG00000097312 | 378,5215611 | -2,988699616 | 0,421771639 | -7,384254516 | 1,5331E-13 | 1,39146E-11 | Gm26870 |
| ENSMUSG00000082876 | 1,962368596 | -2,903292926 | 1,175958037 | -3,159272678 | 0,001581634 | 0,016245242 | Gm11889 |
| ENSMUSG00000105416 | 2,03997496 | -2,847562524 | 1,383928808 | -2,937123141 | 0,003312725 | 0,029147159 | RP24-560A18.3 |
| ENSMUSG00000051985 | 3,234257482 | -2,776017969 | 1,257570634 | -3,031343218 | 0,002434683 | 0,022829704 | Igfn1 |
| ENSMUSG00000104211 | 4,782008324 | -2,653532872 | 0,601127347 | -4,83985438 | 1,29934E-06 | 3,51139E-05 | Gm37985 |
| ENSMUSG00000074623 | 2,380437074 | -2,349426857 | 1,179927792 | -2,794664429 | 0,005195361 | 0,041285385 | Gm826 |
| ENSMUSG00000095547 | 11,90461058 | -2,307676551 | 0,382501061 | -6,349066567 | 2,16625E-10 | 1,19318E-08 | Gm10719 |
| ENSMUSG00000079455 | 3,106188711 | -2,214720733 | 0,794470836 | -3,419879057 | 0,00062649 | 0,007616575 | Gm16026 |
| ENSMUSG00000039328 | 6,841125728 | -2,063941935 | 0,606329431 | -3,934011556 | 8,35398E-05 | 0,001369957 | Rnf122 |
| ENSMUSG00000089940 | 6,049083508 | -1,824443797 | 0,528404478 | -3,982993935 | 6,80525E-05 | 0,001148615 | Gm4117 |
| ENSMUSG00000046958 | 5,988228091 | -1,77844999 | 0,45633085 | -4,371390441 | 1,23458E-05 | 0,000256348 | 4930432E11Rik |
| ENSMUSG00000017417 | 3,431441178 | -1,726342931 | 0,770272597 | -3,004066028 | 0,002663975 | 0,024439812 | Plxdc1 |

| | | | | | | | |
|---|---|---|---|---|---|---|---|
| ENSMUSG00000095186 | 4,614046732 | -1,679315638 | 0,596890559 | -3,448425468 | 0,000563865 | 0,006978676 | Gm10718 |
| ENSMUSG00000002274 | 3,98965478 | -1,662212567 | 0,746705153 | -3,001337758 | 0,002687962 | 0,024591428 | Metrn |
| ENSMUSG00000063415 | 9,030058187 | -1,627370595 | 0,459561968 | -4,053129154 | 5,05371E-05 | 0,000888308 | Cyp26b1 |
| ENSMUSG00000065845 | 22,25192062 | -1,615605243 | 0,712928647 | -3,081540481 | 0,002059325 | 0,020025684 | Gm25189 |
| ENSMUSG00000004996 | 5,718612244 | -1,587057316 | 0,582667494 | -3,373657692 | 0,000741765 | 0,008774113 | Mri1 |
| ENSMUSG00000064941 | 15,02204267 | -1,556052185 | 0,495907458 | -3,714026477 | 0,000203987 | 0,002964494 | Gm23238 |
| ENSMUSG00000032968 | 3,395413276 | -1,547994477 | 0,820474293 | -2,761097186 | 0,005760753 | 0,044747733 | Inha |
| ENSMUSG00000024330 | 9,186584808 | -1,53865032 | 0,510704099 | -3,595472054 | 0,000323804 | 0,004354906 | Col11a2 |
| ENSMUSG00000087177 | 4,928795401 | -1,505991962 | 0,588376128 | -3,230768459 | 0,001234579 | 0,013385583 | E130307A14Rik |
| ENSMUSG00000074224 | 3,972666955 | -1,496873109 | 0,575471996 | -3,286685656 | 0,001013739 | 0,011429118 | 4932431P20Rik |
| ENSMUSG00000103238 | 3,240911943 | -1,465016153 | 0,800201944 | -2,762411569 | 0,00573761 | 0,044635824 | Gm36527 |
| ENSMUSG00000035283 | 11,42674611 | -1,393624481 | 0,404993824 | -3,968372383 | 7,23652E-05 | 0,001210084 | Adrb1 |
| ENSMUSG00000106220 | 7,718174857 | -1,391643627 | 0,443205547 | -3,695651302 | 0,000219324 | 0,003146185 | RP24-488I7.1 |
| ENSMUSG00000052794 | 8,263075449 | -1,329365493 | 0,478163829 | -3,402797128 | 0,000666998 | 0,008034596 | 1700030K09Rik |
| ENSMUSG00000082286 | 14,43793054 | -1,293764382 | 0,391630163 | -3,834810123 | 0,000125661 | 0,001943643 | Pisd-ps1 |
| ENSMUSG00000055994 | 4,51860894 | -1,292423591 | 0,656602009 | -2,804532386 | 0,005038963 | 0,040443419 | Nod2 |
| ENSMUSG00000096211 | 10,36954759 | -1,275902312 | 0,521723858 | -3,140402736 | 0,001687157 | 0,017121917 | Gm23330 |
| ENSMUSG00000040441 | 8,844616458 | -1,273360409 | 0,47300953 | -3,327787616 | 0,000875386 | 0,010106153 | Slc26a10 |
| ENSMUSG00000070802 | 5,713640871 | -1,267674874 | 0,569006242 | -2,982625288 | 0,002857877 | 0,025865001 | Pnmal2 |
| ENSMUSG00000095891 | 5,044765953 | -1,244544085 | 0,567738443 | -2,97059645 | 0,002972221 | 0,0266908 | Gm10717 |
| ENSMUSG00000045350 | 5,109652198 | -1,221682597 | 0,613450936 | -2,825480542 | 0,004720976 | 0,038317434 | Fam186a |
| ENSMUSG00000042099 | 106,7755366 | -1,195702711 | 0,162972104 | -7,565301632 | 3,86966E-14 | 3,77318E-12 | Kank3 |
| ENSMUSG00000002910 | 21,10493489 | -1,187688777 | 0,276631467 | -4,70350124 | 2,55738E-06 | 6,38503E-05 | Arrdc2 |
| ENSMUSG00000104525 | 7,459185737 | -1,182264273 | 0,444339183 | -3,302867217 | 0,000957017 | 0,010891728 | Gm37334 |
| ENSMUSG00000031805 | 39,13036366 | -1,171845439 | 0,220985071 | -5,633869463 | 1,7621E-08 | 7,24471E-07 | Jak3 |
| ENSMUSG00000022546 | 11,42860763 | -1,171681463 | 0,358327519 | -3,796393615 | 0,000146816 | 0,002234922 | Gpt |
| ENSMUSG00000006154 | 11,95616779 | -1,168539457 | 0,385320294 | -3,60492399 | 0,000312244 | 0,004227015 | Eps8l1 |
| ENSMUSG00000081824 | 11,30100734 | -1,149324906 | 0,388251989 | -3,523258207 | 0,000426276 | 0,005527032 | BC002163 |
| ENSMUSG00000024210 | 4,863615246 | -1,142792655 | 0,573084944 | -2,840559513 | 0,004503447 | 0,036988755 | Ip6k3 |
| ENSMUSG00000064702 | 436,9746472 | -1,141761951 | 0,288185097 | -4,403978793 | 1,06283E-05 | 0,00022689 | Gm24950 |
| ENSMUSG00000040146 | 6,814903157 | -1,13033179 | 0,465811199 | -3,121763848 | 0,001797711 | 0,017941054 | Rgl3 |
| ENSMUSG00000072770 | 10,22329823 | -1,128432677 | 0,438094779 | -3,222502841 | 0,001270759 | 0,013694044 | Acrbp |
| ENSMUSG00000095915 | 20,3894278 | -1,119059944 | 0,452666279 | -3,157219805 | 0,001592812 | 0,016312875 | n-R5s115 |
| ENSMUSG00000025358 | 5,950476929 | -1,107059923 | 0,535000258 | -2,860590008 | 0,004228535 | 0,035252449 | Cdk2 |
| ENSMUSG00000034173 | 18,95137885 | -1,104950042 | 0,351781686 | -3,661381292 | 0,000250859 | 0,003511303 | Zbed5 |
| ENSMUSG00000039481 | 6,63697378 | -1,097088133 | 0,531714846 | -2,864632292 | 0,004174937 | 0,034946937 | Nrtn |
| ENSMUSG00000044037 | 36,21074734 | -1,091828648 | 0,220081369 | -5,296000849 | 1,18366E-07 | 4,01916E-06 | Als2cl |
| ENSMUSG00000068205 | 7,598990074 | -1,075973914 | 0,492249006 | -2,94682226 | 0,003210577 | 0,028389605 | Macrod2 |
| ENSMUSG00000038188 | 26,80196938 | -1,069168998 | 0,24865952 | -4,705074803 | 2,53773E-06 | 6,34696E-05 | Scarf1 |
| ENSMUSG00000034255 | 29,33971215 | -1,068040005 | 0,25737838 | -4,559397744 | 5,13005E-06 | 0,000118732 | Arhgap27 |
| ENSMUSG00000022053 | 10,10575806 | -1,049781074 | 0,361204352 | -3,485193241 | 0,000491782 | 0,006230819 | Ebf2 |
| ENSMUSG00000021495 | 46,67640426 | -1,045735733 | 0,224912262 | -5,005370664 | 5,57546E-07 | 1,63869E-05 | Fam193b |
| ENSMUSG00000081583 | 17,78529511 | -1,045723838 | 0,343471906 | -3,600038074 | 0,000318171 | 0,00429114 | Gm14769 |
| ENSMUSG00000040003 | 32,45235095 | -1,044159384 | 0,206700102 | -5,360564555 | 8,29623E-08 | 2,91296E-06 | Magi2 |
| ENSMUSG00000074652 | 328,8100844 | -1,033039627 | 0,097711228 | -10,73967045 | 6,62806E-27 | 2,12554E-24 | Myh7b |
| ENSMUSG00000001943 | 7,563140035 | -1,032460741 | 0,528691623 | -2,782907691 | 0,005387413 | 0,042437642 | Vsig2 |
| ENSMUSG00000015090 | 99,73799482 | -1,029600656 | 0,128739209 | -8,210550926 | 2,20176E-16 | 2,78716E-14 | Ptgds |
| ENSMUSG00000026017 | 13,73039546 | -1,018986024 | 0,312699439 | -3,766035041 | 0,000165861 | 0,002482918 | Carf |
| ENSMUSG00000076433 | 8,903336017 | -1,012539139 | 0,500541681 | -2,812583704 | 0,004914523 | 0,039598818 | BC100451 |
| ENSMUSG00000026857 | 9,45175925 | -1,004532152 | 0,416651314 | -3,090898489 | 0,001995518 | 0,0195767 | Ntmt1 |

**Supplementary table 5)** Unique genes in the heart tissue that were deregulated after delivery of VEGF mRNA to the heart via CPC SEC-EVs, HTB SEC-EVs, HUVEC SEC-EVs and LNPs.

| NA. | baseMean | log2FC.shrink. | lfcSE | stat | pvalue | padj | gene_symbol |
|---|---|---|---|---|---|---|---|
| | | | | LNPs | | | |
| USG000000 | 11,053959 | 1,73908287 | 0,334313 | 5,564009 | 2,63647E-08 | 1,04524E-06 | Slfn4 |
| USG000000 | 15,810415 | 1,37775407 | 0,22954 | 6,289681 | 3,18119E-10 | 1,70661E-08 | Itgb2 |
| USG000000 | 1,7619909 | 1,515961565 | 0,812429 | 2,828925 | 0,004670472 | 0,038014425 | Cdh1 |
| USG000000 | 9,8324759 | 1,192740994 | 0,32648 | 4,138923 | 3,48941E-05 | 0,000643934 | Gm2a |
| USG000000 | 3,8740051 | 1,421410754 | 0,635346 | 3,041206 | 0,002356327 | 0,022268605 | Slc7a7 |
| USG000000 | 18,266138 | 1,197912547 | 0,238543 | 5,37347 | 7,72355E-08 | 2,72515E-06 | S100a6 |
| USG000000 | 13,533453 | 1,400355192 | 0,258383 | 5,75244 | 8,79647E-09 | 3,8351E-07 | Timp1 |
| USG000000 | 8,4867179 | 1,91161164 | 0,361518 | 5,648152 | 1,62181E-08 | 6,70613E-07 | Cnn1 |
| USG000000 | 7,56314 | -1,032460741 | 0,528692 | -2,782908 | 0,005387413 | 0,042437642 | Vsig2 |
| USG000000 | 6,6360051 | 1,201907241 | 0,465152 | 3,273353 | 0,001062798 | 0,011870924 | Spi1 |
| USG000000 | 3,9896548 | -1,662212567 | 0,746705 | -3,001338 | 0,002687962 | 0,024591428 | Metrn |
| USG000000 | 21,104935 | -1,187688777 | 0,276631 | -4,703501 | 2,55738E-06 | 6,38503E-05 | Arrdc2 |
| USG000000 | 7,8565698 | 3,076921815 | 0,401742 | 7,919611 | 2,38255E-15 | 2,72877E-13 | Prkar2b |
| USG000000 | 1,6942138 | 1,408841909 | 0,857018 | 2,730503 | 0,006323782 | 0,048005525 | Fkbp11 |
| USG000000 | 6,6371986 | 2,460428563 | 0,417208 | 6,225428 | 4,80242E-10 | 2,52933E-08 | Slc25a1 |
| USG000000 | 9,0334859 | 1,491105021 | 0,329862 | 4,923309 | 8,50928E-07 | 2,38906E-05 | Ptpn6 |
| USG000000 | 5,7631506 | 1,211929691 | 0,392819 | 3,656312 | 0,00025587 | 0,003567599 | Adgre1 |
| USG000000 | 5,7186122 | -1,587057316 | 0,582667 | -3,373658 | 0,000741765 | 0,008774113 | Mri1 |
| USG000000 | 5,3941451 | 1,018717023 | 0,398192 | 3,22232 | 0,001271569 | 0,013694044 | Cstb |
| USG000000 | 37,110191 | 1,605016137 | 0,157731 | 10,35565 | 3,94484E-25 | 1,11623E-22 | Cd44 |
| USG000000 | 8,5599354 | 2,495795281 | 0,36536 | 7,112437 | 1,14012E-12 | 8,7053E-11 | Slc1a3 |
| USG000000 | 1,7819614 | 4,93061625 | 1,538902 | 3,791503 | 0,000149738 | 0,002274605 | Cyp2a5 |
| USG000000 | 3,5983692 | 1,541156589 | 0,52439 | 3,555095 | 0,000377842 | 0,004993258 | Mmp8 |
| USG000000 | 15,526916 | 1,00278749 | 0,226953 | 4,790099 | 1,66699E-06 | 4,36594E-05 | Rcn1 |
| USG000000 | 77,437731 | 1,716580496 | 0,173564 | 10,08463 | 6,46099E-24 | 1,63576E-21 | Prg4 |
| USG000000 | 11,956168 | -1,168539457 | 0,38532 | -3,604924 | 0,000312244 | 0,004227015 | Eps8l1 |
| USG000000 | 15,081367 | 1,051153571 | 0,270983 | 4,318604 | 1,57019E-05 | 0,00031709 | Crip1 |
| USG000000 | 3,5743761 | 1,707311691 | 0,609793 | 3,465317 | 0,000529606 | 0,006622831 | Aplp1 |
| USG000000 | 20,502272 | 1,166598043 | 0,223579 | 5,558153 | 2,72645E-08 | 1,07208E-06 | Mgst1 |
| USG000000 | 6,782499 | 1,181235699 | 0,40002 | 3,543742 | 0,000394491 | 0,005180068 | Met |
| USG000000 | 18,661579 | 1,167110519 | 0,246335 | 5,112404 | 3,18085E-07 | 9,97887E-06 | Fxyd5 |
| USG000000 | 1,6140385 | 2,0282716 | 0,957676 | 3,003034 | 0,002673025 | 0,024491696 | Exoc3l2 |
| USG000000 | 44,689599 | 1,023043578 | 0,145486 | 7,273294 | 3,50826E-13 | 2,90963E-11 | Steap4 |
| USG000000 | 16,819931 | 1,018881843 | 0,258381 | 4,373081 | 1,22505E-05 | 0,000255275 | Atp6v0d1 |
| USG000000 | 19,835497 | 1,240293202 | 0,229027 | 5,753215 | 8,75622E-09 | 3,82912E-07 | Aldh1a2 |
| USG000000 | 99,737995 | -1,029600656 | 0,128739 | -8,210551 | 2,20176E-16 | 2,78716E-14 | Ptgds |
| USG000000 | 7,2064431 | 1,532525325 | 0,374534 | 4,535939 | 5,73476E-06 | 0,000130947 | Aldh1a3 |
| USG000000 | 7,0001858 | 1,600918087 | 0,343306 | 5,060131 | 4,18968E-07 | 1,27787E-05 | Cybb |
| USG000000 | 4,0536387 | 1,651049231 | 0,604659 | 3,412362 | 0,000644025 | 0,00781659 | Fcgr1 |
| USG000000 | 17,545035 | 1,015975054 | 0,238604 | 4,65728 | 3,20415E-06 | 7,81065E-05 | Ctsz |
| USG000000 | 79,90837 | 1,592613524 | 0,108068 | 14,8836 | 4,21188E-50 | 3,03908E-47 | Sdc4 |
| USG000000 | 3,4314412 | -1,726342931 | 0,770273 | -3,004066 | 0,002663975 | 0,024439812 | Plxdc1 |
| USG000000 | 1,3136588 | 6,497592455 | 1,631725 | 4,020355 | 5,81104E-05 | 0,000994781 | Wfdc2 |
| USG000000 | 2,1170512 | 1,541234894 | 0,655471 | 3,132562 | 0,001732881 | 0,017438777 | Mmp9 |
| USG000000 | 23,655374 | 1,197497397 | 0,238038 | 5,378581 | 7,50752E-08 | 2,65542E-06 | Ptgis |

| | | | | | | | |
|---|---|---|---|---|---|---|---|
| USG000000 | 283,34833 | 1,270030119 | 0,084009 | 15,24244 | 1,84841E-52 | 1,40392E-49 | Gpx3 |
| USG000000 | 3,0439817 | 1,329580964 | 0,538133 | 3,199735 | 0,001375541 | 0,014537676 | Cd68 |
| USG000000 | 25,149748 | 2,567158135 | 0,23983 | 10,90184 | 1,12956E-27 | 3,97576E-25 | Ccl6 |
| USG000000 | 14,998004 | 1,18938533 | 0,255452 | 5,036127 | 4,75045E-07 | 1,42228E-05 | Ltv1 |
| USG000000 | 3,9777023 | 1,011271587 | 0,515758 | 2,822093 | 0,004771128 | 0,038659265 | Actr6 |
| USG000000 | 6,2002834 | 2,771196096 | 0,431433 | 6,722096 | 1,79128E-11 | 1,19676E-09 | Arg1 |
| USG000000 | 10,779972 | 1,119247108 | 0,293347 | 4,268511 | 1,96783E-05 | 0,000384792 | Snrpf |
| USG000000 | 19,107007 | 1,619140699 | 0,209265 | 7,977016 | 1,49913E-15 | 1,74014E-13 | Srgn |
| USG000000 | 21,637777 | 1,596311961 | 0,21128 | 7,788245 | 6,79463E-15 | 7,47372E-13 | Plek |
| USG000000 | 22,085385 | 1,376124934 | 0,226341 | 6,380573 | 1,76426E-10 | 9,86825E-09 | Csrp2 |
| USG000000 | 8,7153393 | 1,781633681 | 0,324386 | 5,833962 | 5,41264E-09 | 2,42577E-07 | Efemp1 |
| USG000000 | 184,80037 | 2,124408012 | 0,095002 | 22,44682 | 1,3746E-111 | 3,3062E-108 | Myl7 |
| USG000000 | 50,442855 | 1,184387558 | 0,147038 | 8,300809 | 1,03405E-16 | 1,36902E-14 | Aebp1 |
| USG000000 | 70,922261 | 2,190382289 | 0,137247 | 16,06833 | 4,25387E-58 | 3,41042E-55 | Acaca |
| USG000000 | 8,0808476 | 1,5345152 | 0,311702 | 5,296285 | 1,18183E-07 | 4,01916E-06 | Id2 |
| USG000000 | 5,5145934 | 1,45459216 | 0,556382 | 3,327292 | 0,000876943 | 0,010116039 | Adcy3 |
| USG000000 | 7,5779377 | 3,101555757 | 0,434788 | 7,414101 | 1,22453E-13 | 1,14007E-11 | Krt19 |
| USG000000 | 49,307518 | 1,261390781 | 0,137164 | 9,343293 | 9,3384E-21 | 1,95308E-18 | Acly |
| USG000000 | 17,902143 | 1,138997336 | 0,219577 | 5,527644 | 3,2456E-08 | 1,23581E-06 | Lsm12 |
| USG000000 | 9,1430168 | 1,971645273 | 0,3353 | 6,208191 | 5,3598E-10 | 2,81263E-08 | Pygl |
| USG000000 | 11,593579 | 1,097157588 | 0,284771 | 4,299722 | 1,71013E-05 | 0,000340474 | Smoc1 |
| USG000000 | 22,929515 | 1,533552546 | 0,206087 | 7,693258 | 1,43434E-14 | 1,48914E-12 | Net1 |
| USG000000 | 46,676404 | -1,045735733 | 0,224912 | -5,005371 | 5,57546E-07 | 1,63869E-05 | Fam193b |
| USG000000 | 95,447293 | 1,255555338 | 0,101184 | 12,55077 | 3,93642E-36 | 2,27226E-33 | Vcan |
| USG000000 | 3,9475509 | 1,923622197 | 0,475304 | 4,516561 | 6,2852E-06 | 0,000141721 | Emb |
| USG000000 | 12,535848 | 1,299143878 | 0,273232 | 5,154326 | 2,54544E-07 | 8,10889E-06 | Fam213a |
| USG000000 | 20,49153 | 2,422172412 | 0,477467 | 5,45683 | 4,8471E-08 | 1,78896E-06 | Itih4 |
| USG000000 | 41,805746 | 1,596270997 | 0,185643 | 8,817742 | 1,16791E-18 | 1,91524E-16 | Tkt |
| USG000000 | 27,178526 | 1,306607873 | 0,196606 | 6,917482 | 4,59742E-12 | 3,28442E-10 | Lcp1 |
| USG000000 | 10,105758 | -1,049781074 | 0,361204 | -3,485193 | 0,000491782 | 0,006230819 | Ebf2 |
| USG000000 | 160,27351 | 1,024323088 | 0,08435 | 12,26554 | 1,38688E-34 | 7,69773E-32 | Pabpc1 |
| USG000000 | 3,1266302 | 1,050905681 | 0,553866 | 2,796097 | 0,005172384 | 0,041125443 | Sqle |
| USG000000 | 19,657974 | 1,072741731 | 0,21699 | 5,289687 | 1,22526E-07 | 4,14091E-06 | Col14a1 |
| USG000000 | 7,8286304 | 1,875400353 | 0,327691 | 6,069302 | 1,28467E-09 | 6,32734E-08 | Enpp2 |
| USG000000 | 9,4616484 | 1,665352844 | 0,344279 | 5,226351 | 1,72888E-07 | 5,70926E-06 | Fgf12 |
| USG000000 | 11,428608 | -1,171681463 | 0,358328 | -3,796394 | 0,000146816 | 0,002234922 | Gpt |
| USG000000 | 8,1047809 | 1,21946783 | 0,321603 | 4,257521 | 2,06706E-05 | 0,000400937 | Alcam |
| USG000000 | 7,2467429 | -4,187572834 | 1,225608 | -3,966039 | 7,30768E-05 | 0,001220569 | Slc15a2 |
| USG000000 | 63,141375 | 1,449116693 | 0,190064 | 7,875458 | 3,39496E-15 | 3,79788E-13 | Gpd1 |
| USG000000 | 3,7826617 | 1,096439497 | 0,501413 | 2,970271 | 0,00297537 | 0,026702466 | Krt7 |
| USG000000 | 8,741868 | 3,211615764 | 0,408041 | 8,109862 | 5,06774E-16 | 6,1456E-14 | Krt18 |
| USG000000 | 19,004584 | 1,304517207 | 0,235797 | 5,862324 | 4,56432E-09 | 2,06482E-07 | Igfbp6 |
| USG000000 | 3,0745 | 2,056398629 | 0,612716 | 3,917654 | 8,9415E-05 | 0,001454733 | Cxcl13 |
| USG000000 | 5,7728385 | 1,128933754 | 0,406016 | 3,403126 | 0,000666195 | 0,008031631 | Tmem176a |
| USG000000 | 2,8410611 | 1,235959497 | 0,621547 | 2,877319 | 0,004010695 | 0,033827201 | Acat2 |
| USG000000 | 7,6604282 | 1,232620781 | 0,337255 | 4,148209 | 3,35087E-05 | 0,000621547 | Pla2g7 |
| USG000000 | 263,94599 | 2,198530822 | 0,089883 | 24,51962 | 9,1249E-133 | 3,292E-129 | C3 |
| USG000000 | 4,8636152 | -1,142792655 | 0,573085 | -2,84056 | 0,004503447 | 0,036988755 | Ip6k3 |
| USG000000 | 9,1865848 | -1,53865032 | 0,510704 | -3,595472 | 0,000323804 | 0,004354906 | Col11a2 |

| | | | | | | | |
|---|---|---|---|---|---|---|---|
| USG000000 | 7,3955981 | 1,301638553 | 0,355152 | 4,152967 | 3,28193E-05 | 0,000609543 | Tmem173 |
| USG000000 | 1,503443 | 3,071538387 | 1,387208 | 3,064283 | 0,002181924 | 0,020960724 | C2 |
| USG000000 | 12,928561 | 1,380926604 | 0,302704 | 4,967261 | 6,7905E-07 | 1,95987E-05 | Cidea |
| USG000000 | 18,418353 | 11,53637667 | 1,204793 | 8,727327 | 2,60762E-18 | 4,00324E-16 | Scgb1a1 |
| USG000000 | 22,931267 | 1,374706289 | 0,189478 | 7,501799 | 6,29479E-14 | 5,93726E-12 | Anxa1 |
| USG000000 | 3,8853694 | 1,413234959 | 0,576138 | 3,179811 | 0,00147371 | 0,015377523 | Ms4a6d |
| USG000000 | 21,965062 | 1,720977814 | 0,203271 | 8,691697 | 3,57065E-18 | 5,25797E-16 | Il33 |
| USG000000 | 30,352454 | 1,167783623 | 0,223851 | 5,553485 | 2,8003E-08 | 1,09515E-06 | Pcx |
| USG000000 | 2,9952478 | 1,061017826 | 0,617816 | 2,723242 | 0,006464472 | 0,048893501 | Fosl1 |
| USG000000 | 13,436164 | 1,331438012 | 0,285113 | 5,050185 | 4,41382E-07 | 1,33534E-05 | Msr1 |
| USG000000 | 1,2611831 | 1,905282173 | 1,298508 | 2,723248 | 0,006464351 | 0,048893501 | Bnc1 |
| USG000000 | 10,158447 | 2,27550228 | 0,30531 | 7,710831 | 1,25001E-14 | 1,31671E-12 | Cbr2 |
| USG000000 | 170,57794 | 3,535071452 | 0,154916 | 22,92014 | 2,9265E-116 | 8,4464E-113 | Fasn |
| USG000000 | 7,3096165 | 1,216447182 | 0,334747 | 4,123354 | 3,73395E-05 | 0,000682083 | Itgb8 |
| USG000000 | 5,9504769 | -1,107059923 | 0,535 | -2,86059 | 0,004228535 | 0,035252449 | Cdk2 |
| USG000000 | 32,049932 | 1,763985557 | 0,169725 | 10,57422 | 3,92433E-26 | 1,20494E-23 | Rps26 |
| USG000000 | 5,6385966 | 2,287001815 | 0,494978 | 5,039836 | 4,65931E-07 | 1,39789E-05 | Cyp2e1 |
| USG000000 | 3,2297379 | 1,227505804 | 0,531355 | 3,071567 | 0,002129385 | 0,020595944 | Ifitm1 |
| USG000000 | 30,127302 | 1,080881356 | 0,172555 | 6,539057 | 6,19078E-11 | 3,85082E-09 | Ifitm3 |
| USG000000 | 6,7392224 | 1,614819968 | 0,345389 | 5,070968 | 3,95797E-07 | 1,22046E-05 | Ccr1 |
| USG000000 | 13,730395 | -1,018986024 | 0,312699 | -3,766035 | 0,000165861 | 0,002482918 | Carf |
| USG000000 | 3,0508733 | 1,167606986 | 0,51581 | 3,027694 | 0,002464275 | 0,02297284 | Il18rap |
| USG000000 | 2,9228136 | 1,212795818 | 0,552591 | 3,005747 | 0,002649292 | 0,024336052 | Il18r1 |
| USG000000 | 3,824296 | 1,430629667 | 0,476625 | 3,613537 | 0,000302048 | 0,00411213 | Sphkap |
| USG000000 | 3,86709 | 2,322865578 | 0,523897 | 4,868361 | 1,12528E-06 | 3,06394E-05 | Cxcr2 |
| USG000000 | 177,12562 | 1,754489926 | 0,080566 | 21,87232 | 4,7673E-106 | 9,8281E-103 | Fn1 |
| USG000000 | 6,7610917 | 1,685447202 | 0,37319 | 4,926914 | 8,35386E-07 | 2,35458E-05 | Tmem163 |
| USG000000 | 363,68259 | 1,140565664 | 0,129834 | 9,019956 | 1,88166E-19 | 3,43724E-17 | Cfh |
| USG000000 | 1,4055854 | 6,136983638 | 1,680092 | 3,828708 | 0,000128818 | 0,001990334 | Marco |
| USG000000 | 22,987662 | 1,212298483 | 0,202163 | 6,296825 | 3,03804E-10 | 1,63589E-08 | Ptprc |
| USG000000 | 4,0734952 | 3,727199171 | 0,704516 | 5,655797 | 1,55125E-08 | 6,46997E-07 | Pigr |
| USG000000 | 11,238577 | 1,268682571 | 0,325002 | 4,361033 | 1,2945E-05 | 0,000268019 | Lamc2 |
| USG000000 | 7,3988508 | 1,921647017 | 0,437836 | 4,822777 | 1,41573E-06 | 3,76251E-05 | F5 |
| USG000000 | 6,9626786 | 1,705673713 | 0,400756 | 4,696441 | 2,64734E-06 | 6,57362E-05 | Sell |
| USG000000 | 7,278462 | 2,22818044 | 0,342791 | 6,790732 | 1,11566E-11 | 7,55874E-10 | Sele |
| USG000000 | 6,5984383 | 1,525736549 | 0,365288 | 4,62099 | 3,81914E-06 | 9,12483E-05 | Fam107b |
| USG000000 | 371,1688 | 1,102693234 | 0,057721 | 19,1846 | 4,97788E-82 | 5,98632E-79 | Vim |
| USG000000 | 13,039036 | 1,247126694 | 0,307177 | 4,503639 | 6,67997E-06 | 0,000149455 | Gpd2 |
| USG000000 | 9,4517593 | -1,004532152 | 0,416651 | -3,090898 | 0,001995518 | 0,0195767 | Ntmt1 |
| USG000000 | 8,2666297 | 1,037133105 | 0,331053 | 3,668454 | 0,000244021 | 0,003445667 | Itga4 |
| USG000000 | 4,896085 | 3,075782098 | 0,594797 | 5,548607 | 2,87954E-08 | 1,12007E-06 | Lrp2 |
| USG000000 | 10,694445 | 1,329316733 | 0,31153 | 4,692883 | 2,69381E-06 | 6,65657E-05 | Ube2l6 |
| USG000000 | 15,454564 | 1,302538694 | 0,247583 | 5,607957 | 2,04729E-08 | 8,29898E-07 | Hsd17b12 |
| USG000000 | 22,496876 | 11,64789288 | 1,196346 | 8,878065 | 6,80337E-19 | 1,18288E-16 | Bpifa1 |
| USG000000 | 5,0453298 | 5,134034219 | 0,745716 | 7,149991 | 8,67836E-13 | 6,73319E-11 | Bpifb1 |
| USG000000 | 8,178802 | 1,030110816 | 0,337364 | 3,60969 | 0,000306563 | 0,004169666 | Pag1 |
| USG000000 | 39,279925 | 1,118328761 | 0,157238 | 7,355454 | 1,90279E-13 | 1,66419E-11 | Fabp5 |
| USG000000 | 13,852808 | 2,159538733 | 0,251328 | 8,784565 | 1,56971E-18 | 2,50762E-16 | S100a11 |
| USG000000 | 34,941698 | 1,263303415 | 0,19715 | 6,686035 | 2,29299E-11 | 1,51096E-09 | Vcam1 |

| | | | | | | | |
|---|---|---|---|---|---|---|---|
| USG000000 | 8,7407423 | 1,085338901 | 0,325738 | 3,850624 | 0,000117817 | 0,001838078 | Ctso |
| USG000000 | 6,5337338 | 1,346536913 | 0,39538 | 3,936324 | 8,27392E-05 | 0,001358373 | Pdgfc |
| USG000000 | 29,335616 | 1,073796398 | 0,222211 | 5,189568 | 2,10782E-07 | 6,7746E-06 | Ecm1 |
| USG000000 | 20,802969 | 1,495694189 | 0,209978 | 7,381284 | 1,56771E-13 | 1,41397E-11 | Gbp2 |
| USG000000 | 17,113993 | 1,032412133 | 0,218935 | 5,073021 | 3,9155E-07 | 1,20995E-05 | Erp44 |
| USG000000 | 33,264067 | 1,086505238 | 0,216954 | 5,356539 | 8,4831E-08 | 2,97135E-06 | Svep1 |
| USG000000 | 11,404154 | 1,793434794 | 0,274313 | 6,820262 | 9,08747E-12 | 6,24482E-10 | Pdpn |
| USG000000 | 48,738999 | 1,10277807 | 0,138629 | 8,198448 | 2,43511E-16 | 3,05575E-14 | Cap1 |
| USG000000 | 8,9386654 | 1,497859879 | 0,409299 | 4,166612 | 3,09161E-05 | 0,000579415 | Csf3r |
| USG000000 | 1,7590365 | 1,744037212 | 0,897381 | 2,885181 | 0,003911883 | 0,033168261 | Rspo1 |
| USG000000 | 4,3402443 | 1,355996545 | 0,464535 | 3,531119 | 0,000413805 | 0,00537984 | Fgr |
| USG000000 | 59,428005 | 1,050996878 | 0,137314 | 7,884006 | 3,17049E-15 | 3,57448E-13 | Rpl22 |
| USG000000 | 128,65397 | 1,395635181 | 0,126957 | 11,13407 | 8,56356E-29 | 3,34002E-26 | Nppb |
| USG000000 | 57,450114 | 1,004677012 | 0,123483 | 8,364685 | 6,02759E-17 | 8,12936E-15 | Tbc1d1 |
| USG000000 | 2,972295 | 1,677224307 | 0,639635 | 3,335608 | 0,000851131 | 0,009873527 | Spp1 |
| USG000000 | 4,6208733 | 3,154854245 | 0,533224 | 6,247592 | 4,16828E-10 | 2,20338E-08 | Ppbp |
| USG000000 | 10,664311 | 1,371392209 | 0,302732 | 4,94619 | 7,568E-07 | 2,15412E-05 | Pf4 |
| USG000000 | 14,479898 | 2,188839977 | 0,253873 | 8,842506 | 9,35953E-19 | 1,57055E-16 | Aacs |
| USG000000 | 18,228005 | 1,030772331 | 0,258879 | 4,39944 | 1,08531E-05 | 0,000230324 | Oasl2 |
| USG000000 | 17,365796 | 1,370747269 | 0,299013 | 4,980585 | 6,33923E-07 | 1,84811E-05 | Tmem176b |
| USG000000 | 4,7331115 | 6,315513147 | 1,12494 | 5,946785 | 2,7346E-09 | 1,29812E-07 | Reg3g |
| USG000000 | 11,960129 | 1,2051065 | 0,322815 | 4,207664 | 2,58024E-05 | 0,000489297 | Aldh1l1 |
| USG000000 | 3,3592568 | 1,450721508 | 0,564228 | 3,285971 | 0,001016314 | 0,011449196 | Olr1 |
| USG000000 | 35,627025 | 1,040141955 | 0,175759 | 6,193918 | 5,86868E-10 | 3,04074E-08 | Emp1 |
| USG000000 | 8,4255965 | 1,157925186 | 0,367217 | 3,700834 | 0,000214892 | 0,003091826 | Arhgdib |
| USG000000 | 7,0912331 | 6,117658593 | 0,884587 | 7,206909 | 5,72364E-13 | 4,51355E-11 | Cidec |
| USG000000 | 23,109926 | 1,285729425 | 0,196638 | 6,810952 | 9,6955E-12 | 6,63108E-10 | Cd9 |
| USG000000 | 4,6697081 | 5,569710141 | 1,105557 | 5,403854 | 6,52241E-08 | 2,35312E-06 | Plin1 |
| USG000000 | 3,981728 | 1,084413791 | 0,538354 | 2,863634 | 0,00418812 | 0,035016666 | 2200002D01Rik |
| USG000000 | 11,311293 | 1,164307681 | 0,315546 | 4,167417 | 3,0807E-05 | 0,000578122 | Pde3b |
| USG000000 | 46,799847 | 1,229965084 | 0,141725 | 8,887367 | 6,25741E-19 | 1,10123E-16 | Stard10 |
| USG000000 | 79,694883 | 1,342093643 | 0,121892 | 11,18046 | 5,08272E-29 | 2,03746E-26 | Dkk3 |
| USG000000 | 38,76329 | 1,083625488 | 0,155264 | 7,258954 | 3,90096E-13 | 3,1805E-11 | Itgam |
| USG000000 | 4,4958776 | 1,017777408 | 0,489497 | 2,894871 | 0,003793143 | 0,032370697 | Rgs10 |
| USG000000 | 3,9018205 | 1,435626652 | 0,547388 | 3,31871 | 0,000904343 | 0,010382314 | Fgfr2 |
| USG000000 | 30,333843 | 1,194875792 | 0,190705 | 6,531703 | 6,50261E-11 | 4,01022E-09 | Sh3bgrl |
| USG000000 | 3,4244174 | 2,519993116 | 0,732137 | 3,989453 | 6,62258E-05 | 0,001120404 | Car5b |
| USG000000 | 8,0762239 | 1,495682095 | 0,338275 | 4,835194 | 1,33016E-06 | 3,58126E-05 | G6pdx |
| USG000000 | 1,6082038 | 2,713399834 | 1,272509 | 2,989552 | 0,002793873 | 0,025453528 | Adrb3 |
| USG000000 | 6,7631985 | 2,073724974 | 0,39084 | 5,667263 | 1,45097E-08 | 6,06925E-07 | Gpm6a |
| USG000000 | 18,733159 | 1,406971266 | 0,213066 | 6,87909 | 6,0236E-12 | 4,21974E-10 | Gsr |
| USG000000 | 9,5633749 | 1,847070236 | 0,367709 | 5,396719 | 6,78706E-08 | 2,43037E-06 | Hpgd |
| USG000000 | 9,5571148 | 1,077083266 | 0,272787 | 4,383424 | 1,16828E-05 | 0,000245408 | Cdh11 |
| USG000000 | 92,569782 | 1,071445327 | 0,111042 | 9,853525 | 6,61812E-23 | 1,64666E-20 | Mt2 |
| USG000000 | 39,130364 | -1,171845439 | 0,220985 | -5,633869 | 1,7621E-08 | 7,24471E-07 | Jak3 |
| USG000000 | 8,6046199 | 1,057700726 | 0,337446 | 3,678893 | 0,000234248 | 0,003325417 | Cotl1 |
| USG000000 | 15,344422 | 2,001636691 | 0,288505 | 7,213142 | 5,46753E-13 | 4,33527E-11 | Smpd3 |
| USG000000 | 7,9704857 | 1,401627101 | 0,339507 | 4,572695 | 4,81491E-06 | 0,000112252 | Adamts8 |
| USG000000 | 1,7505154 | 1,825461787 | 0,861911 | 2,990444 | 0,002785723 | 0,025395305 | Plet1 |

| | | | | | | | |
|---|---|---|---|---|---|---|---|
| USG000000 | 31,132717 | 1,075441212 | 0,185033 | 6,107094 | 1,01461E-09 | 5,04893E-08 | Stt3a |
| USG000000 | 9,4736924 | 1,07179757 | 0,328993 | 3,768467 | 0,000164253 | 0,002472847 | Tmod2 |
| USG000000 | 55,872392 | 1,197869465 | 0,136944 | 8,945208 | 3,71251E-19 | 6,61423E-17 | Anxa2 |
| USG000000 | 4,5329704 | 1,46652433 | 0,509249 | 3,512523 | 0,000443874 | 0,00570904 | Nnmt |
| USG000000 | 6,9587987 | 1,318017822 | 0,350345 | 4,245203 | 2,18395E-05 | 0,000421346 | Ptgs2 |
| USG000000 | 8,0199863 | 9,99870826 | 1,202533 | 7,689598 | 1,47598E-14 | 1,52142E-12 | Ltf |
| USG000000 | 19,009196 | 1,047394402 | 0,21597 | 5,196768 | 2,02783E-07 | 6,57609E-06 | Slco2a1 |
| USG000000 | 28,343398 | 1,688627945 | 0,184946 | 9,3396 | 9,66983E-21 | 1,99351E-18 | Trf |
| USG000000 | 3,3954133 | -1,547994477 | 0,820474 | -2,761097 | 0,005760753 | 0,044747733 | Inha |
| USG000000 | 5,4375045 | 2,783892161 | 0,479553 | 6,132064 | 8,67461E-10 | 4,36179E-08 | Cdo1 |
| USG000000 | 14,220025 | 1,02076229 | 0,256631 | 4,399492 | 1,08504E-05 | 0,000230324 | Nol12 |
| USG000000 | 2,9831487 | 1,940009272 | 0,613505 | 3,758663 | 0,000170824 | 0,002549288 | Rac2 |
| USG000000 | 2,932998 | 3,094816124 | 0,873595 | 4,0805 | 4,4939E-05 | 0,000802617 | Apol6 |
| USG000000 | 3,2606068 | 1,650444464 | 0,533042 | 3,692571 | 0,000221998 | 0,003177815 | Tlr13 |
| USG000000 | 29,339712 | -1,068040005 | 0,257378 | -4,559398 | 5,13005E-06 | 0,000118732 | Arhgap27 |
| USG000000 | 4,9090716 | 1,061121116 | 0,438508 | 3,122159 | 0,001795302 | 0,017941054 | Scara3 |
| USG000000 | 20,734288 | 1,498678749 | 0,192115 | 8,038106 | 9,12377E-16 | 1,07045E-13 | Plvap |
| USG000000 | 3,5307925 | 1,271882036 | 0,549908 | 3,08058 | 0,00206598 | 0,020063366 | Pdk3 |
| USG000000 | 11,426746 | -1,393624481 | 0,404994 | -3,968372 | 7,23652E-05 | 0,001210084 | Adrb1 |
| USG000000 | 8,3215397 | 1,273979935 | 0,32227 | 4,412715 | 1,02082E-05 | 0,000219219 | Ccl7 |
| USG000000 | 11,728038 | 2,005953493 | 0,341342 | 6,201955 | 5,57659E-10 | 2,91579E-08 | Ccl2 |
| USG000000 | 42,716308 | 1,20691288 | 0,152106 | 8,14892 | 3,67189E-16 | 4,56802E-14 | Tgfbi |
| USG000000 | 8,8665563 | 1,570244318 | 0,356958 | 4,82924 | 1,37055E-06 | 3,67628E-05 | Myrf |
| USG000000 | 68,596075 | 1,048541076 | 0,113212 | 9,43653 | 3,85329E-21 | 8,17748E-19 | Calm2 |
| USG000000 | 12,873943 | 1,170996792 | 0,331503 | 4,029146 | 5,598E-05 | 0,000965169 | Lum |
| USG000000 | 28,030234 | 1,013402538 | 0,193419 | 5,558387 | 2,72279E-08 | 1,07208E-06 | Sep15 |
| USG000000 | 513,05534 | 1,160959645 | 0,057978 | 20,10163 | 7,14091E-90 | 9,36823E-87 | Eef1a1 |
| USG000000 | 26,801969 | -1,069168998 | 0,24866 | -4,705075 | 2,53773E-06 | 6,34696E-05 | Scarf1 |
| USG000000 | 16,87633 | 1,058519696 | 0,256208 | 4,552585 | 5,29906E-06 | 0,000122098 | Degs1 |
| USG000000 | 23,241737 | 1,483653631 | 0,189427 | 8,07732 | 6,62055E-16 | 7,83124E-14 | Ctss |
| USG000000 | 33,333211 | 2,093065902 | 0,164649 | 12,86246 | 7,32138E-38 | 4,59369E-35 | Pkhd1l1 |
| USG000000 | 1,5330067 | 6,900294858 | 1,43825 | 4,652037 | 3,28672E-06 | 7,99843E-05 | Scgb3a2 |
| USG000000 | 18,210495 | 1,287265316 | 0,242225 | 5,652316 | 1,58301E-08 | 6,58295E-07 | Bpgm |
| USG000000 | 45,378779 | 1,058232243 | 0,140389 | 7,761601 | 8,38639E-15 | 9,03165E-13 | Rps21 |
| USG000000 | 3,7522303 | 1,709875211 | 0,544693 | 3,736498 | 0,000186601 | 0,002753408 | Stx11 |
| USG000000 | 3,6375174 | 1,39363036 | 0,566905 | 3,200438 | 0,001372187 | 0,014537676 | Isg20 |
| USG000000 | 6,8411257 | -2,063941935 | 0,606329 | -3,934012 | 8,35398E-05 | 0,001369957 | Rnf122 |
| USG000000 | 10,870178 | 1,176959715 | 0,332163 | 4,033158 | 5,50323E-05 | 0,000953387 | Prss23 |
| USG000000 | 6,6369738 | -1,097088133 | 0,531715 | -2,864632 | 0,004174937 | 0,034946937 | Nrtn |
| USG000000 | 12,152446 | 1,118520622 | 0,252263 | 4,831134 | 1,35758E-06 | 3,64827E-05 | Dse |
| USG000000 | 32,452351 | -1,044159384 | 0,2067 | -5,360565 | 8,29623E-08 | 2,91296E-06 | Magi2 |
| USG000000 | 6,8149032 | -1,13033179 | 0,465811 | -3,121764 | 0,001797711 | 0,017941054 | Rgl3 |
| USG000000 | 142,6716 | 1,471985955 | 0,115934 | 12,84404 | 9,28995E-38 | 5,58597E-35 | Thbs1 |
| USG000000 | 6,2183023 | 1,162527558 | 0,464569 | 3,198709 | 0,001380443 | 0,014564483 | Sema3d |
| USG000000 | 8,8446165 | -1,273360409 | 0,47301 | -3,327788 | 0,000875386 | 0,010106153 | Slc26a10 |
| USG000000 | 6,2475956 | 1,140378626 | 0,374524 | 3,615922 | 0,000299281 | 0,004078305 | Ildr2 |
| USG000000 | 5,3085271 | 1,695699682 | 0,380513 | 4,871497 | 1,10756E-06 | 3,02711E-05 | Cd53 |
| USG000000 | 4,9965079 | 2,937613016 | 0,544208 | 5,753523 | 8,74025E-09 | 3,82912E-07 | Chil3 |
| USG000000 | 7,7885705 | 2,898318373 | 0,43704 | 6,927156 | 4,29386E-12 | 3,08282E-10 | Elovl6 |

| | | | | | | | |
|---|---|---|---|---|---|---|---|
| USG000000 | 7,186391 | 1,165228751 | 0,466935 | 3,19962 | 0,001376087 | 0,014537676 | Shtn1 |
| USG000000 | 6,8596653 | 1,246101892 | 0,342695 | 4,120539 | 3,77987E-05 | 0,000686994 | Fmod |
| USG000000 | 5,9018312 | 2,179353784 | 0,497217 | 4,827704 | 1,38116E-06 | 3,69788E-05 | Mbp |
| USG000000 | 1241,8161 | 1,87204061 | 0,054144 | 34,67611 | 1,8053E-263 | 1,3026E-259 | Nppa |
| USG000000 | 4,5153705 | 1,507242841 | 0,570218 | 3,336145 | 0,000849487 | 0,00986822 | Pnpla3 |
| USG000000 | 34,522839 | 1,546848148 | 0,181817 | 8,724177 | 2,68123E-18 | 4,0305E-16 | S100a10 |
| USG000000 | 35,39986 | 2,309968574 | 0,168419 | 13,86073 | 1,09552E-43 | 7,18612E-41 | Sln |
| USG000000 | 106,77554 | -1,195702711 | 0,162972 | -7,565302 | 3,86966E-14 | 3,77318E-12 | Kank3 |
| USG000000 | 4,6562231 | 3,093737113 | 0,616733 | 5,415809 | 6,10122E-08 | 2,21223E-06 | Upk3b |
| USG000000 | 3,6913378 | 2,605336502 | 0,671801 | 4,380811 | 1,18238E-05 | 0,000247648 | Lrrn4 |
| USG000000 | 5,5557571 | 1,646315202 | 0,43139 | 4,2945 | 1,75088E-05 | 0,000346122 | Pyhin1 |
| USG000000 | 36,210747 | -1,091828648 | 0,220081 | -5,296001 | 1,18366E-07 | 4,01916E-06 | Als2cl |
| USG000000 | 27,639036 | 1,800314409 | 0,203078 | 9,070702 | 1,18253E-19 | 2,21624E-17 | Ctla2a |
| USG000000 | 21,12661 | 1,778651876 | 0,237425 | 7,747984 | 9,33628E-15 | 9,98013E-13 | Fam46c |
| USG000000 | 5,1096522 | -1,221682597 | 0,613451 | -2,825481 | 0,004720976 | 0,038317434 | Fam186a |
| USG000000 | 3,3981536 | 1,723446127 | 0,576852 | 3,603458 | 0,000314011 | 0,004246953 | Krt14 |
| USG000000 | 3,3454746 | 2,0958526 | 0,660452 | 3,762925 | 0,000167937 | 0,002508805 | Basp1 |
| USG000000 | 4,2442729 | 1,251660154 | 0,557069 | 3,032037 | 0,002429097 | 0,022792127 | Tusc5 |
| USG000000 | 6,9535728 | 1,328029263 | 0,378153 | 4,02794 | 5,62676E-05 | 0,000967817 | Rpl14-ps1 |
| USG000000 | 2,3020387 | 1,191921129 | 0,708509 | 2,721558 | 0,006497505 | 0,049014898 | Gprc5a |
| USG000000 | 12,5391 | 1,451223927 | 0,316429 | 4,97861 | 6,40427E-07 | 1,85956E-05 | Mpeg1 |
| USG000000 | 1,4950778 | 2,397481163 | 1,378283 | 2,718278 | 0,006562263 | 0,04941971 | Vat1l |
| USG000000 | 7,8645553 | 2,48374707 | 0,335755 | 7,655899 | 1,91965E-14 | 1,95088E-12 | Cd24a |
| USG000000 | 1,4306827 | 1,771656623 | 1,048996 | 2,8076 | 0,004991221 | 0,040149564 | BC048546 |
| USG000000 | 7,9294061 | 1,982395622 | 0,358697 | 5,868617 | 4,39445E-09 | 2,00052E-07 | Cd300lf |
| USG000000 | 5,6321141 | 1,812711365 | 0,481654 | 4,265398 | 1,99546E-05 | 0,000389141 | Osr1 |
| USG000000 | 6,4080055 | 1,273434121 | 0,385028 | 3,845034 | 0,000120536 | 0,001876433 | Ndnf |
| USG000000 | 12,746371 | 1,611593551 | 0,275321 | 6,17118 | 6,7782E-10 | 3,48101E-08 | Ccr2 |
| USG000000 | 25,797688 | 1,075543281 | 0,188597 | 6,000654 | 1,96524E-09 | 9,54895E-08 | Sorl1 |
| USG000000 | 14,940205 | 2,37019468 | 0,322383 | 7,612913 | 2,67986E-14 | 2,66711E-12 | Krt8 |
| USG000000 | 3,9450321 | 1,013508152 | 0,527491 | 2,80158 | 0,005085304 | 0,040679612 | Arhgap15 |
| USG000000 | 8,5308682 | 2,19915363 | 0,345872 | 6,658539 | 2,76563E-11 | 1,79778E-09 | Lgals3 |
| USG000000 | 3,2342575 | -2,776017969 | 1,257571 | -3,031343 | 0,002434683 | 0,022829704 | Igfn1 |
| USG000000 | 83,535477 | 1,073543859 | 0,111787 | 9,782236 | 1,34212E-22 | 3,22803E-20 | Ezr |
| USG000000 | 35,413447 | 1,040269678 | 0,161412 | 6,69977 | 2,08748E-11 | 1,38823E-09 | Rap1b |
| USG000000 | 6,1566443 | 1,155837209 | 0,362555 | 3,739582 | 0,000184327 | 0,002729432 | Trim30b |
| USG000000 | 8,2630754 | -1,329365493 | 0,478164 | -3,402797 | 0,000666998 | 0,008034596 | 1700030K09Rik |
| USG000000 | 3,6466463 | 8,789549852 | 1,266653 | 6,411479 | 1,44115E-10 | 8,28573E-09 | Cyp2f2 |
| USG000000 | 26,75523 | 1,262358651 | 0,198928 | 6,615819 | 3,69501E-11 | 2,38047E-09 | Aldh1a1 |
| USG000000 | 21,180686 | 2,249669263 | 0,236944 | 9,695086 | 3,1637E-22 | 7,24689E-20 | Slc39a8 |
| USG000000 | 3,7419399 | 1,450232759 | 0,591186 | 3,200055 | 0,001374013 | 0,014537676 | Adam12 |
| USG000000 | 8,677428 | 1,013311573 | 0,311492 | 3,771501 | 0,000162269 | 0,002446915 | Mmrn1 |
| USG000000 | 4,7524524 | 1,631312127 | 0,491489 | 3,88114 | 0,000103968 | 0,001654203 | Tmem119 |
| USG000000 | 6,2320355 | 1,185792845 | 0,372393 | 3,747464 | 0,000178631 | 0,002652089 | Lmntd1 |
| USG000000 | 1,668836 | 5,107855121 | 1,416015 | 4,105001 | 4,04313E-05 | 0,000728419 | Sec14l3 |
| USG000000 | 19,115213 | 1,056361578 | 0,204677 | 5,493151 | 3,94826E-08 | 1,47993E-06 | C1ra |
| USG000000 | 9,698514 | 2,229750886 | 0,366329 | 6,402836 | 1,52517E-10 | 8,66526E-09 | Gpc3 |
| USG000000 | 4,5186089 | -1,292423591 | 0,656602 | -2,804532 | 0,005038963 | 0,040443419 | Nod2 |
| USG000000 | 2,7369915 | 2,024484661 | 0,679946 | 3,609025 | 0,00030735 | 0,004176434 | S100a8 |

| | | | | | | | |
|---|---|---|---|---|---|---|---|
| USG000000 | 6,8508538 | 2,802244871 | 0,402312 | 7,233161 | 4,7188E-13 | 3,82567E-11 | S100a9 |
| USG000000 | 8,5875387 | 1,205488282 | 0,355658 | 3,912301 | 9,14208E-05 | 0,001479028 | Pla2g4a |
| USG000000 | 104,16107 | 1,061959969 | 0,116919 | 9,254539 | 2,15156E-20 | 4,22496E-18 | Npm1 |
| USG000000 | 6,1506463 | 1,285128914 | 0,386252 | 3,875379 | 0,000106459 | 0,001691966 | Idi1 |
| USG000000 | 3,7697414 | 1,213116428 | 0,452526 | 3,339725 | 0,000838615 | 0,009759717 | Cxcl2 |
| USG000000 | 110,8371 | 1,131451594 | 0,11165 | 10,30694 | 6,55522E-25 | 1,8192E-22 | Rpl23a |
| USG000000 | 60,96376 | 1,167134801 | 0,138476 | 8,642695 | 5,49023E-18 | 7,8445E-16 | Gda |
| USG000000 | 4,3242927 | 1,834658913 | 0,557656 | 3,863512 | 0,000111768 | 0,001758921 | Fcer1g |
| USG000000 | 12,155886 | 1,262364891 | 0,293119 | 4,720074 | 2,35758E-06 | 5,94795E-05 | Pde1a |
| USG000000 | 8,0632393 | 1,033207884 | 0,363276 | 3,439521 | 0,000582745 | 0,007175421 | Fcgr3 |
| USG000000 | 1,8836797 | 1,245608843 | 0,689097 | 2,785152 | 0,005350271 | 0,042237286 | Hydin |
| USG000000 | 6,0603558 | 1,00994677 | 0,406058 | 3,168229 | 0,001533706 | 0,015900076 | Alox5ap |
| USG000000 | 5,4361037 | 1,120487239 | 0,389191 | 3,485773 | 0,000490717 | 0,00622279 | Cd302 |
| USG000000 | 61,190884 | 1,151685193 | 0,133726 | 8,78374 | 1,58127E-18 | 2,50762E-16 | B2m |
| USG000000 | 1,3957679 | 2,677755808 | 1,011564 | 3,340567 | 0,000836076 | 0,009748081 | Dmkn |
| USG000000 | 4,0810522 | 2,538764333 | 0,671498 | 4,29078 | 1,78046E-05 | 0,000351009 | Cdh3 |
| USG000000 | 41,498029 | 1,779133111 | 0,164161 | 11,01121 | 3,3745E-28 | 1,28151E-25 | Myl1 |
| USG000000 | 9,0300582 | -1,627370595 | 0,459562 | -4,053129 | 5,05371E-05 | 0,000888308 | Cyp26b1 |
| USG000000 | 5,9957431 | 4,906091251 | 0,62496 | 8,094465 | 5,75166E-16 | 6,85968E-14 | Scgb3a1 |
| USG000000 | 404,60974 | 1,015184963 | 0,350033 | 3,486262 | 0,00048982 | 0,006217289 | mt-Atp8 |
| USG000000 | 242,7667 | 1,11093942 | 0,507459 | 2,985772 | 0,002828634 | 0,025705297 | Rny3 |
| USG000000 | 2,9494066 | 1,421071964 | 0,725603 | 2,879516 | 0,003982862 | 0,033671168 | Gm26168 |
| USG000000 | 24,192613 | 7,798597616 | 0,764797 | 10,39106 | 2,7232E-25 | 7,85969E-23 | Muc5b |
| USG000000 | 2,1297412 | 4,735491965 | 1,200585 | 4,406805 | 1,04907E-05 | 0,000224616 | Jchain |
| USG000000 | 4,4746387 | 1,010402415 | 0,465608 | 2,945459 | 0,003224762 | 0,028497577 | H2-Q10 |
| USG000000 | 41,76299 | 1,492440956 | 0,158634 | 9,604035 | 7,68746E-22 | 1,7334E-19 | Myl9 |
| USG000000 | 7,5989901 | -1,075973914 | 0,492249 | -2,946822 | 0,003210577 | 0,028389605 | Macrod2 |
| USG000000 | 4,075512 | 1,848680442 | 0,55503 | 3,903961 | 9,46311E-05 | 0,001522433 | Cgn |
| USG000000 | 51,218191 | 1,475576392 | 0,132732 | 11,29625 | 1,36948E-29 | 5,81264E-27 | Lyz2 |
| USG000000 | 2,8365423 | 1,693856048 | 0,673773 | 3,260496 | 0,001112177 | 0,012308149 | Wfdc17 |
| USG000000 | 44,778373 | 1,483434655 | 0,173226 | 8,783749 | 1,58114E-18 | 2,50762E-16 | Hba-a1 |
| USG000000 | 5,7136409 | -1,267674874 | 0,569006 | -2,982625 | 0,002857877 | 0,025865001 | Pnmal2 |
| USG000000 | 111,62311 | 1,047310475 | 0,101763 | 10,4477 | 1,50129E-25 | 4,42145E-23 | Ppia |
| USG000000 | 14,147871 | 1,064701723 | 0,253559 | 4,611482 | 3,99808E-06 | 9,48953E-05 | Slfn2 |
| USG000000 | 10,223298 | -1,128432677 | 0,438095 | -3,222503 | 0,001270759 | 0,013694044 | Acrbp |
| USG000000 | 24,834261 | 1,100610358 | 0,215757 | 5,43876 | 5,36525E-08 | 1,95475E-06 | Ifi204 |
| USG000000 | 3,972667 | -1,496873109 | 0,575472 | -3,286686 | 0,001013739 | 0,011429118 | 4932431P20Rik |
| USG000000 | 2,3804371 | -2,349426857 | 1,179928 | -2,794664 | 0,005195361 | 0,041285385 | Gm826 |
| USG000000 | 328,81008 | -1,033039627 | 0,097711 | -10,73967 | 6,62806E-27 | 2,12554E-24 | Myh7b |
| USG000000 | 1,7330669 | 1,980008819 | 0,85145 | 3,129053 | 0,001753709 | 0,017574847 | Sirpb1c |
| USG000000 | 1,5731869 | -3,628284491 | 1,443761 | -3,259234 | 0,001117136 | 0,012353551 | Fsip2 |
| USG000000 | 8,903336 | -1,012539139 | 0,500542 | -2,812584 | 0,004914523 | 0,039598818 | BC100451 |
| USG000000 | 4,4361515 | 1,670029628 | 0,441073 | 4,280685 | 1,86319E-05 | 0,000366817 | Slfn1 |
| USG000000 | 7,8654518 | 1,129397254 | 0,362216 | 3,674899 | 0,000237943 | 0,003367679 | 1810022K09Rik |
| USG000000 | 3,0918375 | 1,521499678 | 0,700924 | 3,012324 | 0,002592561 | 0,02390179 | Serpina3m |
| USG000000 | 4,4766271 | 1,214547766 | 0,432579 | 3,431715 | 0,000599778 | 0,007341306 | Ifi27l2a |
| USG000000 | 10,577535 | 1,271898166 | 0,309688 | 4,546294 | 5,45987E-06 | 0,000125464 | Ccr5 |
| USG000000 | 2,1474345 | 1,805582125 | 0,76238 | 3,141593 | 0,001680312 | 0,017072338 | Clec7a |
| USG000000 | 5,9174123 | 1,079677214 | 0,489931 | 2,989039 | 0,002798562 | 0,025480161 | Ms4a6c |

| | | | | | | | |
|---|---|---|---|---|---|---|---|
| USG000000 | 47,83996 | 1,247875007 | 0,148311 | 8,635552 | 5,84443E-18 | 8,26872E-16 | Rpl36a |
| USG000000 | 3,1061887 | -2,214720733 | 0,794471 | -3,419879 | 0,00062649 | 0,007616575 | Gm16026 |
| USG000000 | 2,9646936 | 3,207860562 | 0,762993 | 4,666824 | 3,05891E-06 | 7,49459E-05 | Muc4 |
| USG000000 | 22,023412 | 1,017535741 | 0,200958 | 5,398716 | 6,71194E-08 | 2,40945E-06 | Rpl39 |
| USG000000 | 17,785295 | -1,045723838 | 0,343472 | -3,600038 | 0,000318171 | 0,00429114 | Gm14769 |
| USG000000 | 11,301007 | -1,149324906 | 0,388252 | -3,523258 | 0,000426276 | 0,005527032 | BC002163 |
| USG000000 | 1,7849033 | -4,089346408 | 1,368584 | -3,394399 | 0,000687794 | 0,008252194 | Gm15807 |
| USG000000 | 14,437931 | -1,293764382 | 0,39163 | -3,83481 | 0,000125661 | 0,001943643 | Pisd-ps1 |
| USG000000 | 1,9623686 | -2,903292926 | 1,175958 | -3,159273 | 0,001581634 | 0,016245242 | Gm11889 |
| USG000000 | 30,076177 | 1,062394081 | 0,201615 | 5,593479 | 2,22564E-08 | 8,94658E-07 | Sbk3 |
| USG000000 | 21,87357 | 1,160163746 | 0,299929 | 4,311706 | 1,62E-05 | 0,000325601 | Gm13054 |
| USG000000 | 4,9287954 | -1,505991962 | 0,588376 | -3,230768 | 0,001234579 | 0,013385583 | E130307A14Rik |
| USG000000 | 5,8858142 | 2,765697482 | 0,464993 | 6,269801 | 3,6151E-10 | 1,9322E-08 | Cfb |
| USG000000 | 2,3106153 | 2,857969471 | 0,798339 | 4,113021 | 3,90514E-05 | 0,000707091 | Apol11b |
| USG000000 | 3,3187992 | 1,41499524 | 0,569276 | 3,213036 | 0,001313397 | 0,014018959 | Evi2b |
| USG000000 | 4,6140467 | -1,679315638 | 0,596891 | -3,448425 | 0,000563865 | 0,006978676 | Gm10718 |
| USG000000 | 5,044766 | -1,244544085 | 0,567738 | -2,970596 | 0,002972221 | 0,0266908 | Gm10717 |
| USG000000 | 1,4181118 | -3,369618976 | 1,53719 | -2,960873 | 0,003067681 | 0,027383796 | Gm10721 |
| USG000000 | 63,773598 | 1,307323579 | 0,214443 | 6,398639 | 1,56767E-10 | 8,87181E-09 | Erdr1 |
| USG000000 | 7,734808 | 1,250950658 | 0,397761 | 3,710916 | 0,000206511 | 0,00299513 | Bin2 |
| USG000000 | 1,4794529 | -3,454757001 | 1,454772 | -3,054282 | 0,002255995 | 0,021517694 | Kat6b-ps1 |
| USG000001 | 3,3108364 | -3,434032161 | 1,339331 | -3,159799 | 0,001578779 | 0,016229737 | Trpc2 |
| USG000001 | 3,2409119 | -1,465016153 | 0,800202 | -2,762412 | 0,00573761 | 0,044635824 | Gm36527 |
| USG000001 | 7,4591857 | -1,182264273 | 0,444339 | -3,302867 | 0,000957017 | 0,010891728 | Gm37334 |
| USG000001 | 2,039975 | -2,847562524 | 1,383929 | -2,937123 | 0,003312725 | 0,029147159 | RP24-560A18.3 |
| USG000001 | 7,7181749 | -1,391643627 | 0,443206 | -3,695651 | 0,000219324 | 0,003146185 | RP24-488I7.1 |
| USG000001 | 3,095684 | -4,496178299 | 1,272553 | -4,066399 | 4,77452E-05 | 0,000842863 | RP24-351E22.1 |

**CPC SEC-EVs**

| NA. | baseMean | log2FC.shrink. | lfcSE | stat | pvalue | padj | gene_symbol |
|---|---|---|---|---|---|---|---|
| USG000000 | 5,5957614 | 1,207894494 | 0,443703 | 3,42404 | 0,000616976 | 0,01685303 | Alox12 |
| USG000000 | 10,748636 | -1,001472063 | 0,414327 | -3,149133 | 0,001637558 | 0,0347048 | Snapin |
| USG000000 | 9,0796435 | 1,040109182 | 0,358543 | 3,548238 | 0,000387818 | 0,012315891 | Plag1 |
| USG000000 | 6,5545823 | -1,498280723 | 0,650748 | -3,052614 | 0,002268577 | 0,04343968 | Sla |
| USG000000 | 11,948333 | 1,61292389 | 0,302536 | 5,697875 | 1,2131E-08 | 1,84123E-06 | Ehhadh |
| USG000000 | 12,894354 | -1,172305659 | 0,46805 | -3,216187 | 0,001299059 | 0,029519311 | Ahsg |
| USG000000 | 11,547185 | 1,32165868 | 0,337843 | 4,404795 | 1,05884E-05 | 0,00066232 | Slc25a27 |
| USG000000 | 15,051073 | -1,587219556 | 0,46118 | -3,977474 | 6,96512E-05 | 0,003098578 | Mmp19 |
| USG000000 | 17,738472 | -1,035371121 | 0,363818 | -3,484209 | 0,000493593 | 0,014390886 | Tspan4 |
| USG000000 | 6,8977007 | -1,32118906 | 0,577049 | -3,055726 | 0,002245166 | 0,043117185 | Tma16 |
| USG000000 | 33,98022 | -1,027710409 | 0,237065 | -4,769714 | 1,84488E-06 | 0,000151119 | Lcn2 |
| USG000000 | 13,016436 | 1,032916903 | 0,282937 | 4,175505 | 2,97326E-05 | 0,001557716 | Hmgcs2 |
| USG000000 | 25,744386 | 1,021811406 | 0,20898 | 5,263358 | 1,41448E-07 | 1,75889E-05 | Rorc |
| USG000000 | 9,9774351 | -1,275665799 | 0,562665 | -3,038291 | 0,002379242 | 0,044409839 | Cxcl1 |
| USG000000 | 11,032921 | 1,004668792 | 0,3467 | 3,540928 | 0,000398722 | 0,012508248 | Tyk2 |
| USG000000 | 50,155834 | 1,055535049 | 0,166924 | 6,599249 | 4,13247E-11 | 9,91888E-09 | Flcn |
| USG000000 | 7,5118703 | 1,297248128 | 0,391985 | 3,893317 | 9,88829E-05 | 0,004030008 | Itga2b |
| USG000000 | 10,309856 | 1,307731325 | 0,331244 | 4,438842 | 9,0444E-06 | 0,000572683 | Bach2 |
| USG000000 | 17,564048 | -1,578971162 | 0,374195 | -4,663738 | 3,10517E-06 | 0,000235465 | D1Ertd622e |
| USG000000 | 14,195644 | -1,036891099 | 0,384086 | -3,363897 | 0,000768503 | 0,020115212 | Mapk10 |

| | | | | | | | |
|---|---|---|---|---|---|---|---|
| USG000000 | 46,842666 | 1,284395703 | 0,14428 | 9,115117 | 7,85855E-20 | 4,77107E-17 | Khnyn |
| USG000000 | 140,79239 | 1,093273376 | 0,121524 | 9,201259 | 3,53779E-20 | 2,35353E-17 | Lgals4 |
| USG000000 | 12,164152 | 1,156674104 | 0,312952 | 4,210114 | 2,55242E-05 | 0,001386501 | Pde5a |
| USG000000 | 10,030735 | -1,109539559 | 0,475476 | -3,099176 | 0,0019406 | 0,039272418 | Ado |
| USG000000 | 5,988482 | -1,721973372 | 0,662439 | -3,282831 | 0,001027703 | 0,024840579 | Pclo |
| USG000000 | 8607,414 | -1,025362899 | 0,166713 | -6,461817 | 1,03453E-10 | 2,27179E-08 | mt-Nd5 |
| USG000000 | 7,6555162 | 1,102896574 | 0,433173 | 3,29306 | 0,000991032 | 0,024180706 | Gm26109 |
| USG000000 | 41,222631 | -1,137894672 | 0,198274 | -6,059969 | 1,36148E-09 | 2,60218E-07 | Ppp1r3c |
| USG000000 | 9,9758207 | -1,069776145 | 0,467605 | -3,058097 | 0,002227472 | 0,042971469 | Gm1966 |
| USG000000 | 14,847752 | -1,186185282 | 0,396705 | -3,60927 | 0,00030706 | 0,010157585 | Fign |
| USG000000 | 5,7684936 | 1,112011527 | 0,42079 | 3,363417 | 0,000769839 | 0,020115212 | A930018M24Rik |
| USG000001 | 11,479585 | -1,168882847 | 0,462484 | -3,230611 | 0,001235259 | 0,0285215 | Gm38070 |
| USG000001 | 9,3225727 | -1,158521531 | 0,450876 | -3,277479 | 0,001047387 | 0,025198321 | RP23-458G12.2 |

**HTB SEC-EVs**

| NA. | baseMean | log2FC.shrink. | lfcSE | stat | pvalue | padj | Gene_symbol |
|---|---|---|---|---|---|---|---|
| USG00000( | 2,4018666 | 2,539239026 | 0,897221 | 3,517405 | 0,000435788 | 0,009705943 | Sept1 |
| USG00000( | 26,950555 | -1,015881458 | 0,426061 | -3,007621 | 0,002633009 | 0,039091515 | Rab5b |
| USG00000( | 17,474582 | 1,303839859 | 0,310865 | 4,636954 | 3,53582E-06 | 0,000150075 | Tcirg1 |
| USG00000( | 11,681293 | 1,038961858 | 0,3886 | 3,297184 | 0,000976594 | 0,018446631 | Ift46 |
| USG00000( | 11,735604 | 1,182373744 | 0,35118 | 3,865262 | 0,00011097 | 0,003121651 | Sdf2 |
| USG00000( | 83,572582 | 1,052793472 | 0,163648 | 6,684207 | 2,32179E-11 | 2,35956E-09 | Akap8l |
| USG00000( | 4,3341164 | 1,343346071 | 0,673658 | 2,906817 | 0,00365127 | 0,049755881 | Qtrt1 |
| USG00000( | 15,821668 | 1,211465582 | 0,330739 | 4,135403 | 3,54332E-05 | 0,001172789 | Ercc1 |
| USG00000( | 4,9311703 | 1,375588965 | 0,564376 | 3,174215 | 0,001502426 | 0,025417945 | Trmt10a |
| USG00000( | 15,719537 | 1,153976939 | 0,354633 | 3,778731 | 0,000157629 | 0,004228158 | Pnpla6 |
| USG00000( | 6,549683 | 1,269366042 | 0,496804 | 3,255326 | 0,001132624 | 0,020354785 | Stxbp2 |
| USG00000( | 14,834383 | 1,473635607 | 0,37281 | 4,398876 | 1,08813E-05 | 0,000404712 | Pias4 |
| USG00000( | 4,3240131 | 1,420972555 | 0,676539 | 2,983932 | 0,002845702 | 0,041607215 | Trpm2 |
| USG00000( | 19,123784 | 1,588175939 | 0,298566 | 5,661852 | 1,49748E-08 | 1,05415E-06 | Cdh23 |
| USG00000( | 6,6902433 | 1,195218082 | 0,496111 | 3,125784 | 0,001773315 | 0,029286986 | Tmem208 |
| USG00000( | 138,6425 | 1,192233347 | 0,125413 | 9,683979 | 3,52716E-22 | 1,21192E-19 | Lama5 |
| USG00000( | 3,4895283 | 2,014380874 | 0,684851 | 3,592109 | 0,000328013 | 0,0076219 | Pbld2 |
| USG00000( | 12,047439 | 1,621126643 | 0,352799 | 4,993951 | 5,91564E-07 | 3,0819E-05 | Best3 |
| USG00000( | 28,099122 | 1,165616066 | 0,240583 | 5,195654 | 2,04001E-07 | 1,13666E-05 | Pfas |
| USG00000( | 5,0371348 | 1,750653773 | 0,605211 | 3,54246 | 0,000396413 | 0,009051643 | Higd1b |
| USG00000( | 24,458909 | 1,002985155 | 0,278523 | 4,043198 | 5,27272E-05 | 0,001683419 | Snrnp48 |
| USG00000( | 53,044906 | -1,334497991 | 0,327185 | -4,490077 | 7,11974E-06 | 0,000279959 | Npr3 |
| USG00000( | 9,501847 | 1,407625894 | 0,435226 | 3,785991 | 0,000153097 | 0,004133828 | Nubp1 |
| USG00000( | 21,64752 | -1,476440843 | 0,502337 | -3,484122 | 0,000493755 | 0,010895073 | Ifngr2 |
| USG00000( | 5,4732872 | 1,74376295 | 0,565336 | 3,701353 | 0,000214453 | 0,005437134 | Mx2 |
| USG00000( | 15,645829 | -1,797838612 | 0,751071 | -3,064269 | 0,002182029 | 0,034152782 | Rsph3b |
| USG00000( | 53,562217 | 1,231681244 | 0,189155 | 6,773045 | 1,261E-11 | 1,31866E-09 | Sik1 |
| USG00000( | 2,7859988 | 2,316347154 | 0,803803 | 3,553262 | 0,000380486 | 0,008729397 | Kifc5b |
| USG00000( | 26,030211 | 1,008546642 | 0,27369 | 4,100946 | 4,11465E-05 | 0,001346451 | Lrrc45 |
| USG00000( | 9,1019613 | 1,210946245 | 0,404547 | 3,570061 | 0,000356898 | 0,008240621 | Dpysl4 |
| USG00000( | 10,566884 | 1,372515864 | 0,449198 | 3,642037 | 0,000270489 | 0,006549375 | Dhtkd1 |
| USG00000( | 16,736512 | 1,024739743 | 0,315391 | 3,750401 | 0,000176552 | 0,004632406 | Stk11ip |
| USG00000( | 7,63066 | 1,281064774 | 0,490435 | 3,291028 | 0,000998221 | 0,018708223 | Capn10 |
| USG00000( | 57,97842 | -1,215284436 | 0,301231 | -4,477812 | 7,5412E-06 | 0,000294924 | Elk4 |

| | | | | | | | |
|---|---|---|---|---|---|---|---|
| USG00000( | 20,536769 | -1,339930962 | 0,505932 | -3,258157 | 0,001121385 | 0,020228382 | Srp9 |
| USG00000( | 62,314729 | -1,026795449 | 0,261445 | -4,304711 | 1,67204E-05 | 0,000595783 | Lpgat1 |
| USG00000( | 11,416259 | 1,221842232 | 0,380859 | 3,754791 | 0,000173486 | 0,00457693 | Atf3 |
| USG00000( | 13,413624 | 1,225277406 | 0,343989 | 4,044582 | 5,24167E-05 | 0,001677217 | Tatdn3 |
| USG00000( | 6,129852 | 1,358691312 | 0,498965 | 3,390487 | 0,000697685 | 0,014281263 | Pter |
| USG00000( | 41,494721 | 1,073581446 | 0,218163 | 5,200663 | 1,98579E-07 | 1,11941E-05 | Neb |
| USG00000( | 49,282261 | -1,238122566 | 0,344589 | -4,052719 | 5,06257E-05 | 0,00162351 | Kpna4 |
| USG00000( | 24,730891 | -1,277675111 | 0,478576 | -3,262814 | 0,001103121 | 0,019948787 | Cisd2 |
| USG00000( | 46,420763 | -1,114618708 | 0,328295 | -3,87268 | 0,000107645 | 0,003051917 | Rps20 |
| USG00000( | 89,342906 | -1,427879979 | 0,245939 | -6,105881 | 1,02235E-09 | 8,53037E-08 | Bag1 |
| USG00000( | 19,266214 | 1,335578719 | 0,310704 | 4,705357 | 2,53423E-06 | 0,000112182 | Ccdc107 |
| USG00000( | 5,9722364 | 1,807243057 | 0,548552 | 3,87195 | 0,000107968 | 0,003055068 | Hgf |
| USG00000( | 6,4510671 | -3,375953301 | 1,228765 | -3,282252 | 0,001029817 | 0,019004207 | Angptl7 |
| USG00000( | 15,686295 | 1,133251136 | 0,334883 | 3,885651 | 0,000102056 | 0,002922164 | Miip |
| USG00000( | 3,1451526 | 1,602017467 | 0,771031 | 2,973487 | 0,002944369 | 0,042660836 | Brdt |
| USG00000( | 78,78133 | -1,003008495 | 0,246358 | -4,462618 | 8,09644E-06 | 0,00030992 | Tmed2 |
| USG00000( | 3,6426988 | 2,018479741 | 0,638755 | 3,764068 | 0,000167171 | 0,004451007 | Pop7 |
| USG00000( | 36,315535 | -1,206457591 | 0,370985 | -3,74983 | 0,000176954 | 0,004634535 | Strap |
| USG00000( | 53,743951 | 1,139266804 | 0,193758 | 6,16148 | 7,20682E-10 | 6,22764E-08 | Plekha5 |
| USG00000( | 4,3477878 | 1,718019847 | 0,648367 | 3,369617 | 0,000752726 | 0,01506566 | Lrmp |
| USG00000( | 4,3140777 | 2,503247163 | 0,596167 | 4,657743 | 3,19696E-06 | 0,000137307 | Ttll3 |
| USG00000( | 17,611304 | 1,030999491 | 0,32048 | 3,738252 | 0,000185304 | 0,00481825 | C2cd5 |
| USG00000( | 10,117437 | 1,18600214 | 0,388849 | 3,608105 | 0,000308441 | 0,007296914 | Ttyh1 |
| USG00000( | 35,6837 | -1,025126261 | 0,356136 | -3,418627 | 0,000629379 | 0,013201397 | Prkrir |
| USG00000( | 35,262939 | 1,572322331 | 0,221701 | 7,352015 | 1,95241E-13 | 2,49338E-11 | Vwa3a |
| USG00000( | 41,400905 | -1,040212281 | 0,334497 | -3,603045 | 0,000314511 | 0,007392031 | Amfr |
| USG00000( | 129,66903 | 1,030433937 | 0,136011 | 7,784054 | 7,02369E-15 | 1,10856E-12 | Slc27a1 |
| USG00000( | 12,872329 | 1,22998022 | 0,3881 | 3,709301 | 0,000207832 | 0,005308361 | Pde4c |
| USG00000( | 34,490016 | 1,003468601 | 0,236467 | 4,617117 | 3,89109E-06 | 0,000162194 | Man2c1 |
| USG00000( | 48,907656 | -1,062746011 | 0,307955 | -3,907148 | 9,3392E-05 | 0,002733752 | Tmem30a |
| USG00000( | 27,81054 | -1,120954329 | 0,479331 | -2,987841 | 0,002809555 | 0,041203955 | Atp1b3 |
| USG00000( | 5,5937065 | 1,247909218 | 0,555613 | 3,038773 | 0,002375438 | 0,036429274 | Ttf2 |
| USG00000( | 34,918762 | 1,191056991 | 0,213299 | 5,905155 | 3,52315E-09 | 2,74824E-07 | Myo5c |
| USG00000( | 2,3747229 | 1,943658056 | 0,905987 | 3,028458 | 0,002458053 | 0,03722158 | Egr3 |
| USG00000( | 28,584866 | -1,019174372 | 0,406627 | -3,099996 | 0,001935231 | 0,031379004 | Tmem127 |
| USG00000( | 3,3912945 | 1,503502669 | 0,758975 | 2,928051 | 0,003410943 | 0,047512854 | Isg15 |
| USG00000( | 23,240806 | 1,537681312 | 0,282114 | 5,787812 | 7,13089E-09 | 5,2503E-07 | Hmha1 |
| USG00000( | 2,7590728 | 3,224866523 | 0,944213 | 3,979008 | 6,92035E-05 | 0,002124841 | Rnf39 |
| USG00000( | 3,8294806 | 1,408166935 | 0,693232 | 2,9325 | 0,003362445 | 0,047064447 | Dcaf15 |
| USG00000( | 31,143316 | -1,201736889 | 0,428427 | -3,384213 | 0,000713826 | 0,014508771 | Ndufc1 |
| USG00000( | 7,209914 | 1,605380787 | 0,451338 | 4,079622 | 4,5109E-05 | 0,001462849 | Mospd3 |
| USG00000( | 2,9325469 | 1,976222234 | 0,796467 | 3,270491 | 0,001073608 | 0,019661468 | Traf3ip3 |
| USG00000( | 21,522404 | 1,160436128 | 0,282613 | 4,527167 | 5,97797E-06 | 0,000240301 | Trpm4 |
| USG00000( | 3,4401904 | 1,740937201 | 0,625083 | 3,443863 | 0,000573467 | 0,012278494 | Cdk19os |
| USG00000( | 2,0271561 | 2,213949728 | 1,039443 | 3,043643 | 0,002337325 | 0,035953302 | Susd1 |
| USG00000( | 24,724061 | -1,413608716 | 0,511826 | -3,352997 | 0,000799415 | 0,015868444 | Txndc5 |
| USG00000( | 8,4940904 | 1,052078709 | 0,435446 | 3,094247 | 0,001973132 | 0,03181482 | Naif1 |
| USG00000( | 3,8438677 | 1,828789153 | 0,665615 | 3,425481 | 0,000613713 | 0,01298605 | Arhgap9 |
| USG00000( | 100,61715 | 1,186540385 | 0,138769 | 8,736863 | 2,39673E-18 | 5,40424E-16 | Dgkz |

| | | | | | | | |
|---|---|---|---|---|---|---|---|
| USG00000( | 9,907699 | 1,357133004 | 0,423846 | 3,758399 | 0,000171004 | 0,004519703 | Mthfd1l |
| USG00000( | 15,551792 | 1,30984958 | 0,346391 | 4,244086 | 2,19486E-05 | 0,00076323 | Plekhh2 |
| USG00000( | 17,603189 | 1,07147543 | 0,300545 | 4,023801 | 5,72663E-05 | 0,001812302 | Crocc |
| USG00000( | 10,018574 | 2,30111846 | 0,413226 | 5,909901 | 3,42314E-09 | 2,68474E-07 | Kif21b |
| USG00000( | 3,3716624 | 1,471642771 | 0,640562 | 3,100718 | 0,00193052 | 0,03133784 | Clspn |
| USG00000( | 12,990242 | 1,02597762 | 0,343756 | 3,51738 | 0,000435829 | 0,009705943 | Zfp513 |
| USG00000( | 8,6801551 | 1,146048434 | 0,541043 | 2,91531 | 0,003553352 | 0,048883152 | Gm8399 |
| USG00000( | 80,772303 | 1,054773039 | 0,158303 | 6,896509 | 5,32959E-12 | 5,78281E-10 | Lincpint |
| USG00000( | 4,2717669 | 2,379619644 | 0,594376 | 4,473354 | 7,70023E-06 | 0,00029952 | Gm6430 |
| USG00000( | 51,051336 | -1,12190082 | 0,294025 | -4,238403 | 2,25115E-05 | 0,000780922 | Mylk4 |
| USG00000( | 17,774817 | 1,041725019 | 0,366864 | 3,419686 | 0,000626934 | 0,01316927 | 1110034G24Rik |
| USG00000( | 103,37543 | -1,280238783 | 0,2448 | -5,548504 | 2,88124E-08 | 1,89854E-06 | Rps8 |
| USG00000( | 18,975842 | 1,492771623 | 0,306269 | 5,229715 | 1,69771E-07 | 9,68368E-06 | Atg16l2 |
| USG00000( | 5,4764469 | -2,750369967 | 1,232093 | -3,009403 | 0,002617618 | 0,039028315 | Zbtb33 |
| USG00000( | 4,6042139 | 1,618290764 | 0,597168 | 3,399001 | 0,000676325 | 0,013907092 | Gm9833 |
| USG00000( | 6,3856275 | 1,863066111 | 0,508222 | 4,179188 | 2,92552E-05 | 0,000991038 | Rasd1 |
| USG00000( | 5,7130607 | -2,846288775 | 1,221874 | -3,059315 | 0,002218438 | 0,034610036 | C77080 |
| USG00000( | 25,721041 | 1,023034027 | 0,253799 | 4,427366 | 9,5391E-06 | 0,000356629 | Tbc1d22a |
| USG00000( | 23,509335 | 1,338335177 | 0,291539 | 4,973165 | 6,58686E-07 | 3,35883E-05 | Sec31b |
| USG00000( | 10,29605 | 1,487262472 | 0,419367 | 4,061291 | 4,88021E-05 | 0,001572017 | Tchh |
| USG00000( | 43,302922 | 1,849812175 | 0,190083 | 9,893723 | 4,43231E-23 | 1,59907E-20 | Dancr |
| USG00000( | 29,040733 | -1,054999031 | 0,396271 | -3,240194 | 0,001194484 | 0,021176405 | Spcs3 |
| USG00000( | 5,658316 | 1,362204998 | 0,648576 | 2,931739 | 0,003370695 | 0,047134203 | H2-Q5 |
| USG00000( | 19,652224 | -1,05669913 | 0,468821 | -2,916203 | 0,003543196 | 0,048836548 | St3gal5 |
| USG00000( | 3,052101 | 1,749325298 | 0,68405 | 3,285892 | 0,001016598 | 0,018880986 | C330011M18Rik |
| USG00000( | 6,1072154 | 3,384414846 | 1,287198 | 3,343519 | 0,000827229 | 0,016352238 | Serpina3k |
| USG00000( | 126,94752 | 1,028078338 | 0,126809 | 8,301268 | 1,03006E-16 | 1,93049E-14 | Dot1l |
| USG00000( | 6,3769215 | 1,460086514 | 0,533853 | 3,420884 | 0,00062418 | 0,013130526 | Athl1 |
| USG00000( | 5,068327 | -3,226163641 | 1,231356 | -3,211 | 0,001322738 | 0,022998105 | Gm10115 |
| USG00000( | 180,86918 | 1,025164284 | 0,128852 | 8,154662 | 3,50158E-16 | 6,23843E-14 | Snrnp70 |
| USG00000( | 5,6138198 | 2,361777784 | 0,552377 | 4,729218 | 2,25387E-06 | 0,000101642 | Ccdc146 |
| USG00000( | 10,857079 | 3,564261114 | 0,413172 | 8,849792 | 8,76825E-19 | 2,04088E-16 | Gm26448 |
| USG00000( | 16,734542 | 2,372792206 | 0,330319 | 7,457409 | 8,82405E-14 | 1,20132E-11 | Gm26447 |
| USG00000( | 17,683451 | 2,799415547 | 0,286839 | 9,955545 | 2,38512E-23 | 9,05781E-21 | Snora73a |
| USG00000( | 14,664547 | 2,14175505 | 0,348199 | 6,457459 | 1,06476E-10 | 1,00428E-08 | Gm23928 |
| USG00000( | 21,652257 | 2,490076884 | 0,307631 | 8,329961 | 8,0869E-17 | 1,53555E-14 | Snora23 |
| USG00000( | 11,589533 | 2,463640511 | 0,347389 | 7,380114 | 1,58154E-13 | 2,05614E-11 | n-R5s56 |
| USG00000( | 4,4895612 | 2,147286697 | 0,802289 | 3,39941 | 0,000675313 | 0,013907092 | Gm22457 |
| USG00000( | 5,6032839 | 2,866467458 | 1,168398 | 3,247507 | 0,001164207 | 0,020741563 | Gm22306 |
| USG00000( | 2,4305136 | 2,316923358 | 0,857598 | 3,426963 | 0,000610372 | 0,012934322 | Gm22620 |
| USG00000( | 9,5365568 | 1,895183961 | 0,425524 | 4,873839 | 1,0945E-06 | 5,24742E-05 | Gm23301 |
| USG00000( | 63,336266 | 1,431972456 | 0,310579 | 4,989243 | 6,06164E-07 | 3,13532E-05 | Gm24146 |
| USG00000( | 3,3455244 | 4,353323899 | 0,841954 | 5,548938 | 2,8741E-08 | 1,89854E-06 | Gm24148 |
| USG00000( | 2,4898742 | 1,984157107 | 0,827112 | 3,203918 | 0,001355709 | 0,023402198 | n-R5s62 |
| USG00000( | 3,5171546 | 2,113786566 | 0,685117 | 3,703022 | 0,000213047 | 0,005412807 | Snord96a |
| USG00000( | 2,4631803 | 2,210109786 | 0,92222 | 3,2044 | 0,001353443 | 0,023391061 | Gm24708 |
| USG00000( | 13,248679 | 1,620179667 | 0,360655 | 4,900176 | 9,57508E-07 | 4,66817E-05 | Gm26330 |
| USG00000( | 84,531519 | 2,509619241 | 0,342003 | 7,600148 | 2,95791E-14 | 4,2263E-12 | n-R5s193 |
| USG00000( | 3,4089334 | 3,089394331 | 1,482242 | 3,027972 | 0,00246201 | 0,037227156 | Gm26205 |

| | | | | | | | |
|---|---|---|---|---|---|---|---|
| USG00000( | 2,9267272 | 2,471746499 | 0,810808 | 3,694358 | 0,000220443 | 0,005532552 | n-R5s25 |
| USG00000( | 1,3265566 | 3,478803339 | 1,527931 | 3,101019 | 0,001928562 | 0,03133784 | Snora43 |
| USG00000( | 10,463354 | 1,29354242 | 0,462277 | 3,422048 | 0,000621514 | 0,013112291 | Snord58b |
| USG00000( | 2,8492778 | 4,27569691 | 0,915424 | 5,077805 | 3,8182E-07 | 2,02575E-05 | Gm22042 |
| USG00000( | 5,215525 | 3,426854193 | 0,624548 | 5,843835 | 5,10127E-09 | 3,83419E-07 | Snora61 |
| USG00000( | 253,21406 | 1,863120838 | 0,144396 | 13,0526 | 6,14128E-39 | 6,81729E-36 | Snord15b |
| USG00000( | 7,2953637 | 3,42688076 | 0,591344 | 6,130904 | 8,73812E-10 | 7,46152E-08 | Snora70 |
| USG00000( | 19,960078 | 2,51869224 | 0,300992 | 8,597181 | 8,16979E-18 | 1,70867E-15 | Gm22422 |
| USG00000( | 8,4156975 | 1,760526155 | 0,442267 | 4,453849 | 8,43442E-06 | 0,000319468 | Gm26035 |
| USG00000( | 41,533303 | 2,762983728 | 0,25358 | 11,07229 | 1,70969E-28 | 9,86903E-26 | Gm23511 |
| USG00000( | 4,9642693 | 3,173907014 | 0,596355 | 5,680197 | 1,34539E-08 | 9,56423E-07 | Gm22362 |
| USG00000( | 13,09862 | 1,547126294 | 0,754016 | 2,937803 | 0,003305473 | 0,046627436 | Gm25704 |
| USG00000( | 48,962775 | 1,927268881 | 0,178041 | 10,99249 | 4,15308E-28 | 2,30512E-25 | Gm22884 |
| USG00000( | 316,26312 | 1,809507638 | 0,093544 | 19,43763 | 3,70913E-84 | 1,78422E-80 | Snord22 |
| USG00000( | 24,731277 | 2,258923964 | 1,198463 | 2,956729 | 0,003109216 | 0,044601485 | Snora16a |
| USG00000( | 1,7007733 | 3,799494429 | 1,310058 | 3,566917 | 0,000361206 | 0,008326782 | Gm26132 |
| USG00000( | 2,9816664 | 1,81743935 | 0,740569 | 3,228873 | 0,001242793 | 0,021841481 | n-R5s50 |
| USG00000( | 6,3688914 | 2,585244573 | 1,038543 | 3,265996 | 0,001090798 | 0,019875381 | Snora65 |
| USG00000( | 142,51418 | 1,610502221 | 0,167399 | 9,806756 | 1,053E-22 | 3,70629E-20 | Rnu12 |
| USG00000( | 1,677638 | 3,389088376 | 1,112424 | 3,674392 | 0,000238417 | 0,005891424 | Gm26071 |
| USG00000( | 16,814961 | 2,215495615 | 0,28535 | 8,012893 | 1,12041E-15 | 1,92484E-13 | Gm24616 |
| USG00000( | 2,5852569 | 2,102064089 | 0,971748 | 3,044847 | 0,002327988 | 0,035854002 | Gm25791 |
| USG00000( | 2,6462121 | 2,079729984 | 1,010786 | 3,012389 | 0,002592003 | 0,038721739 | Gm23969 |
| USG00000( | 10,103545 | 2,295377116 | 0,389951 | 6,212083 | 5,22866E-10 | 4,62913E-08 | Snora30 |
| USG00000( | 7,1614716 | 1,365593825 | 0,570952 | 3,154605 | 0,001607153 | 0,026937085 | Gm25128 |
| USG00000( | 7,163762 | 3,399336706 | 0,531383 | 6,706224 | 1,99724E-11 | 2,05873E-09 | Gm24455 |
| USG00000( | 7,1734736 | 2,068598096 | 0,491514 | 4,660368 | 3,15645E-06 | 0,000135972 | Gm23245 |
| USG00000( | 4,2318124 | 3,076268111 | 0,617514 | 5,373969 | 7,7022E-08 | 4,72981E-06 | Gm22058 |
| USG00000( | 1,7630326 | 4,368110555 | 1,558725 | 3,524504 | 0,000424277 | 0,009536982 | Gm24927 |
| USG00000( | 13,512706 | 1,495251057 | 0,356726 | 4,624083 | 3,7626E-06 | 0,000158303 | Snora34 |
| USG00000( | 7,6674837 | 1,591330248 | 0,491971 | 3,808873 | 0,000139602 | 0,003808301 | Gm25894 |
| USG00000( | 55,736803 | 1,536642873 | 0,256606 | 6,285979 | 3,25795E-10 | 2,92021E-08 | Snord33 |
| USG00000( | 27,232737 | 2,210307632 | 0,23754 | 9,508703 | 1,93054E-21 | 5,92759E-19 | Gm26397 |
| USG00000( | 5,9616113 | 2,081128818 | 0,507457 | 4,565516 | 4,98267E-06 | 0,000202549 | Snora69 |
| USG00000( | 20,464792 | 1,978710721 | 0,880862 | 3,091421 | 0,00199201 | 0,032047601 | Snora74a |
| USG00000( | 2,8181401 | 2,444583909 | 0,981546 | 3,266294 | 0,001089649 | 0,019875381 | Snora5c |
| USG00000( | 5,0693753 | 3,255736145 | 0,52565 | 6,50977 | 7,5266E-11 | 7,21425E-09 | Gm26165 |
| USG00000( | 50,38806 | 1,560175012 | 0,222536 | 7,26403 | 3,75724E-13 | 4,59497E-11 | Gm24494 |
| USG00000( | 4,0989388 | 2,236975452 | 0,729507 | 3,686587 | 0,000227282 | 0,005675957 | Gm23344 |
| USG00000( | 64,894462 | 1,72753306 | 0,186578 | 9,454421 | 3,24815E-21 | 9,76543E-19 | Snord15a |
| USG00000( | 15,598311 | 1,767201564 | 0,30326 | 6,143733 | 8,06041E-10 | 6,9238E-08 | Gm26315 |
| USG00000( | 5,6913233 | 2,498902724 | 0,666378 | 4,255356 | 2,08717E-05 | 0,000731066 | Gm24029 |
| USG00000( | 12,607635 | 1,860688294 | 0,436488 | 4,702452 | 2,57055E-06 | 0,000113442 | Gm25682 |
| USG00000( | 2,2175893 | 2,28518024 | 0,842613 | 3,417455 | 0,000632096 | 0,013239162 | n-R5s151 |
| USG00000( | 557,57571 | 1,868621682 | 0,245952 | 7,854018 | 4,02917E-15 | 6,46056E-13 | Gm24447 |
| USG00000( | 29,085617 | 1,762745916 | 0,23532 | 7,729801 | 1,07714E-14 | 1,67143E-12 | n-R5-8s1 |
| USG00000( | 3,150183 | 2,108002772 | 0,781147 | 3,406969 | 0,000656885 | 0,013639585 | Snora2b |
| USG00000( | 8,1857779 | 1,240137731 | 0,499042 | 3,177921 | 0,00148335 | 0,025183801 | Gm6265 |
| USG00000( | 2,1684091 | 1,938571619 | 0,957996 | 2,985636 | 0,002829894 | 0,041460108 | Gm10167 |

| | | | | | | | |
|---|---|---|---|---|---|---|---|
| USG00000( | 10,089631 | 1,325631727 | 0,419268 | 3,725348 | 0,000195046 | 0,005053327 | Syngap1 |
| USG00000( | 33,520782 | -1,26444603 | 0,483329 | -3,221303 | 0,001276093 | 0,022348656 | Mfap1a |
| USG00000( | 7,3176398 | 2,151075469 | 0,493532 | 4,792722 | 1,64534E-06 | 7,56175E-05 | Snora33 |
| USG00000( | 1,5393311 | 4,574926616 | 1,401511 | 3,823573 | 0,000131532 | 0,003615491 | Snora57 |
| USG00000( | 2,471955 | 2,574111747 | 0,99121 | 3,328434 | 0,000873357 | 0,017008656 | n-R5s71 |
| USG00000( | 9,9715866 | 1,079643292 | 0,401551 | 3,304541 | 0,00095132 | 0,018016396 | Gm20634 |
| USG00000( | 7,052958 | 1,108315281 | 0,465449 | 3,089884 | 0,002002349 | 0,032178058 | Sars2 |
| USG00000( | 3,9364812 | 2,006926259 | 0,692956 | 3,572297 | 0,000353864 | 0,008183667 | Ccdc173 |
| USG00000( | 12,691123 | 1,048141002 | 0,401612 | 3,23407 | 0,001220395 | 0,021582742 | Eml3 |
| USG00000( | 34,655425 | -1,616113637 | 0,474787 | -3,903819 | 9,46866E-05 | 0,002754883 | Tuba1a |
| USG00000( | 41,303086 | -1,015854971 | 0,336674 | -3,519479 | 0,000432395 | 0,009661055 | Smim10l1 |
| USG00000( | 4,8308366 | 1,187222877 | 0,578426 | 2,909737 | 0,003617332 | 0,049386683 | 9330160F10Rik |
| USG00000( | 28,688006 | 1,172963352 | 0,265528 | 4,797632 | 1,60552E-06 | 7,42605E-05 | Grk6 |
| USG00000( | 2,3654362 | 2,334452067 | 0,97077 | 3,189288 | 0,001426237 | 0,024415221 | Gm22589 |
| USG00000( | 1,3748512 | 5,820209154 | 1,60305 | 3,819843 | 0,000133537 | 0,003663624 | Igkv5-39 |
| USG00000( | 5,0951595 | 2,153600793 | 0,623473 | 4,010888 | 6,04908E-05 | 0,001899304 | Snora17 |
| USG00000( | 19,341345 | 2,002445853 | 0,289876 | 7,169661 | 7,51839E-13 | 8,82097E-11 | Gm22270 |
| USG00000( | 11,760656 | 1,951468224 | 0,453324 | 4,742321 | 2,11284E-06 | 9,55811E-05 | Snord66 |
| USG00000( | 12,325251 | 2,031569087 | 0,378237 | 5,7099 | 1,13043E-08 | 8,15661E-07 | Gm26079 |
| USG00000( | 29,421079 | 1,635910243 | 0,255187 | 6,692007 | 2,2013E-11 | 2,25297E-09 | Gm23946 |
| USG00000( | 31,797785 | 1,867190755 | 0,260025 | 7,437185 | 1,02853E-13 | 1,37433E-11 | Snora64 |
| USG00000( | 87,32492 | 1,715316097 | 0,149613 | 11,627 | 3,00464E-31 | 2,40888E-28 | Snord17 |
| USG00000( | 36,946857 | -1,285026611 | 0,398589 | -3,740006 | 0,000184016 | 0,00479339 | Psme3 |
| USG00000( | 45,91242 | -1,092107146 | 0,306073 | -4,008953 | 6,09885E-05 | 0,001909164 | Gm14420 |
| USG00000( | 55,939642 | 1,238710451 | 0,187898 | 6,849164 | 7,4283E-12 | 7,94058E-10 | Phyhd1 |
| USG00000( | 6,6354969 | 1,494411348 | 0,494478 | 3,629012 | 0,000284508 | 0,006808842 | Gm14239 |
| USG00000( | 2,7904057 | 1,576214207 | 0,744818 | 3,005665 | 0,002650007 | 0,03926309 | Gm25776 |
| USG00000( | 2,004943 | 4,724648794 | 1,335937 | 4,081436 | 4,47583E-05 | 0,001454746 | Gm22488 |
| USG00000( | 2,3707983 | 2,235750581 | 0,976632 | 3,135929 | 0,001713107 | 0,028383296 | Gm22193 |
| USG00000( | 6,0885099 | 1,4298609 | 0,523932 | 3,398924 | 0,000676514 | 0,013907092 | Gm23991 |
| USG00000( | 3,9044713 | 1,575212845 | 0,63535 | 3,231109 | 0,001233111 | 0,021701243 | Gm11408 |
| USG00000( | 2,8563972 | 1,805411914 | 0,785104 | 3,113735 | 0,001847352 | 0,030294481 | Gm15538 |
| USG00000( | 5,7936145 | -2,579685006 | 1,250544 | -2,929372 | 0,003396479 | 0,047402893 | Tpt1-ps3 |
| USG00000( | 4,457895 | 2,492162818 | 0,59121 | 4,669358 | 3,02143E-06 | 0,000130546 | n-R5s155 |
| USG00000( | 215,04837 | 1,496748825 | 0,164691 | 9,282979 | 1,64805E-20 | 4,57365E-18 | Gm22988 |
| USG00000( | 43,830307 | 1,321507579 | 0,21269 | 6,490523 | 8,55388E-11 | 8,12112E-09 | Gm25291 |
| USG00000( | 6,0726427 | 1,373815442 | 0,529465 | 3,296591 | 0,000978659 | 0,018461483 | Ino80dos |
| USG00000( | 14,52408 | 1,125091355 | 0,331054 | 3,884113 | 0,000102704 | 0,002923323 | Snhg17 |
| USG00000( | 3,0697828 | 2,245261787 | 0,801354 | 3,496198 | 0,000471938 | 0,010461661 | Gm13270 |
| USG00000( | 1,7154705 | 3,694389057 | 1,371471 | 3,381142 | 0,000721853 | 0,014630695 | Gm14236 |
| USG00000( | 1,6185433 | 3,428517163 | 1,069897 | 3,785738 | 0,000153253 | 0,004133828 | Rprl2 |
| USG00000( | 11,865849 | 1,775647325 | 0,421085 | 4,651756 | 3,2912E-06 | 0,000140519 | Gm25117 |
| USG00000( | 163,00818 | 2,802187284 | 0,173558 | 16,27391 | 1,51176E-59 | 2,72703E-56 | Snora81 |
| USG00000( | 27,932388 | 1,785370874 | 0,254764 | 7,268813 | 3,6266E-13 | 4,47311E-11 | Gm25395 |
| USG00000( | 382,2649 | 1,74434324 | 0,2406 | 7,495763 | 6,59136E-14 | 9,14614E-12 | Rmrp |
| USG00000( | 2,3382032 | 1,852068794 | 0,918253 | 2,950543 | 0,003172162 | 0,045324228 | Scarna3b |
| USG00000( | 94,867905 | 2,012064661 | 0,179542 | 11,36893 | 5,97175E-30 | 3,91719E-27 | Scarna2 |
| USG00000( | 2,5549591 | 3,547685497 | 0,8701 | 4,549688 | 5,37256E-06 | 0,000217785 | Gm23751 |
| USG00000( | 3,7053276 | 3,058258909 | 0,887684 | 4,007257 | 6,14281E-05 | 0,001914619 | Gm24016 |

| | | | | | | | |
|---|---|---|---|---|---|---|---|
| USG00000( | 40,594529 | 1,598556538 | 0,381855 | 4,618164 | 3,87149E-06 | 0,000161941 | Scarna13 |
| USG00000( | 6,2569768 | 1,073422422 | 0,508676 | 2,91461 | 0,003561337 | 0,048905917 | Gm24727 |
| USG00000( | 3,8474741 | 3,079166385 | 0,749418 | 4,583502 | 4,57252E-06 | 0,000186401 | Snord11 |
| USG00000( | 5,8142637 | 1,871627255 | 0,542644 | 3,995338 | 6,4602E-05 | 0,002004886 | Gm24130 |
| USG00000( | 3,9334262 | 2,877880512 | 0,691778 | 4,627339 | 3,70394E-06 | 0,000156291 | Gm24679 |
| USG00000( | 80,869305 | 1,369287555 | 0,162552 | 8,632208 | 6,01786E-18 | 1,29618E-15 | Gm25835 |
| USG00000( | 14,076777 | 1,250118755 | 0,355856 | 3,989863 | 6,61115E-05 | 0,00204733 | Gp49a |
| USG00000( | 16,870213 | 1,590469029 | 0,309303 | 5,496411 | 3,87598E-08 | 2,51956E-06 | Gm8113 |
| USG00000( | 2,2893827 | 2,152109413 | 0,780141 | 3,451421 | 0,000557643 | 0,012028908 | 4930524O07Rik |
| USG00000( | 10,707823 | 1,188440985 | 0,393151 | 3,59129 | 0,000329045 | 0,0076219 | Gm16340 |
| USG00000( | 12,843916 | 3,039267414 | 0,337565 | 9,21886 | 3,00275E-20 | 7,87866E-18 | Gm17300 |
| USG00000( | 9,4320241 | 1,21159181 | 0,425946 | 3,45427 | 0,000551784 | 0,011956153 | Mpc1-ps |
| USG00000( | 7,0672223 | 1,431849165 | 0,506011 | 3,479466 | 0,000502413 | 0,011035506 | Gm17229 |
| USG00000( | 9,6597371 | 2,004361044 | 0,437231 | 4,991628 | 5,98725E-07 | 3,10799E-05 | Gm24105 |
| USG00000( | 3,3633756 | 2,321285227 | 1,015922 | 3,133544 | 0,001727087 | 0,028582108 | Gm25744 |
| USG00000( | 3,2300046 | 1,678432728 | 0,800239 | 2,983324 | 0,00285136 | 0,041647744 | Gm24514 |
| USG00000( | 662,83685 | 1,806831091 | 0,356783 | 5,430058 | 5,63356E-08 | 3,50422E-06 | Rn7s6 |
| USG00000( | 3,1476332 | 2,127475952 | 0,71738 | 3,621216 | 0,000293222 | 0,00697115 | Gm26197 |
| USG00000( | 5,4685903 | 3,32007766 | 0,648219 | 5,49421 | 3,92464E-08 | 2,53587E-06 | Gm23639 |
| USG00000( | 1604,561 | 1,03180795 | 0,282426 | 4,10521 | 4,03948E-05 | 0,001324857 | Rpph1 |
| USG00000( | 8,8674861 | 1,427545672 | 0,577307 | 3,196442 | 0,001391336 | 0,023957203 | Gm23806 |
| USG00000( | 5,6147063 | 1,543966612 | 0,575291 | 3,376769 | 0,000733425 | 0,014782201 | Snord8 |
| USG00000( | 22,07709 | 1,956929197 | 0,313019 | 6,549937 | 5,75614E-11 | 5,72875E-09 | Gm22701 |
| USG00000( | 2,7824528 | 4,067704774 | 0,850224 | 5,187959 | 2,12611E-07 | 1,17555E-05 | Gm23799 |
| USG00000( | 4,9390024 | 2,606758307 | 0,592913 | 4,840001 | 1,29839E-06 | 6,18384E-05 | Snora15 |
| USG00000( | 3,9272132 | 3,061891463 | 0,711846 | 4,756309 | 1,97165E-06 | 9,00405E-05 | Gm26505 |
| USG00000( | 223,69999 | 2,821005925 | 0,316771 | 9,115474 | 7,83276E-20 | 1,88391E-17 | Gm24312 |
| USG00000( | 3,2973704 | 1,718824632 | 0,85494 | 2,93698 | 0,003314255 | 0,046661479 | Gm23966 |
| USG00000( | 320,61514 | 3,086946802 | 0,679444 | 4,966183 | 6,82834E-07 | 3,46971E-05 | Gm24924 |
| USG00000( | 454,90408 | 2,370046156 | 0,493768 | 5,196288 | 2,03307E-07 | 1,13666E-05 | Gm24407 |
| USG00000( | 11,836479 | -1,608734172 | 0,707226 | -2,988275 | 0,002805571 | 0,041187385 | Gm21769 |
| USG00000( | 9,8247398 | 2,966038189 | 0,400282 | 7,674019 | 1,66689E-14 | 2,50572E-12 | Gm25360 |
| USG00000( | 2,9882243 | 2,472052091 | 0,855551 | 3,561622 | 0,000368571 | 0,008483013 | Gm24873 |
| USG00000( | 198,86427 | 3,53810153 | 0,728312 | 5,256807 | 1,46578E-07 | 8,46108E-06 | Gm22614 |
| USG00000( | 7,3728345 | 2,770614064 | 0,60796 | 4,974963 | 6,52602E-07 | 3,35149E-05 | n-R5s194 |
| USG00000( | 12,08165 | 1,069812655 | 0,38657 | 3,370726 | 0,000749705 | 0,015047273 | Tubb4b-ps1 |
| USG00000( | 3,3495168 | 2,162626325 | 0,842529 | 3,300948 | 0,000963587 | 0,018224807 | Gm25380 |
| USG00000( | 8,8334499 | 2,270231451 | 0,431116 | 5,628008 | 1,82303E-08 | 1,24095E-06 | n-R5s112 |
| USG00000( | 19,091371 | 2,089440015 | 0,277339 | 7,783272 | 7,06724E-15 | 1,10856E-12 | Gm24305 |
| USG00000( | 6,1289922 | 2,566224766 | 0,501837 | 5,49369 | 3,93621E-08 | 2,53587E-06 | n-R5s136 |
| USG00000( | 22,317128 | 2,504345626 | 0,619588 | 4,513792 | 6,36786E-06 | 0,000254582 | Gm25818 |
| USG00000( | 557,76811 | 1,721402828 | 0,480949 | 4,092277 | 4,27159E-05 | 0,001391496 | Gm24265 |
| USG00000( | 13,381027 | 2,800895004 | 0,379501 | 7,639086 | 2,18769E-14 | 3,15705E-12 | Gm23686 |
| USG00000( | 3,3380478 | 2,33423401 | 0,790193 | 3,606322 | 0,000310568 | 0,007335197 | n-R5s102 |
| USG00000( | 6,6092463 | 1,090406818 | 0,526186 | 2,914409 | 0,003563629 | 0,048905917 | Gm17690 |
| USG00000( | 1,6251815 | 4,334552953 | 1,405496 | 3,677802 | 0,000235252 | 0,005840466 | Gm26674 |
| USG00000( | 101,59187 | 1,646520689 | 0,184102 | 9,142152 | 6,12215E-20 | 1,49744E-17 | Snhg4 |
| USG00000( | 30295,175 | 1,688169137 | 0,200868 | 8,664937 | 4,51763E-18 | 1,00298E-15 | Gm26917 |
| USG00000( | 22,498701 | 1,369143706 | 0,607684 | 3,035006 | 0,002405308 | 0,036770121 | Gm28043 |

| | | | | | | | |
|---|---|---|---|---|---|---|---|
| USG00000( | 119334,88 | 1,958447208 | 0,381831 | 5,48798 | 4,06557E-08 | 2,59603E-06 | Rn7s1 |
| USG00000( | 3,0309689 | 2,818224876 | 0,774711 | 4,168177 | 3,07046E-05 | 0,001032862 | Gm28872 |
| USG00000 | 180,15487 | -1,104354024 | 0,161589 | -7,065392 | 1,60163E-12 | 1,83437E-10 | Kcnq1ot1 |
| USG00000 | 26,726643 | -1,074252645 | 0,386372 | -3,333954 | 0,000856208 | 0,016780305 | 2900097C17Rik |
| USG00000 | 133,46353 | 1,360153073 | 0,12937 | 10,67976 | 1,26597E-26 | 6,29972E-24 | Dst |
| USG00000 | 6,5292663 | -3,451576755 | 1,234101 | -3,304664 | 0,000950904 | 0,018016396 | Gm37296 |
| USG00000 | 58,363965 | -1,008426599 | 0,249579 | -4,425245 | 9,6333E-06 | 0,00035922 | RP23-407I6.6 |
| USG00000 | 5,4910561 | -2,698081908 | 1,22913 | -2,992786 | 0,002764436 | 0,040749306 | RP23-458G12.1 |
| USG00000 | 139027,1 | 1,252213939 | 0,577311 | 2,95325 | 0,003144471 | 0,045017712 | Rn18s-rs5 |
| USG00000 | 1,3922839 | 4,057519462 | 1,769969 | 3,106427 | 0,001893629 | 0,030912844 | SCARNA15 |
| USG00000 | 18,807956 | 2,459426208 | 0,300718 | 8,398683 | 4,51513E-17 | 9,04971E-15 | Snora26 |
| USG00000 | 10,754989 | 1,014834574 | 0,428634 | 3,055174 | 0,002249304 | 0,03495211 | RP23-149A1.2 |
| USG00000 | 4,7403376 | -2,695184123 | 1,240292 | -2,981143 | 0,002871748 | 0,041776408 | RP24-122N24.2 |
| | | | | **HUVEC SEC-EVs** | | | |
| USG000000 | 4,6407187 | 2,473460557 | 0,590782 | 4,650269 | 3,31503E-06 | 0,000254944 | Nmrk2 |
| USG000000 | 16,839854 | 1,341204141 | 0,286806 | 5,081603 | 3,74263E-07 | 3,98533E-05 | Dnajb1 |
| USG000000 | 14,523247 | 1,020816126 | 0,314681 | 3,790257 | 0,000150492 | 0,006218674 | Vac14 |
| USG000000 | 301,96488 | 1,134113561 | 0,101323 | 11,35611 | 6,91542E-30 | 6,83787E-27 | Ctgf |
| USG000000 | 59,193383 | 1,304576791 | 0,177248 | 7,614853 | 2,63992E-14 | 8,70106E-12 | Btg2 |
| USG000000 | 13,862832 | 1,070627645 | 0,318172 | 3,904837 | 9,4289E-05 | 0,004224085 | Myo19 |
| USG000000 | 1,7550396 | 2,215209387 | 1,062178 | 3,036169 | 0,002396052 | 0,049768432 | Cd180 |
| USG000000 | 2,8519799 | 1,570213314 | 0,667358 | 3,167341 | 0,001538399 | 0,035791699 | Gm5457 |
| USG000000 | 57,563845 | 1,008378103 | 0,186484 | 5,735744 | 9,70851E-09 | 1,52722E-06 | Dusp1 |
| USG000000 | 4,5574763 | 1,395251145 | 0,54046 | 3,326291 | 0,0008801 | 0,02398273 | Zfp511 |
| USG000000 | 40,71138 | 1,08013215 | 0,202995 | 5,683026 | 1,32332E-08 | 1,99116E-06 | Spta1 |
| USG000000 | 28,593547 | -1,351210387 | 0,325734 | -4,573079 | 4,80609E-06 | 0,000345157 | Cenpf |
| USG000000 | 91,010019 | 1,444802732 | 0,141862 | 10,37116 | 3,35438E-25 | 2,90217E-22 | Cyr61 |
| USG000000 | 80,824101 | 1,42770033 | 0,141742 | 10,23547 | 1,3752E-24 | 1,05761E-21 | Tpm2 |
| USG000000 | 3,7696129 | 2,437638383 | 0,774449 | 3,777439 | 0,000158449 | 0,006470243 | Myh2 |
| USG000000 | 7,2398372 | 2,550177159 | 0,844812 | 3,672111 | 0,000240555 | 0,008871832 | Fgb |
| USG000000 | 51864,995 | 1,709824644 | 0,741387 | 3,141169 | 0,001682752 | 0,038439492 | Lars2 |
| USG000000 | 3,5534221 | 1,777603484 | 0,767031 | 3,171636 | 0,001515829 | 0,035385544 | Shq1 |
| USG000000 | 15,332504 | 1,042129716 | 0,357657 | 3,530398 | 0,000414935 | 0,013357997 | Zfp692 |
| USG000000 | 4,00159 | 1,93183962 | 0,584846 | 3,8935 | 9,88083E-05 | 0,004308955 | Tacc3 |
| USG000000 | 15,02021 | 1,9952601 | 0,294728 | 7,055953 | 1,71422E-12 | 4,31453E-10 | Nov |
| USG000000 | 85,836972 | 1,044712304 | 0,137518 | 7,848986 | 4,19414E-15 | 1,56918E-12 | Serpine1 |
| USG000000 | 86,297439 | 1,680754916 | 0,124498 | 13,6425 | 2,23763E-42 | 5,16259E-39 | Egr1 |
| USG000000 | 17,814214 | -1,039668385 | 0,445399 | -3,044851 | 0,002327958 | 0,048901256 | C130074G19Rik |
| USG000000 | 9,0799067 | 1,123518172 | 0,402011 | 3,450991 | 0,000558532 | 0,016992888 | Hlx |
| USG000000 | 16,798452 | 1,244151548 | 0,323587 | 4,330492 | 1,48777E-05 | 0,000919426 | Wdr70 |
| USG000000 | 45,266204 | 1,227709789 | 0,210693 | 6,139872 | 8,25881E-10 | 1,56612E-07 | Ppp1r15a |
| USG000000 | 10,720155 | 1,713154935 | 0,385957 | 4,877271 | 1,07564E-06 | 9,79609E-05 | Mfap4 |
| USG000000 | 4,8992507 | 1,333502953 | 0,573177 | 3,161401 | 0,001570124 | 0,036407405 | Ttc41 |
| USG000000 | 5,5707615 | -3,68858834 | 1,219834 | -3,45904 | 0,000542104 | 0,016639359 | Proser2 |
| USG000000 | 7,6331507 | 1,081047089 | 0,490726 | 3,043817 | 0,002335974 | 0,048995279 | Polr2k |
| USG000000 | 25,55155 | 1,791340947 | 0,237138 | 7,795445 | 6,41819E-15 | 2,33808E-12 | Lmod1 |
| USG000000 | 17,328439 | -1,219210294 | 0,46568 | -3,282572 | 0,001028646 | 0,027071394 | Syt12 |
| USG000000 | 1,6580505 | 2,447255353 | 1,079156 | 3,13086 | 0,001742954 | 0,039357901 | Cxx1c |
| USG000000 | 29,098023 | 1,220748193 | 0,234617 | 5,563566 | 2,64318E-08 | 3,73362E-06 | Ier2 |

| | | | | | | | |
|---|---|---|---|---|---|---|---|
| USG000000 | 3,0072257 | 1,622114354 | 0,755574 | 3,057708 | 0,002230368 | 0,047499974 | 6330403A02Rik |
| USG000000 | 5,2039782 | 2,305237371 | 0,568311 | 4,542247 | 5,56577E-06 | 0,000391101 | Myh8 |
| USG000000 | 36,843456 | 1,374977505 | 0,228521 | 6,319265 | 2,62811E-10 | 5,35014E-08 | Ryr3 |
| USG000000 | 546894,17 | 1,002426607 | 0,141947 | 7,310542 | 2,66068E-13 | 7,67328E-11 | mt-Rnr1 |
| USG000000 | 705,35181 | 2,098112558 | 0,543912 | 4,36143 | 1,29215E-05 | 0,000833302 | mt-Tl1 |
| USG000000 | 433,16838 | 2,222829907 | 0,508564 | 4,815653 | 1,46719E-06 | 0,00012694 | mt-Tq |
| USG000000 | 1199,3962 | 2,574338053 | 0,602914 | 4,727589 | 2,27201E-06 | 0,000187211 | mt-Tw |
| USG000000 | 1000,0766 | 2,567354145 | 0,628098 | 4,559206 | 5,13474E-06 | 0,000364514 | mt-Ts1 |
| USG000000 | 293,32897 | 1,071731128 | 0,381241 | 3,452263 | 0,000555906 | 0,016987652 | mt-Td |
| USG000000 | 416,82383 | 1,859260797 | 0,529206 | 4,060771 | 4,8911E-05 | 0,00246209 | mt-Th |
| USG000000 | 135,01852 | 1,832122858 | 0,453807 | 4,513773 | 6,36844E-06 | 0,000440792 | mt-Te |
| USG000000 | 2,4659488 | 3,125918623 | 1,083043 | 3,547207 | 0,000389338 | 0,012711334 | Snord49b |
| USG000000 | 6,5283882 | 1,538876288 | 0,533617 | 3,55045 | 0,000384573 | 0,012585456 | Gm22980 |
| USG000000 | 3,6607321 | 3,992874111 | 0,830809 | 5,214573 | 1,84241E-07 | 2,14324E-05 | Gm22997 |
| USG000000 | 2,0436354 | 4,986725479 | 1,572237 | 3,81657 | 0,00013532 | 0,00570613 | Gm24920 |
| USG000000 | 3,0021983 | 1,925276418 | 0,836448 | 3,146391 | 0,001652989 | 0,037895488 | Gm23680 |
| USG000000 | 7,239988 | 1,670066198 | 0,584543 | 3,529637 | 0,00041613 | 0,013365391 | Gm22744 |
| USG000000 | 3,8238098 | 2,416971791 | 0,799226 | 3,671666 | 0,000240974 | 0,008871832 | Mir133a-2 |
| USG000000 | 6,5478273 | 1,755107048 | 0,726025 | 3,216203 | 0,001298989 | 0,031547209 | Mir24-2 |
| USG000000 | 98,193044 | 1,204674955 | 0,434506 | 3,434882 | 0,000592812 | 0,017878643 | Rny1 |
| USG000000 | 1,9473913 | 7,242960712 | 1,576911 | 4,588283 | 4,46907E-06 | 0,000323902 | C230004F18Rik |
| USG000000 | 224,26998 | 2,924790927 | 0,784505 | 4,254109 | 2,09883E-05 | 0,001241627 | AA474408 |
| USG000000 | 2,7646177 | 2,022797469 | 0,85009 | 3,217038 | 0,001295216 | 0,031520416 | Gm26002 |
| USG000000 | 630,0724 | 1,430723438 | 0,53789 | 3,369037 | 0,000754312 | 0,021158291 | Gm23935 |
| USG000000 | 36,886997 | 1,630272184 | 0,724264 | 3,109265 | 0,001875534 | 0,041492972 | Gm24270 |
| USG000000 | 2,7963497 | 2,565902949 | 0,733491 | 4,068125 | 4,7393E-05 | 0,002394385 | Gm24336 |
| USG000000 | 2,1172667 | 2,889577101 | 1,065214 | 3,420508 | 0,000625044 | 0,018687867 | Gm25506 |
| USG000000 | 4,322694 | 3,186128506 | 0,663479 | 5,217925 | 1,80939E-07 | 2,12265E-05 | Gm24372 |
| USG000000 | 17,287554 | 1,743435317 | 0,681205 | 3,315755 | 0,000913958 | 0,024614622 | Snord12 |
| USG000000 | 4,9594462 | 2,589040432 | 0,702992 | 4,224374 | 2,39606E-05 | 0,00137629 | Snord90 |
| USG000000 | 6,4461438 | 1,430071584 | 0,624341 | 3,116622 | 0,001829361 | 0,040844901 | Gm9531 |
| USG000000 | 1,6583485 | 6,908199648 | 1,743407 | 4,11256 | 3,91296E-05 | 0,002107669 | Olfr650-ps1 |
| USG000000 | 4,4259828 | 1,264930011 | 0,555262 | 3,111378 | 0,001862166 | 0,04137714 | AA415398 |
| USG000000 | 183,40133 | 2,93590232 | 1,136832 | 3,368767 | 0,000755053 | 0,021158291 | Gm24245 |
| USG000000 | 73,138798 | 2,309029994 | 0,647338 | 4,11575 | 3,85922E-05 | 0,002086842 | Gm15662 |
| USG000000 | 20,114469 | 1,216755859 | 0,265796 | 4,980587 | 6,33917E-07 | 6,26808E-05 | Apold1 |
| USG000000 | 12,624253 | 1,187549555 | 0,387079 | 3,680226 | 0,000233027 | 0,008759095 | Hspa1b |
| USG000000 | 14,090925 | 1,08310439 | 0,302786 | 4,09692 | 4,18684E-05 | 0,002195396 | Gm6611 |
| USG000000 | 2,0076613 | 3,131002766 | 1,080244 | 3,560981 | 0,000369471 | 0,012177599 | Gm22265 |
| USG000000 | 1,6078421 | 3,659669955 | 1,462053 | 3,241721 | 0,001188103 | 0,030122542 | Gm22129 |
| USG000000 | 2,4154423 | 2,003675236 | 0,842681 | 3,216951 | 0,001295609 | 0,031520416 | Gm22113 |
| USG000000 | 9,1975452 | -2,99066658 | 1,203655 | -3,219057 | 0,001286131 | 0,031455664 | Gm4673 |
| USG000000 | 941714,76 | 1,545921626 | 0,627927 | 3,227618 | 0,001248254 | 0,031190569 | Yam1 |
| USG000000 | 1728,6197 | 2,633980106 | 0,756497 | 4,047546 | 5,17573E-05 | 0,00257101 | Mir6236 |
| USG000000 | 2,6247261 | 2,324375648 | 0,752453 | 3,729013 | 0,000192231 | 0,00762068 | Gm7889 |
| USG000001 | 2,2289513 | 5,947411832 | 1,629267 | 3,888178 | 0,000101 | 0,004382876 | AC153941.1 |

**Supplementary table 6)** List of inflammatory genes that changed their expression (up/down reglated) after delivery of VEGF-A mRNA to the cardiac tissue via CPC SEC-EVs, HTB SEC-EVs, HUVEC SEC-EVs and LNPs.

| CPC SEC-EVs | | | | | | | | |
|---|---|---|---|---|---|---|---|---|
| **NA.** | **baseMean** | **log2FC** | **lfcSE** | **stat** | **pvalue** | **padj** | **gene_symbol** | **Up/down regulation** |
| ENSMUSG00000019122 | 16,512171 | -1,018199 | 0,432508 | -3,0958523 | 0,0019625 | 0,039406 | Ccl9 | Down reglated |
| ENSMUSG00000026580 | 19,719917 | -1,175982 | 0,426357 | -3,3962217 | 0,0006832 | 0,0183 | Selp | Down reglated |
| ENSMUSG00000029380 | 9,9774351 | -1,275666 | 0,562665 | -3,0382908 | 0,0023792 | 0,04441 | Cxcl1 | Down reglated |
| ENSMUSG00000044768 | 17,564048 | -1,578971 | 0,374195 | -4,663738 | 3,105E-06 | 0,000235 | D1Ertd622e | Down reglated |

| HTB SEC-EVs | | | | | | | | |
|---|---|---|---|---|---|---|---|---|
| **NA.** | **baseMean** | **log2FC** | **lfcSE** | **stat** | **pvalue** | **padj** | **gene_symbol** | |
| ENSMUSG00000001750 | 17,474582 | 1,3038399 | 0,310865 | 4,6369535 | 3,536E-06 | 0,00015 | Tcirg1 | up-reglated |

| HUVEC SEC-EVs | | | | | | | | |
|---|---|---|---|---|---|---|---|---|
| ENSMUSG00000021624 | 1,7550396 | 2,2152094 | 1,062178 | 3,0361688 | 0,0023961 | 0,049768 | Cd180 | up-reglated |
| ENSMUSG00000050440 | 10,025181 | 1,0697504 | 0,471444 | 3,0986266 | 0,0019442 | 0,042585 | Hamp | up-reglated |

| LNPs | | | | | | | | |
|---|---|---|---|---|---|---|---|---|
| **NA.** | **baseMean** | **log2FC** | **lfcSE** | **stat** | **pvalue** | **padj** | **gene_symbol** | |
| ENSMUSG00000030017 | 4,7331115 | 6,3155131 | 1,12494 | 5,9467853 | 2,735E-09 | 1,3E-07 | Reg3g | up-reglated |
| ENSMUSG00000025153 | 170,57794 | 3,5350715 | 0,154916 | 22,920138 | 2,93E-116 | 8,4E-113 | Fasn | up-reglated |
| ENSMUSG00000029372 | 4,6208733 | 3,1548542 | 0,533224 | 6,2475924 | 4,168E-10 | 2,2E-08 | Ppbp | up-reglated |
| ENSMUSG00000040809 | 4,9965079 | 2,937613 | 0,544208 | 5,7535231 | 8,74E-09 | 3,83E-07 | Chil3 | up-reglated |
| ENSMUSG00000056071 | 6,8508538 | 2,8022449 | 0,402312 | 7,2331609 | 4,719E-13 | 3,83E-11 | S100a9 | up-reglated |
| ENSMUSG00000018927 | 25,149748 | 2,5671581 | 0,23983 | 10,901835 | 1,13E-27 | 3,98E-25 | Ccl6 | up-reglated |
| ENSMUSG00000050440 | 10,025181 | 2,2972826 | 0,327779 | 7,2840185 | 3,24E-13 | 2,7E-11 | Hamp | up-reglated |
| ENSMUSG00000024164 | 263,94599 | 2,1985308 | 0,089883 | 24,51962 | 9,12E-133 | 3,3E-129 | C3 | up-reglated |
| ENSMUSG00000023078 | 3,0745 | 2,0563986 | 0,612716 | 3,9176538 | 8,942E-05 | 0,001455 | Cxcl13 | up-reglated |
| ENSMUSG00000056054 | 2,7369915 | 2,0244847 | 0,679946 | 3,6090247 | 0,0003074 | 0,004176 | S100a8 | up-reglated |
| ENSMUSG00000035385 | 11,728038 | 2,0059535 | 0,341342 | 6,2019553 | 5,577E-10 | 2,92E-08 | Ccl2 | up-reglated |
| ENSMUSG00000026580 | 19,719917 | 1,9152799 | 0,205713 | 9,5034812 | 2,03E-21 | 4,37E-19 | Selp | up-reglated |
| ENSMUSG00000019122 | 16,512171 | 1,6830914 | 0,223238 | 7,7874743 | 6,836E-15 | 7,47E-13 | Ccl9 | up-reglated |
| ENSMUSG00000033777 | 3,2606068 | 1,6504445 | 0,533042 | 3,6925714 | 0,000222 | 0,003178 | Tlr13 | up-reglated |
| ENSMUSG00000025804 | 6,7392224 | 1,61482 | 0,345389 | 5,0709679 | 3,958E-07 | 1,22E-05 | Ccr1 | up-reglated |
| ENSMUSG00000049103 | 12,746371 | 1,6115936 | 0,275321 | 6,1711804 | 6,778E-10 | 3,48E-08 | Ccr2 | up-reglated |
| ENSMUSG00000015340 | 7,0001858 | 1,6009181 | 0,343306 | 5,0601311 | 4,19E-07 | 1,28E-05 | Cybb | up-reglated |
| ENSMUSG00000040152 | 142,6716 | 1,471986 | 0,115934 | 12,844038 | 9,29E-38 | 5,59E-35 | Thbs1 | up-reglated |
| ENSMUSG00000030162 | 3,3592568 | 1,4507215 | 0,564228 | 3,2859713 | 0,0010163 | 0,011449 | Olr1 | up-reglated |
| ENSMUSG00000024659 | 22,931267 | 1,3747063 | 0,189478 | 7,5017988 | 6,295E-14 | 5,94E-12 | Anxa1 | up-reglated |
| ENSMUSG00000029373 | 10,664311 | 1,3713922 | 0,302732 | 4,9461902 | 7,568E-07 | 2,15E-05 | Pf4 | up-reglated |
| ENSMUSG00000032487 | 6,9587987 | 1,3180178 | 0,350345 | 4,2452031 | 2,184E-05 | 0,000421 | Ptgs2 | up-reglated |
| ENSMUSG00000035373 | 8,3215397 | 1,2739799 | 0,32227 | 4,4127152 | 1,021E-05 | 0,000219 | Ccl7 | up-reglated |
| ENSMUSG00000079227 | 10,577535 | 1,2718982 | 0,309688 | 4,5462942 | 5,46E-06 | 0,000125 | Ccr5 | up-reglated |
| ENSMUSG00000023913 | 7,6604282 | 1,2326208 | 0,337255 | 4,1482087 | 3,351E-05 | 0,000622 | Pla2g7 | up-reglated |
| ENSMUSG00000058427 | 3,7697414 | 1,2131164 | 0,452526 | 3,3397247 | 0,0008386 | 0,00976 | Cxcl2 | up-reglated |
| ENSMUSG00000026070 | 2,9228136 | 1,2127958 | 0,552591 | 3,0057471 | 0,0026493 | 0,024336 | Il18r1 | up-reglated |
| ENSMUSG00000026068 | 3,0508733 | 1,167607 | 0,51581 | 3,0276939 | 0,0024643 | 0,022973 | Il18rap | up-reglated |
| ENSMUSG00000026365 | 363,68259 | 1,1405657 | 0,129834 | 9,0199556 | 1,882E-19 | 3,44E-17 | Cfh | up-reglated |
| ENSMUSG00000028108 | 29,335616 | 1,0737964 | 0,222211 | 5,1895683 | 2,108E-07 | 6,77E-06 | Ecm1 | up-reglated |
| ENSMUSG00000055994 | 4,5186089 | -1,292424 | 0,656602 | -2,8045324 | 0,005039 | 0,040443 | Nod2 | Down reglated |
| ENSMUSG00000063415 | 9,0300582 | -1,627371 | 0,459562 | -4,0531292 | 5,054E-05 | 0,000888 | Cyp26b1 | Down reglated |

**Supplementary table 7)** List of Canonical Pathways mediated by VEGF-A mRNA to cardiac tissue via CPC SEC-EVs, HTB SEC-EVs, HUVEC SEC-EVs and LNPs.

| | (-log(P-value)) | | | | z-score | | | |
|---|---|---|---|---|---|---|---|---|
| **Canonical Pathways:** | **CPC SEC-EVs** | **HTB SEC-EVs** | **HUVEC SEC-EVs** | **LNPs** | **CPC SEC-EVs** | **HTB SEC-EVs** | **HUVEC SEC-EVs** | **LNPs** |
| Cardiomyocyte Differentiation via BMP Receptors (Bone morphogenetic protein receptor) | 1,353839234 | 0 | 0 | 1,27622688 | N/A | N/A | N/A | N/A |
| Apelin Cardiomyocyte Signaling Pathway | 0,751134418 | 0,286508027 | 0 | 1,11167269 | N/A | N/A | N/A | 2 |
| Cardiac β-adrenergic Signaling | 1,311119948 | 0 | 0 | 0,77304954 | N/A | N/A | N/A | 1,342 |
| **Factors Promoting Cardiogenesis in Vertebrates** | **1,443492586** | **0** | **0** | **0** | N/A | N/A | N/A | N/A |
| Nitric Oxide Signaling in the Cardiovascular System | 1,632449307 | 0,231911837 | 0,39838973 | 3,11614561 | N/A | N/A | N/A | 2,121 |
| Cardiac Hypertrophy Signaling (Enhanced) | 1,043789891 | 0 | 0,394411459 | 4,53065122 | N/A | N/A | N/A | 3,5 |

**Supplementary table 8)** Top10 enriched GO Biological Processes 2021 associated with the list of DEGs (differentially expressed genes), and top 10 enriched MGI Mammalian Phenotype Level 4 2021 associated with the list of DEGs.

| **Top10 enriched GO Biological Processes 2021 associated with the list of DEGs (differentially expressed genes)** | | |
|---|---|---|
| **CPC SEC-EVs** | | |
| **Index** | **Name** | **P-value** |
| 1 | **regulated exocytosis (GO:0045055)** | 7,44E-07 |
| 2 | **platelet degranulation (GO:0002576)** | 0,00001255 |
| 3 | **cardiac muscle cell development (GO:0055013)** | 0,004322 |
| 4 | in utero embryonic development (GO:0001701) | 0,004686 |
| 5 | synaptic vesicle cycle (GO:0099504) | 0,006705 |
| 6 | proton transmembrane transport (GO:1902600) | 0,01008 |
| 7 | lysosome localization (GO:0032418) | 0,01008 |
| 8 | fatty acid derivative biosynthetic process (GO:1901570) | 0,01171 |
| 9 | synaptic vesicle exocytosis (GO:0016079) | 0,01171 |
| 10 | signal release from synapse (GO:0099643) | 0,01345 |
| **HTB SEC-EVs** | | |
| **Index** | **Name** | **P-value** |
| 1 | **endothelial cell chemotaxis (GO:0035767)** | 0,0008209 |
| 2 | **positive regulation of glucose transmembrane transport (GO:0010828)** | 0,005226 |
| 3 | **cell migration involved in sprouting angiogenesis (GO:0002042)** | 0,006876 |
| 4 | equilibrioception (GO:0050957) | 0,006922 |
| 5 | regulation of type B pancreatic cell proliferation (GO:0061469) | 0,006922 |
| 6 | blood vessel endothelial cell migration (GO:0043534) | 0,01375 |
| 7 | positive regulation of fatty acid oxidation (GO:0046321) | 0,0279 |
| 8 | negative regulation of hydrogen peroxide-induced cell death (GO:1903206) | 0,03249 |
| 9 | positive regulation of histone deacetylation (GO:0031065) | 0,03249 |
| 10 | neuromuscular process controlling balance (GO:0050885) | 0,03736 |
| **HUVEC SEC-EVs** | | |
| **Index** | **Name** | **P-value** |
| 1 | **regulation of biosynthetic process (GO:0009889)** | 0,000259 |
| 2 | **regulated exocytosis (GO:0045055)** | 0,0003251 |
| 3 | **fibrinolysis (GO:0042730)** | 0,0006895 |
| 4 | platelet degranulation (GO:0002576) | 0,0007696 |
| 5 | muscle contraction (GO:0006936) | 0,0009266 |
| 6 | regulation of type B pancreatic cell proliferation (GO:0061469) | 0,002067 |
| 7 | induction of bacterial agglutination (GO:0043152) | 0,002067 |
| 8 | negative regulation of low-density lipoprotein receptor activity (GO:1905598) | 0,00287 |
| 9 | regulation of heterotypic cell-cell adhesion (GO:0034114) | 0,003191 |
| 10 | negative regulation of transport (GO:0051051) | 0,003201 |
| **LNPs** | | |
| **Index** | **Name** | **P-value** |
| 1 | **neutrophil degranulation (GO:0043312)** | 1,85E-16 |
| 2 | **neutrophil activation involved in immune response (GO:0002283)** | 2,53E-16 |

| | | |
|---|---|---|
| 3 | **neutrophil mediated immunity (GO:0002446)** | 3,19E-16 |
| 4 | extracellular structure organization (GO:0043062) | 5,18E-15 |
| 5 | external encapsulating structure organization (GO:0045229) | 5,86E-15 |
| 6 | extracellular matrix organization (GO:0030198) | 1,41E-13 |
| 7 | cytokine-mediated signaling pathway (GO:0019221) | 3,33E-10 |
| 8 | granulocyte chemotaxis (GO:0071621) | 5,25E-10 |
| 9 | platelet degranulation (GO:0002576) | 1,99E-09 |
| 10 | positive regulation of metabolic process (GO:0009893) | 3,28E-09 |

| **Top 10 enriched MGI Mammalian Phenotype Level 4 2021 associated with the list of DEGs** | | |
|---|---|---|
| **CPC SEC-EVs** | | |
| **Index** | **Name** | **P-value** |
| 1 | **subcutaneous hemorrhage MP:0031152** | 0,0003441 |
| 2 | **decreased CD8-positive, naive alpha-beta T cell number MP:0013435** | 0,0007283 |
| 3 | **skin hemorrhage MP:0011514** | 0,001067 |
| 4 | abnormal blood coagulation MP:0002551 | 0,0012 |
| 5 | abnormal visceral yolk sac blood island morphology MP:0011204 | 0,00217 |
| 6 | increased bleeding time MP:0005606 | 0,00221 |
| 7 | eye inflammation MP:0001851 | 0,002714 |
| 8 | hemorrhage MP:0001914 | 0,004047 |
| 9 | myeloid hyperplasia MP:0010373 | 0,007149 |
| 10 | increased fat cell size MP:0006094 | 0,009051 |
| **HTB SEC-EVs** | | |
| **Index** | **Name** | **P-value** |
| 1 | **abnormal cochlear hair cell inter-stereocilial links morphology MP:0004577** | 0,006922 |
| 2 | **decreased granulocyte monocyte progenitor cell number MP:0013697** | 0,006922 |
| 3 | **delayed chorioallantoic fusion MP:0009656** | 0,006922 |
| 4 | decreased hippocampus pyramidal cell number MP:0009971 | 0,009549 |
| 5 | abnormal hair cell mechanoelectric transduction MP:0004431 | 0,0159 |
| 6 | abnormal hippocampus pyramidal cell layer MP:0008284 | 0,02129 |
| 7 | abnormal defecation MP:0003866 | 0,0279 |
| 8 | abnormal orientation of inner hair cell stereociliary bundles MP:0004492 | 0,0279 |
| 9 | abnormal orientation of cochlear hair cell stereociliary bundles MP:0004522 | 0,03736 |
| 10 | absent teeth MP:0030610 | 0,03736 |
| **HUVEC SEC-EVs** | | |
| **Index** | **Name** | **P-value** |
| 1 | **uterine hemorrhage MP:0004898** | 0,002067 |
| 2 | **renal necrosis MP:0003946** | 0,00287 |
| 3 | **abnormal circulating alanine transaminase level MP:0001573** | 0,00287 |
| 4 | abnormal vascular wound healing MP:0004883 | 0,003681 |
| 5 | skin hemorrhage MP:0011514 | 0,008673 |
| 6 | abnormal uterine environment MP:0004014 | 0,008673 |
| 7 | abnormal energy homeostasis MP:0005448 | 0,01017 |
| 8 | decreased left ventricle systolic pressure MP:0003822 | 0,01017 |
| 9 | focal hepatic necrosis MP:0001656 | 0,01017 |
| 10 | anisopoikilocytosis MP:0002641 | 0,01718 |

| LNPs | | |
|---|---|---|
| **Index** | **Name** | **P-value** |
| 1 | **abnormal leukocyte adhesion MP:0003628** | 2,34E-11 |
| 2 | **increased susceptibility to bacterial infection MP:0002412** | 4,34E-11 |
| 3 | **abnormal neutrophil physiology MP:0002463** | 1,60E-09 |
| 4 | abnormal leukocyte physiology MP:0002442 | 3,21E-09 |
| 5 | abnormal cellular extravasation MP:0009858 | 3,46E-09 |
| 6 | abnormal leukocyte migration MP:0003156 | 5,22E-09 |
| 7 | impaired neutrophil recruitment MP:0008719 | 2,46E-07 |
| 8 | increased eosinophil cell number MP:0005011 | 8,72E-07 |
| 9 | delayed wound healing MP:0002908 | 9,90E-07 |
| 10 | impaired macrophage chemotaxis MP:0003799 | 0,000001079 |

**Supplementary table 9)** List of IPA top 3 Networks caused by VEGF-A mRNA to the cardiac tissue via CPC-EVs, HTB-EVs, and LNPs.

**Caused by CPC-EVs VEGF-A mRNA delivery**

| ID | Molecules in Network | Score | Focus Molecules | Top Diseases and Functions | |
|---|---|---|---|---|---|
| 1 | AHSG,Akt,ALB,ALOX12,BMP10,Collagen(s),Fcer1,FGA,Fibrin,Fibrinogen,FLCN,GPIIB-IIIA,Growth hormone,HBA1/HBA2,HDL,HDL-cholesterol,hemoglobin,IL1,IL12 (complex),Immunoglobulin,Integrin,ITGA2B,LDL,MSLN,MT-ATP6,MT-ND5,PLAG1,Pro-inflammatory Cytokine,SELP,Stat3-Stat3,STAT5a/b,TSPAN4,TUBB1,UCP3,VEGFA | 39 | 17 | [Cardiovascular Disease, Cancer, Organismal Injury and Abnormalities] | 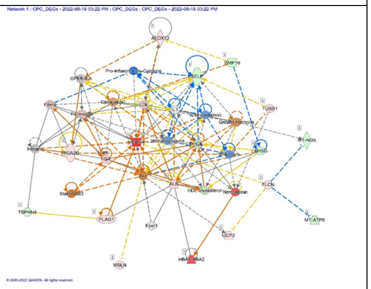 |
| 2 | AMPK,Ap1,BACH2,Ccl9,CD3,chemokine,Collagen Alpha1,collagen type i (family),CXCL2,EHHADH,ERK,ERK1/2,HMGCS2,Ige,IgG,IL12 (family),Interferon alpha,LCN2,LGALS4,Lh,Mapk,Mek,MMP19,MTORC1,P38 MAPK,PDE5A,PI3K (complex),PPP1R3C,Rac,RNA polymerase II,RORC,SLA,TCR,Tgf beta,Vegf | 25 | 12 | [Cellular Movement, Hematological System Development and Function, Immune Cell Trafficking] | 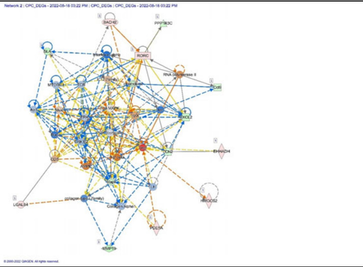 |
| 3 | ADAMTS9,ANXA8/ANXA8L1,DTX1,EGFR,EIF2AK2,ELAVL1,epoprostenol,ERAP1,FIGN,GATA3,Gvin1 (includes others),Hbb-b2,HSD17B1,IFNG,IL12B,IL18BP,IRF2BP2,KHNYN,LAG3,MACIR,Mir155hg,NOX1,PIM1,PNPLA7,PTEN,PTPRB,RAMP1,SH2D2A,SMTN,Snord57,Snord89,TEK,TMA16,TNFRSF1A,TRPM2 | 18 | 9 | [Cellular Development, Cellular Growth and Proliferation, Hematological System Development and Function] | 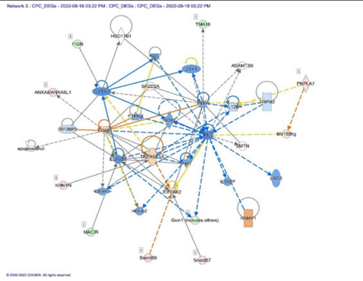 |

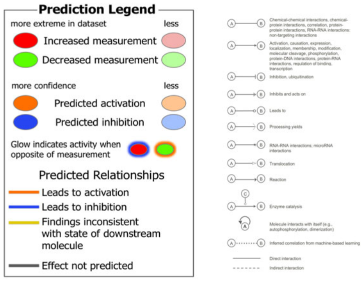

**Caused by HTB-EVs VEGF-A mRNA delivery**

| ID | Molecules in Network | Score | Focus Molecules | Top Diseases and Functions | |
|---|---|---|---|---|---|
| 1 | 40S Ribosome-eIF3-mRNA-eIF4A-eIF4B-eIF4E-eIF4G,43S Translation Preinitiation,48s,ATP1B3,ELK4,H2-Q5,IFI16,Ifn,IFN alpha/beta,IFN Beta,Ifnar,IFT46,IL12 (complex),ISG15,JAK,KIFC1,KPNA4,MT-RNR2,MT-TK,MT-TL2,MT-TM,MT-TP,MT-TT,MT-TY,Mx2,NFkB (complex),NUBP1,PIAS4,Ribosomal 40s subunit,RMRP,RNA polymerase II,Rnr,SARS2,STAT,THAP12 | 38 | 21 | [Hereditary Disorder, Metabolic Disease, Neurological Disease] | 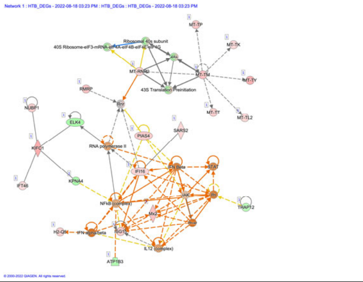 |
| 2 | ALAS2,ALB,Alpha tubulin,ANGPTL7,BET,BETA TUBULIN,BRDT,Cbp/p300,CDH23,chymotrypsin,CROCC,ERK1/2,Fcer1,Ferritin,Growth hormone,HBA1/HBA2,HBB,HDL,hemoglobin,HP,MIIP,NDUFC1,Nr1h,PFAS,Pro-inflammatory Cytokine,RASD1,Rsk,RSPH3,SLC4A1,STAT5a/b,STXBP2,TRAF3IP3,TRPM2,TUBB1,USH1C | 36 | 20 | [Cardiovascular Disease, Cell Death and Survival, Connective Tissue Disorders] | 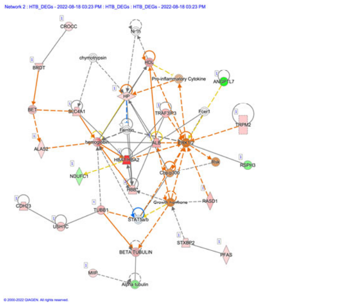 |

| | | | | | |
|---|---|---|---|---|---|
| 3 | Actin,B-cell receptor,Cofilin,Collagen Alpha1,DGKZ,EGR3,ERK,GOT,GRK6,IFNGR2,IgG,IgG1,IgG2a,IGHM,IGKC,Igm,Immunoglobulin,Ldh (complex),LDL,Mlc,MYL4,NEB,Notch,PI3K (family),Rap1,RNU11,RNU12,Snord17,snRNP,snRNP-IgG Immune complex,SNRNP48,SNRNP70,STRAP,SYNGAP1,TCIRG1 | 27 | 16 | [Dermatological Diseases and Conditions, Immunological Disease, Inflammatory Disease] | 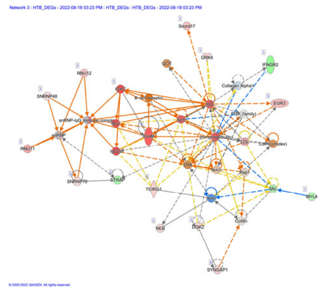 |

| Caused by LNPs VEGF-A mRNA delivery | | | | | |
|---|---|---|---|---|---|
| ID | Molecules in Network | Score | Focus Molecules | Top Diseases and Functions | |
| 1 | ADCY3,Adducin,Akt,Atrial Natriuretic Peptide,BANK1,BPIFA1,BPIFB1,CIDEC,Collagen type VI,cyclooxygenase,EBF2,HP,IP6K3,LTF,LYZ,MARCO,Muc4,MUC5B,Mucin,N-Cadherin,NDNF,NRTN,PIGR,PTGDS,Rpl23a,SCAVENGER receptor CLASS A,SCGB3A1,SCGB3A2,SEMA3D,SIRPB1,STEAP4,T3-TR-RXR,TMEM176A,TMEM176B,WFDC2 | 43 | 26 | [Cell-To-Cell Signaling and Interaction, Cellular Development, Cellular Growth and Proliferation] | 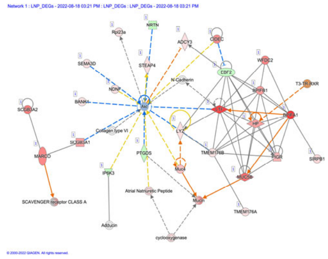 |
| 2 | ALAS2,Apol11b (includes others),BNC1,CD300LF,Cpla2,Ctla2a/Ctla2b,DMKN,DNA-methyltransferase,EGLN,EMP1,GPD1,GPD2,Hamp/Hamp2,HBA1/HBA2,HBB,HDL,hemoglobin,Il8r,Integrin α5-β,INTERLEUKIN,ITGB3,mir-486,PAM,PF4,PLVAP,PYGL,RGS10,RNF122,SELENOF,SLC39A8,SLC4A1,Srgn,STX11,UPK3B,Vegf | 43 | 26 | [Cardiovascular Disease, Hematological Disease, Organismal Injury and Abnormalities] | 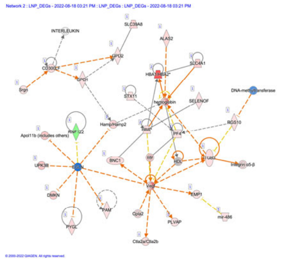 |
| 3 | 60S ribosomal subunit,ACTA2,Adaptor protein 1,Alpha catenin,C3,Ccl6,Ccl7,Ccl9,CDH1,CG,chemokine,COL11A2,Collagen type I (complex),CRIP1,CXCL3,GM2A,ID2,Ifitm1,IL33,Mir122a,b,MMP9,Notch,Ppbp,Relaxin,RPL22,Rpl36a,RPL39,S100A6,S100A8,S100A9,SHTN1,SLCO2A1,TIMP1,VCAN,Wnt | 41 | 25 | [Cellular Movement, Hematological System Development and Function, Immune Cell Trafficking] | 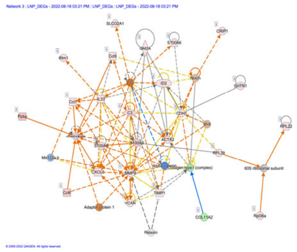 |
